# A Highly Selective, Cell-Permeable Fluorescent Probe for Imaging Histone Deacetylase 6 in Live Cells

**DOI:** 10.64898/2025.12.09.692542

**Authors:** Văn Thắng Nguyễn, Tanja Koenen, Jonas Bucevičius, Gražvydas Lukinavičius

**Author notes:** **Corresponding Author** Gražvydas Lukinavičius – Chromatin Labeling and Imaging Group, Department of NanoBiophotonics, Max Planck Institute for Multidisciplinary Sciences, Am Fassberg 11, 37077 Göttingen, Germany.

## Abstract

Histone Deacetylase 6 (HDAC6) plays a crucial role in diverse cellular processes, including cytoskeletal regulation, protein quality control, and stress responses; and its dysregulation is linked to multiple cancers and neurodegenerative disorders, making it a key therapeutic target. However, a detailed understanding of its dynamic functions has been limited by the lack of chemical tools for its visualization in living systems. Utilizing the highly selective inhibitor Nexturastat A as a targeting scaffold, we developed **6SiR-C3-NextA** probe incorporating a bright, photostable, far-red silicon-rhodamine fluorophore. Biochemical and cellular assays show that **6SiR-C3-NextA** binds to HDAC6 with high affinity (K_d_^app^ = 21 ± 4 nM) and selectivity over other HDAC enzymes, as validated using a developed panel of engineered cell lines expressing individual human HDACs. The probe is cell-permeable, exhibits low cytotoxicity, compatible with super-resolution technique and enables the visualization of endogenous HDAC6 across multiple cell lines. We demonstrate its utility by performing imaging of HDAC6’s association with the microtubule network and its dynamic recruitment to stress granules in living cells.

## INTRODUCTION

Histone deacetylases (HDACs) are a family of enzymes that catalyze the removal of acetyl groups from lysine residues on histone and non-histone protein substrates, playing a central role in regulating gene expression and cellular function^1^. Among the 18 human HDACs, HDAC6 stands out due to its distinct localization and domain architecture, which includes two different catalytic domains and a C-terminal zinc finger ubiquitin-binding domain. This particular structure enables HDAC6 to regulate multiple critical cellular pathways, including cytoskeleton dynamics, protein quality control, and stress responses^2^.

Given its central role in maintaining cellular homeostasis, the dysregulation of HDAC6 is linked to multiple human diseases. In oncology, HDAC6 promotes cellular transformation and enhances tumor cell proliferation^3^. In the central nervous system, its activity is critical for axonal transport and synaptic plasticity, and its dysfunction is implicated in neurodegenerative diseases like Alzheimer’s and Parkinson’s disease^4^. Consequently, the development of selective HDAC6 inhibitors has been an area of intense research, with several compounds entering clinical trials^5^.

Despite this therapeutic interest, studying the dynamic spatial and temporal regulation of HDAC6 in its native cellular environment remains a significant challenge. Although genetic tagging with fluorescent proteins (FPs) has provided valuable insights, this approach can introduce artifacts. The large size of FPs can lead to steric hindrance and functional perturbations, and the overexpression typically required for visualization can cause the fusion protein to mislocalize, failing to represent the true distribution of the native protein^6^. Small-molecule fluorescent probes offer a compelling alternative, providing minimal perturbation and superior photophysical properties. Several fluorescent probes for HDAC6 have been developed, typically by conjugating a known inhibitor to a fluorophore. However, existing probes suffer from significant drawbacks that limit their utility. A primary concern for many reported probes is the challenge of achieving and comprehensively demonstrating high selectivity, particularly over all other HDACs; this can lead to ambiguous biological data if off-target binding occurs^7, 8^. In addition, many naphthalimide and early Scriptaid analogues emit at shorter wavelengths, which suffer from cellular autofluorescence, limited tissue penetration for in vivo work, and potential phototoxicity^7^. While activatable probes such as HDAC-MB and CyAc-RGD aim to address these issues by generating fluorescence upon interaction or enzymatic processing, their slow kinetics limits the ability to track the highly dynamic processes involving HDAC6^9^.

To address this gap, we aimed to develop a fluorescent probe for HDAC6 suitable for live-cell imaging and advanced microscopy applications. We specified a probe with high affinity and selectivity, far-red emission to minimize autofluorescence and phototoxicity, excellent photostability, and high cell permeability. Herein, we describe the successful development of such a probe, **6SiR-C3-NextA**, through a systematic design, synthesis, and imaging-based assay. We demonstrate its utility in visualizing the dynamics of endogenous HDAC6 in different cell lines, including its association with the microtubule network and its recruitment to stress granules during the cellular stress response.

## RESULTS AND DISCUSSION

### Principle of Imaging-based assay

A critical challenge in probe development is validating selectivity within the complex environment of living cells. Conventional approaches reported for HDAC6 probes have largely relied on *in-vitro* binding assays using purified HDAC proteins. While informative, such methods cannot account for cell permeability, probe stability, or potential off-target interactions with the thousands of other components inside cell^10^. To overcome these limitations and rigorously assess probe performance in a more physiologically relevant context, we developed a robust, cell-based co-localization approach. A panel of U-2 OS cell lines was engineered to inducibly express individual HDACs (HDAC1-8) as fusions with HaloTag at N-terminus. This selection was based on the classification of human HDACs; the panel includes the most well-characterized members of the classical, zinc-dependent Class I (HDAC1, 2, 3, 8) and Class II (HDAC4, 5, 6, 7) enzymes, which represent the most likely off-targets for a probe based on a hydroxamic acid inhibitor. The mechanistically and structurally distinct Class III HDACs (sirtuins) were excluded as they are NAD□-dependent and lack the catalytic zinc ion targeted by the class I and II inhibitors^11^. The cell lines were constructed using a single-plasmid system containing an Epstein-Barr virus origin of replication for stable episomal maintenance. The plasmid also includes a cytomegalovirus-tetracycline operator (CMV-TetO□) promoter, which allows for tight, doxycycline-inducible control of gene expression^12^. The coding sequence of each human HDAC was inserted into this vector upstream of a C-terminal HaloTag using Gateway cloning, with the genes of interest PCR-amplified as listed in **Table S1**. All final constructs were confirmed by sequencing. In this system, the HaloTag is covalently labeled with a fluorescent ligand, providing a ground-truth reference for the location of each HDAC. The selectivity of a candidate probe can then be quantified by measuring the Pearson’s Correlation Coefficient (PCC) between its signal and the HaloTag reference signal (**Figure 1a**). This approach provides a direct, quantitative measure of on-target engagement while simultaneously considering any significant off-target binding within the cellular environment. Furthermore, this versatile assay is not limited to HDAC6 and can be readily adapted for the discovery and validation of selective fluorescent probes targeting other HDAC enzymes.

**Figure 1.**
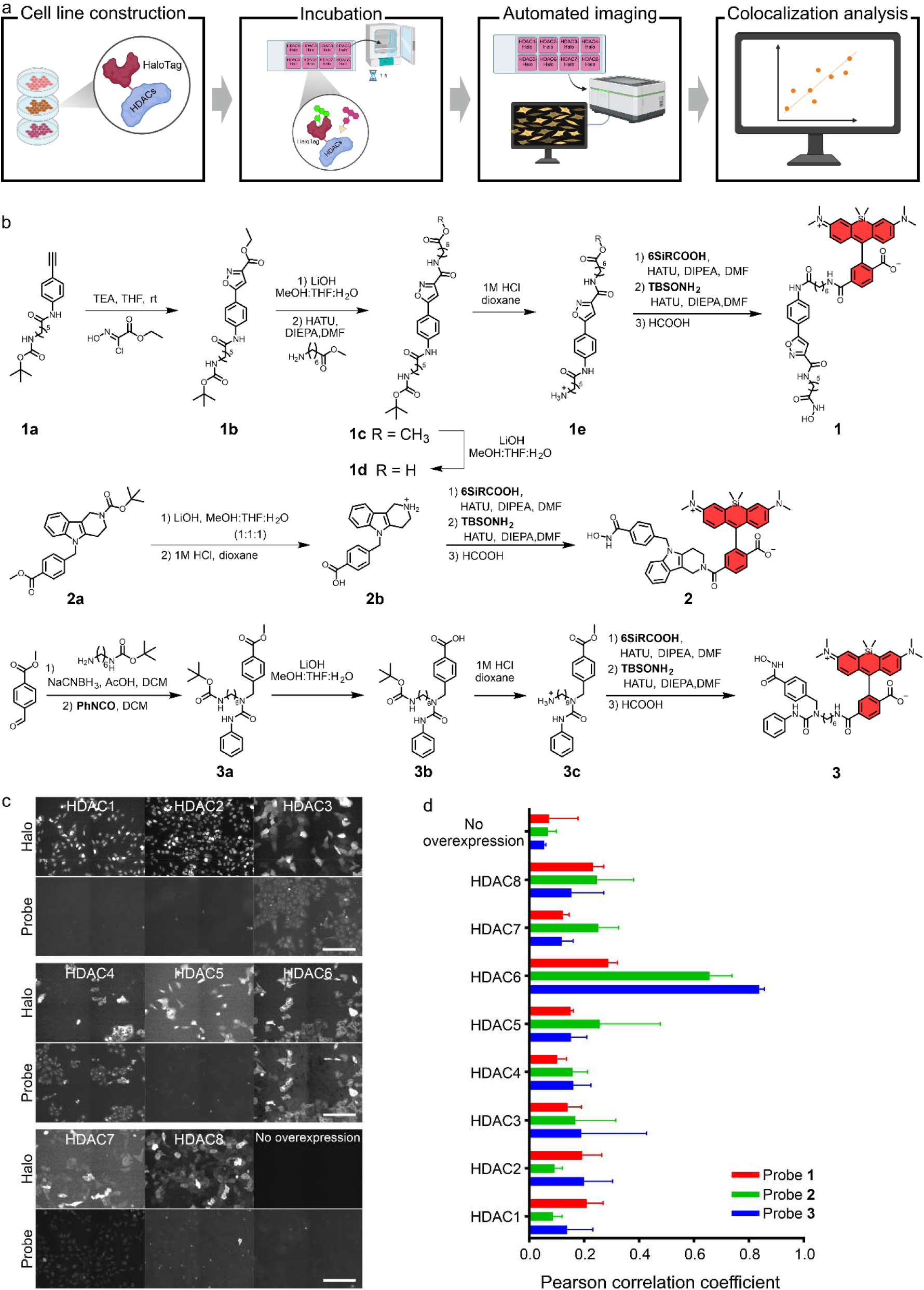
Imaging-based assay for HDAC6 Probe selection and optimization. (a) Schematic of the assay. U-2 OS cell lines engineered to express individual HDAC-HaloTag fusions are incubated with a HaloTag ligand and the candidate probe, followed by automated widefield imaging and co-localization analysis. (b) Synthetic schemes for the SiR-conjugated probes based on (1) CAY10603, (2) Tubastatin, and (3) Nexturastat A. (c) Representative fluorescence images from the assay of probe 1. Scale bar: 200 µm. (d) Quantitative co-localization analysis for probes 1, 2, and 3. Pearson’s Correlation Coefficient (PCC) between the probe and HaloTag signals. Data are presented as mean ± SD from three independent experiments. Figure 1a was created in Biorender.

#### Probe design and Structure-Activity Relationship (SAR) study

Our strategy was based on a modular probe design tethering a recognition moiety to a fluorophore via a chemical linker. For the recognition moiety, we selected three well-characterized inhibitors known for their high selectivity for HDAC6: Nexturastat A (IC_50_ 5.02 ± 0.06 nM), CAY10603 (IC_50_ 0.002 nM), and Tubastatin A (IC_50_ 15□± □1.0 nM)^13^. The primary challenge was to ensure that this selectivity was maintained after conjugation to a bulky fluorophore.

For our probe synthesis, we chose rhodamine class as the fluorophore. Rhodamine dyes are exceptionally well-suited for demanding live-cell imaging applications and compatible with super resolution techniques due to their combination of high brightness, excellent photostability. A key feature contributing to their utility is their excellent cell permeability, which is governed by reversible equilibrium between a fluorescent, zwitterionic form and a non-fluorescent, neutral spirolactone form. The charge-neutral spirolactone readily crosses the cell membrane, after which the intracellular environment shifts the equilibrium back towards the fluorescent zwitterionic state, leading to “turn-on” of the signal^14^. Thus, we synthesized first-generation probes, **1, 2** and **3**, by conjugating these HDAC6 selective inhibitor to a cell-permeable, far-red SiR (silicon rhodamine) fluophore (**Figure 1b**)^15^. Following the imaging-based assay, we found that there are significant differences in HDAC6 selectivity among the probes. The CAY10603-based probe, **1**, lost its desired binding profile as shown by low correlation value toward HDAC6 cell staining (PCC = 0.29 ± 0.03). The Tubastatin A-based probe **2** maintained its original selectivity but suffered from off-target binding (PCC = 0.66 ± 0.08) (**Figure 1c**). In contrast, the Nexturastat A-based probe **3** demonstrated exceptional selectivity, yielding a high PCC of0.84 ± 0.02 (**Figure 1c, 1d**). This finding validates our design strategy by confirming that the Nexturastat A scaffold maintains its high selectivity when conjugated to a large fluorophore.

With the selectivity of the conjugated Nexturastat A scaffold confirmed, we performed a systematic SAR study to optimize the probe’s performance by modifying the fluorophore, linker, and binding moiety (**Figure 2**).

**Figure 2.**
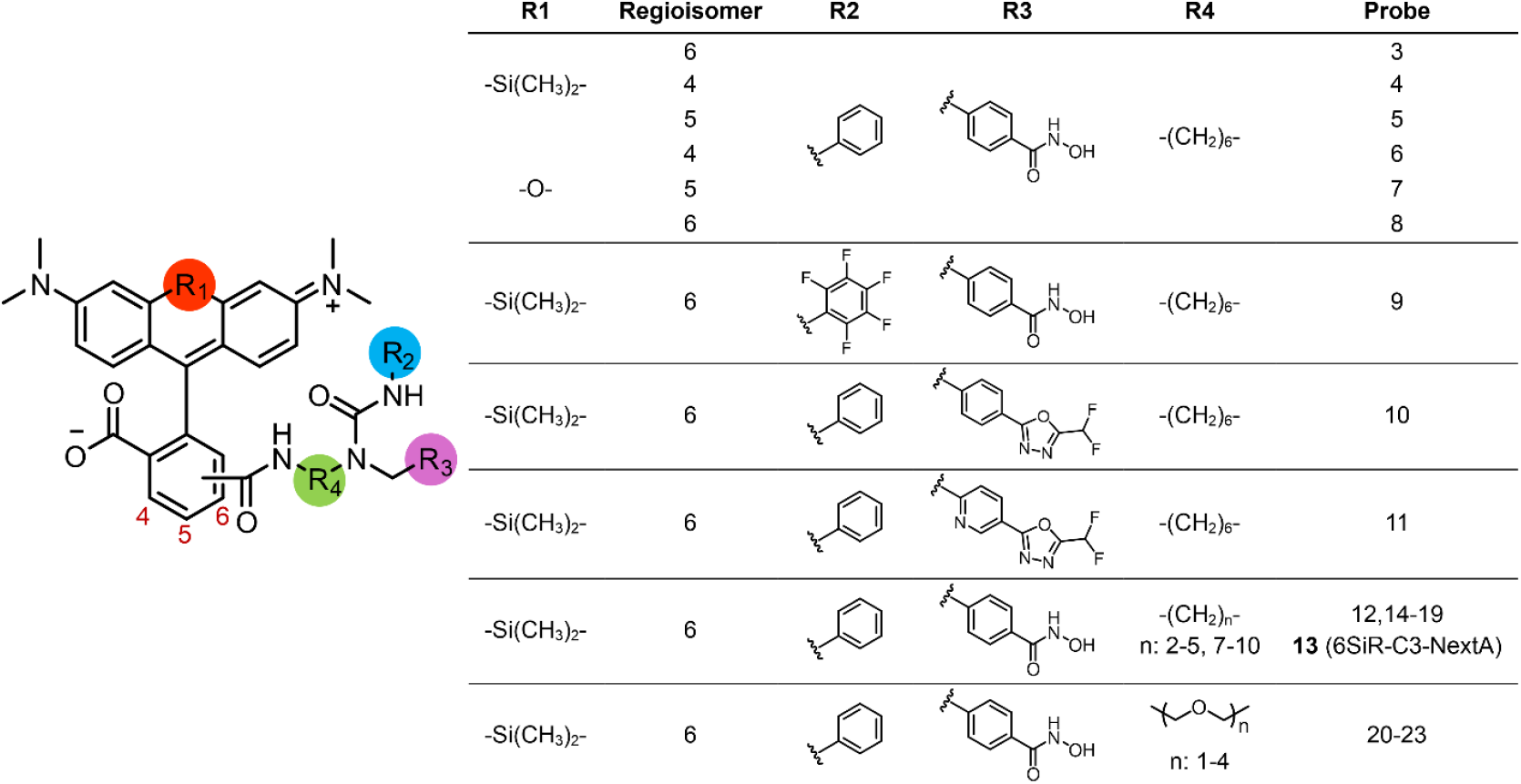
HDAC6 fluorescent probes used for the systematic structure-activity relationship (SAR) study.

#### Fluorophore

The choice of fluorophore is critical as its physicochemical properties can strongly influence the overall performance of the probe. We initiated our optimization by comparing our lead SiR-based probe with analogues constructed from tetramethylrhodamine (TMR). This comparison was designed to assess the impact of the fluorophore’s core structure and polarity on cellular performance. SiR dyes are generally more hydrophobic compared to the more polar TMR, a difference illustrated by their calculated partition coefficients of their zwitterion form (*c*Log*P* of ∼-0.88 for SiR vs. ∼-3.44 for TMR) and spiro-lactone form (*c*Log*P* of ∼5.86 for SiR vs. ∼4.24 for TMR), properties which often correlate with probe solubility and cell permeability. Furthermore, we synthesized and evaluated SiR’s isomers-4/5/6 (probes **3, 4** and **5)** and different constitutional TMR’s isomers-4/5/6 (probes **6, 7** and **8**), varying the attachment point of the linker to the fluorophore’s xanthene core (**Figure 2**). The specific conjugation site is known to be a critical determinant of probe performance, as it can alter the electronic structure of the fluorophore, thereby affecting its brightness and quantum yield, and can also introduce steric hindrance that impacts binding affinity or leads to non-specific interactions^16^. Following the imaging-based assessment, The SiR-based probes **3, 4** and **5** all showed strong co-localization with the HDAC6-Halo reference signal, as indicated by their high PCC values. In contrast, the TMR-based probes **6, 7** and **8** exhibited a significant loss of binding affinity, with dramatically lower PCCs (**Figure S1b**). Among the SiR probes, probe **3** was identified as the lead candidate due to its superior signal intensity, confirming 6SiR as the fluorophore of choice for this application (**Figure S1c**). Its far-red emission profile is particularly advantageous for live-cell imaging as it minimizes interference from cellular autofluorescence and reduces the laser power, thereby lowering phototoxicity^17^.

#### Scaffold Fluorination

To investigate if modulating the inhibitor’s physicochemical properties could improve imaging performance, we synthesized and tested an analogue in which the phenyl group of the Nexturastat A core was perfluorinated^18^. Contrary to the hypothesis that this might improve cell permeability, fluorinated probe **9** proved detrimental, leading to a significant decrease in both the signal intensity and the signal-to-background ratio (**Figure S2**). This suggests that fluorination negatively impacts either the probe’s affinity for HDAC6 or its cellular accumulation, confirming the non-fluorinated scaffold as superior.

#### Zinc binding group (ZBG)

We investigated alternative ZBGs, such as difluoromethyloxadiazole (DFMO), which have been reported as irrevesible HDAC6 binder with unprecedented isotype selectivity^19^. However, probes **10** and **11** incorporating DFMO failed to show better binding affinity in our imaging-based assay as shown by lower PCC values. This result confirms that the classical ZBG hydroxamic acid of Nexturastat A is essential for potent target engagement within our probe design (**Figure S3**).

#### Linker

The linker length and composition were systematically varied. A series of probes **12** - **23** with alkyl chains of C2 to C10 and polyethylene glycol (PEG) linkers of 1-4 units were synthesized and tested. This analysis revealed a optimal probe, with the C3 alkyl linker **13** (**6SiR-C3-NextA**) providing the highest signal intensity and best signal-to-background ratio (**Figure S4**). This suggests an optimal distance and flexibility is required to allow the SiR fluorophore to sit outside the binding pocket without disrupting the inhibitor’s key interactions. Performance decreased with both shorter (C2) and longer (C4-C10) alkyl chains, as well as with all PEG-based linkers. In addition, the optimal probe **6SiR-C3-NextA** still maintained the desired selectivity profile from the parent probe **3** (**Figure S5**).

This systematic imaging-based optimization revealed **6SiR-C3-NextA** as the lead candidate, possessing the optimal combination of a SiR fluorophore, a C3 alkyl linker, and the Nexturastat A hydroxamic acid ZBG.

#### Biochemical and Cellular Characterization of 6SiR-C3-NextA

To quantify the binding affinity of **6SiR-C3-NextA**, we performed a Homogeneous Time-Resolved Fluorescence (HTRF) saturation binding assay. This assay, which relies on Förster Resonance Energy Transfer (FRET) between a long-lifetime terbium donor on the protein and the probe’s SiR acceptor, provides a highly sensitive and robust measure of binding^20^. The assay measured a high-affinity interaction between the probe and HDAC6, yielding an apparent dissociation constant (K_d_ ^app^) of 21 ± 4 nM. In contrast, no significant binding was observed for HDAC1 or HDAC7, which represent Class I and Class IIa isoforms respectively, confirming the probe’s high selectivity (**Figure 3a**).

**Figure 3.**
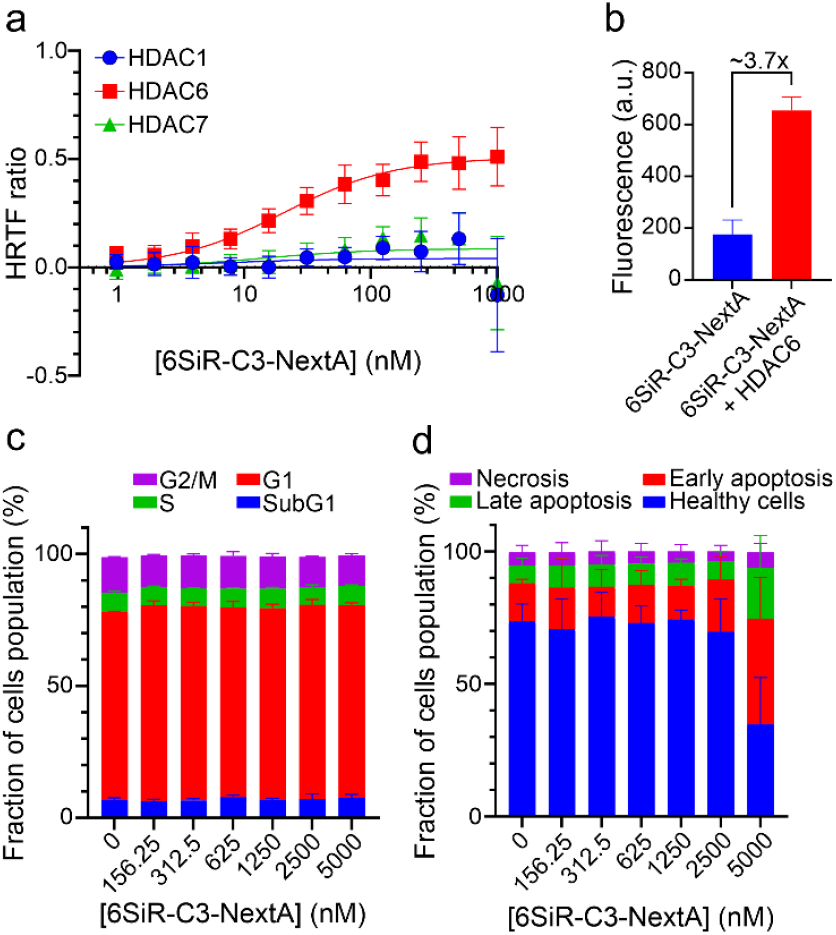
*In vitro* and *in cellulo* characterization of **6SiR-C3-NextA**. (a) Saturation binding curve of HTRF binding assay of **SiR-C3-NextA** against HDAC6, with HDAC1 and HDAC7 included as controls for selectivity. b) In vitro fluorogenicity of the 6SiR-C3-NextA probe. **6SiR-C3-NextA** (5 nM) was incubated for 1 hour at room temperature in PBS in either the absence or presence of cell lysate containing HDAC6 (30 nM). c) Cell cycle perturbation induced by **6SiR-C3-NextA** (1 µM in DMEM) after 24h incubation. d) Result of Annexin V Apoptosis Assay of SUP-M2 cells after a 24 hour treatment with increasing concentrations of **6SiR-C3-NextA**. Data points are mean□± □SD, n□= □3 independent experiments.

A highly desirable characteristic for a fluorescent probe is fluorogenicity, the ability to increase its fluorescence intensity upon binding to its target. This property significantly enhances the signal-to-background ratio by ensuring that unbound probe molecules in the cellular environment contribute minimally to the overall fluorescence. To assess this property, we measured the fluorescence emission of **6SiR-C3-NextA** in the presence and absence of cell lysate containing HDAC6 protein. Upon incubation with excess amount of HDAC6, the probe exhibited a significant, approximately 3.7-fold increase in fluorescence intensity compared to the probe alone in buffer (**Figure 3b**). This confirms that **6SiR-C3-NextA** is a modestly fluorogenic probe.

Crucially for live-cell imaging, **6SiR-C3-NextA** exhibited minimal cytotoxicity. Cell cycle analysis of HeLa cell showed no significant perturbations at concentrations up to 5 µM after 24 hours of treatment (**Figure 3c**). Furthermore, an Annexin V apoptosis assay with SUP-M2 cells revealed no significant increase in cell death at concentrations well above those required for effective imaging (≤1.25 µM) (**Figure 3d**). This low toxicity is a critical feature, ensuring that observations reflect native cellular processes rather than a response to a cytotoxic agent.

A key consideration for probes derived from inhibitors is their potential to perturb the biological system under observation. To evaluate the impact of **6SiR-C3-NextA** on its target’s enzymatic activity in cells, we measured its effect on the acetylation level of Lys40 of α-tubulin, the primary substrate of HDAC6^21^. HeLa cells were treated with increasing concentrations of the **6SiR-C3-NextA** and the parent inhibitor Nexturastat A, then acetylated α-tubulin levels were quantified by immunofluorescence imaging. A dose-dependent increase in acetylated α-tubulin was observed, consistent with the inhibitory activity of the parent compound, Nexturastat A (**Figure 4, S7**). At the typically concentration used for imaging (e.g., 100 nM), **6SiR-C3-NextA** did not cause a significant increase in tubulin acetylation, even after prolonged incubation. These results demonstrate that while the probe retains the inhibitory potential of its scaffold, it can be used for imaging under conditions that minimally perturb the endogenous enzymatic activity of HDAC6, ensuring that the visualized dynamics reflect the native state of the cell.

**Figure 4.**
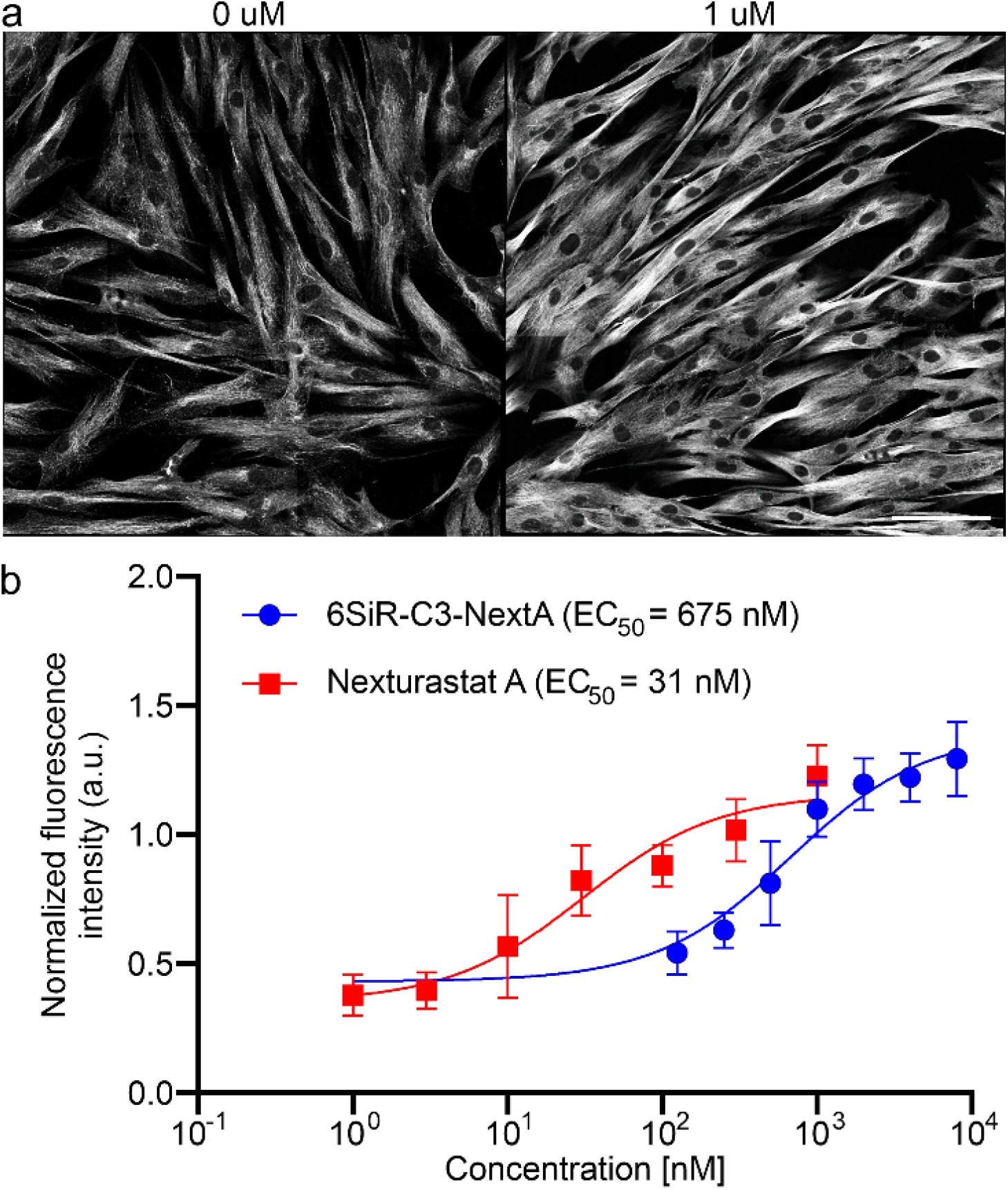
Quantification of probe-induced HDAC6 inhibition and tubulin hyperacetylation in living cells. (a) Representative immunofluorescence images of Human fibroblast cells treated for 24 hours with and without **6SiR-C3-NextA**. Cells were co-stained with antibodies for acetylated α-tubulin. Scale bar: 100 µm. Image acquisition settings are indicated in **Table S2**. (b) Dose-response curves plotting the normalized ratio of acetylated tubulin to β-tubulin. Data points are mean□± □SD, n□= □3 independent experiments.

The probe’s on-target specificity in cells was validated through co-localization experiments. In induced HDAC6 overexpressing U-2 OS cells fixed with paraformaldehyde, the signal from **6SiR-C3-NextA** showed excellent correlation with that of an anti-HDAC6 antibody (**Figure S6a-c**) (PCC = 0.85 ± 0.04). In live cells, the probe’s signal co-localized perfectly with an orthogonally labeled HaloTag-HDAC6 fusion protein (PCC =0.90 ± 0.03) (**Figure S6b-c**). This robust performance was consistent across a diverse panel of cell lines (HeLa, Huh-7, SH-SY5Y, 3T3, HEK293T, SUP-M2 and human dermal fibroblasts), where the probe signal was effectively competed away by an excess of a non-fluorescent inhibitor in live cell staining experiment (**Figure S7a**), and further validated with immunostaining, confirming its utility for imaging endogenous HDAC6 in various cell lines (**Figure S7b**). Notably, the probe remains functional after cell fixation with PFA, demonstrating its compatibility with immunofluorescence protocols.

#### Modulation of Probe Performance by Efflux Pump Inhibition

The intracellular concentration of a small-molecule probe is determined by the balance between membrane permeability and active removal by efflux pumps. To assess the impact of efflux effect on **6SiR-C3-NextA** performance, we quantified its intracellular accumulation in the presence and absence of known efflux pump inhibitors across our diverse cell panel. Our results indicate that the intracellular accumulation of **6SiR-C3-NextA** is significantly limited by active transport, as the probe is a substrate for efflux pumps. In six of the seven cell lines tested, co-incubation with an efflux pump inhibitor resulted in a substantial, up to 9-fold increase in the intracellular fluorescence signal (**Figure 5**). For example, 3T3 and Huh-7, showed a dramatic signal increase with the potent third-generation inhibitors Tariquidar^22^ and Zosuquidar^23^. Interestingly, the optimal inhibitor varied, as HeLa cells and H. Fibroblasts responded more strongly to the first-generation inhibitor Verapamil^24^. These findings confirm that **6SiR-C3-NextA** is an efflux pump substrate and, more importantly, offer a practical strategy to significantly improve its performance by co-incubating with an appropriate inhibitor to maximize the signal-to-noise ratio.

**Figure 5.**
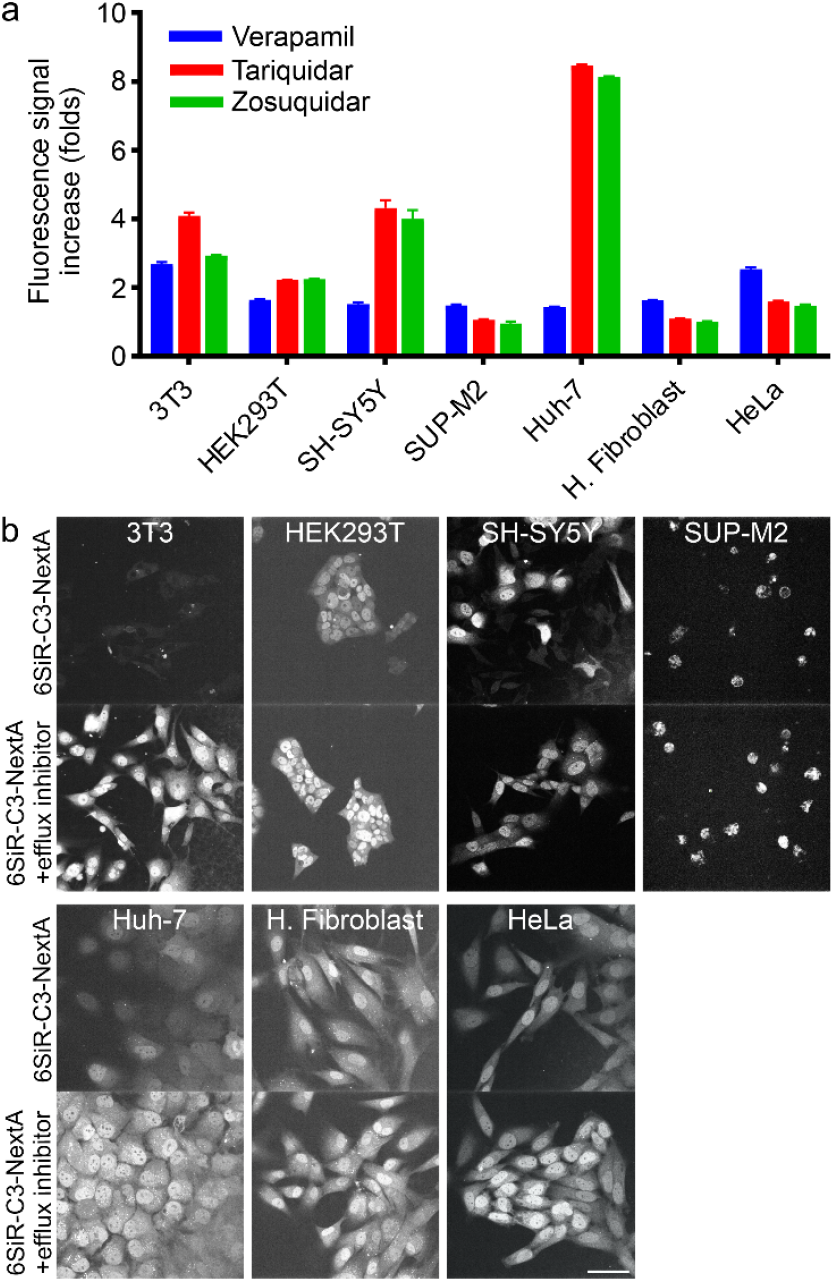
Efflux of **6SiR-C3-NextA** in living cells. (a) FACS analysis of efflux-inhibited staining experiments. Cells were stained at 37°C for 1h. Results are averages of three independent experiments (N=3) and presented as means with standard deviations. (b) Representative fluorescence microscopy images demonstrating active efflux on **6SiR-C3-NextA** staining performance. Seven cell lines from were stained with the probe alone (100 nM) or co-incubated with the probe and most effective efflux inhibitor as determined from FACS analysis. Scale bar: 50 µm. Image acquisition settings are indicated in **Table S2**.

#### Application in Live-Cell Imaging

Having validated its performance, we applied **6SiR-C3-NextA** to visualize the dynamics of HDAC6 in various cellular contexts.

#### Microtubule Association and Stress Granule Dynamics

As a primary tubulin deacetylase and a key component of the cellular stress response, HDAC6 is known to associate with microtubules and relocalize to stress granules (SGs) during stress^25, 26^. To investigate this phenomenon, we induced osmotic stress in U-2 OS cells that were overexpressing HDAC6 labeled with our **6SiR-C3-NextA** probe. This treatment triggered the rapid and dynamic recruitment of HDAC6 from a diffuse cytoplasmic pattern into distinct, punctate SGs. Using STED microscopy, we clearly observed this SG formation before and after the induction of stress. These results aligned with previous reports identifying HDAC6 as an integral SG component and additionally confirm the compatibility of our probe with super-resolution imaging techniques (**Figure 6a, Video S1 and Table S2**).

**Figure 6.**
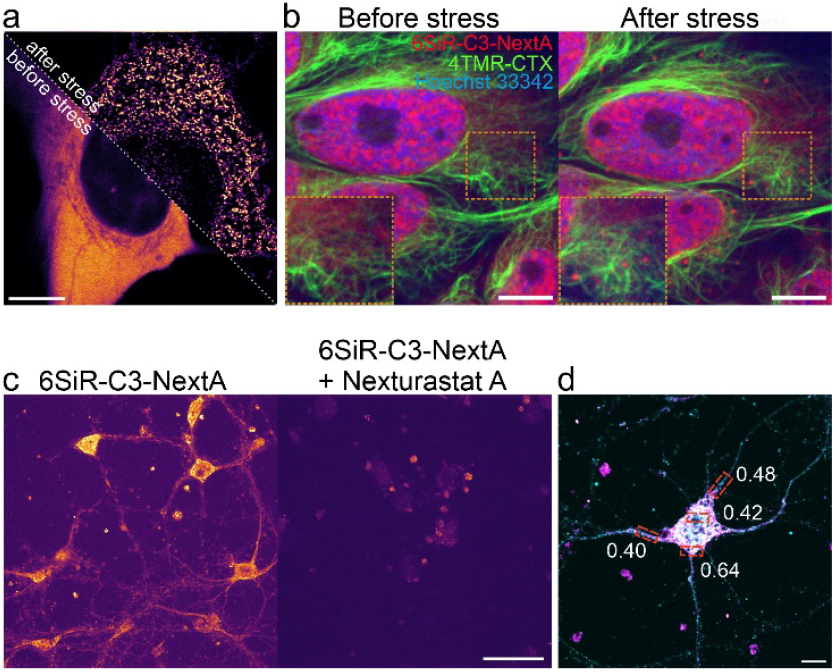
Fluorescence microscopy imaging of HDAC6 probe stained cells. (a) Representative STED imaging of induced HDAC6 expressing cell line stained with 100 nM of **6SiR-C3-NextA** in DMEM growth medium for 1h at 37°C, washed with HBSS, then applied 150 mM NaCl 30 mins before imaging. Scale bar: 10 μm. (b) Representative confocal image of living HeLa cell stained with 100 nM of **6SiR-C3-NextA**, 10 nM of tubulin probe (4TMR-CTX), 1 μg/mL of Hoechst 33342 and 10 μM of verapamil in DMEM growth medium for 1h at 37°C, washed with HBSS. Sig-nal was recorded from Cy5 channel before stress and after inducing osmotic stress with 150 mM NaCl in DMEM growth medium for 30mins. Scale bar: 10 μm. (c) Primary rat neuron cells were stain with 100 nM of probe (left) or co-staining with 100 µM of Nexturastat A for 1h at 37°C. Representative confocal image of immunostaining experiment. Scale bar: 100 μm. d) Representative confocal image of immunostaining experiment with 6SiR-C3-NextA (magneta) and HDAC6 antibody (cyan). Numbers show the Pearson correlation between the two channels in the red-boxed region. Scale bar: 10 μm. Image acquisition settings are indicated in **Table S2**.

To confirm this behavior at endogenous expression levels, we repeated the experiment in HeLa cells, co-incubating with the efflux pump inhibitor verapamil to enhance the signal, and live cell compatible microtubules probe (4TMR-CTX), reported previously by our lab. We again observed the robust recruitment of diffuse cytoplasmic HDAC6 into SGs upon stress induction, and also found that these granules forming along with microtubules, confirming its interaction with microtubules network, this was in line with published finding^26^. Interestingly, we also noted that the fraction of HDAC6 located within the nucleus appeared to relocalize towards the nuclear inner membrane under these stress conditions. This nuclear retention is consistent with the known regulation of the HDAC6 nuclear localization signal (NLS)^27^. Given that acetylation of NLS inhibits its function and results in cytoplasmic retention, the observed nuclear localization implies that this fraction of HDAC6 possesses a non-acetylated, functional NLS. This state would prevent its net export, thereby promoting its accumulation at the nuclear envelope. Collectively, this real-time visualization in multiple cell models confirms the probe’s ability to track the translocation of endogenous HDAC6 in response to physiological stimuli (**Figure 6b, Video S2)**.

#### Imaging in Primary Neurons

Given the importance of HDAC6 in neuronal health, we tested the probe in cultured primary rat neurons. The probe successfully entered the neurons and stained HDAC6; this signal was abolished by competition with excess Nexturastat A, confirming a specific binding mechanism (**Figure 6c**). However, co-localization experiments with an anti-HDAC6 antibody yielded only partial correlation (PCC ≈ 0.4–0.6). While this confirms that the probe binds to endogenous HDAC6 in neurons, it also suggests the presence of some off-target interactions in this complex cellular environment, highlighting a potential area for future probe refinement (**Figure 6d**). This observation is not entirely unexpected, as the highly complex and lipid-rich environment of neurons can sometimes lead to non-specific partitioning of hydrophobic dyes^28^.

## CONCLUSION

In summary, we have developed **6SiR-C3-NextA**, a novel, high-performance fluorescent probe for imaging endogenous HDAC6 in living cells. Through a rational design and systematic optimization process, we created a tool that exhibits high affinity, exceptional selectivity among tested HDAC cell lines. The probe is cell-permeable and well-suited for long-term and super-resolution imaging, as the working concentrations are over an order of magnitude lower than those found to induce cytotoxicity. We have demonstrated its utility in visualizing the enzyme’s association with microtubules and its recruitment to stress granules, highlighting its value for investigating the fundamental biology of HDAC6. This probe represents a significant addition to the chemical biology toolbox and is expected to facilitate new discoveries regarding HDAC6’s role in neurodegenerative pathologies, particularly those driven by protein aggregation and cytoskeletal dysfunction, such as Alzheimer’s and Parkinson’s disease.

## AUTHOR INFORMATION

### Author Contributions

V.T and G.L. conceived and planned the study; V.T. and G.L performed the experiments and analyzed the data. T.K. constructed and maintained cell lines. J.B., and G.L. established the screening approach. V.T and G.L wrote the initial draft; all authors contributed to the final version of the manuscript.

### Funding Sources

Any funds used to support the research of the manuscript should be placed here (per journal style).

### Notes

G.L. is a co-inventor on the patent (EP2748173B1 and US9346957B2, applicant EPFL) describing SiR and its derivatives.

## ACKNOWLEDGMENT

The authors are grateful to Dr. Vladimir Belov, Jan Seikowski, Jens Schimpfhauser, and Jürgen Bienert (Facility for Synthetic Chemistry, MPI-NAT), Dr. H. Frauendorf, and the central analytics’ team (Institute for Organic and Biomolecular Chemistry, Georg-August University, Göttingen) for acquiring NMR and ESI-MS spectra, to Dr. Peter Lenart, Dr. Antonio Politi and Jasmin Jakobi (Facility for Light Microscopy, MPINAT) for the possibility of performing live-cell spinning disk confocal microscopy and flow cytometry. Văn Thắng Nguyễn was supported by the Ph.D. program Genome Science—International Max Planck Research School.

## ABBREVIATIONS

(FPs): Fluorescent proteins
HTRF: (Homogeneous Time Resolved Fluorescence)
HDAC: (Histone deacetylase)
SiR: (Silicon Rhodamine)
TMR: (Carboxytetramethylrhodamine)
(PCC): Pearson’s Correlation Coefficient
SAR: (Structure-Activity Relationship)
(SGs): stress granules

