## Supplementary information for "A Highly Selective, Cell-Permeable Fluorescent Probe for Imaging Histone Deacetylase 6 in Live Cells"

#### Supplementary Figures

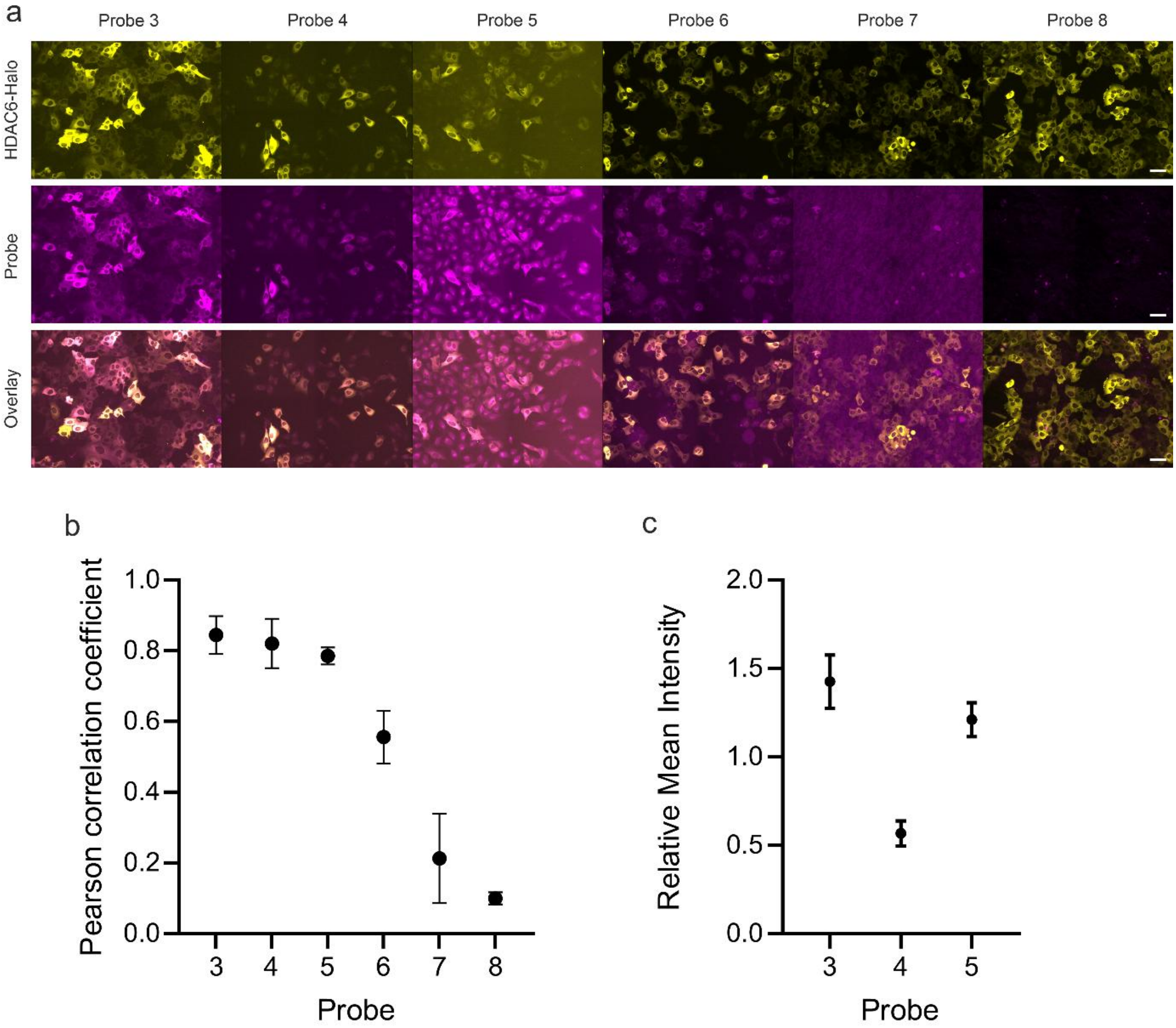

**Figure S1. Optimization of fluorophores of choice.** (a) Representative fluorescence images from the assay of SiR-based probes (**3, 4, 5**) and TMR-based analogues (**6, 7, 8**) with live U2OS cells expressing HDAC6-HaloTag. Scale bar: 100 µm. (b) Quantification of probe performance by measuring the Pearson's Correlation Coefficient (PCC) for co-localization with the HDAC6-HaloTag reference signal in live cells. (c) Comparison of signal intensity for the SiR-based probes. Results are averages of three independent experiments (N=3) and presented as means with standard deviations

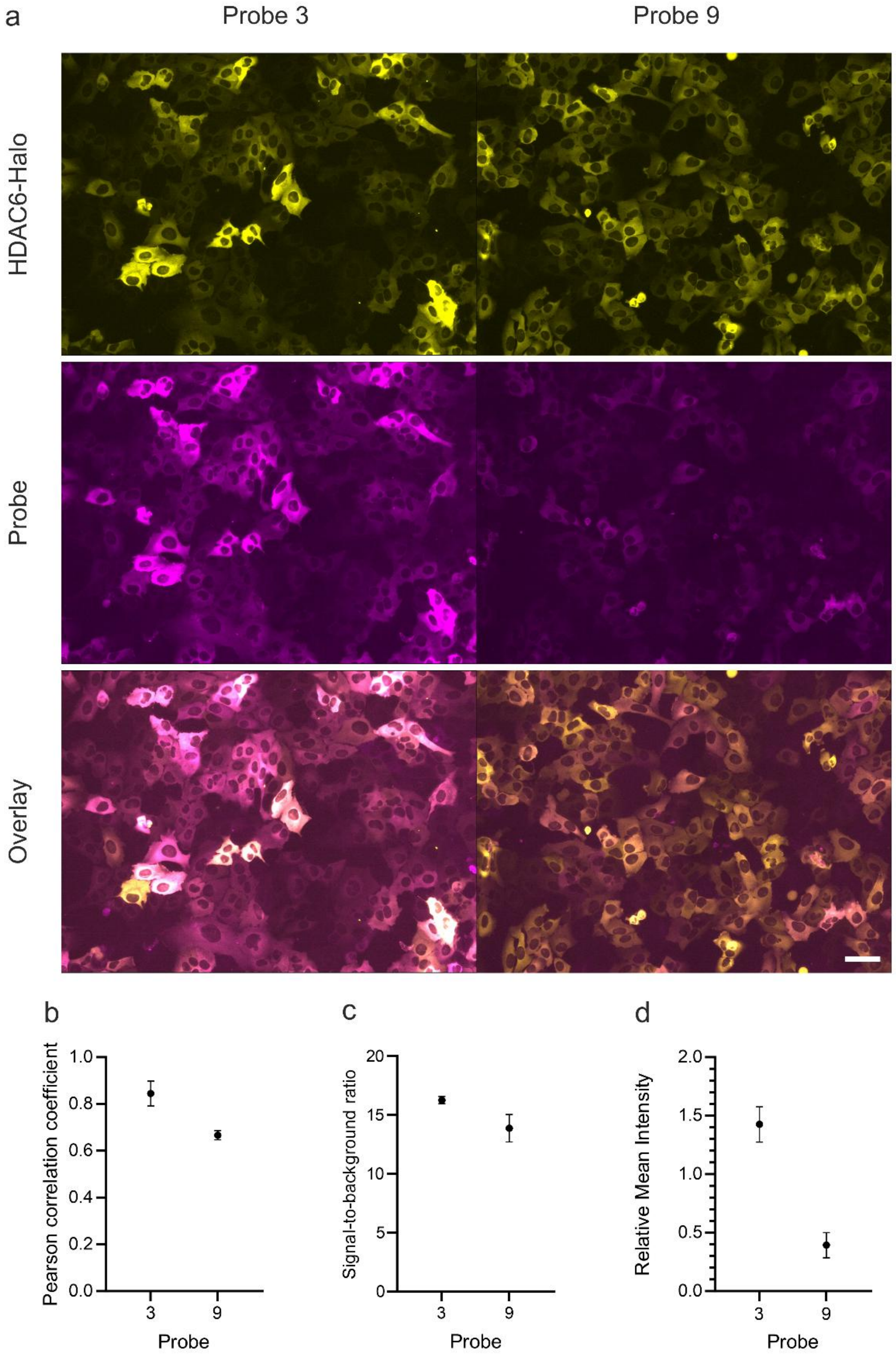

**Figure S2. Evaluation of scaffold fluorination on probe performance.** (a) Representative fluorescence images of live U2OS cells expressing HDAC6-HaloTag, treated with the probe **3** or its perfluorinated **9**. Scale bar: 100 µm. Quantitative comparison of the probes: (b) Pearson's Correlation Coefficient (PCC) with the HDAC6-Halo reference, (c) signal-to-background ratio, and (d) relative mean signal intensity. Results are averages of three independent experiments (N=3) and presented as means with standard deviations.

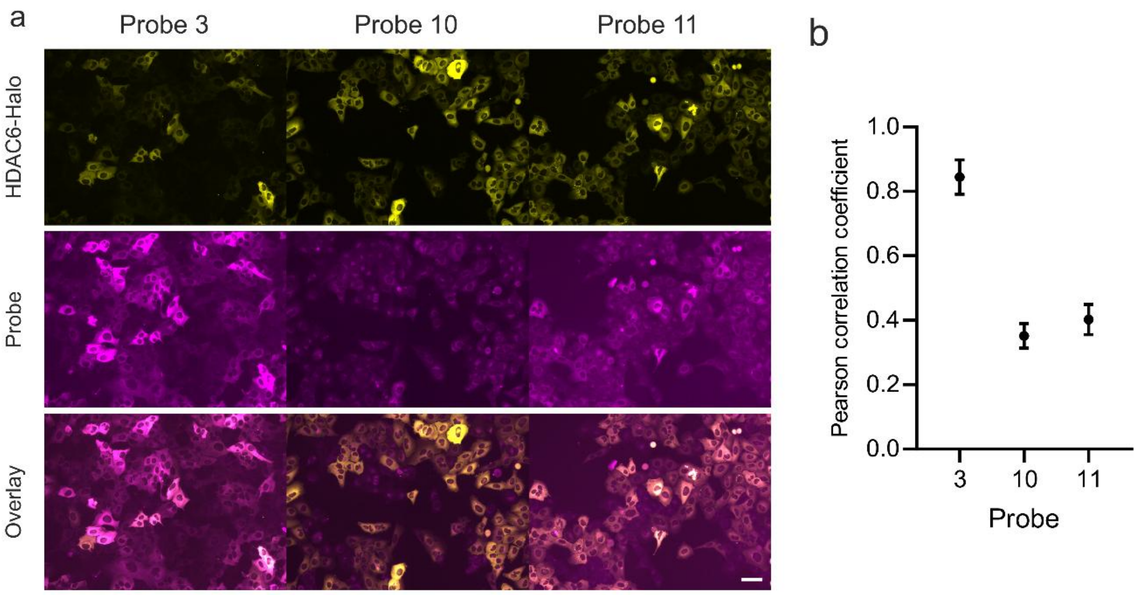

**Figure S3. Evaluation of alternative zinc-binding groups (ZBGs) on probe performance.** (a) Representative fluorescence images comparing the probe **3** with analogues **10** and **11**, which incorporate a difluoromethyloxadiazole (DFMO) ZBG, in live U2OS cells expressing HDAC6-HaloTag. Scale bar: 100 µm (b) Quantification of co-localization via Pearson's Correlation Coefficient (PCC). Results are averages of three independent experiments (N=3) and presented as means with standard deviations.

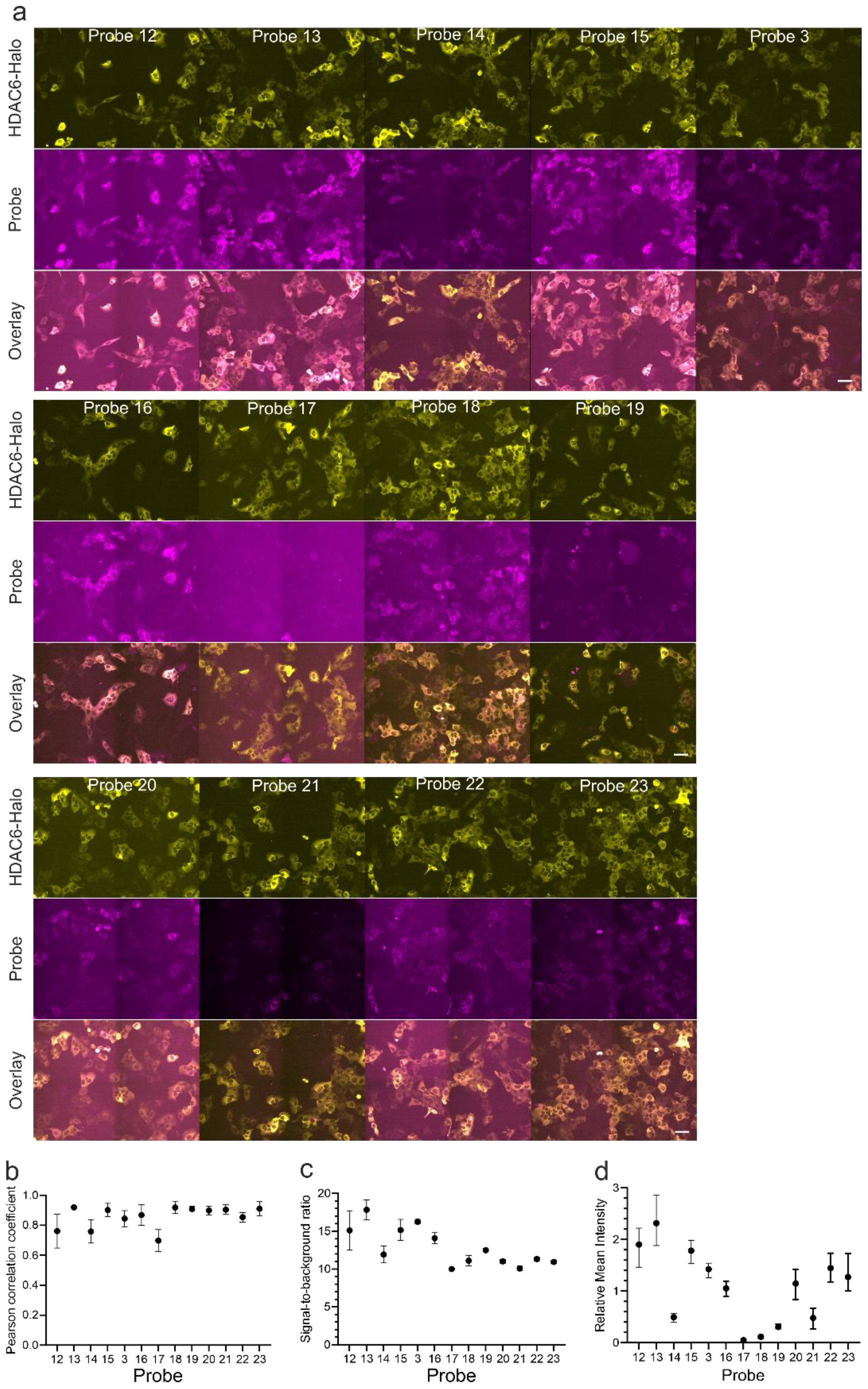

**Figure S4. Systematic optimization of the linker for the HDAC6 probe.** (a) Representative fluorescence images of live U2OS cells expressing HDAC6-HaloTag, treated with a panel of probes synthesized with varying linker lengths and compositions. Scale bar: 100 μm. (b-d) Quantitative analysis of probe performance by the (b) Pearson's Correlation Coefficient, (c) the signal-to-background ratio and (d) relative mean intensity. Results are averages of three independent experiments (N=3) and presented as means with standard deviations.

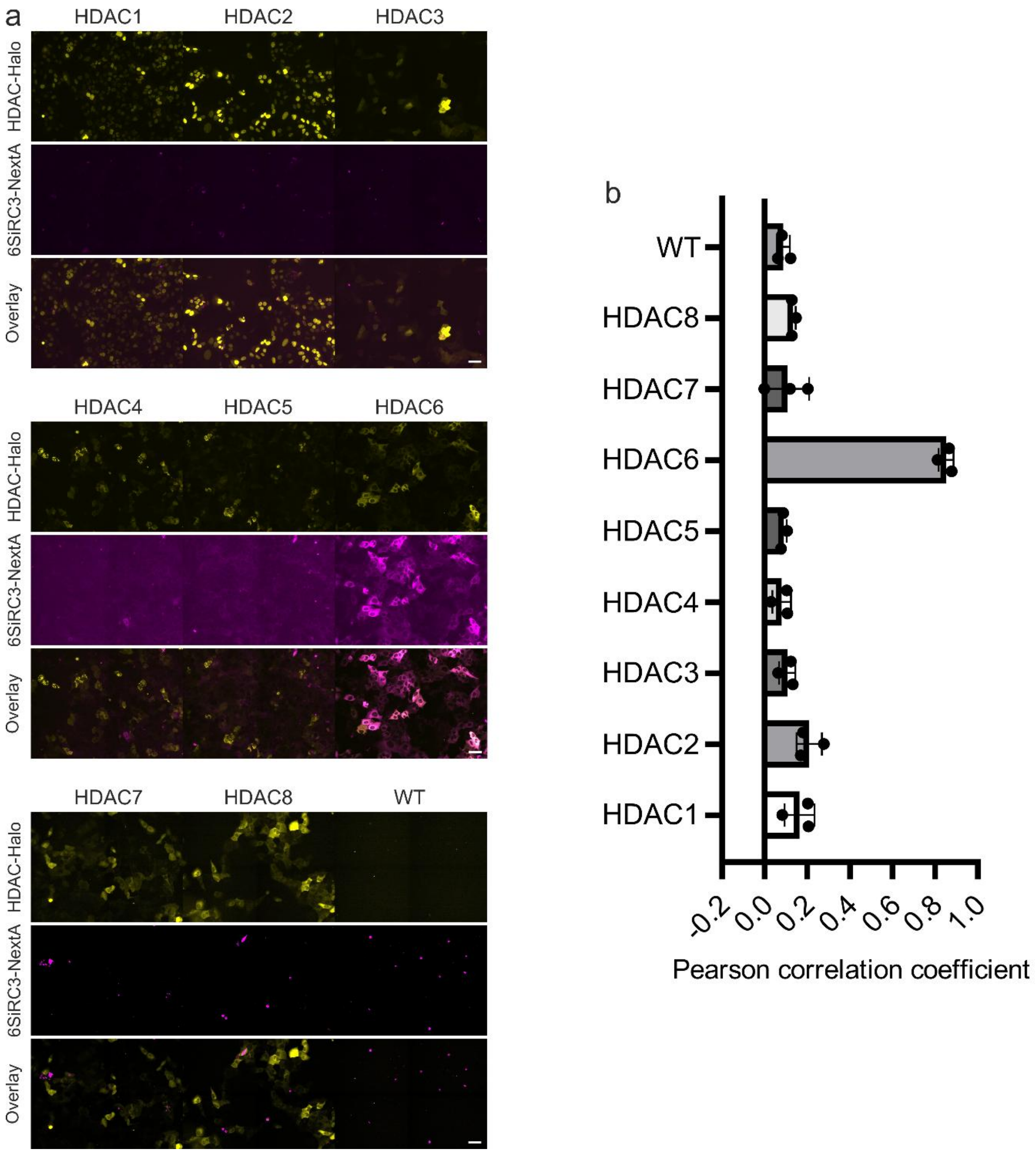

**Figure S5. Evaluation of selectivity of 6SiR-C3-NextA.** a) Representative fluorescence images from the assay. Scale bar: 200 µm. (b) Quantitative co-localization analysis. Pearson's Correlation Coefficient (PCC) between the probe and HaloTag signals. Results are averages of three independent experiments (N=3) and presented as means with standard deviations.

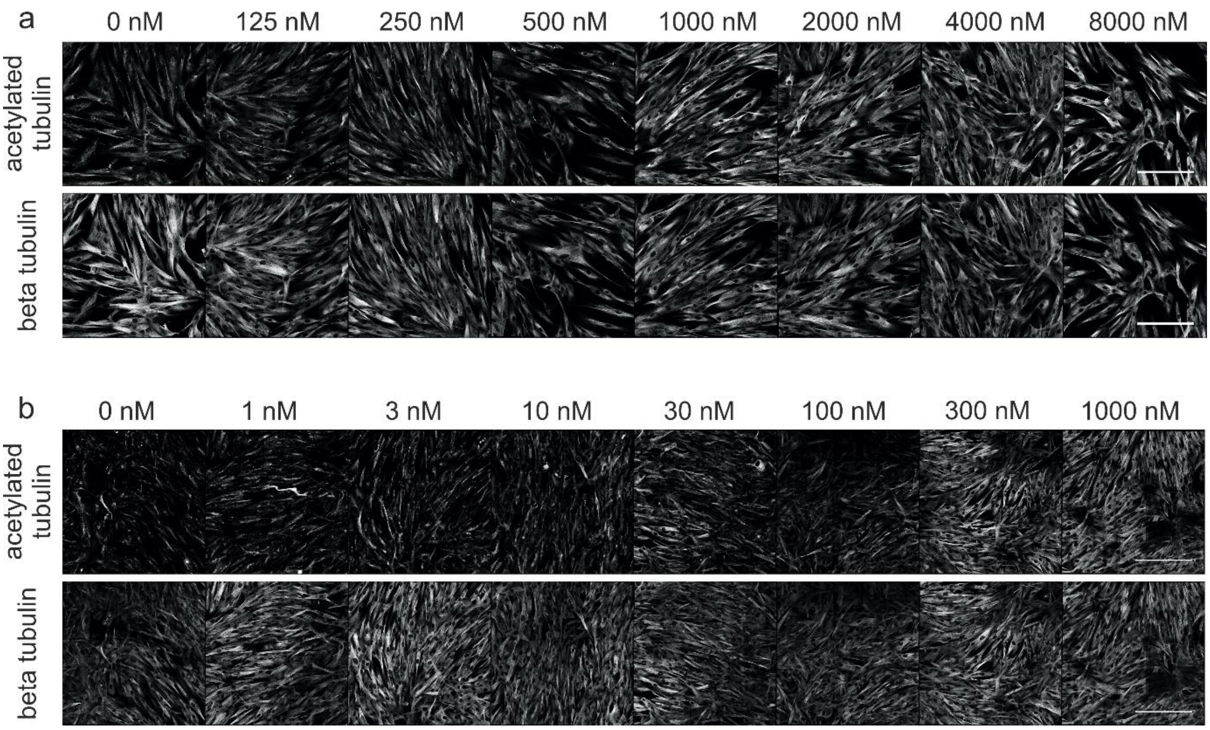

**Figure S6. Quantification of Cellular Target Engagement by Measuring Probe-Induced Tubulin Hyperacetylation.** Representative immunofluorescence images of Human fibroblast cells treated for 24 hours with increasing concentrations of the probe (a) 6SiR-C3-NextA and (b) Nexturastat A. Cells were co-stained with antibodies for acetylated α-tubulin (top row) and β-tubulin (bottom row).

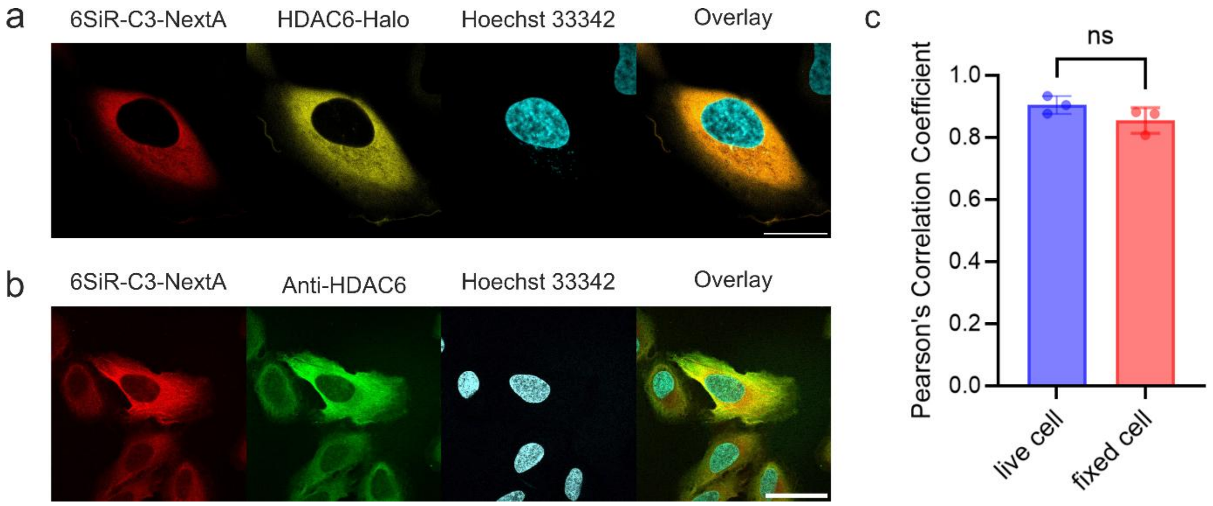

**Figure S7. Validation of Probe Specificity for HDAC6 in living and fixed cells.** (a) Representative confocal image of living U2OS cells expressing HaloTag-HDAC6 cell stained with 100 nM of 6SiR-C3-NextA, 100 nM of Halotag substrate 6TMR-PEG2-Halo, and 1 µg/mL of Hoechst 33342 in DMEM growth medium for 1h at 37°C, washed with HBSS. (b) Representative confocal image of fixed U2OS cells expressing HaloTag-HDAC6 cell. Cells were stained with 100 nM 6SiR-C3-NextA and 1 µg/mL of Hoechst 33342 in DMEM growth medium for 1h at 37°C, fixed with 4% paraformaldehyde (PFA). (c) Quantitative co-localization analysis of probe and reference signal. Results are averages of three independent experiments (N=3) and presented as means with standard deviations. Scale bar: 20 µm

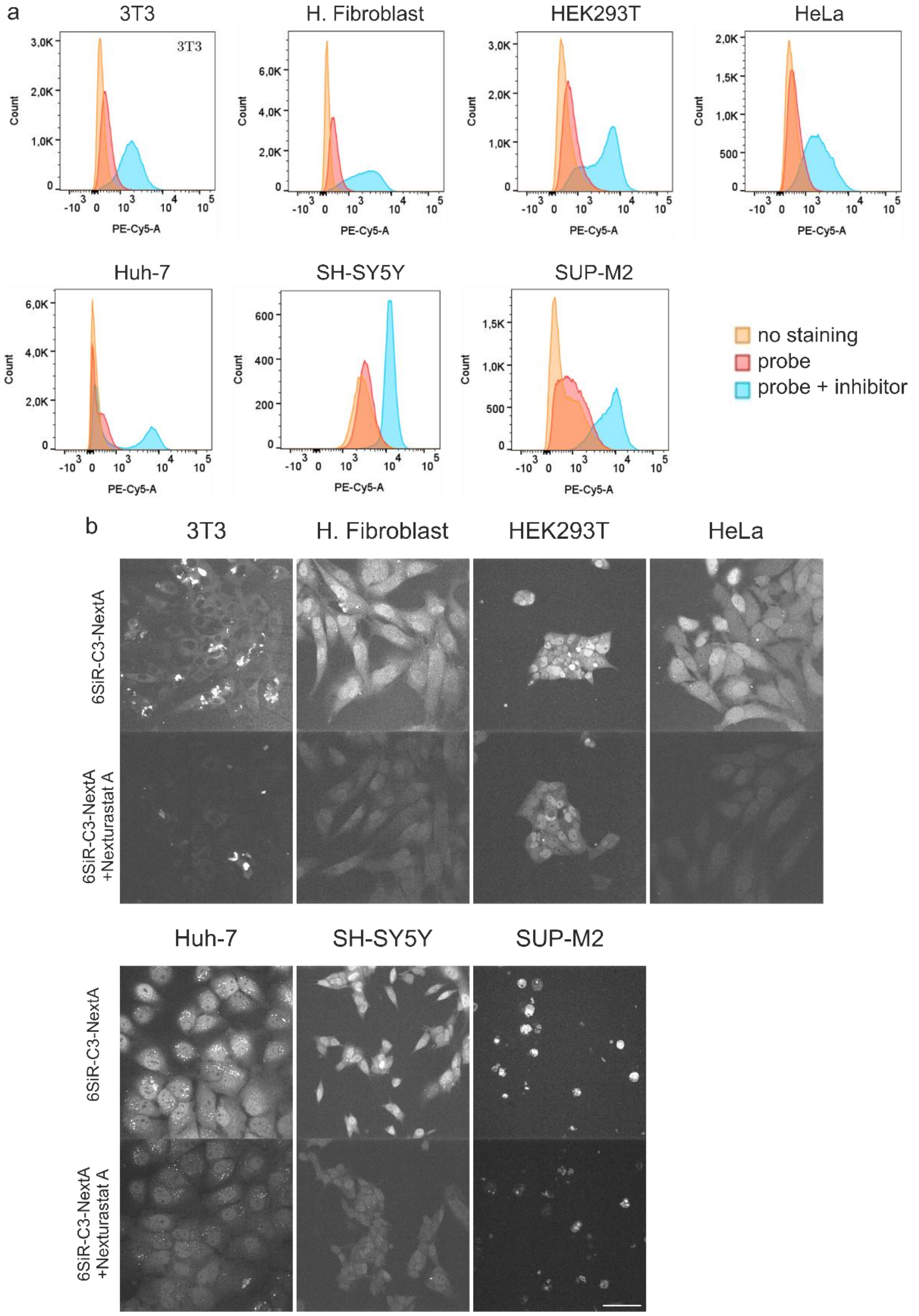

**Figure S8. Validation of Probe Specificity by Competitive Inhibition Across Diverse Cell Lines.** (a) Representative flow cytometry histograms show the fluorescence intensity of cell populations. (b) Representative fluorescence microscopy images demonstrating the on-target specificity of 6SiR-C3-NextA. Cell lines were stained with the probe alone (100 nM) or co-incubated with the probe and an excess of a non-fluorescent competitive inhibitor (10 uM). Scale bar: 50 µm.

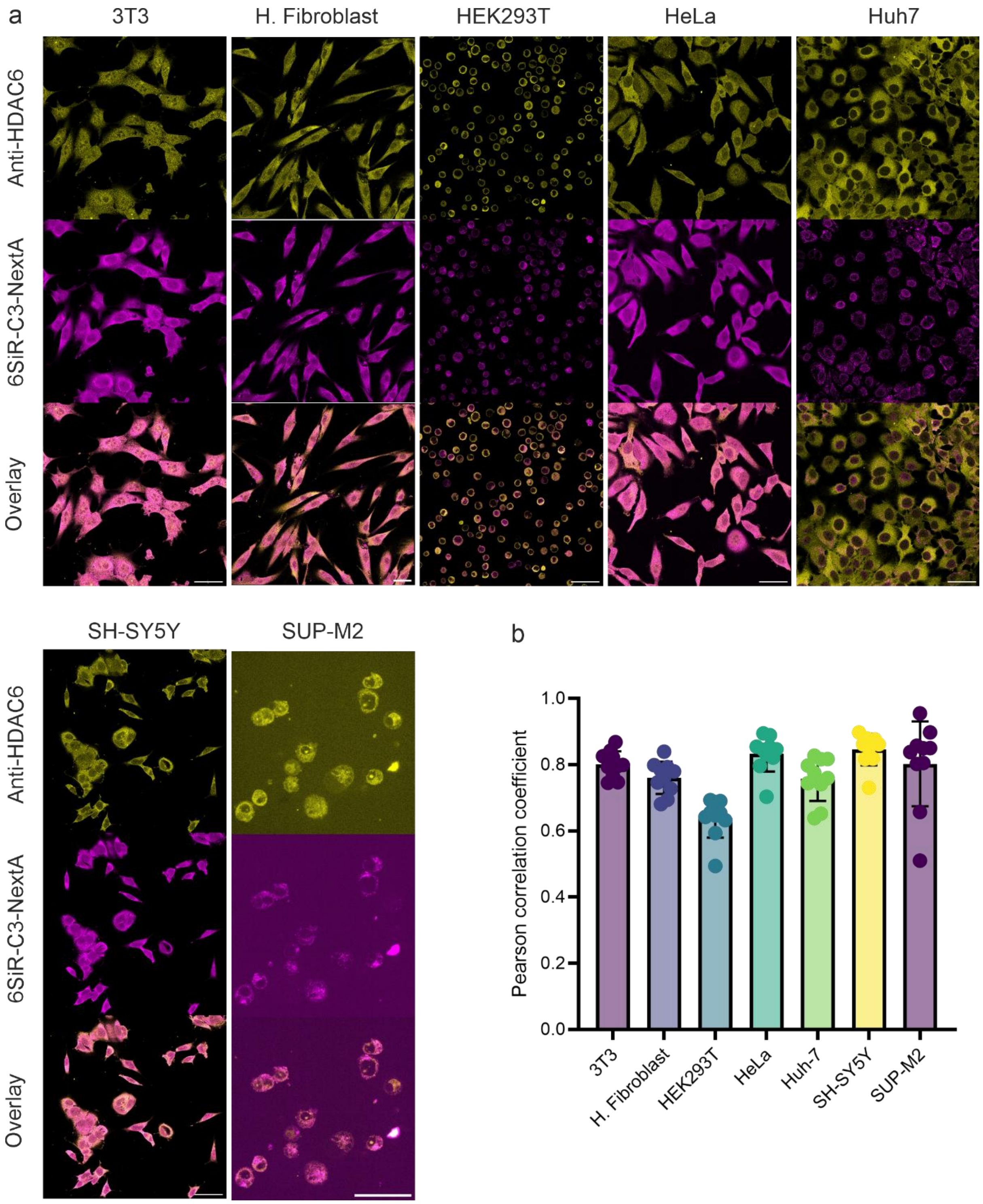

**Figure S9. Validation of Probe Specificity for Endogenous HDAC6 Across a Diverse Panel of Cell Lines.** (a) Representative confocal images of cells co-stained with 6SiR-C3-NextA and a specific anti-HDAC6 antibody. (b) Quantitative analysis of co-localization across the entire panel of different cell lines. Each point on the graph represents an individual cell, with bars indicating the mean ± SD. Scale bar: 50 µm.

#### Supplementary Tables

#### Table S1. Primers used for Gateway cloning of HDAC genes.

| Cell lines | Prime sequences |
|---|---|
| HDAC1-Halo | Attb_Insert_FOR:<br>GGGGACAAGTTTGTACAAAAAAGCAGGCTTAACCATGGCCCAGACCCAGGG<br>attB_Insert_REV:<br>GGGGACCACTTTGTACAAGAAAGCTGGGTTGGCCAGCTTCACCTCCTCC |
| HDAC2-Halo | Attb_Insert_FOR:<br>GGGGACAAGTTTGTACAAAAAAGCAGGCTTAACCATGGCCTACAGCCAGGG<br>attB_Insert_REV:<br>GGGGACCACTTTGTACAAGAAAGCTGGGTTAGGATTAGACAGCTGCTCGGAC |
| HDAC3-Halo | Attb_Insert_FOR:<br>GGGGACAAGTTTGTACAAAAAAGCAGGCTTAACCATGGCCAAGACCGTGGC<br>attB_Insert_REV:<br>GGGGACCACTTTGTACAAGAAAGCTGGGTTGATCTCCACATCGCTCTCCTTATCG |
| HDAC4-Halo | Attb_Insert_FOR:<br>GGGGACAAGTTTGTACAAAAAAGCAGGCTTAACCATGGCTTCGTCAGGGTCCCC<br>attB_Insert_REV:<br>GGGGACCACTTTGTACAAGAAAGCTGGGTGGGCTGGGGGAGGGGGAGC |
| HDAC5-Halo | Attb_Insert_FOR:<br>GGGGACAAGTTTGTACAAAAAAGCAGGCTTAACCATGAATAGCCCCAATGAATCTGAT<br>attB_Insert_REV:<br>GGGGACCACTTTGTACAAGAAAGCTGGGTCAGCGCTGGCTCCTGCTCCATGGCAGG |
| HDAC6-Halo | Attb_Insert_FOR:<br>GGGGACAAGTTTGTACAAAAAAGCAGGCTTAACCATGTCCAACGGGCAGCCTCC<br>attB_Insert_REV:<br>GGGGACCACTTTGTACAAGAAAGCTGGGTCTCGTAGGCATCATCCTG |
| HDAC7-Halo | Attb_Insert_FOR:<br>GGGGACAAGTTTGTACAAAAAAGCAGGCTTAACCATGGCCAACGTGGCCAAGCC<br>attB_Insert_REV:<br>GGGGACCACTTTGTACAAGAAAGCTGGGTATACTGCGGTGTGTACC |
| HDAC8-Halo | Attb_Insert_FOR:<br>GGGGACAAGTTTGTACAAAAAAGCAGGCTTAACCATGGAGGAGCCAGAGGAGC<br>attB_Insert_REV:<br>GGGGACCACTTTGTACAAGAAAGCTGGGTTCACCACGTGCTTCAGATTGC |

**Table S2. Image acquisition parameters of confocal and STED microscopes.**

| Figure | Microscope | Laser power (%) | STED 775 nm laser power (%) | Emission filter or monochromator settings (nm) | Exposure (ms) | Pixel dwell time (µs) | Pixel size xy (nm) | Comment |
|---|---|---|---|---|---|---|---|---|
| Fig 5b | Nikon Spinning-disk confocal | 640 (20-80) | - | ET665LP | 100 | - | 111 x 111 nm | |
| Fig.6b | | 405 (20);<br>561 (5);<br>640 (20) | | ET460/50<br>609/52<br>ET665LP | | | | |
| Fig.6c | | 640 (20) | | ET665LP | | | | |
| Fig.6d | | 561 (10);<br>640 (20) | | 609/52<br>ET665LP | | | | |
| Fig.S8b | | 640 (20-50) | | ET665LP | | | | |
| Fig.S9 | | 561 (5);<br>640 (20-50) | | 609/52<br>ET665LP | | | | |
| Video S1 | | 405 (20);<br>561 (5);<br>640 (20) | | ET460/50<br>609/52<br>ET665LP | | | | Interval time: 10s, Z-stack 3 planes × 200nm |
| Video S2 | | 405 (20);<br>561 (5);<br>640 (20) | | ET460/50<br>609/52<br>ET665LP | | | | Interval time: 10s, Z-stack 3 planes × 200nm |
| Fig.4a | Abberior Facility Line | 561 (20);<br>640 (20) | - | 580 – 630<br>650 – 720 | - | 1 | 80 | |
| Fig.6a | | 640 (20) | 10 | 650 – 720 | | 5 | 20 | Line accum.: 2 |
| Fig.S6 | | 561 (20);<br>640 (20); | - | 580 – 630<br>650 – 720 | | 1 | 80 | |
| Fig.S7a | | 405 (2);<br>561 (10);<br>640 (10) | - | 440 – 550<br>580 – 630<br>650 – 720 | | 1 | 80 | |
| Fig.S7b | | 405 (2);<br>561 (10);<br>640 (10) | - | 440 – 550<br>580 – 630<br>650 – 720 | | 1 | 80 | |

**Supplementary Videos**

**Video S1. Time-lapse series of living U2OS induced HDAC6 expressing cells under osmotic stress.** Cells were stained in DMEM for 1 h at 37°C with 100 nM 6SiR-C3-NextA and 0.1 μg/mL Hoechst 33342, washed with HBSS, and then treated with DMEM containing 150 mM NaCl to induce osmotic stress. Scale bar 20 μm.

**Video S2. Time-lapse series of living HeLa under osmotic stress.** Cells were stained in DMEM for 1 h at 37°C with 100 nM 6SiR-C3-NextA, 10 nM 4TMR-CTX, 0.1 μg/mL Hoechst 33342, and 10 μM verapamil, washed with HBSS, and then treated with DMEM containing 150 mM NaCl to induce osmotic stress. Scale bar 10 μm.

#### Supplementary Methods

#### Molecular biology and biochemical methods

*Cell line construction*

The pHDACx-Halo plasmid was constructed using a Gateway-compatible destination vector system which was characterized previously. Plasmids contain a cytomegalovirus-tetracycline operator (CMV-TetO2) promoter which is “switched on” by the addition of doxycycline, along with puromycin resistance and EBNA-1 cassettes for selection and episomal maintenance. The human HDACx coding gene was PCR amplified using specific primers (see **Table S1**) and cloned into the destination vector containing a C-terminal Halo-tag by Gateway LR. The final construct sequence has been verified by sequencing.

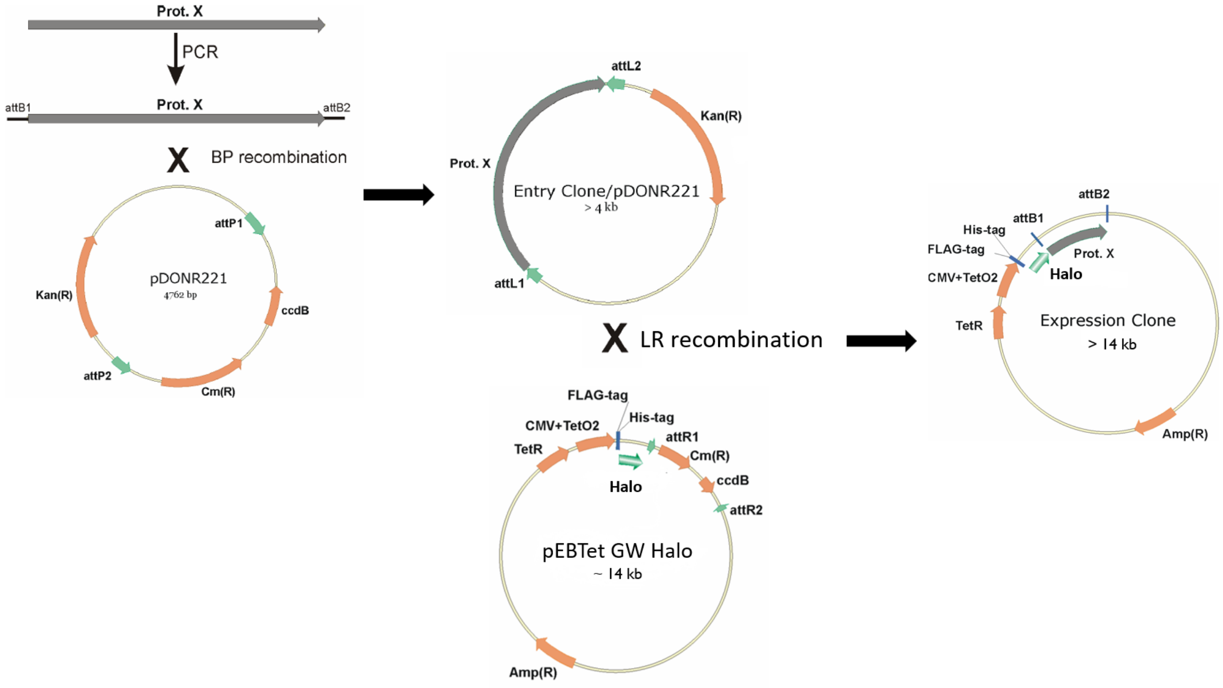

General strategy of Halo-tagged proteins construction

#### HDAC1

ATGGCCCAGACCCAGGGCACACGGAGAAAGGTGTGCTACTATTACGACGGCGATG
TGGGCAATTATTACTATGGCCAGGGCCACCCCATGAAGCCTCACAGGATCCGCATG
ACCCACAATCTGCTGCTGAACTACGGCCTGTATCGGAAGATGGAGATCTATAGACC
CCACAAGGCCAACGCCGAGGAGATGACAAAGTATCACTCTGACGATTACATCAAG
TTCCTGCGGAGCATCAGACCAGACAATATGTCTGAGTACAGCAAGCAGATGCAGC
GGTTCAACGTGGGCGAGGACTGCCCCGTGTTTGATGGCCTGTTCGAGTTTTGTCA
GCTGTCCACCGGAGGATCCGTGGCATCTGCCGTGAAGCTGAATAAGCAGCAGACA
GACATCGCAGTGAACTGGGCAGGAGGACTGCACCACGCCAAGAAGAGCGAGGC
CTCCGGCTTCTGTTATGTGAATGATATCGTGCTGGCCATCCTGGAGCTGCTGAAGT
ATCACCAGAGGGTGCTGTACATCGACATCGATATCCACCACGGCGACGGAGTGGA
GGAGGCCTTTTACACCACAGATCGCGTGATGACCGTGTCTTTCCACAAGTACGGC
GAGTATTTTCCTGGAACAGGCGACCTGAGGGATATCGGAGCAGGCAAGGGCAAGT
ACTATGCCGTGAATTATCCACTGAGAGACGGCATCGACGATGAGAGCTACGAGGC

CATCTTCAAGCCTGTGATGTCCAAAGTGATGGAGATGTTTCAGCCATCTGCCGTGG
TGCTGCAGTGCGGATCCGACTCTCTGAGCGGCGATAGGCTGGGCTGCTTTAACCT
GACCATCAAGGGCCACGCCAAGTGCGTGGAGTTCGTGAAGAGCTTTAATCTGCCA
ATGCTGATGCTGGGAGGAGGAGGATATACCATCAGGAACGTGGCCCGCTGTTGGA
CCTACGAGACAGCCGTGGCCCTGGATACAGAGATCCCTAATGAGCTGCCATACAA
CGACTATTTCGAGTACTTTGGCCCAGATTTCAAGCTGCACATCTCCCCCTCTAATAT
GACCAACCAGAATACAAACGAGTACCTGGAGAAGATCAAGCAGAGGCTGTTTGA
GAACCTGAGGATGCTGCCACACGCACCAGGAGTGCAGATGCAGGCAATCCCAGA
GGACGCAATCCCTGAGGAGAGCGGCGACGAGGATGAGGACGATCCTGACAAGCG
GATCTCCATCTGCAGCTCCGATAAGAGAATCGCCTGTGAGGAGGAGTTCAGCGAT
TCCGAGGAGGAGGGAGAGGGAGGAAGGAAGAATTCTAGCAACTTTAAGAAGGCC
AAGCGCGTGAAGACCGAGGACGAGAAGGAGAAGGATCCCGAGGAGAAGAAGGA
GGTGACCGAGGAGGAGAAGACAAAGGAGGAGAAGCCTGAGGCCAAGGGCGTGA
AGGAGGAGGTGAAGCTGGCC

**HDAC2**

ATGGCCTACAGCCAGGGCGGCGGCAAGAAGAAGGTGTGCTACTATTACGACGGCG
ATATCGGCAACTATTACTATGGCCAGGGCCACCCCATGAAGCCTCACAGGATCCGC
ATGACCCACAACCTGCTGCTGAATTACGGCCTGTATAGGAAGATGGAGATCTATAG
GCCACACAAGGCAACCGCAGAGGAGATGACAAAGTATCACAGCGACGAGTACAT
CAAGTTCCTGCGGTCCATCAGACCCGATAACATGAGCGAGTACTCCAAGCAGATG
CAGCGGTTCAATGTGGGCGAGGACTGCCCCGTGTTCGATGGCCTGTTCGAGTTTT
GTCAGCTGTCCACCGGAGGATCTGTGGCAGGAGCAGTGAAGCTGAACAGGCAGC
AGACAGACATGGCAGTGAATTGGGCAGGAGGACTGCACCACGCCAAGAAGTCTG
AGGCCAGCGGCTTCTGCTATGTGAATGATATCGTGCTGGCCATCCTGGAGCTGCTG
AAGTATCACCAGCGGGTGCTGTACATCGACATCGATATCCACCACGGCGACGGAG
TGGAGGAGGCCTTTTACACCACAGATAGAGTGATGACCGTGTCCTTCCACAAGTA
CGGCGAGTATTTTCCAGGCACAGGCGACCTGAGGGATATCGGCGCCGGCAAGGGC
AAGTACTATGCCGTGAACTTCCCCATGCGCGACGGCATCGACGATGAGTCTTATGG
CCAGATCTTTAAGCCAATCATCTCTAAAGTGATGGAGATGTACCAGCCAAGCGCCG
TGGTGCTGCAGTGTGGAGCAGACAGCCTGTCCGGCGATAGACTGGGCTGCTTCAA
TCTGACCGTGAAGGGCCACGCCAAGTGCGTGGAGGTGGTGAAGACCTTCAACCT
GCCTCTGCTGATGCTGGGAGGAGGAGGCTATACCATCAGGAATGTGGCCCGCTGC
TGGACCTACGAGACAGCCGTGGCCCTGGATTGTGAGATCCCTAACGAGCTGCCAT
ACAATGACTATTTCGAGTACTTTGGCCCCGATTTCAAGCTGCACATCTCTCCTAGC
AACATGACCAACCAGAATACACCAGAGTACATGGAGAAGATCAAGCAGCGGCTG
TTTGAGAATCTGAGAATGCTGCCACACGCACCAGGAGTGCAGATGCAGGCAATCC
CTGAGGACGCCGTGCACGAGGATTCCGGCGACGAGGATGGCGAGGACCCAGATA
AGCGGATCAGCATCAGAGCCTCCGACAAGAGGATCGCCTGTGATGAGGAGTTTTC
CGACTCTGAGGATGAGGGAGAGGGAGGCCGGAGAAACGTGGCAGACCACAAGA
AGGGCGCCAAGAAGGCCCGCATCGAGGAGGATAAGAAGGAGACCGAGGACAAG
AAGACAGATGTGAAGGAGGAGGACAAGTCTAAGGATAACAGCGGCGAGAAGAC
CGACACAAAGGGCACAAAGTCCGAGCAGCTGTCTAATCCT

**HDAC3**

ATGGCCAAGACCGTGGCCTACTTCTATGACCCCGATGTGGGCAACTTTCACTACGG
AGCAGGACACCCAATGAAGCCTCACAGGCTGGCCCTGACACACTCTCTGGTGCTG
CACTACGGCCTGTATAAGAAGATGATCGTGTTCAAGCCATATCAGGCCAGCCAGCA
CGACATGTGCAGGTTCCACTCCGAGGACTACATCGATTTCTGCAGCGCGTGTCTC
CCACCAATATGCAGGGCTTCACAAAGAGCCTGAACGCCTTTAATGTGGGCGACGA
TTGCCCAGTGTTTCCCGGCCTGTTCGAGTTTTGTAGCAGGTATACCGGAGCATCCC
TGCAGGGAGCAACACAGCTGAACAATAAGATCTGCGACATCGCCATCAACTGGGC
AGGAGGACTGCACCACGCCAAGAAGTTCGAGGCCAGCGGCTTTTGTTACGTGAAT
GATATCGTGATCGGCATCCTGGAGCTGCTGAAGTATCACCCTCGGGTGCTGTACAT
CGACATCGATATCCACCACGGCGATGGCGTGCAGGAGGCCTTCTATCTGACCGAC
AGAGTGATGACAGTGTCCTTTCACAAGTACGGCAACTATTTCTTTCCCGGCACCGG
CGATATGTACGAAGTGGGCGCCGAGAGCGGCAGGTACTATTGTCTGAACGTGCCT
CTGCGCGACGGCATCGACGATCAGTCCTATAAGCACCTGTTCCAGCCTGTGATCAA
TCAGGTGGTGGACTTCTACCAGCCAACATGCATCGTGCTGCAGTGTGGAGCAGAT
TCCCTGGGATGCGACAGGCTGGGCTGTTTCAATCTGTCTATCAGAGGCCACGGCG
AGTGCGTGGAGTACGTGAAGTCCTTTAACATCCCTCTGCTGGTGCTGGGAGGAGG
AGGATATACCGTGCGGAATGTGGCCAGATGTTGGACCTACGAGACATCTCTGCTGG
TGGAGGAGGCCATCTCTGAGGAGCTGCCATACAGCGAGTATTTCGAGTACTTTGCC
CCCGATTTCACCCTGCACCCTGACGTGTCCACAAGGATCGAGAACCAGAATTCTC
GCCAGTATCTGGACCAGATCCTGCAGACCATCTTTGAGAACCTGAAGATGCTGAAT
CACGCCCCATCCGTGCAGATCCACGATGTGCCAGCAGACCTGCTGACCTACGACA
GGACAGATGAGGCAGACGCAGAGGAGAGGGGACCAGAGGAGAACTATTCTAGAC
CTGAGGCCCCAAATGAGTTCTACGACGGCGATCACGACAACGATAAGGAGAGCG
ATGTGGAGATC

**HDAC4**

HDAC4 Flag was a gift from Eric Verdin (Addgene plasmid # 13821 ; http://n2t.net/addgene:13821 ; RRID:Addgene_13821)

**HDAC5**

HDAC5 Flag was a gift from Eric Verdin (Addgene plasmid # 13822 ; http://n2t.net/addgene:13822 ; RRID:Addgene_13822)

**HDAC6**

HDAC6 Flag was a gift from Eric Verdin (Addgene plasmid # 13823 ; http://n2t.net/addgene:13823 ; RRID:Addgene_13823)

**HDAC7**

HDAC7 Flag was a gift from Eric Verdin (Addgene plasmid # 13824 ; http://n2t.net/addgene:13824 ; RRID:Addgene_13824)

**HDAC8**

ATGGAGGAGCCAGAGGAGCCTGCAGACTCTGGACAGAGCCTGGTGCCCGTGTAC
ATCTATTCTCCTGAGTACGTGAGCATGTGCGATTCCCTGGCCAAGATCCCAAAGAG
GGCCTCTATGGTGCACAGCCTGATCGAGGCCTATGCCCTGCACAAGCAGATGCGC
ATCGTGAAGCCCAAGGTGGCCTCCATGGAGGAGATGGCCACCTTCCACACAGACG
CCTACCTGCAGCACCTGCAGAAGGTGAGCCAGGAGGGCGACGATGACCACCCCG
ATTCCATCGAGTACGGCCTGGGCTATGACTGCCCTGCCACCGAGGGCATCTTTGAT
TATGCAGCAGCAATCGGAGGAGCAACCATCACAGCCGCCCAGTGCCTGATCGATG
GCATGTGCAAGGTGGCCATCAACTGGTCCGGAGGATGGCACCACGCCAAGAAGG
ACGAGGCCTCTGGCTTCTGTTACCTGAATGATGCCGTGCTGGGAATCCTGAGGCTG
AGGAGAAAGTTTGAGAGAATCCTGTATGTGGATCTGGACCTGCACCACGGCGATG
GAGTGGAGGACGCCTTCTCCTTTACCTCTAAAGTGATGACAGTGTCCCTGCACAA
GTTCTCTCCAGGCTTCTTTCCAGGAACAGGCGACGTGAGCGACGTGGGACTGGGC
AAGGGATGGTACTATAGCGTGAACGTGCCTATCCAGGATGGCATCCAGGACGAGA
AGTACTATCAGATCTGTGAGAGCGTGCTGAAGGAGGTGTACCAGGCCTTCAACCC
AAAGGCAGTGGTGCTGCAGCTGGGAGCAGATACCATCGCAGGCGACCCAATGTG
CTCCTTTAATATGACACCCGTGGGCATCGGCAAGTGTCTGAAGTATATCCTGCAGT
GGCAGCTGGCCACCCTGATCCTGGGAGGAGGAGGATACAACCTGGCCAATACAGC
CCGGTGCTGGACCTATCTGACAGGCGTGATCCTGGGCAAGACCCTGAGCTCCGAG
ATCCCTGACCACGAGTTCTTTACCGCCTACGGCCCAGATTATGTGCTGGAGATCAC
ACCTAGCTGTAGGCCAGACCGCAATGAGCCCCACCGGATCCAGCAGATCCTGAAC
TACATCAAGGGCAATCTGAAGCACGTGGTG

Inducible U-2 OS cell line was generated by transiently transfecting cells with pHDACx-Halo expression vector at ~70% confluence using Lipofectamine 2000 (Thermo Fisher Scientific, #11668027) following manufacturer's recommendations. Transfected cells were cultivated in DMEM (Thermo Fisher Scientific, #31053-028) supplemented with 10% FBS (Thermo Fisher Scientific, #10082139) and 1 μg/ml puromycin (Sigma Aldrich, #P9620) for 2 weeks until growth rate of the cells reached similar rate to untransfected cells in media without antibiotics. Thereafter, selected cells were frozen in 10% DMSO and stored at -80°C. Expression of transgene was induced using 0.1 μg/ml doxycycline (Sigma Aldrich, #D9891) for 48 - 72 h before imaging experiment.

*Maintenance and preparation of cells*

Human primary dermal fibroblasts (Lonza, #CC-2511) cells were cultured in high-glucose DMEM (Thermo Fisher, #31053044) with 10% FBS (Thermo Fisher, #10082147) supplemented with 1 mM Sodium pyruvate (Sigma, #S8636), 1% GlutaMax (Thermo Fisher, #35050038) and 1% Penicillin-Streptomycin (Sigma, #P0781) in a humidified 5% $CO_2$ incubator at 37 °C. The cells were split every 3-4 days or at confluence.

HeLa (ATCC, CCL-2) cells were cultured in high-glucose DMEM (Thermo Fisher, #31966047) with 10% FBS (BioSELL, #S0615) supplemented with 1% Penicillin-Streptomycin (Sigma, #P0781) in a humidified 5% $CO_2$ incubator at 37 °C. The cells were split every 3-4 days or at confluence.

SUP-M2 (DSMZ, #ACC-509) cells were cultured in RPMI 1640 basic medium (CLS, #820700a) with 10% FBS Superior (Bio&SELL, #S0615) supplemented with 1% Penicillin-

Streptomycin (Sigma, #P0781) in a humidified 5% $CO_2$ incubator at 37 °C. The cells were split every 3-4 days or at confluence.

Huh-7 (Cytion, # 300156) cells were cultured in RPMI 1640 basic medium (CLS, #820700a) with 10% FBS Superior (Bio&SELL, #S0615) supplemented with 1% Penicillin-Streptomycin (Sigma, #P0781) in a humidified 5% $CO_2$ incubator at 37 °C. The cells were split every 3-4 days or at confluence.

HEK293T (DSMZ, #ACC-635) cells were cultured in high-glucose DMEM (Thermo Fisher, #31966047) with 10% FBS Superior (Bio&SELL, #S0615) supplemented with 1% Penicillin-Streptomycin (Sigma, #P0781) in a humidified 5% $CO_2$ incubator at 37 °C. The cells were split every 3-4 days or at confluence.

SH-SY5Y (DSMZ, #ACC-209) cells were cultured in DMEM/F-12 1:1 medium (Cytiva, #SH30271.01) with 10% FBS Superior (Bio&SELL, #S0615) supplemented with 1% Penicillin-Streptomycin (Sigma, #P0781) in a humidified 5% $CO_2$ incubator at 37 °C. The cells were split every 3-4 days or at confluence.

U2OS (ECACC, #92022711) cells were cultured in McCoy's medium (Thermo Fisher, #16600082) with 10% FBS Superior (Bio&SELL, #S0615) supplemented with 1% Penicillin-Streptomycin (Sigma, #P0781) and 1% Sodium Pyruvate (Sigma, #S8636) in a humidified 5% $CO_2$ incubator at 37 °C. The cells were split every 3-4 days or at confluence.

3T3 (ECACC, #85022108) cells were cultured in high-glucose DMEM (Thermo Fisher, #31053044) with 10% HI FBS (Thermo Fisher, #10082147) supplemented with 1% Penicillin-Streptomycin (Sigma, #P0781), 1% Sodium Pyruvate (Sigma, #S8636), and 1% GlutaMax (Thermo Fisher, #35050038) in a humidified 5% $CO_2$ incubator at 37 °C. The cells were split every 3-4 days or at confluence.

Primary Rat Hippocampus Neurons (Gibco, #A36513) were thawed according to the manufacturer's protocol and seeded at a density of $6.25 \times 10^4$ cells per well onto Poly-D-Lysine (>4.5µg/cm$^2$) coated Ibidi 8-well slides. The cells were cultured in Neurobasal Plus Medium (ThermoFisher, #A3582901) supplemented with 2% B27 Plus Supplement (ThermoFisher, #A3582801) and 0.5mM GlutaMax (ThermoFisher, #35050038). On the second day, 1% Penicillin-Streptomycin (Sigma, #P0781) was added, and on the fourth day, 25µM L-Glutamine was added. The neurons were maintained in a humidified 5% $CO_2$ incubator at 37 °C, with a partial medium change performed every 3-4 days.

*Determination of Total Protein Concentration*

The total protein concentration of samples was determined using the Bradford protein assay kit (Bio-Rad, #5000201). A standard curve was generated using serial dilutions of Bovine Serum Albumin (Sigma Aldrich, #A7030) prepared in the appropriate sample buffer to final concentrations ranging from 0 to 25 µg/mL. In a 96-well microplate, 5 µL of each standard or unknown protein sample was added to individual wells in triplicate. Subsequently, 200 µL of the Bradford dye reagent was added to each well, and the plate was incubated for 10 minutes at room temperature, protected from light. The absorbance was measured at 595 nm using a microplate spectrophotometer (Tecan Spark 20M). After subtracting the average absorbance of the blank (0 µg/mL standard) from all readings, the concentrations of the unknown samples were calculated by interpolating their absorbance values against the linear regression of the BSA standard curve.

##### Spectroscopic Analysis

All absorption and fluorescence spectra were acquired at room temperature (25°C) using a Spark® 20M multiwell plate reader (Tecan) in 96-well glass bottom plates.

Absorption spectra were recorded from 320 nm to 850 nm with a 1 nm wavelength step size. A background spectrum, measured in wells containing only the experimental buffer and an equivalent concentration of DMSO, was subtracted from all sample spectra.

Fluorescence emission spectra were recorded with an emission bandwidth of 5 nm and a step size of 2 nm. The specific excitation parameters were as follows:

- For tetramethylrhodamine (TMR)-based probes, samples were excited at 495 nm (15 nm bandwidth), and emission was collected from 520 nm to 850 nm.

- For silicon-rhodamine (SiR)-based probes, samples were excited at 595 nm (15 nm bandwidth), with emission collected from 620 nm to 850 nm.

All samples were prepared and measured in technical triplicate. The entire experiment was repeated three times independently on different days to ensure reproducibility.

##### Lifetime measurement

The fluorescence decay characteristics of the solution samples in PBS buffer (Lonza™, 17517Q) at concentrations ranging from $10^{-6}$ to $10^{-7}$ M were recorded using a fluorescence lifetime measurement system (Quantaurus-Tau, Hamamatsu Photonics) in 3 mL high performance quartz glass cuvettes (Hellma Analytics Art. No. 101-10-K-40). The decay profile was registered for 53 ns interval after excitation and the experiment was continued until 10 000 peak count was reached. The instrument response function was obtained by using diluted LUDOX® TM-50 colloidal silica (Sigma Aldrich, #420778). The analysis of the obtained fluorescence decay profile was performed using the instrument software. Reported values are averages (n = 3) with standard deviation.

##### Quantum Yield Measurement

All reported absolute fluorescence quantum yield values (Φ) were measured using a Quantaurus-QY spectrometer (Hamamatsu Photonics, #C11374-01). This instrument uses an integrating sphere to determine photons absorbed and emitted by a sample. Measurements were carried out using dilute samples in air saturated solvents at 25°C at concentrations ranging from $10^{-6}$ to $10^{-7}$ M (A < 0.1 as indicated in the user's manual) and by using 3 mL quartz cuvettes (Hamamatsu Photonics Art. #A10095-02) provided by the instrument supplier. The fluorescence quantum yields were measured in PBS buffer (Lonza™, 17517Q). Self-absorption corrections, if needed, were performed using the instrument software. Reported values are averages (n = 3) with standard deviation.

**HTRF Saturation Binding Assay**

Cell lysates were prepared by incubating cells with Cellytic™ M lysis buffer (Sigma Aldrich, #C2978) containing a protease inhibitor cocktail (Sigma Aldrich, #11873580001) for 15 minutes at room temperature. The concentration of HaloTag-HDAC protein in the lysate was quantified by SDS-PAGE and subsequent fluorescence scanning, using a standard curve generated from purified JF646-Halo protein. Subsequently, the HaloTag-HDAC protein in the lysate was covalently labeled with the HTRF donor, Halo-Lumi4 (a $Tb^{3+}$ cryptate), by incubation for 1 hour at 37°C.

The labeled lysate was diluted in Tag-lite buffer (Cisbio, #LABMED) and dispensed into half-area, black polystyrene 96-well plates (Sigma Aldrich, #CLS3693) at 20 µL per well. A serial dilution of a fluorescent probe in 20 µL of Tag-lite buffer was added per well. Nonspecific binding was determined in parallel wells containing the fluorescent probe and a high concentration (10 µM) of the competitor **Nexturastat A**. All dilutions were prepared from DMSO stock solutions, ensuring a constant final DMSO concentration across all wells.

After a 1-hour incubation at room temperature, the Homogeneous Time-Resolved Fluorescence (HTRF) signal was measured on a Spark 20M multiwell plate reader (Tecan). The $Tb^{3+}$ donor was excited at 340 nm, and emission was recorded at 620 nm (donor) and 665 nm (acceptor, FRET signal). The HTRF ratio for each well was calculated by dividing the acceptor signal by the donor signal after background subtraction, according to the equation (1):

$$Ratio = \frac{\text{Signal 665 nm} - \text{Background Signal}}{\text{Signal 620 nm} - \text{Background Signal}} \quad (1)$$

Luminescence background signal was measured in wells containing 40 µl of Tag-lite buffer only. Specific binding was calculated by subtracting nonspecific binding ratio from total binding ratio at each fluorescent ligand concentration. Kd app is determined by fitting data using GraphPad Prism 10 to one site specific binding with Hill slope equation (2):

$$Specific\ signal\ ratio = \frac{\text{A}*[\text{probe}]^{\text{h}}}{(K_d^{app})^{\text{h}} + [\text{probe}]^{\text{h}}} \qquad (2)$$

A is amplitude of specific signal ratio, [probe] - probe concentration, $K_d^{app}$ - the ligand concentration needed to achieve a half-maximum binding at equilibrium, h - the Hill slope.

**Cell Cycle Profile Investigation**

HeLa cells were grown in 6-well plates (250.000 cells per well) for 24 h in the presence of the fluorescent probe in variable concentrations. The probes were dissolved in DMSO, thus control samples were prepared by adding only DMSO. We found that HeLa cells do not adhere strongly to the plastic bottom of the 6-well plate and thus the trypsinisation step could be omitted. The cells were simply washed off and suspended in 1 ml of the growth medium by intensively pipetting up and down. Next, the cells were processed according to the NucleoCounter® NC-3000™ two-step cell cycle analysis protocol. Specifically, ~500.000 cells were harvested by centrifuging at room temperature for 5 min at 400g. Afterwards, the cells were resuspended in 250 µl lysis solution (Solution 10, Chemometec, #910-3010) supplemented with 10 µg ml−1 DAPI (Solution 12, Chemometec, #910-3012), incubated at 37 °C for 5 min. Then 250 µl of stabilization solution (Solution 11, Chemometec, #910-3011) was added. Cells were counted on a NucleoCounter® NC-3000™ in NC-Slide A2™ slides (Chemometec, Cat. no. 942-0001) loaded with ~30 µl of each of the cell suspensions into the chambers of the slide. Each time, ~10.000 cells in total were measured, and the obtained cell cycle histograms were analysed with ChemoMetec NucleoView NC-3000 software, version 2.1.25.8. All experiments were repeated three times and the results are presented as means with standard deviations. The obtained mean values were compared by running multiple t-tests on GraphPad Prism version 10.5 software.

**Annexin V Apoptosis Assay**

SUP-M2 cells were seeded in 6-well plates at a density of $2.5 \times 10^5$ cells per well. Cells were then treated for 24 hours with various concentrations of probe or with an equivalent volume of DMSO as a negative control. Following treatment, cells were harvested and counted using a NucleoCounter® with a Via1-Casette™ (Chemometec, #941-0012). For staining, cells were washed twice with PBS by centrifugation at 400 × g for 5 minutes. An aliquot of $2–4 \times 10^5$ cells was resuspended in 100 µL of 1X Annexin V binding buffer (Biotium, #99902). Cells were stained by adding 2 µL of Annexin V-CF488A conjugate (Biotium, #29005) and 2 µL of Hoechst 33342 stock solution (500 µg/mL) to achieve a final concentration of approximately 10 µg/mL. The staining reaction was performed in the dark at 37°C for 15 minutes. After staining, cells were washed twice with 300 µL of Annexin V binding buffer. Finally, the cell pellet was resuspended in 100 µL of Annexin V binding buffer supplemented with PI to a final concentration of 10 µg/mL. Samples were analyzed immediately on a NucleoCounter® NC-3000™ (Chemometec). Approximately 30 µL of each cell suspension was loaded into an NC-Slide A2™, and the pre-set "Annexin V Assay" protocol was executed. The resulting scatter plots and histograms of Annexin V-CF488A and PI fluorescence intensity were used to quantify the percentages of live, early apoptotic, and late apoptotic/necrotic cells.

**Fluorescence Activated Cell Sorting**

Cells were treated with indicated solution 1 hour at 37°C in a 5% $CO_2$. For adherent cells, detachment was achieved by incubation with Accutase (Gibco, #A1110501) for 2-4 minutes at 37°C. Following incubation, cells were washed twice with PBS, pelleting the cells by centrifugation at 250 × g for 2 minutes between washes. After the final wash, cells were resuspended in fresh PBS. Immediately prior to analysis, the cell suspension was passed through a 35 µm cell strainer cap (Falcon, #352235) to ensure a single-cell suspension. Samples were analyzed using a BD FACSAria™ III High-Speed Cell Sorter equipped with a 100 µm nozzle and three lasers (405 nm, 488 nm, and 633 nm). Data were acquired in FCS v. 3.1 format and subsequently analyzed using FlowJo™ software v. 10.5.3 (BD Life Sciences).

**Microscopy**

*Sample Preparation*

Cells were seeded in glass bottom 12-well (MatTek, #P12G-1.5-14-F) or 24-well (MatTek, #P24G-1.5-13-F) plates for wide-field imaging experiments. Confocal and STED microscopy experiments were performed using µ-Slide 8 Well Glass Bottom dishes (Ibidi, #80827).

*Live cell staining*

U2OS-HDACs-Halo cells were plated and induced with doxycline (Sigma Aldrich, #D9891) 24h before imaging. For staining, the cells were incubated for 2 h at 37 °C in DMEM containing 50 ng/mL Hoechst 33342, 1 µM screening probe and 100 nM accordingly Halotag substrate. The microscope was pre-heated to 37°C and 5% $CO_2$ was supplied. After the incubation, the cells were washed once with DMEM and once with Hank's Balanced Salt Solution (HBSS, Gibco, # 14025092). Fresh DMEM was added, and images were recorded.

*Immunofluorescence Staining*

PFA (paraformaldehyde) fixation

Cells were fixed with 4% (w/v) paraformaldehyde (Thermo Fisher Scientific, #28908) in PBS for 5 min. The solution was removed, and 1 ml of 30 mM glycine in PBS (pH 7.4) was added for 5 min. Following three 5-minute washes in PBS, permeabilisation was performed using 0.5% (v/v) Triton-X-100 (1.08603, Millipore) in PBS for 5 mins. After three 5-mines washing steps with PBS, Cell were then incubated overnight at 4°C with a primary antibody cocktail containing mouse HDAC6 Monoclonal Antibody (1:1000; Invitrogen, #MA5-25359) diluted in blocking buffer (1% BSA in PBS). The next day, cells were washed three times with PBS for 5 minutes each. They were then incubated for 1 hour at room temperature with a secondary antibody solution containing donkey anti-mouse IgG conjugated to Alexa Fluor 488 (1:1000; Invitrogen, #A32766). After secondary antibody incubation, cells were washed three times with PBS for 20 minutes each, with the final wash step performed overnight at 4°C.

Methanol fixation for tubulin

Cells were incubated with fresh medium containing the indicated concentrations of probe or Nexturastat A (Selleckchem, #S1030). Following a 24-hour incubation at 37°C, cells were rinsed once with phosphate-buffered saline (PBS) and fixed with cold methanol and containing 1 mM EGTA for 4 minutes. After fixation, cells were washed three times with PBS and blocked for 30 minutes at room temperature with 1% bovine serum albumin (BSA) in PBS. Samples were then incubated overnight at 4°C with a primary antibody cocktail containing mouse anti-acetylated tubulin (1:1000; Invitrogen, #32270-A546) and rabbit anti-β-tubulin (1:1000; Cell Signaling Technology, #2128) diluted in blocking buffer. The next day, samples were washed three times with PBS for 5 minutes each. They were then incubated for 1 hour at room temperature in the dark with a secondary antibody solution containing donkey anti-rabbit IgG conjugated to Alexa Fluor 488 (1:1000; Invitrogen, #A-21206) and donkey anti-mouse IgG conjugated to Alexa Fluor 568 (1:1000; Invitrogen, #A10037). After secondary antibody incubation, cells were washed three times with PBS for 5 minutes each, with the final washing step performed overnight at 4°C.

*Microscope*

Wide field microscope: For wide-field fluorescence microscopy images were acquired using Biotek Lionheart FX Automated Microscope equipped with Olympus dry 20× NA 0.45 objective and incubator set to +37°C. During acquisition cells were kept under atmosphere containing 5% of $CO_2$. The follosing settings were used for imaging:

| **Channel** | **LED** | **Filter cube** | **LED intensity (a.u.)** | **Integration time (ms)** | **Camera gain** |
|---|---|---|---|---|---|
| Hoechst | 365 nm | DAPI 377/447 | 4 | 10 | 27 |
| RFP channel | 523 nm | RFP | 5 | 25 | 30 |
| Cy5 channel | 623 nm | Cy5 | 7 | 50 | 30 |

Confocal/STED microscopes: Images were acquired using Abberior STED Facility Line scanning (Abberior Instruments GmbH). Abberior STED Facility Line is equipped with 488, 515, 561, 640 and 700 nm 40 MHz pulsed excitation lasers, a pulsed 775 nm 40 MHz 3W STED laser, and an UPlanSApo 60x/1.40 Oil objective. For all measurements pinhole was set to 1AU. Microscope has two APD and two MATRIX detectors which can be tuned to any detection window in the range 400 – 800 nm. Pixel size was 30 nm in the xy plane was used for 2D STED images and 80 nm in the xy plane for large field of view images. Laser powers were optimized for each sample. Detailed image acquisition settings are listed in **Table S2**.

Spining disk confocal microscopy imaging was performed on an inverted microscope (Nikon Ti2) equipped with the following excitation lasers: 405nm (150 mW),488nm (150 mW), 561nm (150 mW), 640nm (150 mW) in confocal, 640nm (200 mW) in TIRF mode and the oil-immersion objective Plan Apo Lambda 60x Oil NA 1.4 (MRD01605, Nikon). All measurements utilized disk with 50 µm pinholes. Images registered using back illuminated sCMOS camera Teledyne Photometrics Prime BSI with pixel size 6.5 x 6.5 µm corresponding to image pixel size 111 x 111 nm. Detailed image acquisition settings are listed in **Table S2**.

Multi-color time-lapse movie: For recording multicolor movies, the cells were incubated with the indicated probes for 1h at 37°C, washed twice with HBSS, then applied DMEM for imaging. The time-lapse series were recorded on the spining disk confocal microscope. 5 field of view were acquired as z-stacks with 500 nm step size for every time point. 200 frames win 10 sec interval were acquired for each movie and the final videos were rendered at 10 fps. Detailed image acquisition settings are listed in **Table S2**.

*Image Processing and Analysis*

All acquired or reconstructed images were processed and visualized using Fiji. Stitching and deconvolution of images, Pearson correlation coefficient analysis were performed using SVI Huygens Essential software package.

*Image Analysis Pipeline for Cytoplasmic Signal Quantification*

This image analysis workflow was designed using **CellProfiler** software (version 4.2.6) to automatically segment nuclei, identify the cytoplasm, and quantify the intensity of a HDAC6 fluorescent probe within this specific subcellular compartment [1].

*Image Analysis Pipeline for Tubulin Filament Quantification*

This image analysis workflow was developed in **CellProfiler** (version 4.2.6) to segment and quantify the signal from fluorescently-labeled tubulin filaments in microscopy images. The pipeline employs a threshold-based method to identify the filamentous structures and subsequently measures their total intensity.

#### Chemical synthesis

*General procedures*

Reagents were purchased as reagent-grade (from Sigma-Aldrich, Thermo Fisher, BLD Pharm, abcr, TCI, Aaron Chem and Selleck Chem) and used without further purification. 4SiR-COOH, 5SiR-COOH, 6SiR-COOH, 4TMR-COOH were synthesized according to published procedure[2]. Flash chromatographies were carried out on Biotage® Selekt with Biotage® Sfär Silica HC (20 µm) columns (Biotage, #FSUS-0443).

NMR spectra were recorded at 25 °C with an Agilent 400-MR spectrometer at 400 MHz ($^1H$) and 101 MHz ($^{13}C$). Each sample was measured once (n=1). Chemical shifts (δ), which are expressed in part per million (ppm), were determined relative to residual non-deuterated solvent as an internal reference: $CDCl_3$ ($^{13}C$ NMR: δ = 77.16 ppm; $^1H$ NMR: δ = 7.26 ppm) and $(CD_3)_2SO$ ($^{13}C$ NMR: δ = 39.52 ppm; $^1H$ NMR: δ = 2.50 ppm) or with an external reference ($^{19}F$: δ = 0 ppm for $CFCl_3$). Multiplicities of signals are described as follows: s = singlet, d = doublet, t = triplet, q = quartet, m = multiplet or overlap of non-equivalent resonances; br = broad signal. Coupling constants (*J*) are given in Hz.

ESI-MS were recorded on a Varian 500-MS spectrometer (Agilent). ESI-HRMS were recorded on a MICROTOF spectrometer (Bruker) equipped with ESI ion source (Apollo) and direct injector with LC autosampler Agilent RR 1200. Each sample was measured once (n=1).

Analytical LC–MS analysis was performed on an Agilent 1260 Infinity II LC/MS system equipped with an Autosampler (G7129A), Binary Pump (G7112B), Diode Array Detector WR (G7115A), and Single Quadrupole MSD XT (G6135B). Analysis was done by using a Ascentis® Express ES-Cyano (Supelco, #577307), 2 µm, 50 × 2.1 mm, 5 cm x 2.1 mm with Mobile phase A: 25 mM $HCOONH_4$ (pH = 3.5) aqueous buffer/ Mobile Phase B: MeOH.

Preparative HPLC was performed on a combined Agilent1260/1290 Infinity II preparative system equipped with a 1290 Infinity II open-bed sampler (G7169B)/fraction collector (G7159B), 1260 Infinity II multiple wavelength detector (G7165A) and with:

- Device A: 1260 Infinity II preparative binary pump (G7161A) and Agilent 5 Prep-$C_{18}$, 5 µm, 100 x 50 mm preparative column.
- Device B: 1290 Infinity II preparative binary pump (G7161B) and Agilent Pursuit 10 $C_{18}$, 10 µm, 250 X 50 mm preparative column.

Reaction yield determination by absorption spectroscopy: After the purification and removal of the solvent, the obtained fluorescent probes were dissolved in a precise (700 µL) volume of the *d6*-DMSO (Deutero, #00908) solvent and were transferred to NMR tube (Aldrich, # Z569364) to obtain $^1H$ spectra. Afterwards the contents of the NMR tubes were transferred to an eppendorf and was considered as stock solution. Five 2 µL samples were taken from stock solution and were diluted in five separate eppendorf's with 98 µL of PBS + 0.1% SDS (50-fold dilution), vigorously mixed and aged for 30 min to dissolve aggregates. Then absorption of 2 µL of the diluted samples were measured on nanodrop (Nanodrop 1000, Peqlab) with 1 mm optical path. The measured absorption intensity values at the dyes absorption maxima value were averaged and concentration of the stock solution was determined according to the equation (3):

$$C=(dilution*A)/(\varepsilon*l) \quad (3)$$

where C –concentration of stock solution; A – sample absorption, ε - extinction coefficient of the dye in PBS containing 0.1% SDS, l – path length

Once the concentration of the stock solution is measured the mass of the obtained fluorescent conjugate could be calculated by following equation (4):

$$m=C*MW*V \quad (4)$$

where C –concentration of stock solution; MW – molecular weight of the compound; V-volume of stock solution.

Finally, the yield of the synthesis step could be determined by the classical equation (5):

$$Yield\ \% = m/m_{teor} * 100 \quad (5)$$

Where m – obtained mass of the isolated product; $m_{teor}$ – maximal theoretical mass of the product in the reaction.

<u>ClogP</u> was estimated using Chemdraw Professional 15.0

*Chemical synthesis and charachterization of the probes*

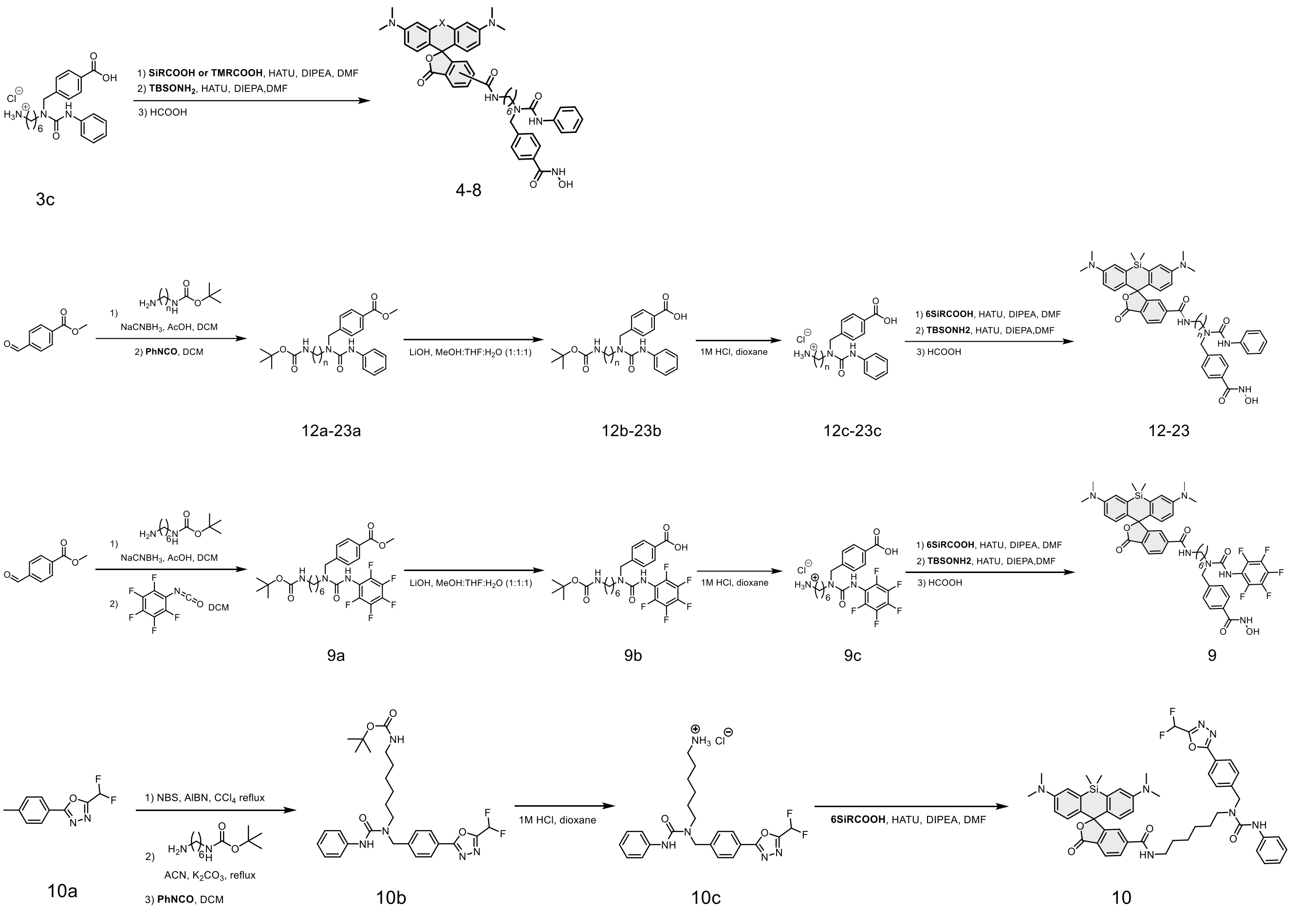

**General procedure A** for the compounds **3a, 9a, 12a-23a**:

A round-bottom flask charged with methyl 4-formylbenzoate (1 equiv) and corresponding amine (1.1 equiv) was taken up in a solution of 5% AcOH in DCM. After 5 min, $NaCNBH_3$ (2 equiv) was added in portions and the resulting mixture was stirred at room temperature under an atmosphere of argon overnight. The reaction was quenched with 1 N NaOH (10 mL) and the aqueous layer extracted with Ethyl acetate (3 × 20 mL). The combined organic extracts were washed with brine, dried over sodium sulfate, concentrated in vacuo and dissolved in DCM (10 mL). To this solution, was added the appropriate isocyanate (1.2 equiv) at room temperature under an atmosphere of argon, and the resulting solution was stirred overnight.

The reaction was quenched with saturated sodium bicarbonate (10 mL) and extracted with DCM (3 × 10 mL). The combined organics were washed with brine (15 mL), dried over sodium sulfate, concentrated in vacuo, and purified via flash chromatography, affording the product as a colorless liquid.

**General procedure B** for the compounds **3b, 9b, 12b-23b**:

Compound from previous step was dissolved in solvent combination of MeOH: THF: $H_2O$ (1:1:1). To this solution, LiOH (2 equiv) was added, and the resulting mixture was stirred at room temperature overnight. The reaction was acidified with 1 N HCl and the aqueous layer extracted with Ethyl acetate (3 x 10mL). The combined organic extracts were washed with brine, dried over sodium sulfate, concentrated in vacuo and use for the next step without purification.

**General procedure C** for the probe **1-9, 12-23**:

Compound from previous step was dissolved in dioxane and 5 eq of HCL 4 N in dioxane was added to this solution. The resulting solution was stirred at room temperature and monitor by TLC until the all starting material was consumed. The reaction was then concentrated and lyophilized to afford white solid compound. This product was then used for the next step without purification.

2 mg of the dye was dissolved in 100 uL of dry DMF in an eppendorf and 5 µL of DIPEA was added. To this solution 1.05 equiv of HATU solution in 100 µL of dry DMF was added. The eppendorf was shaked by a microspin at room temperature for 1 min and the resulting solution was transferred to another eppendorf containing 1.5 eq of corresponding amine from Procedure C and 10 µL of DIPEA in 100 µL of dry DMF and the resulting solution was sonicated for 5 mins. The obtained products were purified by preparative HPLC (preparative column: Agilent 5 Prep-C18, 5 µm, 100 x 50 mm flow rate: 40 mL/min, solvent A: MeCN, solvent B: $H_2O$ + 0.1% HCOOH; temperature 25°C, gradient A:B - 2 min 20:80 isocratic, 2-20 min 20:80 to 100:0 gradient and 20-24min 100:0 isocratic). The product fraction was concentrated in a 20 mL vial using Biotage V-10 touch evaporator (Biotage, # V10-2SC).

The resulting compound was then dissolved in 0.5 mL of dry DMF, following by adding 20 µL of DIPEA, 2 equiv of O-(tert-Butyldimethylsilyl)hydroxylamine and handly mixed. 1.5 equiv of HATU in 100 µL of dry DMF was then added. The vial was then shaked handily for 30 seconds and allow to stand for 5 mins. After that, 1mL of TFA was added to the reaction vial. The vial was shaked handily and the reaction was monitored by LCMS. Final product was purified by preparative HPLC (preparative column: Agilent 5 Prep-C18, 5 µm, 100 x 50 mm flow rate: 40 mL/min, solvent A: MeCN, solvent B: $H_2O$ + 0.1% HCOOH; temperature 25°C, gradient A:B - 2 min 20:80 isocratic, 2-20 min 20:80 to 100:0 gradient and 20-24min 100:0 isocratic). The HPLC fractions of product were then lyophilized.

Compound **1a**:

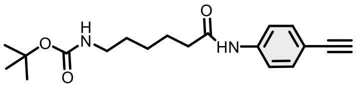

To a solution of 4-Ethynylaniline (2g, 17 mmol) and 6-(Boc-amino)caproic acid (4.7g, 20.4 mmol) in DMF (20 mL) and DIPEA (50 mmol) was added HATU (20 mmol) then the mixture was stirred at room temperature overnight. Saturated ammonium chloride solution was added to the solution and extracted with DCM. The combined organic layers were washed with brine and dried over $Na_2SO_4$ and concentrated in vacuo. The residue was purified by column chromatography to afford 2a (4.6g, 82%).

$^1$H NMR (400 MHz, $CDCl_3$) δ 8.63 (s, 1H), 7.55 – 7.44 (m, 2H), 7.42 – 7.28 (m, 2H), 4.85 (t, $J$ = 5.9 Hz, 1H), 3.12 – 2.94 (m, 3H), 2.28 (t, $J$ = 7.5 Hz, 2H), 1.62 (p, $J$ = 7.6 Hz, 2H), 1.39 (s, 11H), 1.25 (tt, $J$ = 9.5, 5.8 Hz, 2H). $^{13}$C NMR (101 MHz, $CDCl_3$) δ 172.01, 156.27, 138.89, 132.70, 119.57, 117.23, 83.48, 79.14, 40.32, 37.20, 29.73, 28.42, 26.30, 25.08. HRMS (ESI) $m/z$ calculated 331.2021 for $C_{19}H_{27}N_2O_3$ [M+H]$^+$, found 331.2016.

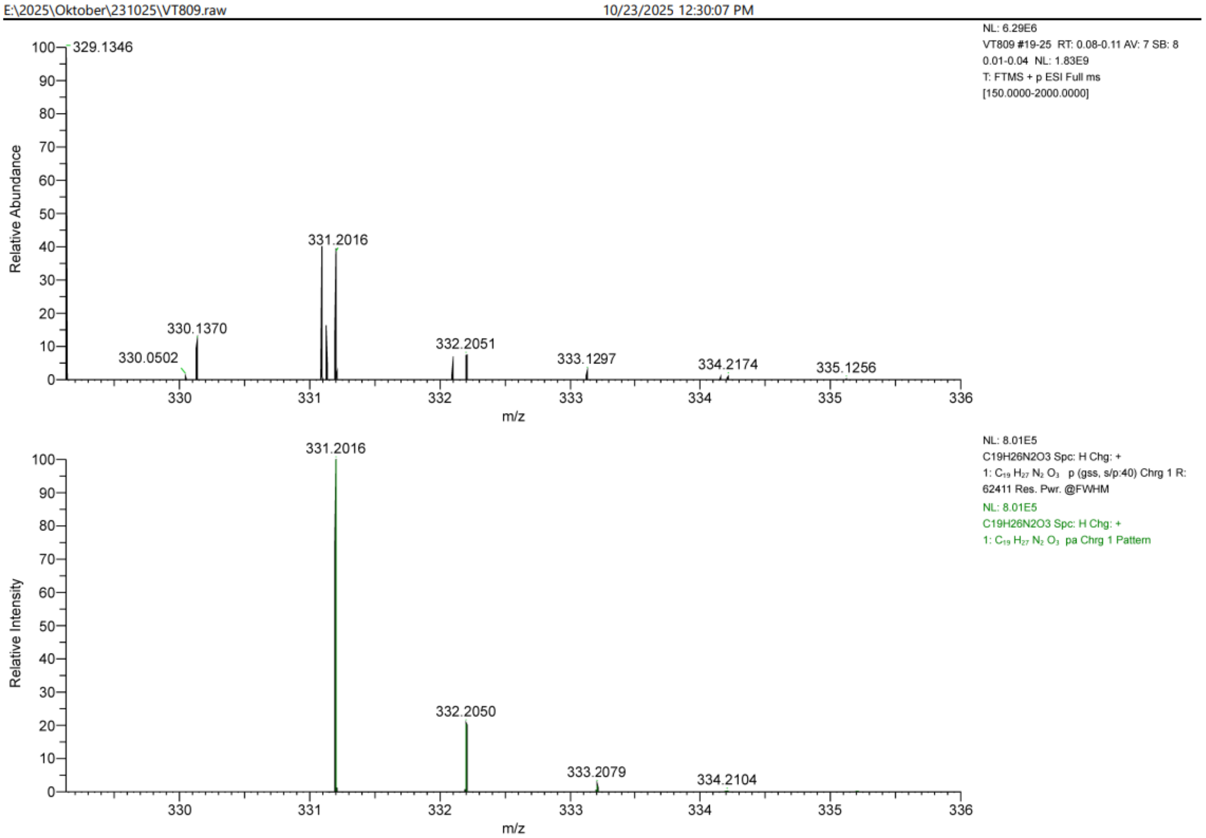

Compound **2a**:

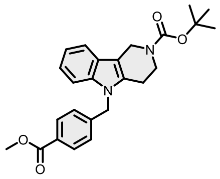

Compound **2a** was synthesized according to reported procedure[3].

Compound **3a:**

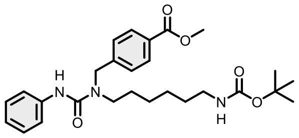

Purified by flash column chromatography (Biotage® Sfär Silica HC 25 g, gradient 2% to 50% hexane – EtOAc). Obtained 1073 mg of target compound in 73% yield as a colorless liquid by general method A.

$^{1}$H NMR (400 MHz, $CDCl_3$) δ 8.12 – 7.94 (m, 2H), 7.43 – 7.23 (m, 6H), 7.04 (t, J = 6.9 Hz, 1H), 6.53 (s, 1H), 4.65 (s, 2H), 3.94 (d, J = 2.5 Hz, 3H), 3.49 – 3.26 (m, 2H), 3.11 (t, J = 6.6 Hz, 2H), 1.74 – 1.56 (m, 2H), 1.46 (d, J = 2.4 Hz, 11H), 1.35 (p, J = 3.5 Hz, 4H). $^{13}$C NMR (101 MHz, $CDCl_3$) δ 166.79, 156.08, 155.38, 143.11, 138.96, 130.19, 129.53, 128.85, 127.07, 123.16, 120.04, 79.17, 52.19, 50.56, 47.90, 40.50, 29.97, 28.45, 28.31, 26.57, 26.43. HRMS (ESI) *m*/*z* calculated 484.2811 for $C_{27}H_{38}N_3O_5$ $[M+H]^+$, found 484.2819.

#### Display Report

**Analysis Info**

| | | | |
|---|---|---|---|
| Analysis Name | Z:\Data\2024\2408\sam280824\VTE16_14_01_127262.d | Acquisition Date | 28.08.2024 20:21:35 |
| Method | hystar_pl.m | Operator | BDAL@DE |
| Sample Name | VTE16 | Instrument / Ser# | micrOTOF 10237 |
| Comment | | | |

**Acquisition Parameter**

| | | | | | |
|---|---|---|---|---|---|
| Source Type | ESI | Ion Polarity | Positive | Set Nebulizer | 1.2 Bar |
| Focus | Not active | | | Set Dry Heater | 180 °C |
| Scan Begin | 50 m/z | Set Capillary | 4500 V | Set Dry Gas | 4.0 l/min |
| Scan End | 1600 m/z | Set End Plate Offset | -500 V | Set Divert Valve | Source |

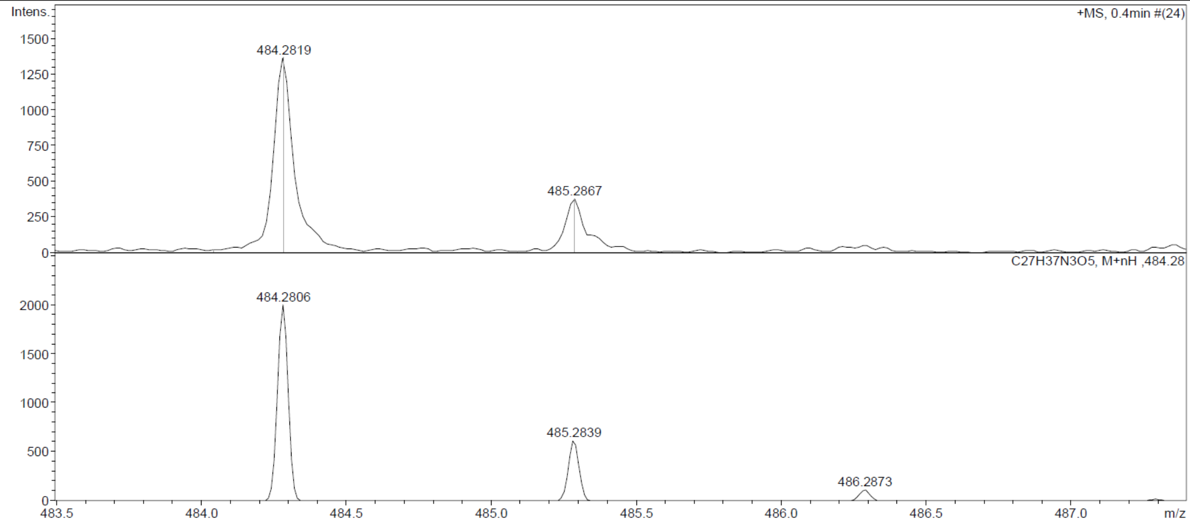

Compound **9a:**

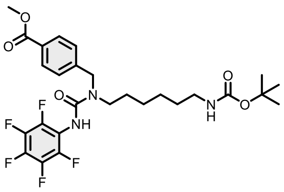

Purified by flash column chromatography (Biotage® Sfär Silica HC 25 g, gradient 2% to 50% hexane – EtOAc). Obtained 1101 mg of target compound in 63% yield as a colorless liquid by general method A.

$^{1}$H NMR (400 MHz, $CDCl_3$) δ 8.07 – 7.95 (m, 2H), 7.32 (d, $J$ = 8.4 Hz, 2H), 6.34 (s, 1H), 4.61 (s, 2H), 3.90 (s, 3H), 3.33 (t, $J$ = 7.6 Hz, 2H), 3.04 (q, $J$ = 6.2 Hz, 2H), 1.61 (s, 2H), 1.40 (s, 11H), 1.30 (h, $J$ = 4.6 Hz, 4H). $^{13}$C NMR (101 MHz, $CDCl_3$) δ 166.72, 156.07, 154.76, 144.28, 142.36, 141.84, 130.16, 129.59, 126.91, 113.98, 113.83, 79.10, 52.14, 50.58, 47.91, 40.25, 29.84, 28.33, 27.94, 26.33, 26.26. HRMS (ESI) *m/z* calculated 574.2340 for $C_{27}H_{33}F_5N_3O_5$ $[M+H]^+$, found 574.2348.

#### Display Report

**Analysis Info**

| | | | |
|---|---|---|---|
| Analysis Name | Z:\Data\2024\2409\sam040924\VTE4_1_01_127405.d | Acquisition Date | 04.09.2024 17:53:03 |
| Method | hystar_pl.m | Operator | BDAL@DE |
| Sample Name | VTE4 | Instrument / Ser# | micrOTOF 10237 |
| Comment | | | |

**Acquisition Parameter**

| | | | | | |
|---|---|---|---|---|---|
| Source Type | ESI | Ion Polarity | Positive | Set Nebulizer | 1.2 Bar |
| Focus | Not active | | | Set Dry Heater | 180 °C |
| Scan Begin | 50 m/z | Set Capillary | 4500 V | Set Dry Gas | 4.0 l/min |
| Scan End | 1600 m/z | Set End Plate Offset | -500 V | Set Divert Valve | Source |

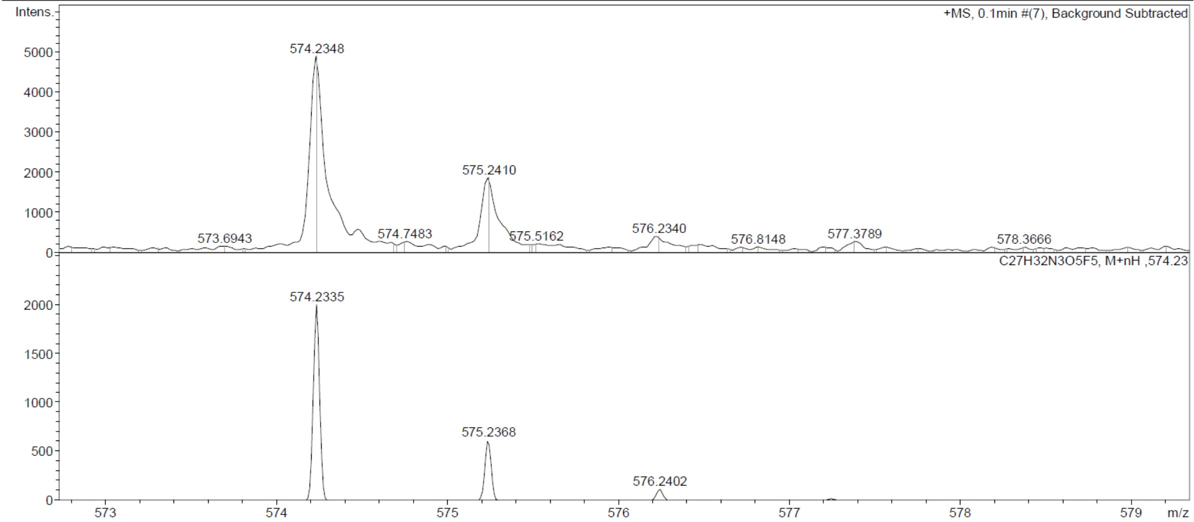

Compound **10a**:

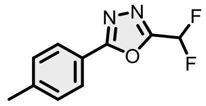

Compound **10a** was synthesized according to reported procedure[4].

Compound **11a**:

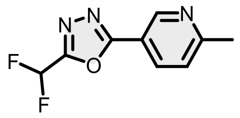

6-Methyl-3-pyridinecarboxylic acid hydrazide (2g, 13.23 mmol), 2,2-difluoroacetic anhydride (40 mmol), and Burgess reagent (40 mmol) were combined in 20 mL THF at rt. The resulting mixture was refluxed under an atmosphere of nitrogen overnight. The reaction mixture was concentrated under reduced pressure and partitioned between EtOAc and water. The organic solution was washed consecutively with water and brine, before drying over $Na_2SO_4$. The organic phased was concentrated to dryness under reduced pressure; and purified by column chromatography to afford **11a** (1.98g, 71%).

$^1$H NMR (400 MHz, $CDCl_3$) δ 9.15 (dd, *J* = 2.3, 0.8 Hz, 1H), 8.23 (dd, *J* = 8.2, 2.3 Hz, 1H), 7.33 (ddq, *J* = 8.2, 1.0, 0.5 Hz, 1H), 6.92 (t, *J* = 51.7 Hz, 1H), 2.63 (d, *J* = 0.5 Hz, 3H). $^{13}$C NMR (101 MHz, $CDCl_3$) δ 164.38, 163.18, 158.33, 134.90, 116.51, 108.11, 105.71, 103.32, 24.69. HRMS (ESI) *m/z* calculated 212.0630 for $C_9H_8F_2N_3O$, found 212.0631 ($[M+H]^+$).

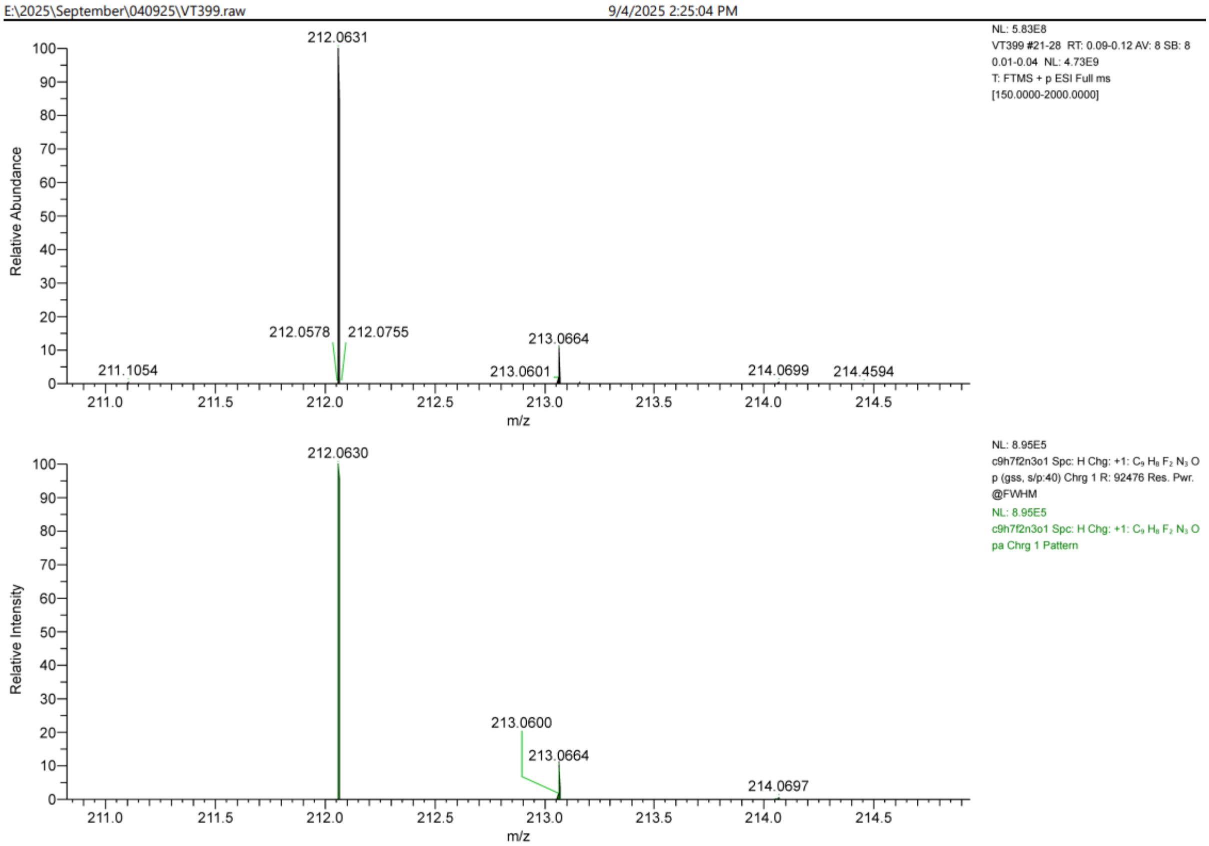

Compound **12a:**

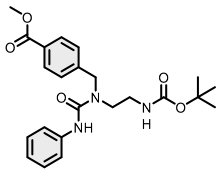

Purified by flash column chromatography (Biotage® Sfär Silica HC 25 g, gradient 2% to 50% hexane – EtOAc). Obtained 1066 mg of target compound in 82% yield as a colorless liquid by general method A.

$^{1}$H NMR (400 MHz, $CDCl_3$) δ 8.30 (s, 1H), 8.08 – 7.98 (m, 2H), 7.66 (d, $J$ = 8.0 Hz, 2H), 7.42 – 7.36 (m, 2H), 7.35 – 7.28 (m, 2H), 7.05 (tt, $J$ = 7.3, 1.1 Hz, 1H), 4.67 (s, 2H), 3.94 (s, 3H), 3.43 (dd, $J$ = 8.7, 6.3 Hz, 2H), 3.15 (dd, $J$ = 8.8, 6.1 Hz, 2H), 1.48 (s, 9H). $^{13}$C NMR (101 MHz, $CDCl_3$) δ 166.87, 157.12, 155.78, 143.79, 139.94, 130.05, 129.40, 128.72, 127.56, 122.64, 119.70, 80.48, 52.16, 50.75, 46.26, 39.80, 28.38. HRMS (ESI) *m*/*z* calculated 428.2185 for $C_{23}H_{30}N_3O_5$ [M+H]$^+$, found 428.2193.

#### Display Report

**Analysis Info**

| | | | |
|---|---|---|---|
| Analysis Name | Z:\Data\2024\2408\sam280824\VTE15_13_01_127261.d | Acquisition Date | 28.08.2024 20:16:58 |
| Method | hystar_pl.m | Operator | BDAL@DE |
| Sample Name | VTE15 | Instrument / Ser# | micrOTOF 10237 |
| Comment | | | |

**Acquisition Parameter**

| | | | | | |
|---|---|---|---|---|---|
| Source Type | ESI | Ion Polarity | Positive | Set Nebulizer | 1.2 Bar |
| Focus | Not active | | | Set Dry Heater | 180 °C |
| Scan Begin | 50 m/z | Set Capillary | 4500 V | Set Dry Gas | 4.0 l/min |
| Scan End | 1600 m/z | Set End Plate Offset | -500 V | Set Divert Valve | Source |

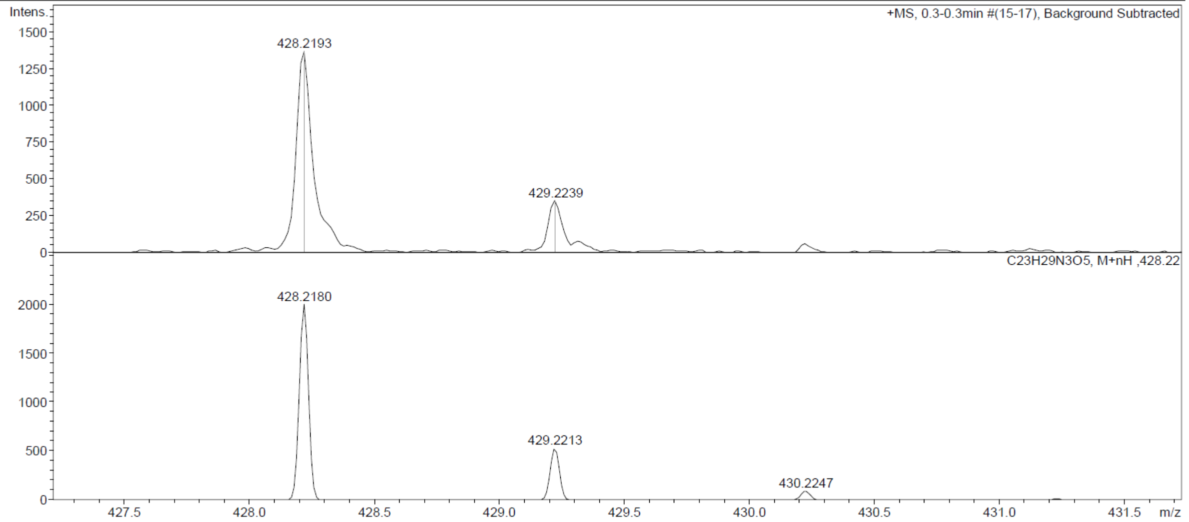

Compound **13a:**

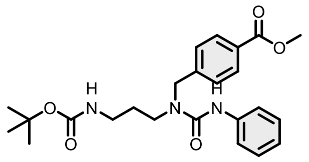

Purified by flash column chromatography (Biotage® Sfär Silica HC 25 g, gradient 2% to 50% hexane – EtOAc). Obtained 1020 mg of target compound in 76% yield as a colorless liquid by general method A.

$^{1}$H NMR (400 MHz, $CDCl_3$) δ 8.12 – 8.00 (m, 2H), 7.42 – 7.34 (m, 4H), 7.32 – 7.24 (m, 2H), 7.09 – 7.01 (m, 1H), 6.94 (s, 1H), 4.64 (s, 2H), 3.95 (s, 3H), 3.49 (t, $J$ = 6.8 Hz, 2H), 3.24 – 3.12 (m, 2H), 1.85 – 1.73 (m, 2H), 1.47 (s, 9H). $^{13}$C NMR (101 MHz, $CDCl_3$) δ 166.75, 156.52, 155.98, 142.77, 139.04, 130.28, 129.63, 128.79, 126.92, 123.19, 120.29, 79.54, 52.21, 50.27, 45.15, 37.73, 28.91, 28.44. HRMS (ESI) $m/z$ calculated 442.2341 for $C_{24}H_{32}N_3O_5$ $[M+H]^+$, found 442.2330.

Display Report

**Analysis Info**

| | | | |
|---|---|---|---|
| Analysis Name | Z:\Data\2024\2408\sam280824\VTE20_15_01_127263.d | Acquisition Date | 28.08.2024 20:26:13 |
| Method | hystar_pl.m | Operator | BDAL@DE |
| Sample Name | VTE20 | Instrument / Ser# | micrOTOF 10237 |
| Comment | | | |

**Acquisition Parameter**

| | | | | | |
|---|---|---|---|---|---|
| Source Type | ESI | Ion Polarity | Positive | Set Nebulizer | 1.2 Bar |
| Focus | Not active | | | Set Dry Heater | 180 °C |
| Scan Begin | 50 m/z | Set Capillary | 4500 V | Set Dry Gas | 4.0 l/min |
| Scan End | 1600 m/z | Set End Plate Offset | -500 V | Set Divert Valve | Source |

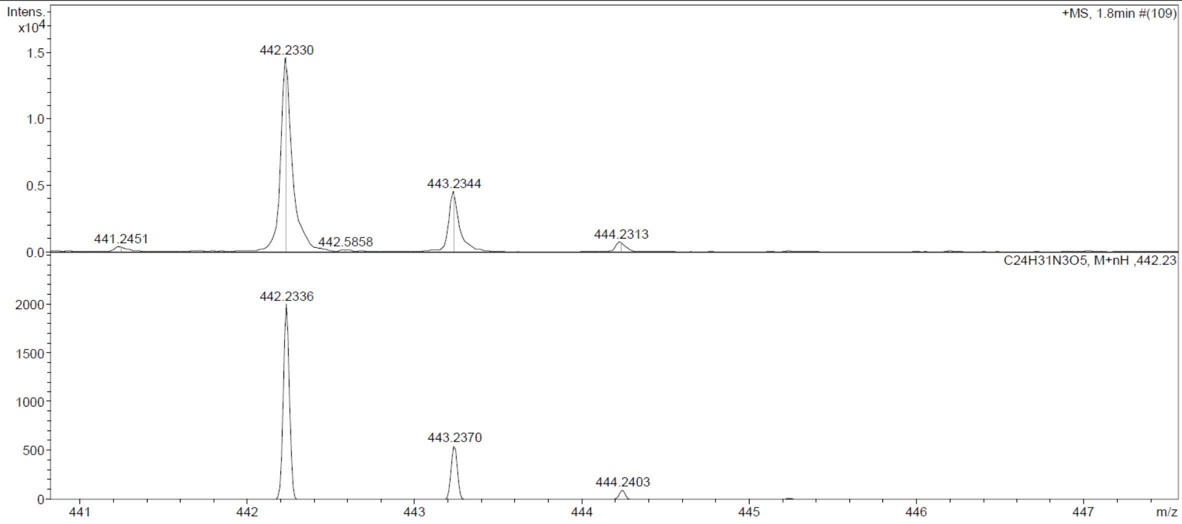

Compound **14a:**

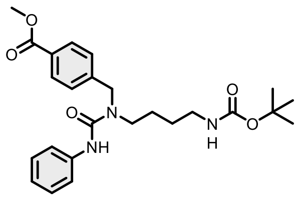

Purified by flash column chromatography (Biotage® Sfär Silica HC 25 g, gradient 2% to 50% hexane – EtOAc). Obtained 1150 mg of target compound in 83% yield as a colorless liquid by general method A.

$^{1}$H NMR (400 MHz, $CDCl_3$) δ 8.07 – 7.95 (m, 2H), 7.47 (d, $J$ = 7.9 Hz, 2H), 7.41 – 7.33 (m, 2H), 7.30 – 7.21 (m, 3H), 7.07 – 6.96 (m, 1H), 4.64 (s, 2H), 3.92 (s, 3H), 3.45 – 3.29 (m, 2H), 3.15 (t, $J$ = 6.4 Hz, 2H), 2.06 (s, 1H), 1.71 – 1.61 (m, 2H), 1.49 (q, $J$ = 6.6 Hz, 2H), 1.41 (s, 9H). $^{13}$C NMR (101 MHz, $CDCl_3$) δ 166.87, 156.67, 155.68, 143.55, 139.36, 130.04, 129.29, 128.63, 127.19, 122.96, 120.54, 79.42, 60.41, 52.13, 50.00, 47.04, 28.32, 28.02, 24.75. HRMS (ESI) $m/z$ calculated 456.2498 for $C_{25}H_{34}N_3O_5$ $[M+H]^+$, found 456.2510.

#### Display Report

**Analysis Info**

| | | | |
|---|---|---|---|
| Analysis Name | Z:\Data\2024\2408\sam280824\VTE9_8_01_127256.d | Acquisition Date | 28.08.2024 19:53:41 |
| Method | hystar_pl.m | Operator | BDAL@DE |
| Sample Name | VTE9 | Instrument / Ser# | micrOTOF 10237 |
| Comment | | | |

**Acquisition Parameter**

| | | | | | |
|---|---|---|---|---|---|
| Source Type | ESI | Ion Polarity | Positive | Set Nebulizer | 1.2 Bar |
| Focus | Not active | | | Set Dry Heater | 180 °C |
| Scan Begin | 50 m/z | Set Capillary | 4500 V | Set Dry Gas | 4.0 l/min |
| Scan End | 1600 m/z | Set End Plate Offset | -500 V | Set Divert Valve | Source |

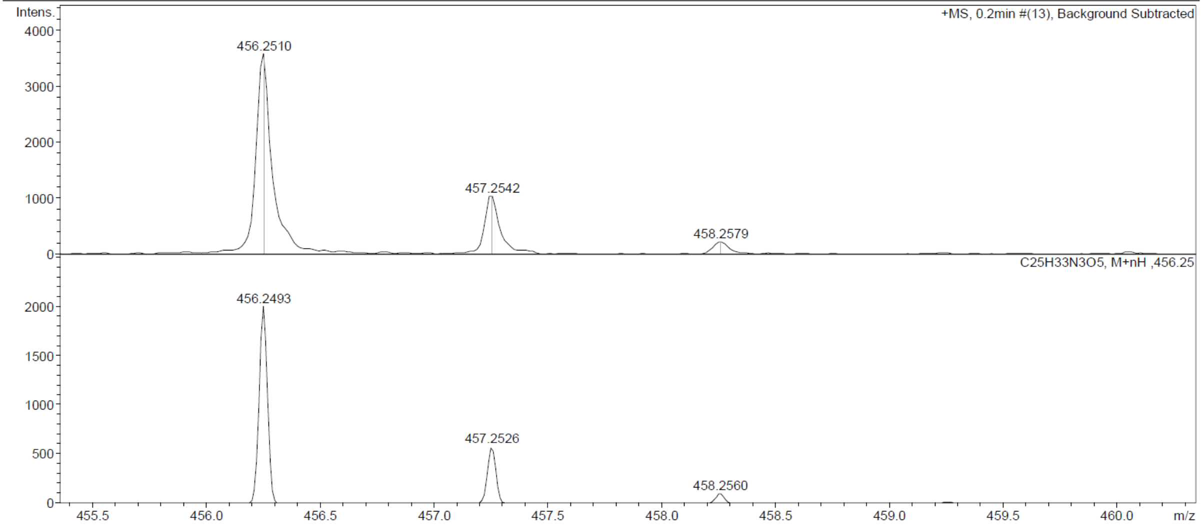

Compound **15a:**

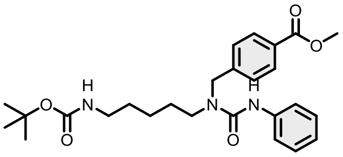

Purified by flash column chromatography (Biotage® Sfär Silica HC 25 g, gradient 2% to 50% hexane – EtOAc). Obtained 971 mg of target compound in 68% yield as a colorless liquid by general method A.

$^1$H NMR (400 MHz, $CDCl_3$) δ 8.13 – 8.00 (m, 2H), 7.42 (dd, $J$ = 9.0, 2.9 Hz, 4H), 7.33 – 7.26 (m, 2H), 7.06 (ddt, $J$ = 8.2, 7.0, 1.1 Hz, 1H), 6.70 (s, 1H), 4.67 (s, 2H), 3.96 (s, 3H), 3.41 – 3.30 (m, 2H), 3.17 (t, $J$ = 6.7 Hz, 2H), 1.71 (s, 2H), 1.53 (p, $J$ = 7.0 Hz, 2H), 1.46 (s, 9H), 1.36 (h, $J$ = 6.3 Hz, 2H). $^{13}$C NMR (101 MHz, $CDCl_3$) δ 166.82, 156.27, 155.43, 143.23, 139.07, 130.18, 129.52, 128.78, 127.18, 123.12, 120.25, 79.33, 52.18, 50.58, 47.89, 39.67, 29.75, 28.44, 27.38, 23.40. HRMS (ESI) $m/z$ calculated 470.2654 for $C_{26}H_{36}N_3O_5$ [M+H]$^+$, found 470.2634.

Display Report

**Analysis Info**

| | | | |
|---|---|---|---|
| Analysis Name | Z:\Data\2024\2408\sam280824\VTE10_9_01_127257.d | Acquisition Date | 28.08.2024 19:58:20 |
| Method | hystar_pl.m | Operator | BDAL@DE |
| Sample Name | VTE10 | Instrument / Ser# | micrOTOF 10237 |
| Comment | | | |

**Acquisition Parameter**

| | | | | | |
|---|---|---|---|---|---|
| Source Type | ESI | Ion Polarity | Positive | Set Nebulizer | 1.2 Bar |
| Focus | Not active | | | Set Dry Heater | 180 °C |
| Scan Begin | 50 m/z | Set Capillary | 4500 V | Set Dry Gas | 4.0 l/min |
| Scan End | 1600 m/z | Set End Plate Offset | -500 V | Set Divert Valve | Source |

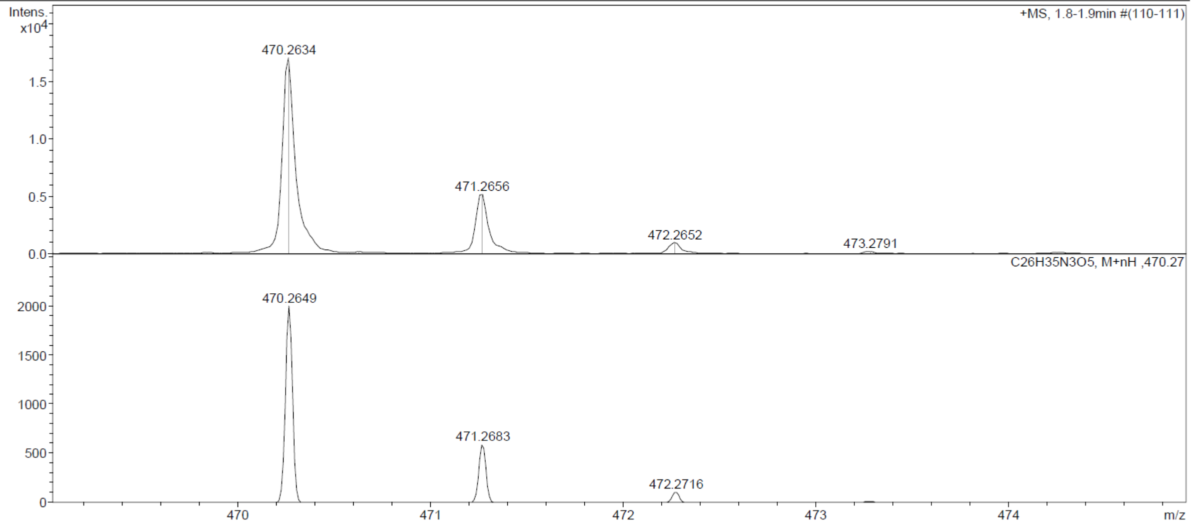

Compound **16a:**

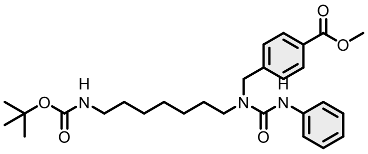

Purified by flash column chromatography (Biotage® Sfär Silica HC 25 g, gradient 2% to 50% hexane – EtOAc). Obtained 1059 mg of target compound in 70% yield as a colorless liquid by general method A.

$^{1}$H NMR (400 MHz, $CDCl_3$) δ 8.13 – 7.98 (m, 2H), 7.45 – 7.38 (m, 2H), 7.37 – 7.24 (m, 5H), 7.11 – 7.02 (m, 1H), 6.43 (s, 1H), 4.66 (s, 2H), 3.95 (s, 3H), 3.42 – 3.31 (m, 2H), 3.12 (t, $J$ = 7.1 Hz, 2H), 2.08 (s, 1H), 1.66 (s, 2H), 1.47 (s, 11H), 1.36 – 1.31 (m, 6H). $^{13}$C NMR (101 MHz, $CDCl_3$) δ 166.79, 156.07, 155.33, 143.09, 138.91, 130.20, 128.88, 127.11, 123.19, 119.98, 119.92, 79.17, 52.19, 50.59, 48.01, 40.61, 29.99, 28.97, 28.45, 28.34, 26.89, 26.64. HRMS (ESI) $m/z$ calculated 498.2967 for $C_{28}H_{40}N_3O_5$ $[M+H]^+$, found 498.2972.

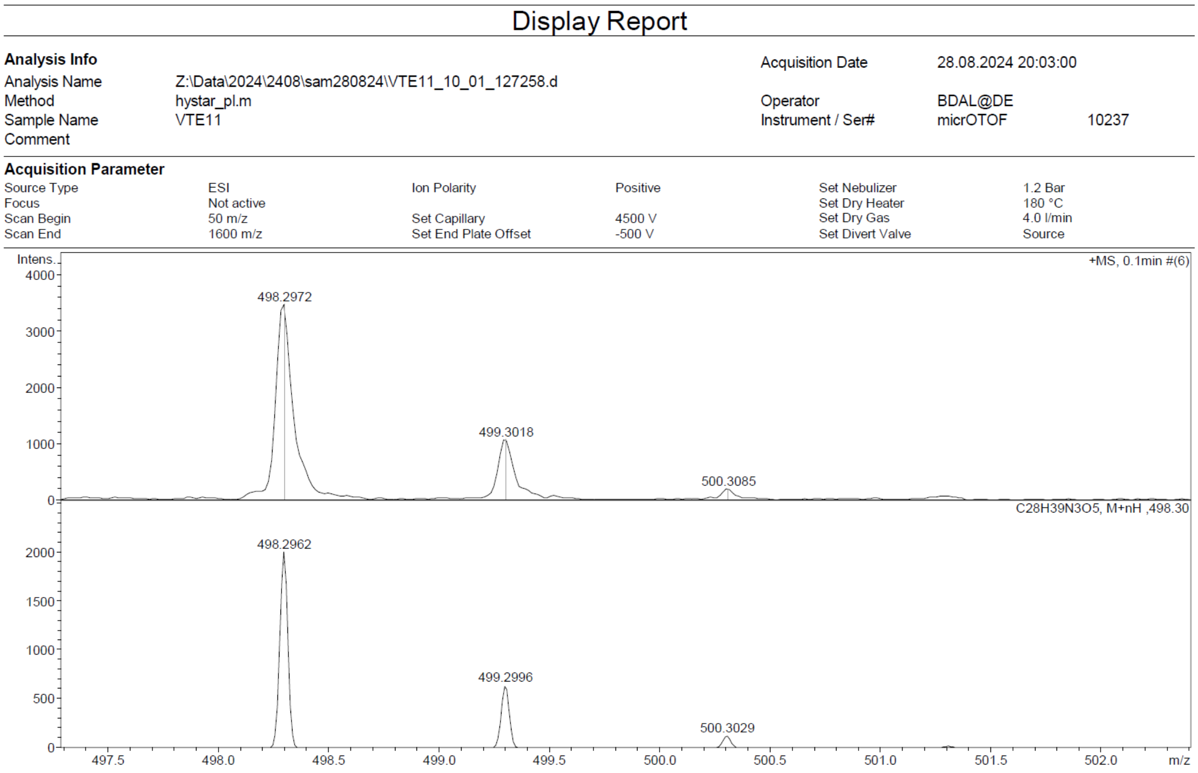

Compound **17a:**

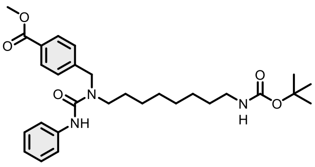

Purified by flash column chromatography (Biotage® Sfär Silica HC 25 g, gradient 2% to 50% hexane – EtOAc). Obtained 1245 mg of target compound in 88% yield as a colorless liquid by general method A.

$^{1}$H NMR (400 MHz, $CDCl_3$) δ 8.13 – 7.96 (m, 2H), 7.44 – 7.32 (m, 4H), 7.32 – 7.24 (m, 2H), 7.04 (tt, $J$ = 7.3, 1.3 Hz, 1H), 6.54 (s, 1H), 4.65 (s, 2H), 3.94 (s, 3H), 3.43 – 3.29 (m, 2H), 3.10 (t, $J$ = 7.1 Hz, 2H), 1.64 (dd, $J$ = 10.0, 4.9 Hz, 2H), 1.46 (s, 11H), 1.34 – 1.27 (m, 8H). $^{13}$C NMR (101 MHz, $CDCl_3$) δ 166.82, 156.08, 155.39, 143.26, 139.01, 130.15, 129.47, 128.84, 127.13, 123.12, 120.03, 79.09, 52.17, 50.51, 47.95, 40.65, 30.03, 29.23, 29.11, 28.46, 28.36, 26.87, 26.63. HRMS (ESI) *m*/*z* calculated 512.3124 for $C_{29}H_{42}N_3O_5$ [M+H]$^{+}$, found 512.3129.

#### Display Report

**Analysis Info**

| | | | |
|---|---|---|---|
| Analysis Name | Z:\Data\2024\2408\sam280824\VTE12_11_01_127259.d | Acquisition Date | 28.08.2024 20:07:42 |
| Method | hystar_pl.m | Operator | BDAL@DE |
| Sample Name | VTE12 | Instrument / Ser# | micrOTOF 10237 |
| Comment | | | |

**Acquisition Parameter**

| | | | | | |
|---|---|---|---|---|---|
| Source Type | ESI | Ion Polarity | Positive | Set Nebulizer | 1.2 Bar |
| Focus | Not active | | | Set Dry Heater | 180 °C |
| Scan Begin | 50 m/z | Set Capillary | 4500 V | Set Dry Gas | 4.0 l/min |
| Scan End | 1600 m/z | Set End Plate Offset | -500 V | Set Divert Valve | Source |

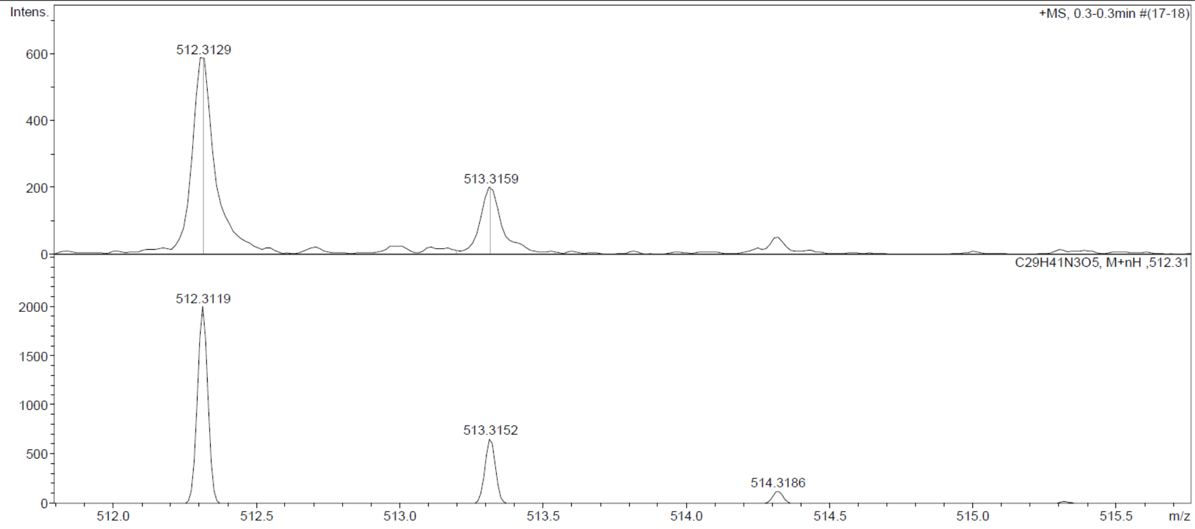

Compound **18a:**

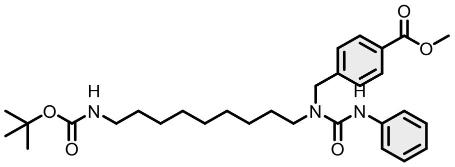

Purified by flash column chromatography (Biotage® Sfär Silica HC 25 g, gradient 2% to 50% hexane – EtOAc). Obtained 396 mg of target compound in 62% yield as a colorless liquid by general method A.

$^{1}$H NMR (400 MHz, $CDCl_3$) δ 8.07 – 7.96 (m, 2H), 7.40 – 7.29 (m, 4H), 7.28 – 7.20 (m, 2H), 7.03 – 6.97 (m, 1H), 6.50 (s, 1H), 4.62 (s, 2H), 3.90 (s, 3H), 3.39 – 3.24 (m, 2H), 3.07 (t, $J$ = 7.1 Hz, 2H), 1.60 (s, 2H), 1.43 (s, 11H), 1.30 – 1.24 (m, 10H). $^{13}$C NMR (101 MHz, $CDCl_3$) δ 166.78, 156.06, 155.36, 143.22, 138.96, 130.11, 129.44, 128.81, 127.10, 123.12, 120.02, 79.06, 52.13, 50.48, 47.94, 40.73, 30.01, 29.37, 29.22, 29.11, 28.42, 28.36, 26.90, 26.68. HRMS (ESI) $m/z$ calculated 526.3281 for $C_{30}H_{44}N_3O_5$, found 526.3280 $[M+H]^+$).

#### Display Report

**Analysis Info**

| | | | |
|---|---|---|---|
| Analysis Name | Z:\Data\2024\2408\sam280824\VTE1_3_01_127251.d | Acquisition Date | 28.08.2024 19:30:26 |
| Method | hystar_pl.m | Operator | BDAL@DE |
| Sample Name | VTE1 | Instrument / Ser# | micrOTOF 10237 |
| Comment | | | |

**Acquisition Parameter**

| | | | | | |
|---|---|---|---|---|---|
| Source Type | ESI | Ion Polarity | Positive | Set Nebulizer | 1.2 Bar |
| Focus | Not active | | | Set Dry Heater | 180 °C |
| Scan Begin | 50 m/z | Set Capillary | 4500 V | Set Dry Gas | 4.0 l/min |
| Scan End | 1600 m/z | Set End Plate Offset | -500 V | Set Divert Valve | Source |

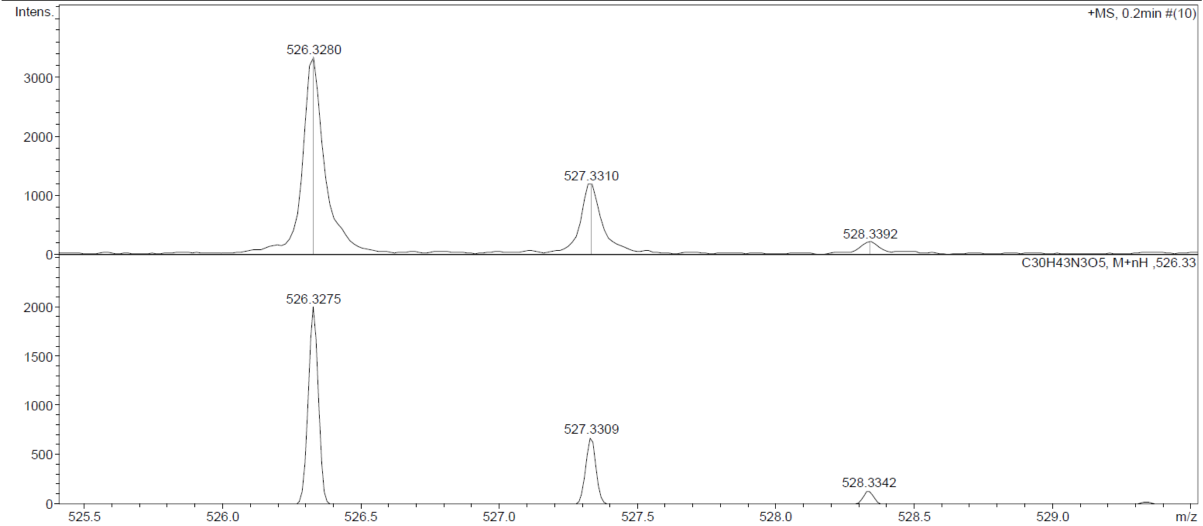

**Compound 19a:**

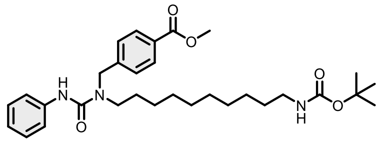

Purified by flash column chromatography (Biotage® Sfär Silica HC 25 g, gradient 2% to 50% hexane – EtOAc). Obtained 485 mg of target compound in 74% yield as a colorless liquid by general method A.

$^1$H NMR (400 MHz, $CDCl_3$) δ 7.41 – 7.18 (m, 8H), 7.00 (tt, $J$ = 7.2, 1.3 Hz, 1H), 6.56 (s, 1H), 4.65 (s, 2H), 4.55 (s, 2H), 3.41 – 3.29 (m, 2H), 3.09 (t, $J$ = 7.1 Hz, 2H), 1.62 (s, 2H), 1.46 (s, 11H), 1.37 – 1.20 (m, 12H). $^{13}$C NMR (101 MHz, $CDCl_3$) δ 156.14, 155.63, 140.83, 139.16, 136.63, 128.76, 127.44, 127.11, 122.93, 120.01, 79.04, 64.51, 50.46, 47.83, 40.62, 30.04, 29.44, 29.34, 29.23, 28.47, 28.37, 26.95, 26.77. HRMS (ESI) $m/z$ calculated 562.3251 for $C_{31}H_{46}NaN_3O_5$ $[M+Na]^+$, found 562.3253.

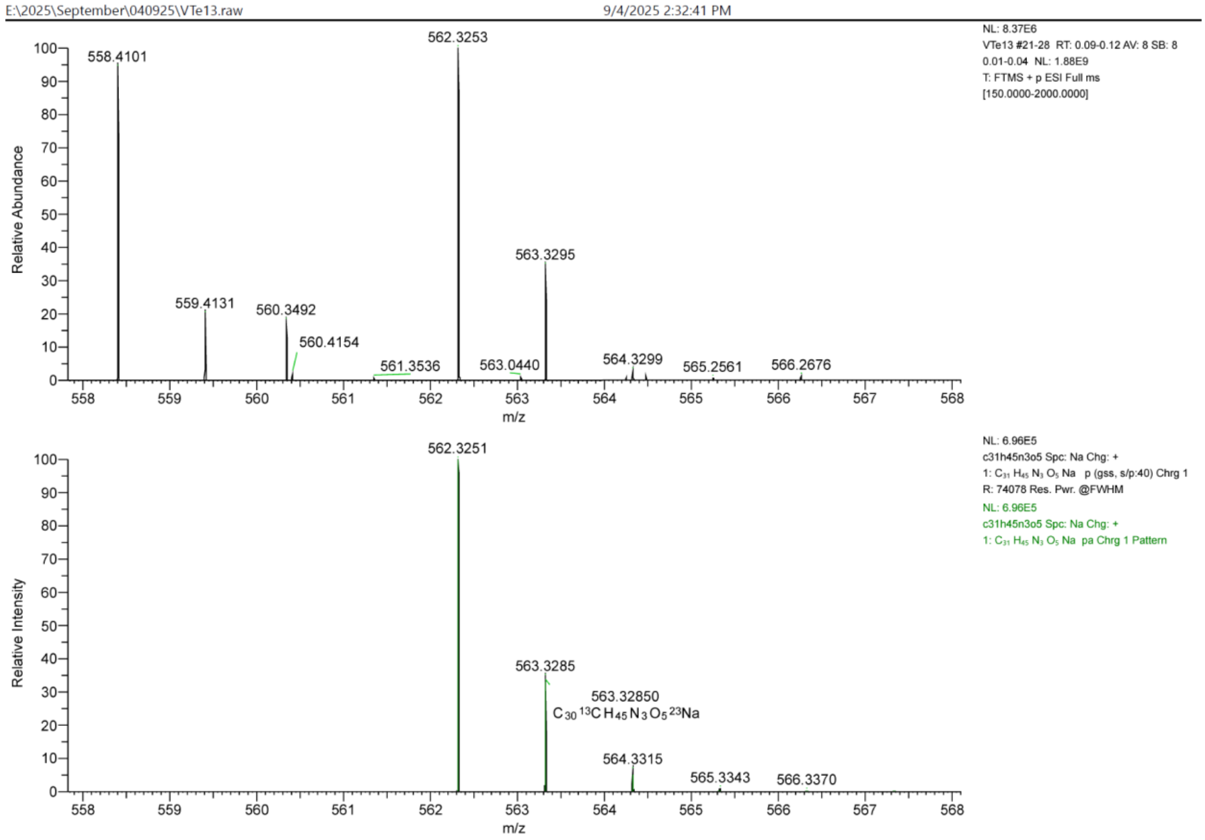

Compound **20a:**

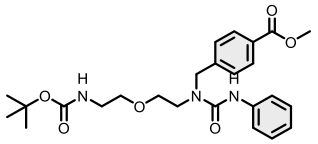

Purified by flash column chromatography (Biotage® Sfär Silica HC 25 g, gradient 10% to 80% hexane – EtOAc). Obtained 1061 mg of target compound in 80% yield as a colorless liquid by general method A.

$^{1}$H NMR (400 MHz, $CDCl_3$) δ 8.11 – 7.96 (m, 3H), 7.46 – 7.35 (m, 4H), 7.31 (dd, $J$ = 8.6, 7.3 Hz, 2H), 7.09 – 6.98 (m, 1H), 4.70 (s, 2H), 3.94 (s, 3H), 3.60 (dt, $J$ = 7.0, 5.2 Hz, 4H), 3.56 – 3.50 (m, 2H), 3.40 (t, $J$ = 5.5 Hz, 2H), 1.46 (s, 9H). $^{13}$C NMR (101 MHz, $CDCl_3$) δ 166.87, 156.87, 155.93, 143.66, 139.68, 130.02, 129.39, 128.98, 127.68, 122.59, 119.12, 79.72, 70.85, 70.56, 52.14, 50.85, 48.33, 40.22, 28.37. HRMS (ESI) $m/z$ calculated 472.2447 for $C_{25}H_{34}N_3O_6$ $[M+H]^+$, found 472.2459.

#### Display Report

**Analysis Info**

| | | | |
|---|---|---|---|
| Analysis Name | Z:\Data\2024\2408\sam280824\VTE5_4_01_127252.d | Acquisition Date | 28.08.2024 19:35:06 |
| Method | hystar_pl.m | Operator | BDAL@DE |
| Sample Name | VTE5 | Instrument / Ser# | micrOTOF 10237 |
| Comment | | | |

**Acquisition Parameter**

| | | | | | |
|---|---|---|---|---|---|
| Source Type | ESI | Ion Polarity | Positive | Set Nebulizer | 1.2 Bar |
| Focus | Not active | | | Set Dry Heater | 180 °C |
| Scan Begin | 50 m/z | Set Capillary | 4500 V | Set Dry Gas | 4.0 l/min |
| Scan End | 1600 m/z | Set End Plate Offset | -500 V | Set Divert Valve | Source |

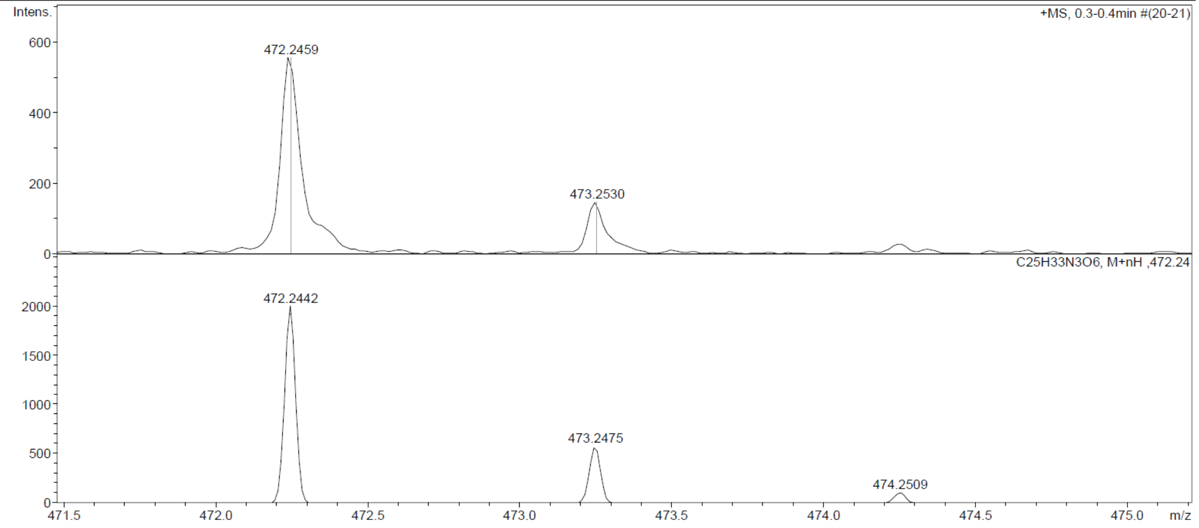

Compound **21a:**

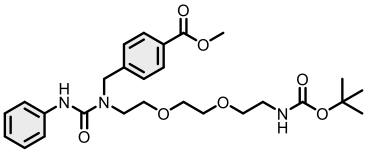

Purified by flash column chromatography (Biotage® Sfär Silica HC 25 g, gradient 10% to 80% hexane – EtOAc). Obtained 1035 mg of target compound in 66% yield as a colorless liquid by general method A.

$^{1}$H NMR (400 MHz, $CDCl_3$) δ 8.33 (s, 1H), 8.10 – 7.96 (m, 2H), 7.49 – 7.38 (m, 4H), 7.35 – 7.25 (m, 2H), 7.07 – 6.96 (m, 1H), 4.70 (s, 2H), 3.94 (s, 3H), 3.72 – 3.64 (m, 4H), 3.64 – 3.58 (m, 2H), 3.53 (t, $J$ = 5.1 Hz, 4H), 3.28 (t, $J$ = 5.3 Hz, 2H), 1.45 (s, 9H). $^{13}$C NMR (101 MHz, $CDCl_3$) δ 166.90, 157.13, 155.94, 143.84, 139.98, 129.96, 129.31, 128.82, 127.78, 122.37, 119.09, 79.35, 71.06, 70.86, 70.30, 69.97, 52.13, 50.79, 48.40, 40.30, 28.41. HRMS (ESI) $m/z$ calculated 516.2709 for $C_{27}H_{38}N_3O_7$ $[M+H]^+$, found 516.2725.

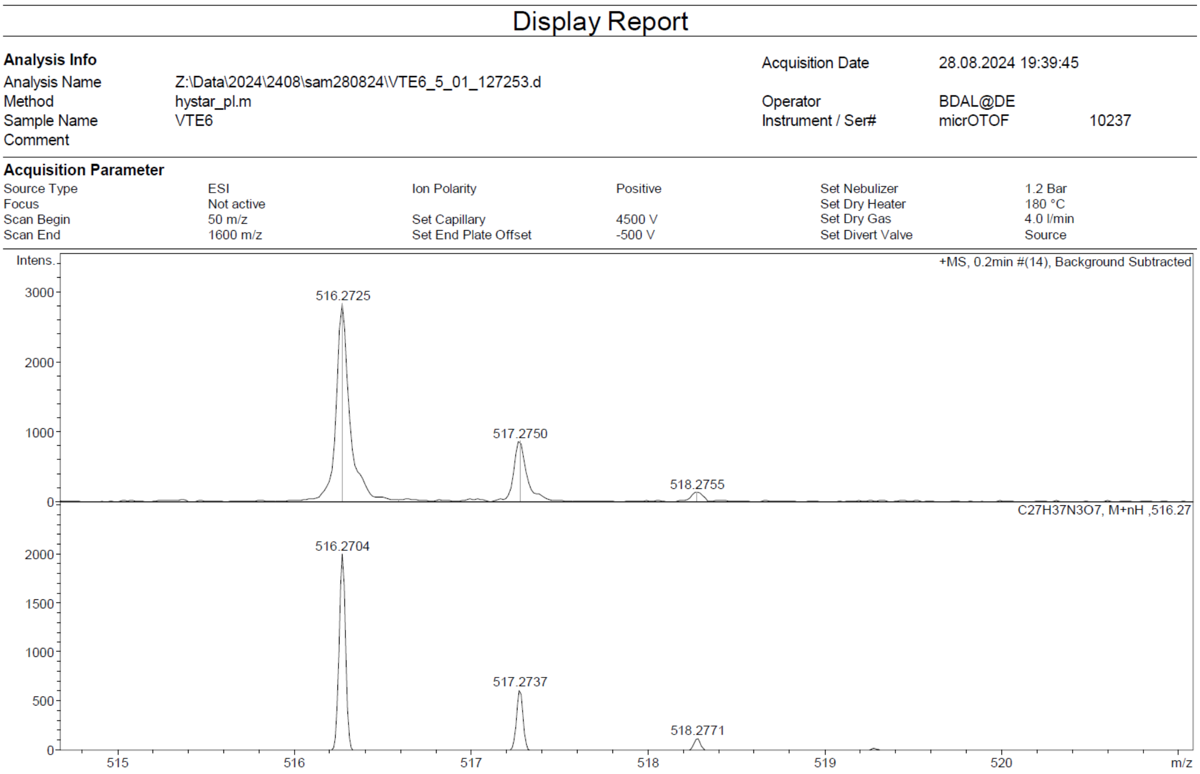

Compound **22a:**

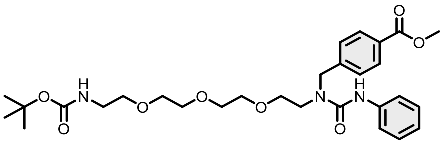

Purified by flash column chromatography (Biotage® Sfär Silica HC 25 g, gradient 10% to 80% hexane – EtOAc). Obtained 1242 mg of target compound in 73% yield as a colorless liquid by general method A.

$^{1}$H NMR (400 MHz, $CDCl_3$) δ 8.36 (s, 1H), 8.13 – 7.95 (m, 2H), 7.51 – 7.39 (m, 4H), 7.37 – 7.22 (m, 2H), 7.02 (tt, $J$ = 7.3, 1.2 Hz, 1H), 4.70 (s, 2H), 3.94 (s, 3H), 3.72 (s, 4H), 3.67 – 3.46 (m, 11H), 3.29 (t, $J$ = 5.2 Hz, 2H), 1.46 (s, 9H). $^{13}$C NMR (101 MHz, $CDCl_3$) δ 166.92, 157.16, 155.98, 143.89, 140.05, 129.95, 129.28, 128.78, 127.78, 122.28, 119.11, 79.25, 71.04, 70.96, 70.54, 70.31, 70.27, 70.09, 52.13, 50.78, 48.43, 40.45, 28.43. HRMS (ESI) $m/z$ calculated 560.2971 for $C_{29}H_{42}N_3O_8$ [M+H]$^+$, found 560.2977.

#### Display Report

**Analysis Info**

| | | | |
|---|---|---|---|
| Analysis Name | Z:\Data\2024\2408\sam280824\VTE7_6_01_127254.d | Acquisition Date | 28.08.2024 19:44:23 |
| Method | hystar_pl.m | Operator | BDAL@DE |
| Sample Name | VTE7 | Instrument / Ser# | micrOTOF 10237 |
| Comment | | | |

**Acquisition Parameter**

| | | | | | |
|---|---|---|---|---|---|
| Source Type | ESI | Ion Polarity | Positive | Set Nebulizer | 1.2 Bar |
| Focus | Not active | | | Set Dry Heater | 180 °C |
| Scan Begin | 50 m/z | Set Capillary | 4500 V | Set Dry Gas | 4.0 l/min |
| Scan End | 1600 m/z | Set End Plate Offset | -500 V | Set Divert Valve | Source |

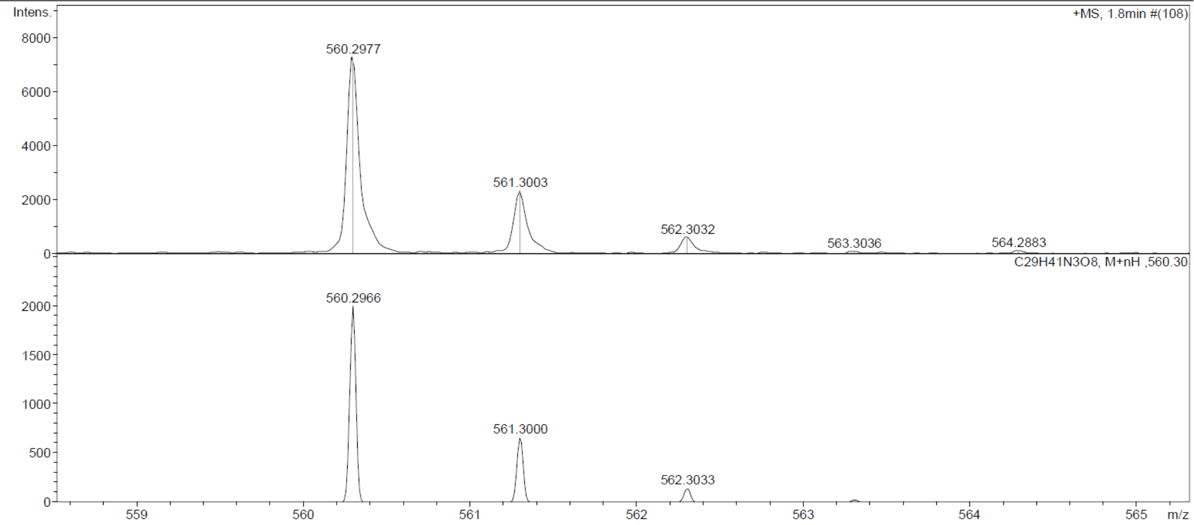

Compound **23a:**

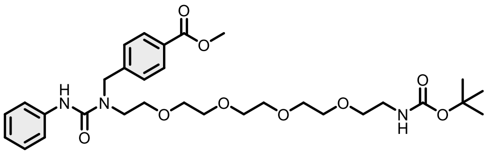

Purified by flash column chromatography (Biotage® Sfär Silica HC 25 g, gradient 10% to 80% hexane – EtOAc). Obtained 753 mg of target compound in 41% yield as a colorless liquid by general method A.

$^{1}$H NMR (400 MHz, $CDCl_3$) δ 8.36 (s, 1H), 8.09 – 7.97 (m, 2H), 7.52 – 7.37 (m, 4H), 7.36 – 7.25 (m, 2H), 7.01 (tt, $J$ = 7.3, 1.2 Hz, 1H), 4.69 (s, 2H), 3.94 (s, 3H), 3.72 (s, 4H), 3.69 – 3.56 (m, 11H), 3.55 – 3.51 (m, 4H), 3.31 (t, $J$ = 5.2 Hz, 2H), 1.47 (s, 9H). $^{13}$C NMR (101 MHz, $CDCl_3$) δ 166.91, 157.14, 155.99, 143.88, 140.06, 129.95, 129.28, 128.78, 127.77, 122.24, 119.13, 79.24, 71.00, 70.96, 70.58, 70.56, 70.52, 70.49, 70.30, 70.20, 52.13, 50.77, 48.42, 48.22, 28.45. HRMS (ESI) $m/z$ calculated 604.3234 for $C_{31}H_{46}N_3O_9$ $[M+H]^+$, found 604.3215.

#### Display Report

**Analysis Info**

| | | | |
|---|---|---|---|
| Analysis Name | Z:\Data\2024\2408\sam280824\VTE8_7_01_127255.d | Acquisition Date | 28.08.2024 19:49:01 |
| Method | hystar_pl.m | Operator | BDAL@DE |
| Sample Name | VTE8 | Instrument / Ser# | micrOTOF 10237 |
| Comment | | | |

**Acquisition Parameter**

| | | | | | |
|---|---|---|---|---|---|
| Source Type | ESI | Ion Polarity | Positive | Set Nebulizer | 1.2 Bar |
| Focus | Not active | | | Set Dry Heater | 180 °C |
| Scan Begin | 50 m/z | Set Capillary | 4500 V | Set Dry Gas | 4.0 l/min |
| Scan End | 1600 m/z | Set End Plate Offset | -500 V | Set Divert Valve | Source |

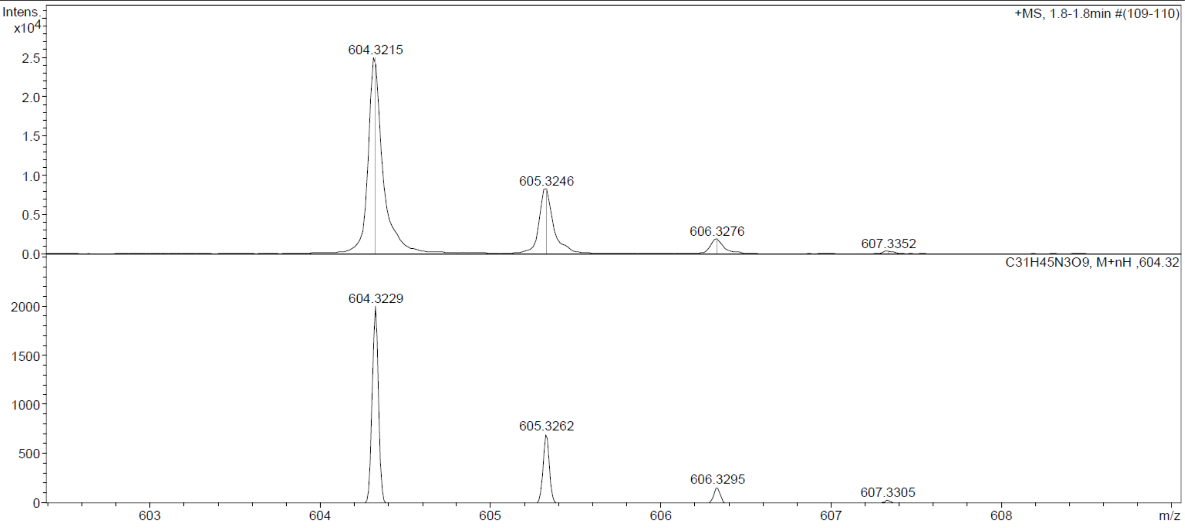

Compound **1b**:

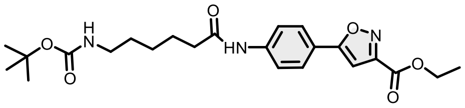

Compound **1a** (2.0 g, 6 mmol) and ethyl 2-chloro-2(hydroxyimino)acetate (2.7 g, 18 mmol) were mixed in 20 mL of anhydrous THF and triethylamine (24 mmol) was added slowly over 6 h by syringe pump. The mixture was stirred overnight and then diluted with 100 mL of ethyl acetate and 50 mL of 1 M HCl. The two layers were separated, and the aqueous layer was extracted with ethyl acetate (2 × 50 mL). The combined organic layers were washed with water and brine, dried over anhydrous $Na_2SO_4$, filtered, and concentrated in vacuo. Purification of the crude reaction mixture by column chromatography afforded the title compound (1.29g, 48 %) as a white solid.

$^1$H NMR (400 MHz, $CDCl_3$) δ 8.59 (d, $J$ = 11.2 Hz, 1H), 7.82 – 7.58 (m, 4H), 6.81 (d, $J$ = 0.8 Hz, 1H), 4.41 (qd, $J$ = 7.2, 1.1 Hz, 2H), 3.06 (q, $J$ = 6.7 Hz, 2H), 2.37 (t, $J$ = 7.5 Hz, 2H), 1.70 (p, $J$ = 7.5 Hz, 2H), 1.53 – 1.43 (m, 2H), 1.43 – 1.16 (m, 13H). $^{13}$C NMR (101 MHz, $CDCl_3$) δ 172.05, 171.43, 159.97, 156.89, 156.25, 140.85, 126.67, 121.80, 119.88, 99.13, 79.18, 62.20, 40.27, 37.27, 29.71, 28.39, 26.27, 24.98, 14.11. HRMS (ESI) $m/z$ calculated 446.2291 for $C_{23}H_{32}N_3O_6$ [M+H]$^+$, found 446.2289.

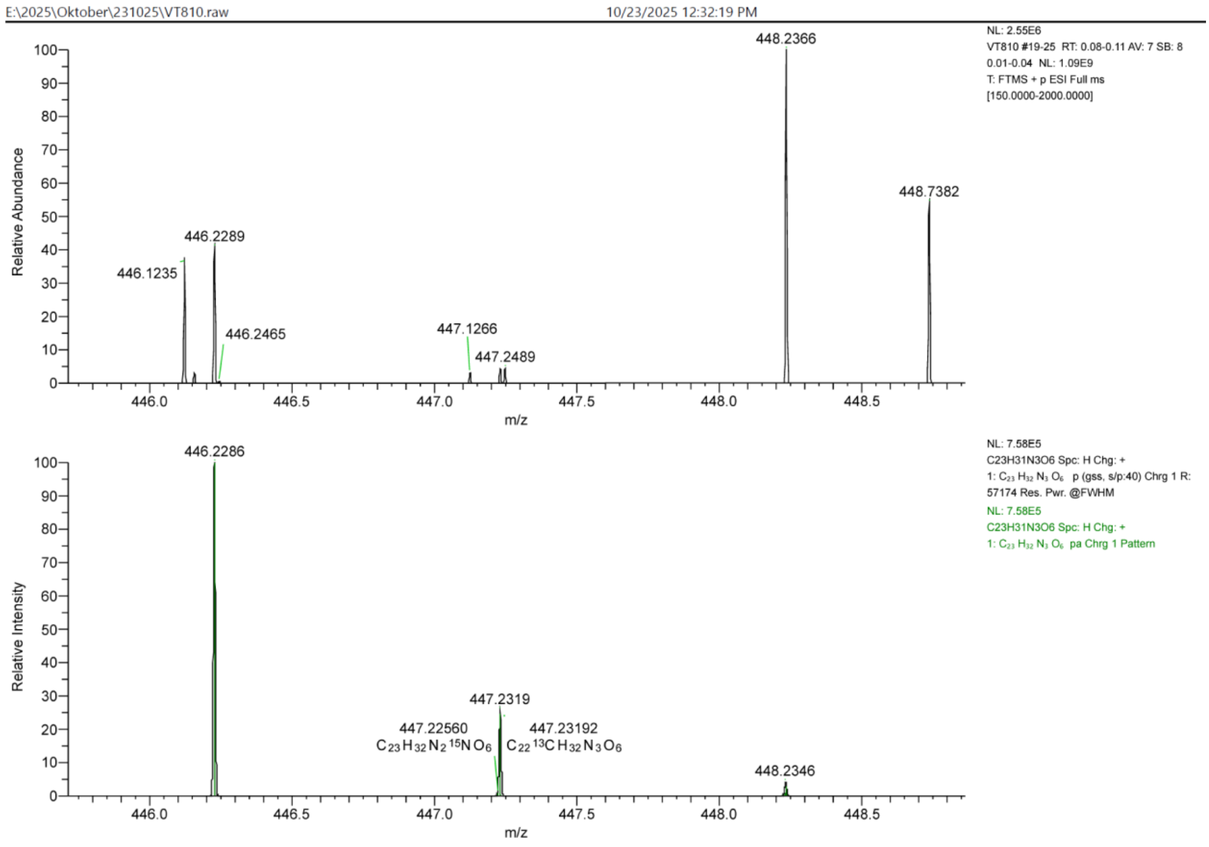

Compound **3b:**

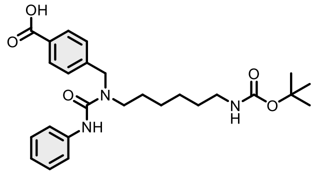

Evaporated ethyl acetate to give 691 mg of targeted compound in 89% yield as a white solid by general method B.

$^{1}$H NMR (400 MHz, DMSO-$d_6$) δ 12.80 (s, 1H), 8.38 (s, 1H), 7.98 – 7.87 (m, 2H), 7.52 – 7.43 (m, 2H), 7.42 – 7.35 (m, 2H), 7.31 – 7.18 (m, 2H), 7.02 – 6.90 (m, 1H), 4.66 (s, 2H), 3.31 (t, *J* = 7.6 Hz, 2H), 2.88 (q, *J* = 6.6 Hz, 2H), 1.50 (t, *J* = 7.4 Hz, 2H), 1.36 (s, 11H), 1.23 (p, *J* = 3.6 Hz, 4H). $^{13}$C NMR (101 MHz, DMSO-$d_6$) δ 167.64, 156.03, 155.69, 144.77, 140.90, 129.96, 129.89, 128.66, 127.60, 122.32, 120.60, 77.73, 60.21, 49.62, 46.88, 29.89, 28.72, 28.24, 26.56, 26.39. HRMS (ESI) *m*/*z* calculated 470.2654 for $C_{26}H_{36}N_3O_5$ [M+H]$^+$, found 470.2653.

Display Report

**Analysis Info**

| | | | |
|---|---|---|---|
| Analysis Name | Z:\Data\2024\2408\sam290824\VTA16_31_01_127303.d | Acquisition Date | 29.08.2024 22:05:21 |
| Method | hystar_pl.m | Operator | BDAL@DE |
| Sample Name | VTA16 | Instrument / Ser# | micrOTOF 10237 |
| Comment | | | |

**Acquisition Parameter**

| | | | | | |
|---|---|---|---|---|---|
| Source Type | ESI | Ion Polarity | Positive | Set Nebulizer | 1.2 Bar |
| Focus | Not active | | | Set Dry Heater | 180 °C |
| Scan Begin | 50 m/z | Set Capillary | 4500 V | Set Dry Gas | 4.0 l/min |
| Scan End | 1600 m/z | Set End Plate Offset | -500 V | Set Divert Valve | Source |

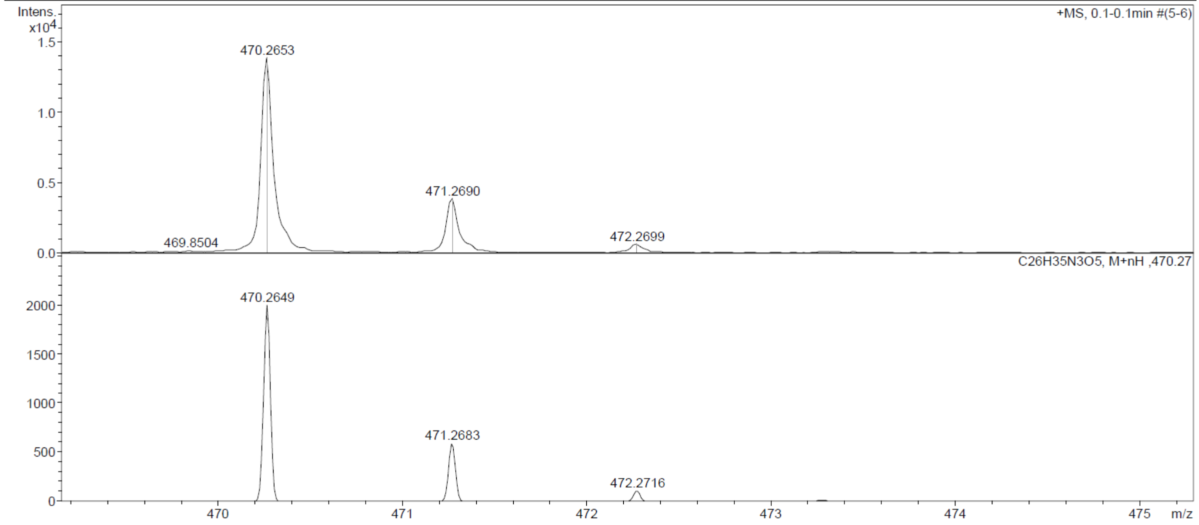

Compound **9b:**

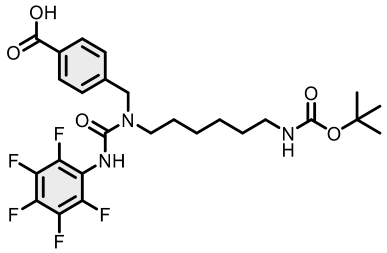

Evaporated ethyl acetate to give 887 mg of targeted compound in 91% yield as a white solid by general method B.

$^{1}$H NMR (400 MHz, DMSO-$d_6$) δ 12.77 (s, 1H), 8.59 (s, 1H), 8.03 – 7.83 (m, 2H), 7.43 – 7.30 (m, 2H), 6.73 (s, 1H), 4.62 (s, 2H), 3.27 (t, $J$ = 7.6 Hz, 2H), 2.87 (q, $J$ = 6.6 Hz, 2H), 1.51 (d, $J$ = 7.8 Hz, 2H), 1.35 (s, 11H), 1.21 (p, $J$ = 4.1 Hz, 4H). $^{13}$C NMR (101 MHz, DMSO-$d_6$) δ 167.59, 156.02, 155.18, 144.12, 130.04, 129.95, 127.56, 115.77, 77.71, 60.19, 49.83, 47.11, 29.88, 28.68, 28.04, 26.51, 26.24, 21.47, 21.18, 14.51. HRMS (ESI) *m/z* calculated 560.2183 for $C_{26}H_{31}F_5N_3O_5$, found 560.2184 ([M+H]$^+$).

#### Display Report

**Analysis Info**

| | | | |
|---|---|---|---|
| Analysis Name | Z:\Data\2024\2409\sam040924\VTA4_3_01_127407.d | Acquisition Date | 04.09.2024 18:02:25 |
| Method | hystar_pl.m | Operator | BDAL@DE |
| Sample Name | VTA4 | Instrument / Ser# | micrOTOF 10237 |
| Comment | | | |

**Acquisition Parameter**

| | | | | | |
|---|---|---|---|---|---|
| Source Type | ESI | Ion Polarity | Positive | Set Nebulizer | 1.2 Bar |
| Focus | Not active | | | Set Dry Heater | 180 °C |
| Scan Begin | 50 m/z | Set Capillary | 4500 V | Set Dry Gas | 4.0 l/min |
| Scan End | 1600 m/z | Set End Plate Offset | -500 V | Set Divert Valve | Source |

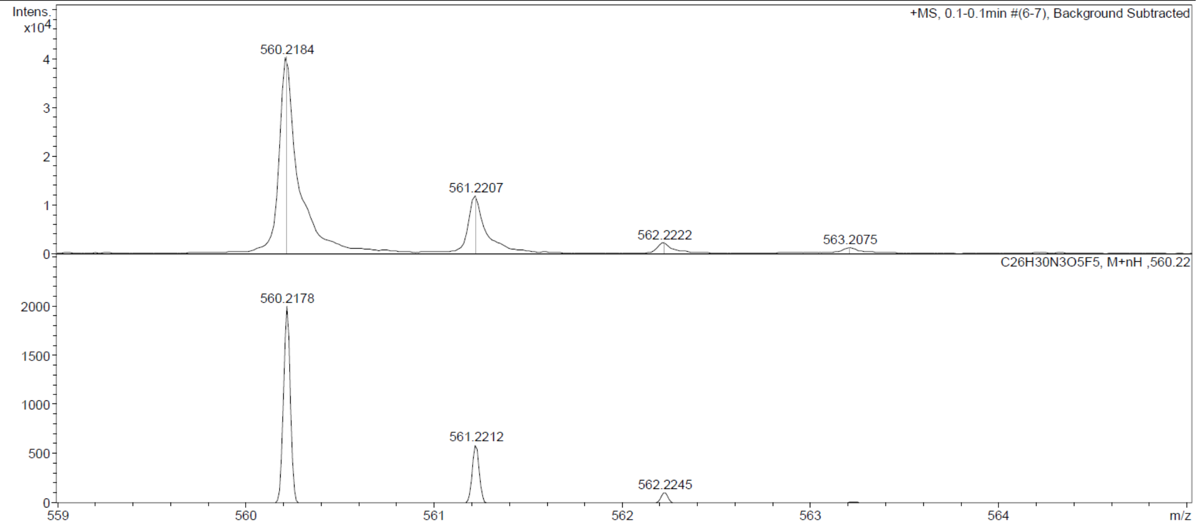

Compound **10b:**

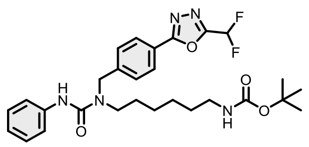

Compound 10a (1g, 4.76 mmol) and NBS (5.71 mmol) were combined in 20mL $CCl_4$ at rt. A catalytic amount of AIBN (0.47 mmol) was added, and the resulting mixture was refluxed, until LCMS analysis indicated that the reaction was completd. The reaction mixture was concentrated under reduced pressure and partitioned between EtOAc and water. The organic solution was washed consecutively with water and brine, before drying over $Na_2SO_4$. The suspension was filtered and concentrated to dryness under reduced pressure.

The crude, N-Boc-1,6-diaminohexane (9.52 mmol) and K2CO3 (14.2 mmol) were dissolved in 10mL of acetonitrile. The reaction mixture was then refluxed overnight. The reaction mixture was concentrated under reduced pressure and partitioned between EtOAc and water. The organic solution was washed consecutively with water and brine, before drying over $Na_2SO_4$. The organic phase was concentrated to dryness under reduced pressure. The crude was then dissolved in 10 mL of DCM. PhNCO (9.52 mmol) was added and the reaction was stirred at rt overnight. The reaction mixture was concentrated under reduced pressure and purified by column chromatography to afford **10b** (465 mg, 18%).

$^1$H NMR (400 MHz, DMSO-*d*$_6$) δ 8.39 (s, 1H), 8.08 – 8.00 (m, 2H), 7.67 – 7.42 (m, 5H), 7.26 – 7.19 (m, 2H), 6.94 (tt, *J* = 7.4, 1.2 Hz, 1H), 6.72 (t, *J* = 5.8 Hz, 1H), 4.68 (s, 2H), 3.32 (d, *J* = 7.8 Hz, 2H), 2.87 (d, *J* = 6.5 Hz, 2H), 1.54 – 1.47 (m, 2H), 1.34 (s, 11H), 1.25 – 1.20 (m, 4H). $^{13}$C NMR (101 MHz, DMSO-*d*$_6$) δ 165.71, 156.03, 155.72, 144.96, 140.84, 128.65, 128.60, 127.72, 122.37, 121.38, 120.64, 109.46, 107.08, 104.71, 77.72, 60.20, 49.69, 47.03, 29.88, 28.69, 28.29, 26.55, 26.37. HRMS (ESI) *m*/*z* calculated 544.2735 for $C_{28}H_{36}F_2N_5O_4$, found 544.2728 ($[M+H]^+$).

### Display Report

**Analysis Info**

| | | | |
|---|---|---|---|
| Analysis Name | Z:\Data\2024\2409\sam040924\VTA18_5_01_127429.d | Acquisition Date | 04.09.2024 22:25:47 |
| Method | hystar_pl.m | Operator | BDAL@DE |
| Sample Name | VTA18 | Instrument / Ser# | micrOTOF 10237 |
| Comment | | | |

**Acquisition Parameter**

| | | | | | |
|---|---|---|---|---|---|
| Source Type | ESI | Ion Polarity | Positive | Set Nebulizer | 1.2 Bar |
| Focus | Not active | | | Set Dry Heater | 180 °C |
| Scan Begin | 50 m/z | Set Capillary | 4500 V | Set Dry Gas | 4.0 l/min |
| Scan End | 1600 m/z | Set End Plate Offset | -500 V | Set Divert Valve | Source |

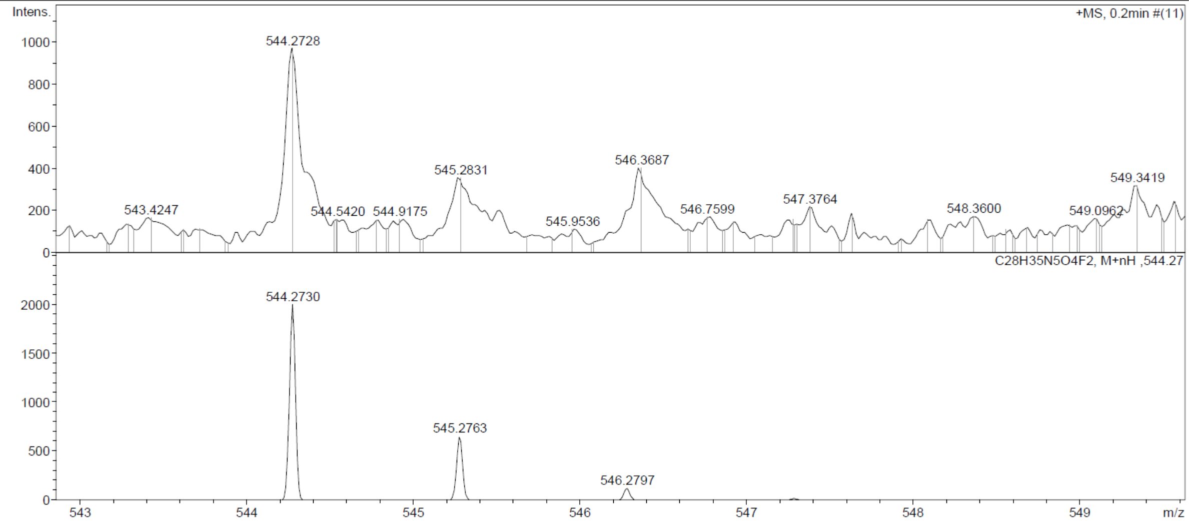

Compound **11b**:

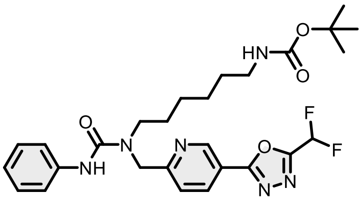

Compound 11a (1g, 4.74 mmol) and NBS (5.68 mmol) were combined in 20mL $CCl_4$ at rt. A catalytic amount of AIBN (0.47 mmol) was added, and the resulting mixture was refluxed, until LCMS analysis indicated that the reaction was completd. The reaction mixture was concentrated under reduced pressure and partitioned between EtOAc and water. The organic solution was washed consecutively with water and brine, before drying over $Na_2SO_4$. The suspension was filtered and concentrated to dryness under reduced pressure.

The crude, N-Boc-1,6-diaminohexane (9.47 mmol) and K2CO3 (14.2 mmol) were dissolved in 10mL of acetonitrile. The reaction mixture was then refluxed overnight. The reaction mixture was concentrated under reduced pressure and partitioned between EtOAc and water. The organic solution was washed consecutively with water and brine, before drying over $Na_2SO_4$. The organic phase was concentrated to dryness under reduced pressure. The crude was then dissolved in 10 mL of DCM. PhNCO (9.52 mmol) was added and the reaction was stirred at rt overnight. The reaction mixture was concentrated under reduced pressure and purified by column chromatography to afford **10b** (283 mg, 11%).

$^1$H NMR (400 MHz, $CDCl_3$) δ 9.24 (dd, $J$ = 2.2, 0.8 Hz, 1H), 8.43 (s, 1H), 8.32 (dd, $J$ = 8.1, 2.2 Hz, 1H), 7.49 (dd, $J$ = 8.2, 0.9 Hz, 1H), 7.40 – 7.31 (m, 2H), 7.25 – 7.14 (m, 2H), 7.11 – 6.73 (m, 2H), 4.77 (d, $J$ = 6.4 Hz, 1H), 4.60 (s, 2H), 3.43 – 3.30 (m, 2H), 2.99 (q, $J$ = 6.6 Hz, 2H), 1.53 (h, $J$ = 7.3 Hz, 2H), 1.37 (s, 11H), 1.23 (h, $J$ = 3.2 Hz, 4H). $^{13}$C NMR (101 MHz, $CDCl_3$) δ 163.81, 162.48, 156.17, 156.06, 147.65, 139.66, 135.90, 128.68, 122.81, 122.57, 119.73, 118.40, 108.12, 105.72, 103.32, 78.83, 60.31, 52.99, 48.25, 29.86, 28.35, 28.33, 26.35. HRMS (ESI) *m*/*z* calculated 545.2682 for $C_{27}H_{35}F_2N_6O_4$, found 545.2683 ($[M+H]^+$).

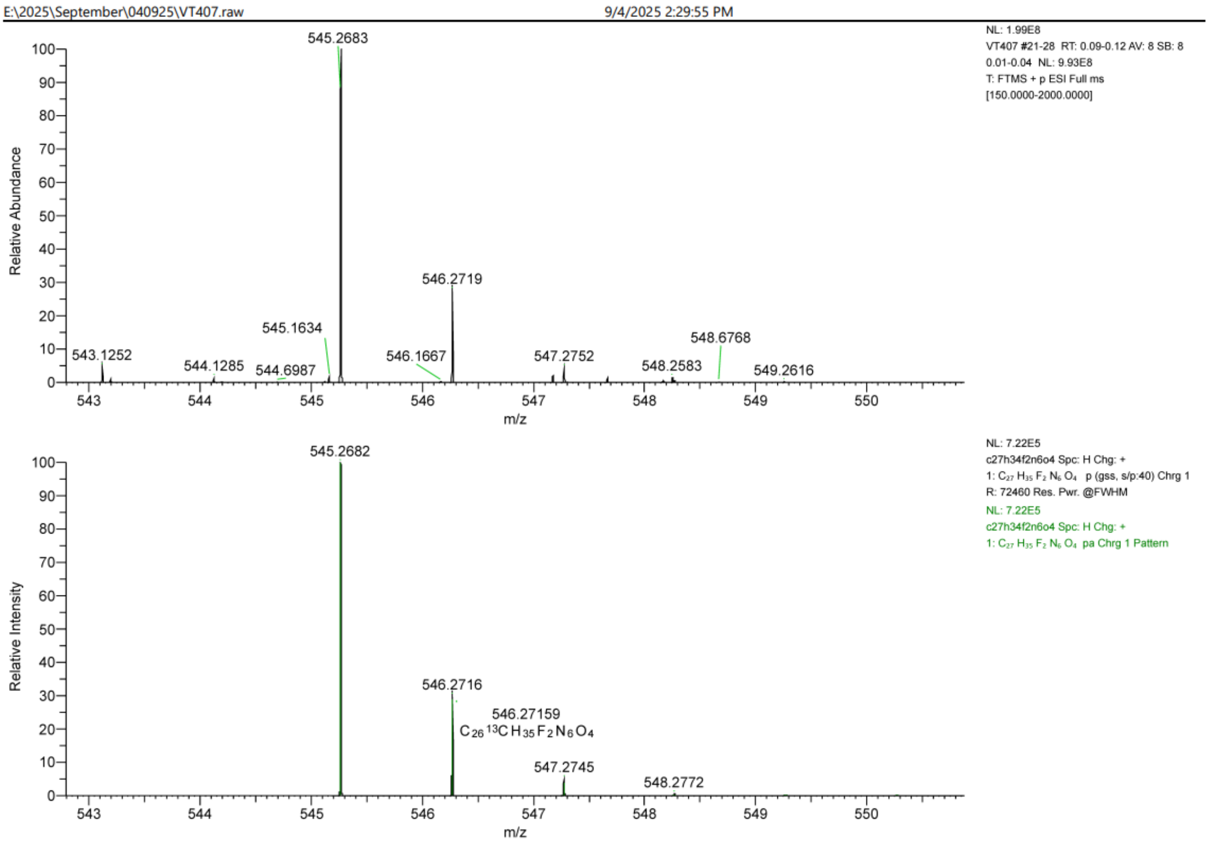
E:\2025\September\040925\VT407.raw
9/4/2025 2:29:55 PM
NL: 1.99E8
VT407 #21-28 RT: 0.09-0.12 AV: 8 SB: 8
0.01-0.04 NL: 9.93E8
T: FTMS + p ESI Full ms
[150.0000-2000.0000]
Relative Abundance
545.2683
546.2719
545.1634
548.6768
543.1252
546.1667
547.2752
544.1285
544.6987
548.2583
549.2616
m/z
NL: 7.22E5
c27h34f2n6o4 Spc: H Chg: +
1: C27 H35 F2 N6 O4 p (gss, s/p:40) Chrg 1
R: 72460 Res. Pwr. @FWHM
NL: 7.22E5
c27h34f2n6o4 Spc: H Chg: +
1: C27 H35 F2 N6 O4 pa Chrg 1 Pattern
Relative Intensity
545.2682
546.2716
546.27159
C26 13C H35 F2 N6 O4
547.2745
548.2772
m/z

Compound **12b:**

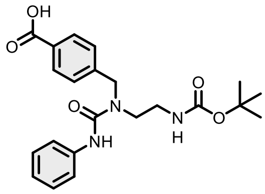

Evaporated ethyl acetate to give 688 mg of targeted compound in 89% yield as a white solid by general method B.

$^{1}$H NMR (400 MHz, DMSO-$d_6$) δ 12.88 (s, 1H), 8.49 (s, 1H), 7.94 (d, $J$ = 8.1 Hz, 2H), 7.53 (d, $J$ = 8.0 Hz, 2H), 7.38 (d, $J$ = 8.1 Hz, 2H), 7.31 – 7.20 (m, 2H), 7.08 – 6.88 (m, 2H), 4.68 (s, 2H), 3.38 (d, $J$ = 6.7 Hz, 2H), 3.11 (q, $J$ = 6.4 Hz, 2H), 1.37 (s, 9H). $^{13}$C NMR (101 MHz, DMSO-$d_6$) δ 167.65, 156.56, 155.69, 144.56, 140.87, 130.01, 129.95, 128.72, 127.64, 122.38, 120.31, 78.50, 50.06, 46.33, 39.21, 28.65. HRMS (ESI) $m/z$ calculated 414.2028 for $C_{22}H_{28}N_3O_5$ $[M+H]^+$, found 414.2036.

Display Report

**Analysis Info**

| | | | |
|---|---|---|---|
| Analysis Name | Z:\Data\2024\2408\sam290824\VTA15_30_01_127302.d | Acquisition Date | 29.08.2024 22:00:42 |
| Method | hystar_pl.m | Operator | BDAL@DE |
| Sample Name | VTA15 | Instrument / Ser# | micrOTOF 10237 |
| Comment | | | |

**Acquisition Parameter**

| | | | | | |
|---|---|---|---|---|---|
| Source Type | ESI | Ion Polarity | Positive | Set Nebulizer | 1.2 Bar |
| Focus | Not active | | | Set Dry Heater | 180 °C |
| Scan Begin | 50 m/z | Set Capillary | 4500 V | Set Dry Gas | 4.0 l/min |
| Scan End | 1600 m/z | Set End Plate Offset | -500 V | Set Divert Valve | Source |

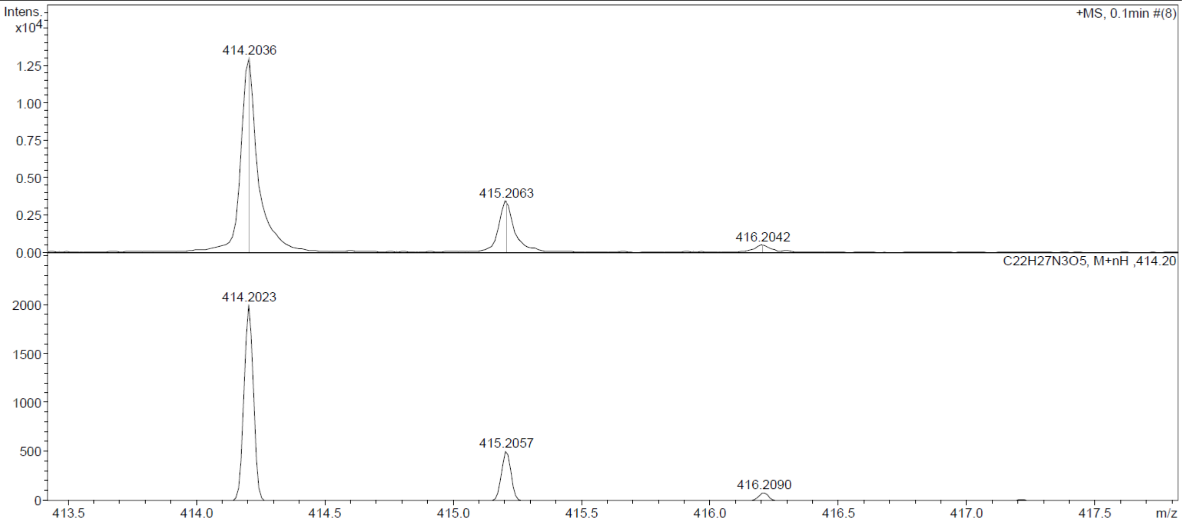

**Compound 13b:**

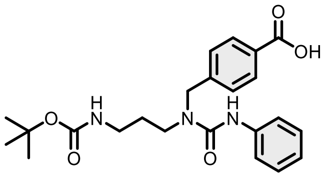

Evaporated ethyl acetate to give 619 mg of targeted compound in 80% yield as a white solid by general method B.

$^{1}$H NMR (400 MHz, DMSO-$d_6$) δ 12.84 (s, 1H), 8.40 (s, 1H), 8.04 – 7.84 (m, 2H), 7.54 – 7.44 (m, 2H), 7.43 – 7.33 (m, 2H), 7.30 – 7.19 (m, 2H), 7.01 – 6.90 (m, 1H), 6.81 (t, $J$ = 5.5 Hz, 1H), 4.66 (s, 2H), 3.33 (s, 2H), 2.97 (q, $J$ = 6.4 Hz, 2H), 1.66 (p, $J$ = 6.9 Hz, 2H), 1.36 (s, 9H). $^{13}$C NMR (101 MHz, DMSO-$d_6$) δ 167.63, 156.07, 155.69, 144.64, 140.80, 129.99, 129.90, 128.68, 127.57, 122.44, 120.69, 77.99, 49.67, 44.66, 37.99, 28.68, 28.63. HRMS (ESI) $m/z$ calculated 428.2185 for $C_{23}H_{30}N_3O_5$ $[M+H]^+$, found 428.2193.

Display Report

**Analysis Info**

| | | | |
|---|---|---|---|
| Analysis Name | Z:\Data\2024\2408\sam290824\VTA20_32_01_127304.d | Acquisition Date | 29.08.2024 22:09:58 |
| Method | hystar_pl.m | Operator | BDAL@DE |
| Sample Name | VTA20 | Instrument / Ser# | micrOTOF 10237 |
| Comment | | | |

**Acquisition Parameter**

| | | | | | |
|---|---|---|---|---|---|
| Source Type | ESI | Ion Polarity | Positive | Set Nebulizer | 1.2 Bar |
| Focus | Not active | | | Set Dry Heater | 180 °C |
| Scan Begin | 50 m/z | Set Capillary | 4500 V | Set Dry Gas | 4.0 l/min |
| Scan End | 1600 m/z | Set End Plate Offset | -500 V | Set Divert Valve | Source |

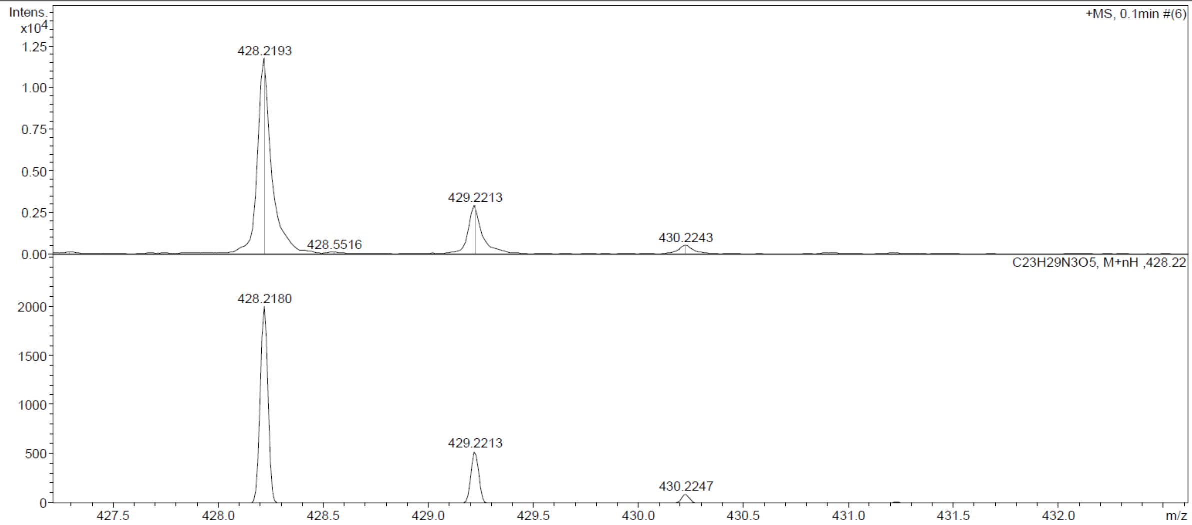

Compound **14b:**

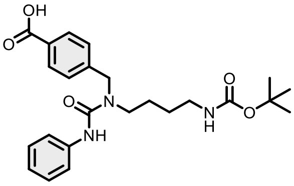

Evaporated ethyl acetate to give 643 mg of targeted compound in 83% yield as a white solid by general method B.

$^{1}$H NMR (400 MHz, DMSO-$d_6$) δ 12.53 (s, 1H), 8.37 (s, 1H), 7.94 (d, $J$ = 8.0 Hz, 2H), 7.49 (d, $J$ = 8.0 Hz, 2H), 7.38 (d, $J$ = 8.2 Hz, 2H), 7.23 (t, $J$ = 7.9 Hz, 2H), 6.95 (t, $J$ = 7.3 Hz, 1H), 6.81 (t, $J$ = 5.9 Hz, 1H), 4.66 (s, 2H), 3.32 (d, $J$ = 7.6 Hz, 2H), 2.92 (q, $J$ = 6.5 Hz, 2H), 1.51 (dq, $J$ = 14.6, 7.3 Hz, 2H), 1.36 (s, 11H). $^{13}$C NMR (101 MHz, DMSO-$d_6$) δ 172.48, 167.65, 156.13, 155.66, 144.78, 140.86, 129.96, 129.87, 128.65, 127.58, 122.36, 120.65, 77.84, 49.58, 46.68, 28.70, 27.37, 25.60. HRMS (ESI) $m/z$ calculated 442.2341 for $C_{24}H_{32}N_3O_5$ [M+H]$^+$, found 442.2346.

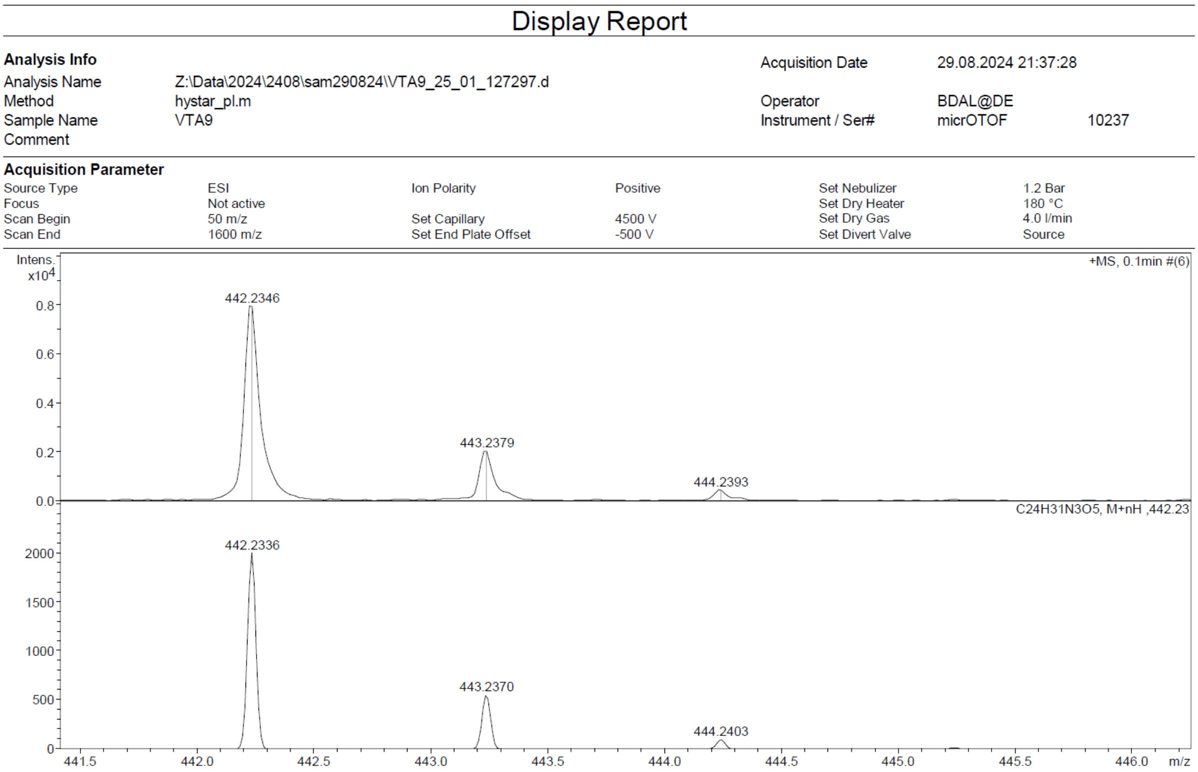

Compound **15b:**

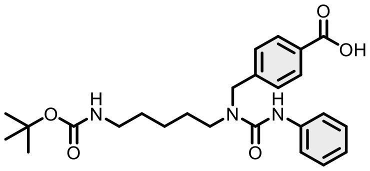

Evaporated ethyl acetate to give 706 mg of targeted compound in 91% yield as a white solid by general method B.

$^{1}$H NMR (400 MHz, DMSO-$d_6$) δ 8.39 (s, 1H), 7.91 (d, $J$ = 7.8 Hz, 2H), 7.57 – 7.42 (m, 2H), 7.32 (d, $J$ = 8.0 Hz, 2H), 7.23 (t, $J$ = 7.9 Hz, 2H), 6.94 (t, $J$ = 7.3 Hz, 1H), 6.77 (t, $J$ = 5.8 Hz, 1H), 4.64 (s, 2H), 3.29 (s, 2H), 2.89 (q, $J$ = 6.6 Hz, 2H), 1.51 (dq, $J$ = 15.1, 7.3 Hz, 2H), 1.37 (s, 11H), 1.22 (td, $J$ = 8.3, 4.0 Hz, 2H). $^{13}$C NMR (101 MHz, DMSO-$d_6$) δ 168.76, 156.07, 155.68, 143.15, 140.95, 132.99, 129.87, 128.65, 127.24, 122.28, 120.61, 77.77, 49.53, 46.70, 29.76, 28.73, 27.86, 23.93, 21.89. HRMS (ESI) $m/z$ calculated 456.2498 for $C_{25}H_{34}N_3O_5$ $[M+H]^+$, found 456.2495.

#### Display Report

**Analysis Info**

| | | | |
|---|---|---|---|
| Analysis Name | Z:\Data\2024\2408\sam290824\VTA10_26_01_127298.d | Acquisition Date | 29.08.2024 21:42:06 |
| Method | hystar_pl.m | Operator | BDAL@DE |
| Sample Name | VTA10 | Instrument / Ser# | micrOTOF 10237 |
| Comment | | | |

**Acquisition Parameter**

| | | | | | |
|---|---|---|---|---|---|
| Source Type | ESI | Ion Polarity | Positive | Set Nebulizer | 1.2 Bar |
| Focus | Not active | | | Set Dry Heater | 180 °C |
| Scan Begin | 50 m/z | Set Capillary | 4500 V | Set Dry Gas | 4.0 l/min |
| Scan End | 1600 m/z | Set End Plate Offset | -500 V | Set Divert Valve | Source |

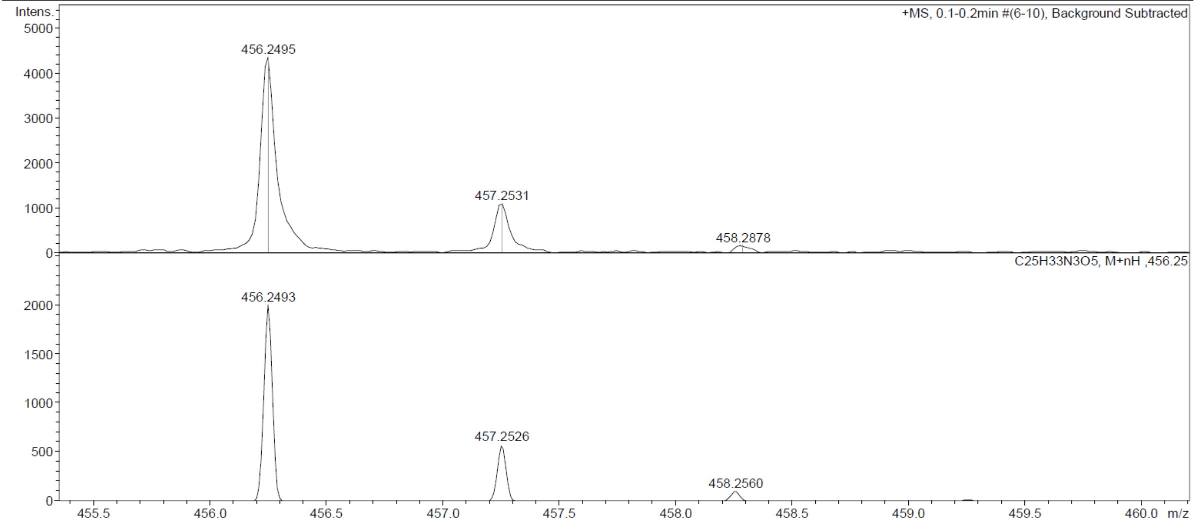

Compound **16b:**

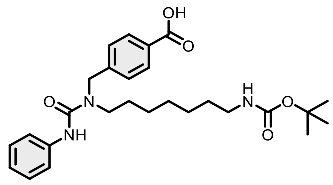

Evaporated ethyl acetate to give 559 mg of targeted compound in 72% yield as a white solid by general

$^{1}$H NMR (400 MHz, DMSO-$d_6$) δ 12.52 (s, 1H), 8.37 (s, 1H), 7.99 – 7.84 (m, 2H), 7.50 – 7.44 (m, 2H), 7.41 – 7.35 (m, 2H), 7.27 – 7.19 (m, 2H), 7.00 – 6.88 (m, 1H), 6.74 (t, $J$ = 5.8 Hz, 1H), 4.65 (s, 2H), 3.31 (t, $J$ = 7.5 Hz, 3H), 2.88 (q, $J$ = 6.6 Hz, 2H), 1.50 (dq, $J$ = 13.8, 6.1 Hz, 2H), 1.36 (s, 11H), 1.21 (dq, $J$ = 10.8, 5.7 Hz, 6H). $^{13}$C NMR (101 MHz, DMSO-$d_6$) δ 167.64, 156.03, 155.69, 144.80, 140.91, 129.96, 129.88, 128.67, 127.62, 122.32, 120.57, 77.73, 49.60, 46.91, 29.89, 29.02, 28.73, 28.20, 26.72, 26.65, 21.51. HRMS (ESI) *m/z* calculated 484.2811 for $C_{27}H_{38}N_3O_5$ $[M+H]^+$, found 484.2825.

#### Display Report

**Analysis Info**

| | | | |
|---|---|---|---|
| Analysis Name | Z:\Data\2024\2408\sam290824\VTA11_27_01_127299.d | Acquisition Date | 29.08.2024 21:46:44 |
| Method | hystar_pl.m | Operator | BDAL@DE |
| Sample Name | VTA11 | Instrument / Ser# | micrOTOF 10237 |
| Comment | | | |

**Acquisition Parameter**

| | | | | | |
|---|---|---|---|---|---|
| Source Type | ESI | Ion Polarity | Positive | Set Nebulizer | 1.2 Bar |
| Focus | Not active | | | Set Dry Heater | 180 °C |
| Scan Begin | 50 m/z | Set Capillary | 4500 V | Set Dry Gas | 4.0 l/min |
| Scan End | 1600 m/z | Set End Plate Offset | -500 V | Set Divert Valve | Source |

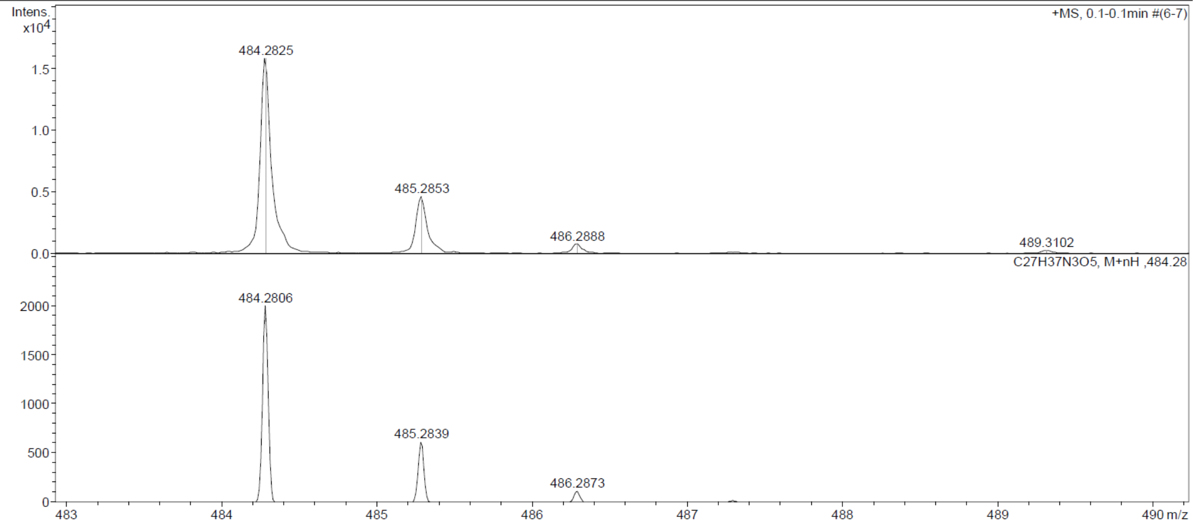

Compound **17b:**

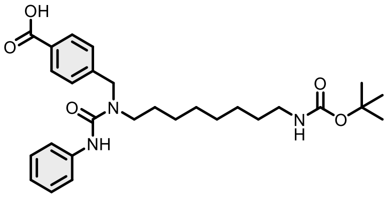

Evaporated ethyl acetate to give 622 mg of targeted compound in 80% yield as a white solid by general method B.

$^{1}$H NMR (400 MHz, DMSO-$d_6$) δ 12.81 (s, 1H), 8.38 (s, 1H), 8.00 – 7.88 (m, 2H), 7.52 – 7.44 (m, 2H), 7.39 (d, $J$ = 8.1 Hz, 2H), 7.27 – 7.19 (m, 2H), 6.99 – 6.89 (m, 1H), 6.73 (t, $J$ = 5.7 Hz, 1H), 4.66 (s, 2H), 3.31 (t, $J$ = 7.5 Hz, 2H), 2.89 (q, $J$ = 6.6 Hz, 2H), 1.50 (h, $J$ = 5.5 Hz, 2H), 1.37 (s, 11H), 1.26 – 1.18 (m, 8H). $^{13}$C NMR (101 MHz, DMSO-$d_6$) δ 167.65, 156.03, 155.70, 144.78, 141.08, 140.91, 129.96, 129.89, 128.65, 127.62, 122.30, 120.56, 77.71, 49.62, 46.92, 29.93, 29.29, 29.18, 28.72, 28.24, 26.69, 26.62. HRMS (ESI) *m*/*z* calculated 498.2967 for $C_{28}H_{40}N_3O_5$, found 498.2974 ([M+H]$^+$).

#### Display Report

**Analysis Info**

| | | | |
|---|---|---|---|
| Analysis Name | Z:\Data\2024\2408\sam290824\VTA12_28_01_127300.d | Acquisition Date | 29.08.2024 21:51:23 |
| Method | hystar_pl.m | Operator | BDAL@DE |
| Sample Name | VTA12 | Instrument / Ser# | micrOTOF 10237 |
| Comment | | | |

**Acquisition Parameter**

| | | | | | |
|---|---|---|---|---|---|
| Source Type | ESI | Ion Polarity | Positive | Set Nebulizer | 1.2 Bar |
| Focus | Not active | | | Set Dry Heater | 180 °C |
| Scan Begin | 50 m/z | Set Capillary | 4500 V | Set Dry Gas | 4.0 l/min |
| Scan End | 1600 m/z | Set End Plate Offset | -500 V | Set Divert Valve | Source |

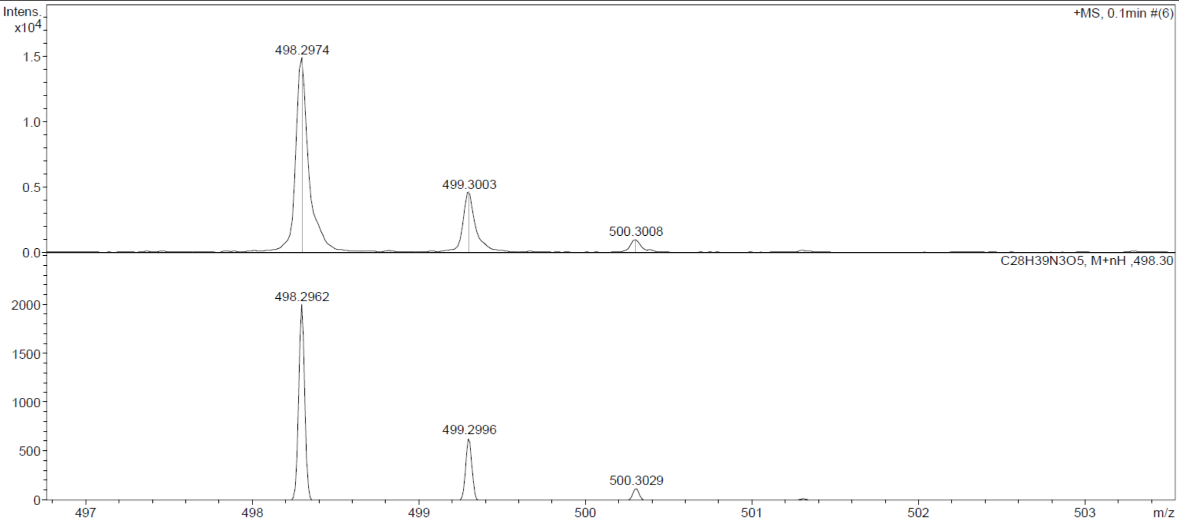

Compound **18b:**

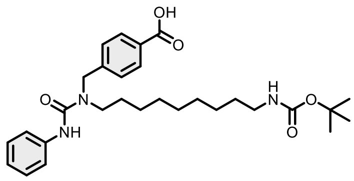

Evaporated ethyl acetate to give 167 mg of targeted compound in 86% yield as a white solid by general method B.

$^{1}$H NMR (400 MHz, DMSO-$d_6$) δ 12.61 (s, 1H), 8.38 (s, 1H), 8.00 – 7.83 (m, 2H), 7.50 – 7.44 (m, 2H), 7.41 – 7.35 (m, 2H), 7.27 – 7.18 (m, 2H), 6.94 (tt, *J* = 7.3, 1.2 Hz, 1H), 6.73 (t, *J* = 5.7 Hz, 1H), 4.65 (s, 2H), 3.31 (t, *J* = 7.5 Hz, 2H), 2.88 (q, *J* = 6.7 Hz, 2H), 1.49 (p, *J* = 5.7 Hz, 2H), 1.37 (s, 11H), 1.26 – 1.17 (m, 10H). $^{13}$C NMR (101 MHz, DMSO-$d_6$) δ 167.64, 156.03, 155.70, 144.82, 140.91, 129.96, 129.87, 128.67, 127.64, 122.31, 120.55, 77.72, 49.63, 46.93, 29.93, 29.43, 29.23, 29.11, 28.73, 28.24, 26.70, 26.63, 21.52. HRMS (ESI) *m*/*z* calculated 512.3124 for $C_{29}H_{42}N_3O_5$ [M+H]$^+$, found 512.3129.

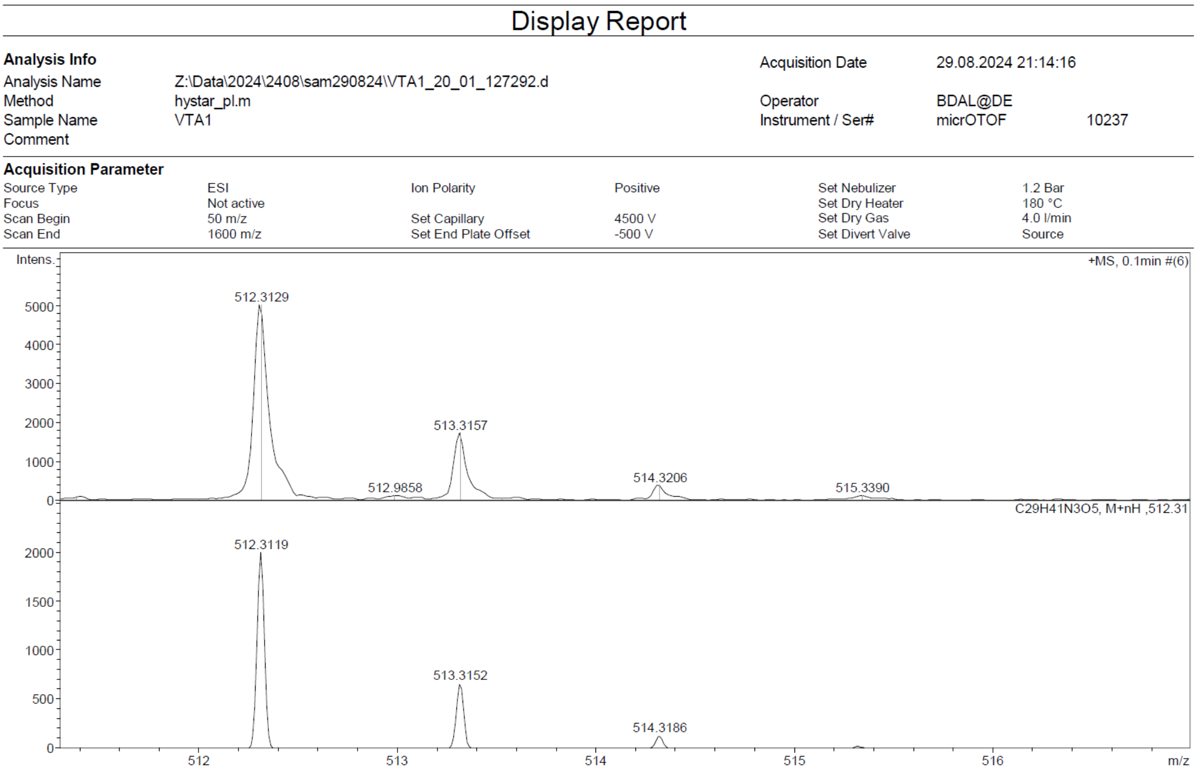

Compound **19b**:

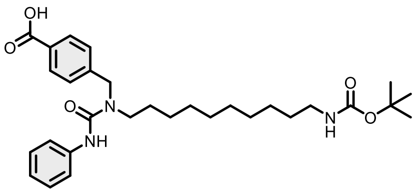

Evaporated ethyl acetate to give 190 mg of targeted compound in 65% yield as a white solid by general method B.

$^{1}$H NMR (400 MHz, DMSO-$d_6$) δ 8.34 (s, 1H), 7.57 – 7.44 (m, 2H), 7.30 (d, $J$ = 8.0 Hz, 2H), 7.27 – 7.19 (m, 4H), 6.98 – 6.90 (m, 1H), 6.74 (t, $J$ = 5.7 Hz, 1H), 4.50 (s, 2H), 3.29 (s, 2H), 2.91 (q, $J$ = 6.6 Hz, 2H), 1.51 (q, $J$ = 6.9 Hz, 2H), 1.38 (s, 11H), 1.23 (d, $J$ = 4.7 Hz, 12H). $^{13}$C NMR (101 MHz, DMSO-$d_6$) δ 172.46, 156.05, 155.71, 141.69, 141.06, 137.62, 128.63, 127.44, 127.02, 122.15, 120.44, 77.70, 63.20, 49.42, 46.45, 29.98, 29.45, 29.33, 29.22, 28.72, 28.15, 26.77, 26.72, 21.49. HRMS (ESI) $m/z$ calculated 526.3275 for $C_{30}H_{44}N_3O_5$, found 526.3279.

#### Display Report

**Analysis Info**

| | | | |
|---|---|---|---|
| Analysis Name | Z:\Data\2024\2409\sam300924\VTA13_15_01_127692.d | Acquisition Date | 30.09.2024 14:00:24 |
| Method | hystar_pl.m | Operator | BDAL@DE |
| Sample Name | VTA13 | Instrument / Ser# | micrOTOF 10237 |
| Comment | | | |

**Acquisition Parameter**

| | | | | | |
|---|---|---|---|---|---|
| Source Type | ESI | Ion Polarity | Positive | Set Nebulizer | 1.2 Bar |
| Focus | Not active | | | Set Dry Heater | 180 °C |
| Scan Begin | 50 m/z | Set Capillary | 4500 V | Set Dry Gas | 4.0 l/min |
| Scan End | 1600 m/z | Set End Plate Offset | -500 V | Set Divert Valve | Source |

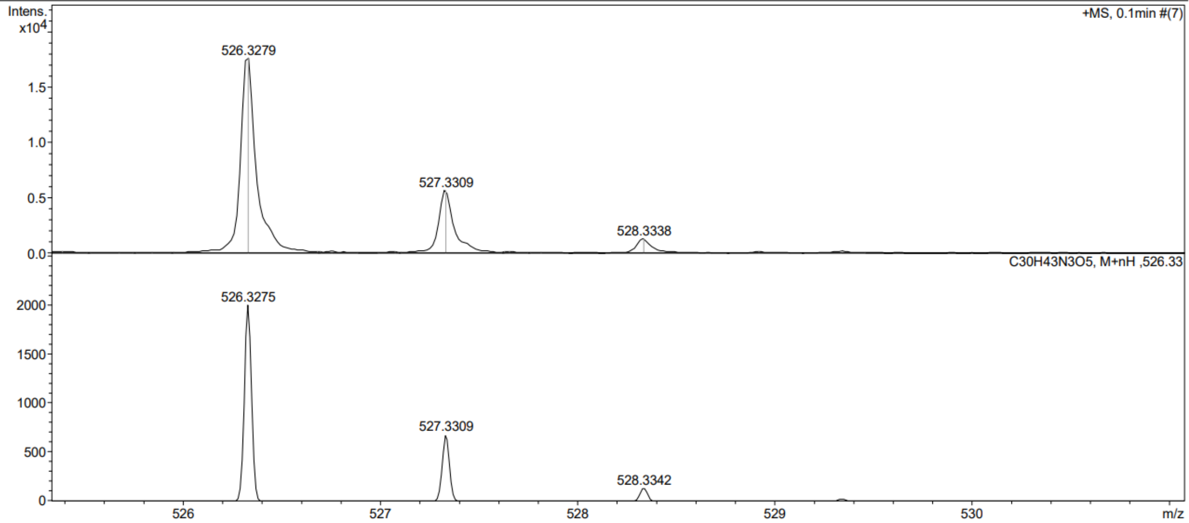

Bruker Compass DataAnalysis 4.0 printed: 30.09.2024 13:57:41 Page 1 of 1

Compound **20b:**

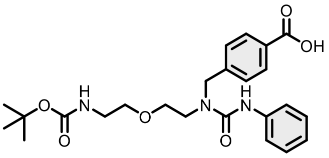

Evaporated ethyl acetate to give 636 mg of targeted compound in 82% yield as a white solid by general method B.

$^{1}$H NMR (400 MHz, DMSO-$d_6$) δ 12.61 (s, 1H), 8.42 (s, 1H), 8.01 – 7.87 (m, 2H), 7.50 – 7.35 (m, 4H), 7.29 – 7.18 (m, 2H), 7.00 – 6.90 (m, 1H), 6.84 (t, $J$ = 5.7 Hz, 1H), 4.70 (s, 2H), 3.59 – 3.47 (m, 4H), 3.42 (t, $J$ = 5.9 Hz, 2H), 3.12 (q, $J$ = 5.9 Hz, 2H), 1.37 (s, 9H). $^{13}$C NMR (101 MHz, DMSO-$d_6$) δ 167.66, 156.13, 156.01, 144.68, 140.77, 129.95, 129.89, 128.77, 127.66, 122.31, 120.18, 78.10, 69.89, 69.54, 60.21, 50.37, 47.03, 28.67. HRMS (ESI) $m/z$ calculated 458.2291 for $C_{24}H_{32}N_3O_6$ [M+H]$^+$, found 458.2295.

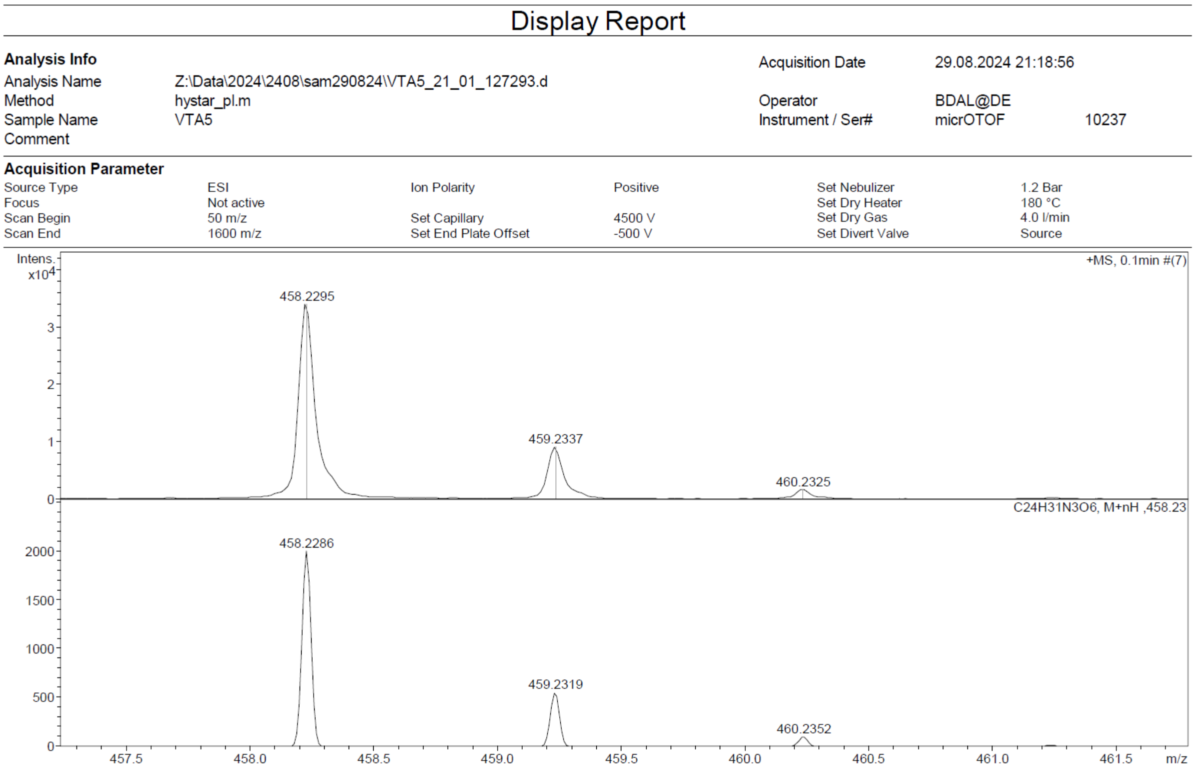

Compound **21b:**

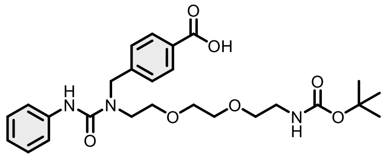

Evaporated ethyl acetate to give 723 mg of targeted compound in 93% yield as a white solid by general method B.

$^{1}$H NMR (400 MHz, DMSO-$d_6$) δ 12.58 (s, 1H), 8.49 (s, 1H), 8.01 – 7.87 (m, 2H), 7.42 (ddd, $J$ = 15.8, 7.7, 1.5 Hz, 4H), 7.31 – 7.18 (m, 2H), 6.95 (tt, $J$ = 7.2, 1.2 Hz, 1H), 6.74 (t, $J$ = 5.7 Hz, 1H), 4.69 (s, 2H), 3.61 – 3.48 (m, 8H), 3.39 (t, $J$ = 6.2 Hz, 2H), 3.07 (q, $J$ = 6.1 Hz, 2H), 1.36 (s, 9H). $^{13}$C NMR (101 MHz, DMSO-$d_6$) δ 167.66, 156.12, 156.05, 144.65, 140.78, 129.95, 129.89, 128.81, 127.69, 122.26, 119.97, 78.05, 70.44, 70.01, 69.90, 69.68, 60.21, 50.34, 47.20, 28.66. HRMS (ESI) $m/z$ calculated 502.2553 for $C_{26}H_{36}N_3O_7$ [M+H]$^+$, found 502.2566.

#### Display Report

**Analysis Info**

| | | | |
|---|---|---|---|
| Analysis Name | Z:\Data\2024\2408\sam290824\VTA6_22_01_127294.d | Acquisition Date | 29.08.2024 21:23:33 |
| Method | hystar_pl.m | Operator | BDAL@DE |
| Sample Name | VTA6 | Instrument / Ser# | micrOTOF 10237 |
| Comment | | | |

**Acquisition Parameter**

| | | | | | |
|---|---|---|---|---|---|
| Source Type | ESI | Ion Polarity | Positive | Set Nebulizer | 1.2 Bar |
| Focus | Not active | | | Set Dry Heater | 180 °C |
| Scan Begin | 50 m/z | Set Capillary | 4500 V | Set Dry Gas | 4.0 l/min |
| Scan End | 1600 m/z | Set End Plate Offset | -500 V | Set Divert Valve | Source |

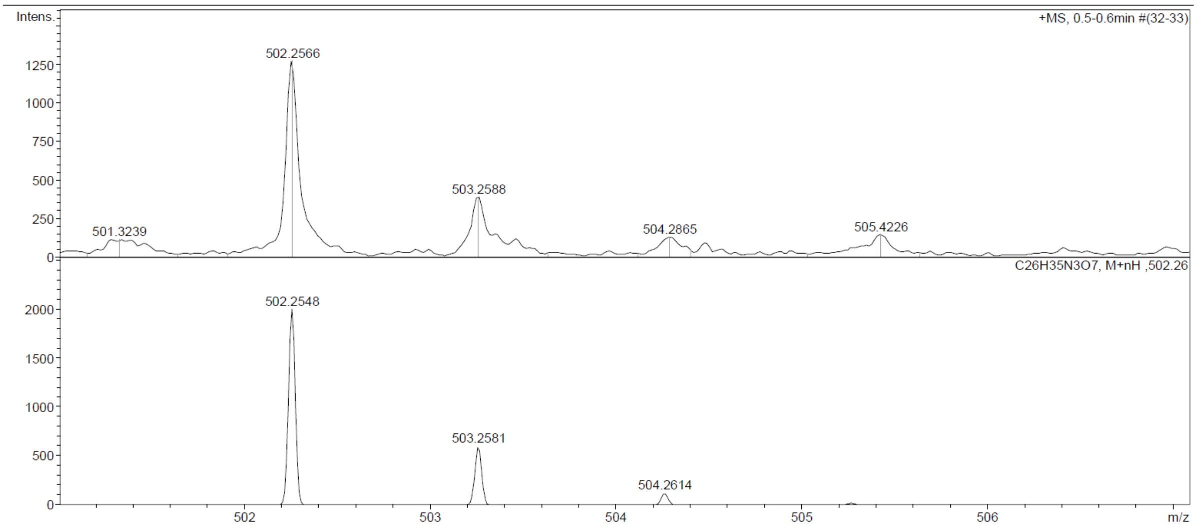

Compound **22b:**

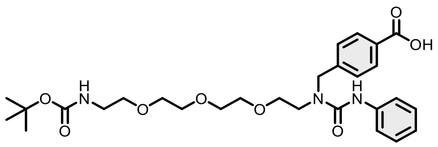

Evaporated ethyl acetate to give 327 mg of targeted compound in 42% yield as a white solid by general method B.

$^1$H NMR (400 MHz, DMSO-$d_6$) δ 12.56 (s, 1H), 8.49 (s, 1H), 8.00 – 7.87 (m, 2H), 7.49 – 7.35 (m, 4H), 7.30 – 7.19 (m, 2H), 6.95 (tt, $J$ = 7.3, 1.2 Hz, 1H), 6.72 (t, $J$ = 5.8 Hz, 1H), 4.68 (s, 2H), 3.59 – 3.45 (m, 12H), 3.36 (s, 2H), 3.05 (q, $J$ = 6.0 Hz, 2H), 1.37 (s, 9H). $^{13}$C NMR (101 MHz, DMSO-$d_6$) δ 167.66, 156.13, 156.03, 144.68, 140.80, 129.95, 129.88, 128.80, 127.68, 122.25, 119.96, 119.84, 78.03, 70.48, 70.20, 70.18, 70.00, 69.92, 69.64, 50.34, 47.26, 28.67. HRMS (ESI) $m/z$ calculated 546.2815 for $C_{28}H_{40}N_3O_8$ [M+H]$^+$, found 546.2812.

Display Report

**Analysis Info**

| | | | |
|---|---|---|---|
| Analysis Name | Z:\Data\2024\2408\sam290824\VTA7_23_01_127295.d | Acquisition Date | 29.08.2024 21:28:11 |
| Method | hystar_pl.m | Operator | BDAL@DE |
| Sample Name | VTA7 | Instrument / Ser# | micrOTOF 10237 |
| Comment | | | |

**Acquisition Parameter**

| | | | | | |
|---|---|---|---|---|---|
| Source Type | ESI | Ion Polarity | Positive | Set Nebulizer | 1.2 Bar |
| Focus | Not active | | | Set Dry Heater | 180 °C |
| Scan Begin | 50 m/z | Set Capillary | 4500 V | Set Dry Gas | 4.0 l/min |
| Scan End | 1600 m/z | Set End Plate Offset | -500 V | Set Divert Valve | Source |

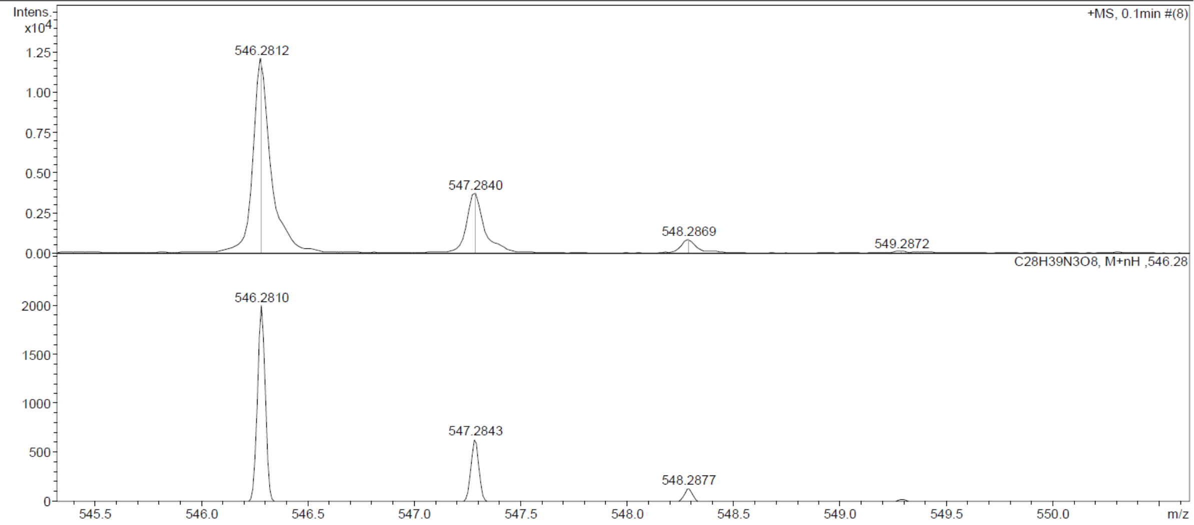

Compound **23b:**

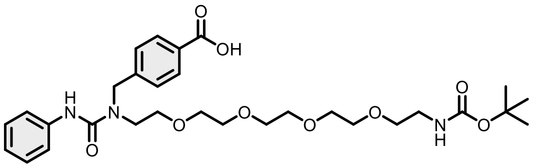

Evaporated ethyl acetate to give 387 mg of targeted compound in 66% yield as a white solid by general method B.

$^{1}$H NMR (400 MHz, DMSO-$d_6$) δ 12.50 (s, 1H), 8.49 (s, 1H), 8.00 – 7.85 (m, 2H), 7.52 – 7.34 (m, 4H), 7.32 – 7.19 (m, 2H), 7.00 – 6.88 (m, 1H), 6.73 (t, $J$ = 5.8 Hz, 1H), 4.68 (s, 2H), 3.59 – 3.46 (m, 16H), 3.36 (s, 2H), 3.06 (q, $J$ = 6.0 Hz, 2H), 1.37 (s, 9H). $^{13}$C NMR (101 MHz, DMSO-$d_6$) δ 172.48, 167.65, 156.12, 156.03, 144.69, 140.80, 129.94, 129.87, 128.80, 127.69, 122.24, 119.95, 78.03, 70.48, 70.26, 70.20, 70.18, 69.99, 69.93, 69.62, 50.34, 47.27, 41.33, 28.67. HRMS (ESI) $m/z$ calculated 590.3077 for $C_{30}H_{44}N_3O_9$ $[M+H]^+$, found 590.3085.

#### Display Report

**Analysis Info**

| | | | | |
|---|---|---|---|---|
| Analysis Name | Z:\Data\2024\2408\sam290824\VTA8_24_01_127296.d | Acquisition Date | 29.08.2024 21:32:49 | |
| Method | hystar_pl.m | Operator | BDAL@DE | |
| Sample Name | VTA8 | Instrument / Ser# | micrOTOF | 10237 |
| Comment | | | | |

**Acquisition Parameter**

| | | | | | |
|---|---|---|---|---|---|
| Source Type | ESI | Ion Polarity | Positive | Set Nebulizer | 1.2 Bar |
| Focus | Not active | | | Set Dry Heater | 180 °C |
| Scan Begin | 50 m/z | Set Capillary | 4500 V | Set Dry Gas | 4.0 l/min |
| Scan End | 1600 m/z | Set End Plate Offset | -500 V | Set Divert Valve | Source |

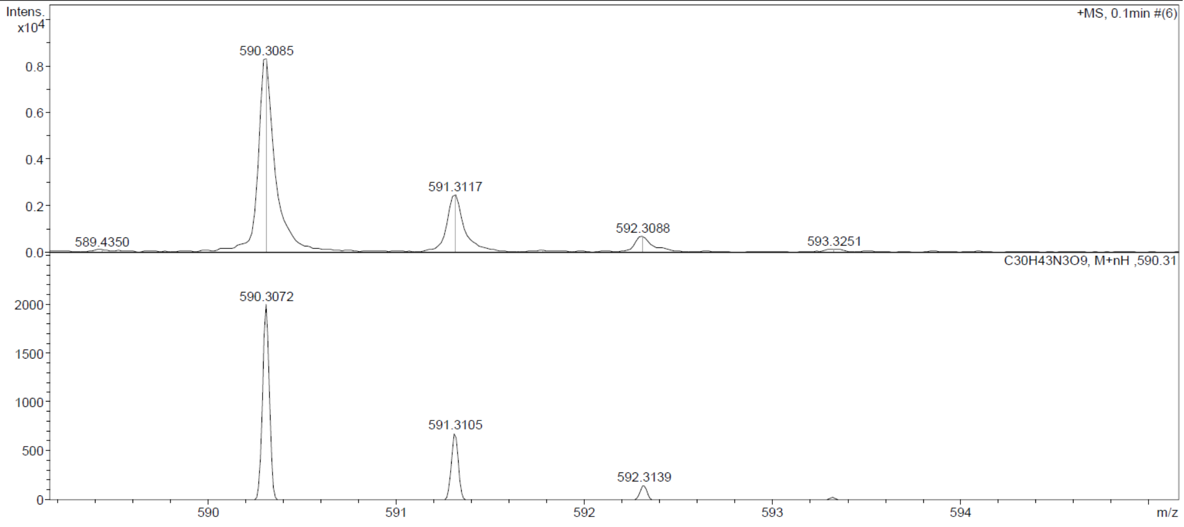

Compound **1c**:

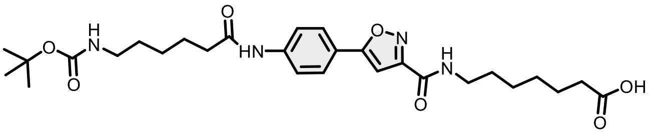

Compound **1b** was dissolved in solvent combination of MeOH: THF: $H_2O$ (1:1:1). To this solution, LiOH (2 equiv) was added, and the resulting mixture was stirred at room temperature overnight. The reaction was acidified with 1 N HCl and the aqueous layer extracted with Ethyl acetate (3 x 10mL). The combined organic extracts were washed with brine, dried over sodium sulfate, concentrated in vacuo and use for the next step without purification.

To a solution of compound from previous step (1g, 2.25 mmol) and Methyl 7-aminoheptanoate hydrochloride (528 mg, 2.7 mmol) in DMF (20 mL) and DIPEA (10 mmol) was added HATU (3 mmol) then the mixture was stirred at room temperature overnight. Saturated ammonium chloride solution was added to the solution and extracted with DCM. The combined organic layers were washed with brine and dried over Na2SO4 and concentrated in vacuo. The crude was dissolve in solvent combination of MeOH: THF: $H_2O$ (1:1:1). To this solution, LiOH (3 equiv) was added, and the resulting mixture was stirred at room temperature overnight. The reaction was acidified with 1 N HCl and the aqueous layer extracted with Ethyl acetate (3 x 10mL). The combined organic extracts were washed with brine, dried over sodium sulfate, concentrated in vacuo, and purified by column chromatography to afford **1c** (4.6g, 56%).

$^1$H NMR (400 MHz, DMSO-$d_6$) δ 11.95 (s, 1H), 10.13 (s, 1H), 8.81 – 8.67 (m, 1H), 7.88 – 7.80 (m, 2H), 7.80 – 7.72 (m, 2H), 7.18 (s, 1H), 6.74 (t, *J* = 5.7 Hz, 1H), 3.31 – 3.17 (m, 2H), 2.90 (q, *J* = 6.6 Hz, 2H), 2.32 (t, *J* = 7.4 Hz, 2H), 2.18 (t, *J* = 7.4 Hz, 2H), 1.66 – 1.39 (m, 6H), 1.34 (s, 9H), 1.32 – 1.19 (m, 6H). $^{13}$C NMR (101 MHz, DMSO-$d_6$) δ 174.91, 172.10, 170.66, 160.11, 158.89, 156.02, 151.53, 141.97, 129.27, 126.97, 121.27, 121.13, 119.58, 99.08, 77.73, 36.91, 34.06, 29.78, 29.18, 28.69, 26.56, 26.44, 25.20, 24.89. (ESI) *m*/*z* calculated 545.2975 for $C_{28}H_{41}N_4O_7$ $[M+H]^+$, found 545.2972.

E:\2025\Oktober\231025\VT812.raw 10/23/2025 12:36:26 PM

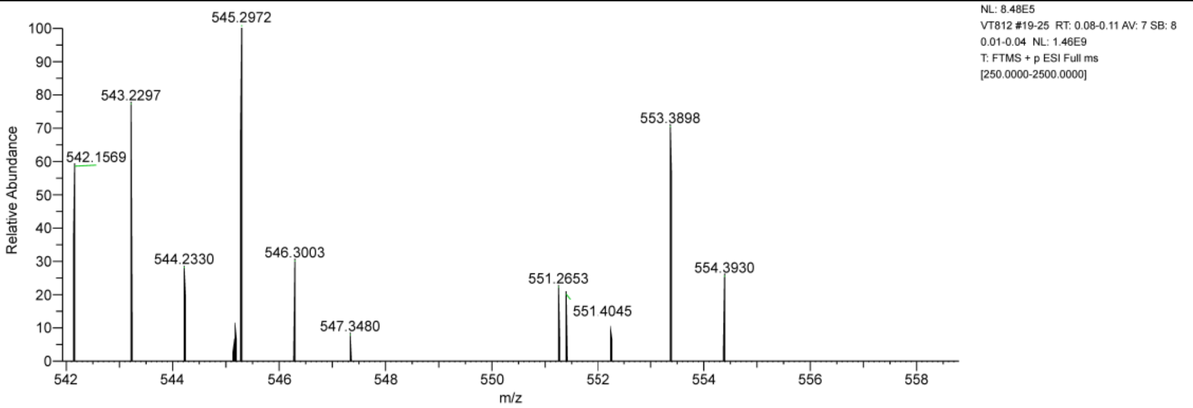

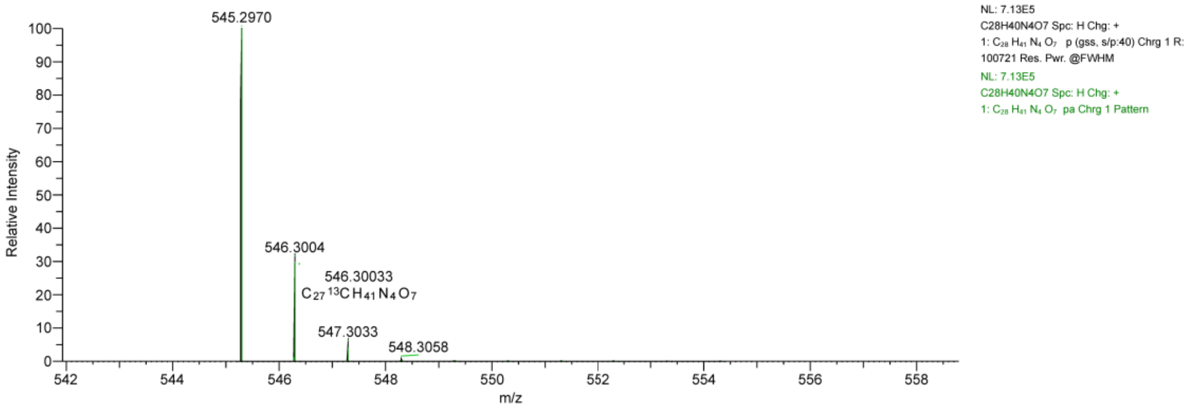

Probe **1**:

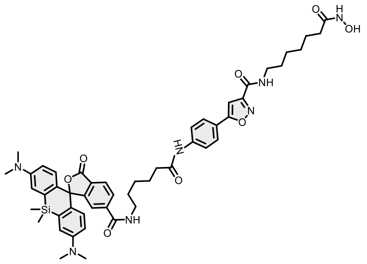

Reaction was conducted and purified following general method C. Fractions containing the product were collected, evaporated and lyophilized from acetonitrile: water mixture. The obtained solid was dissolved in 700 µL of $d_6$-DMSO. The samples from DMSO solution were diluted x100 in PBS +0.1%SDS and concentration was determined spectroscopically with Nanodrop. Determined stock concentration was 1.58 mM which constitutes to 26% yield (1.2 mg).

$^1$H NMR (400 MHz, DMSO-$d_6$) δ 10.31 (s, 1H), 10.16 (s, 1H), 8.76 (dt, *J* = 11.6, 5.7 Hz, 2H), 8.64 (s, 1H), 8.39 (dd, *J* = 1.6, 0.7 Hz, 1H), 8.20 (dd, *J* = 8.1, 1.6 Hz, 1H), 7.87 – 7.81 (m, 2H), 7.78 – 7.73 (m, 2H), 7.33 (dd, *J* = 8.1, 0.7 Hz, 1H), 7.16 (s, 1H), 7.00 (dd, *J* = 2.1, 1.2 Hz, 2H), 6.61 – 6.60 (m, 3H), 3.23 (q, *J* = 6.9 Hz, 2H), 2.91 (s, 12H), 2.36 (t, *J* = 7.4 Hz, 2H), 1.93 (t, *J* = 7.4 Hz, 2H), 1.62 (dp, *J* = 22.3, 7.3 Hz, 5H), 1.48 (p, *J* = 7.5 Hz, 4H), 1.39 (q, *J* = 8.0 Hz, 2H), 1.33 – 1.20 (m, 5H), 0.62 (s, 3H), 0.51 (s, 3H). HRMS (ESI) *m*/*z* calculated 914.4272 for $C_{50}H_{60}N_7O_8Si$ $[M+H]^+$, found 914.4269.

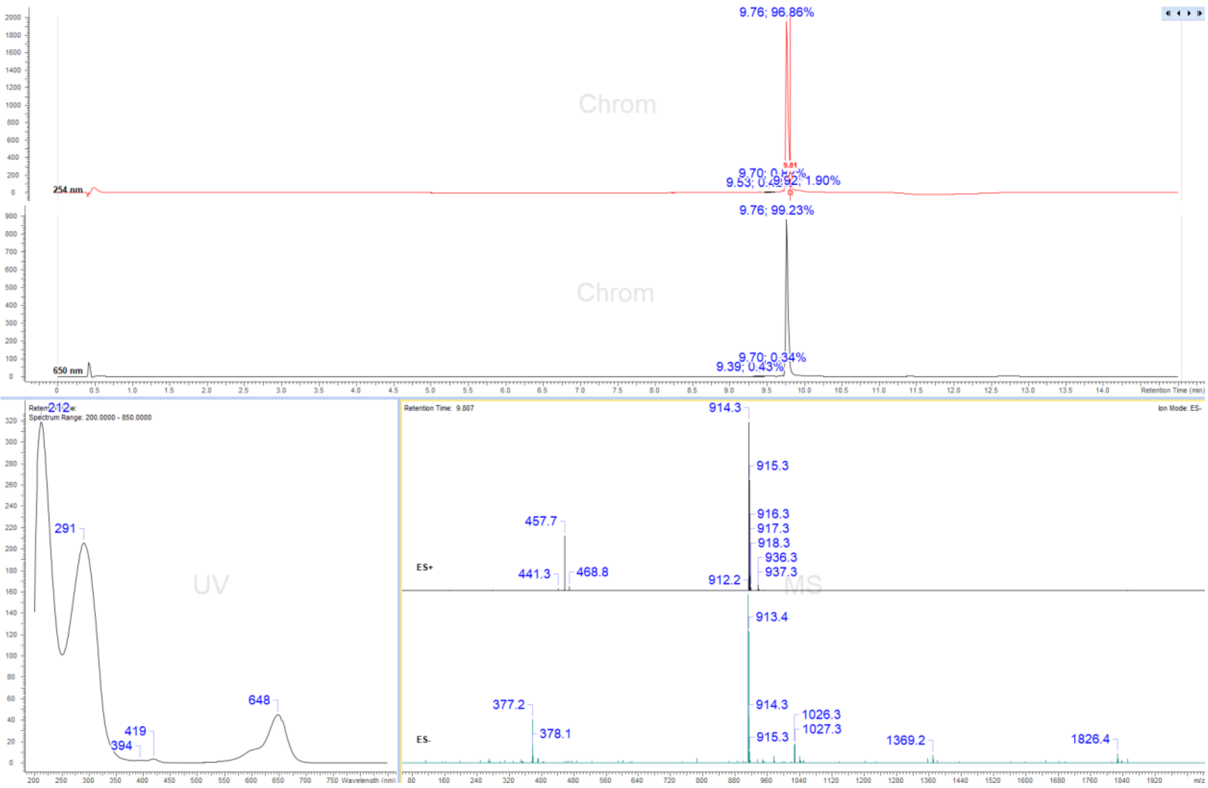

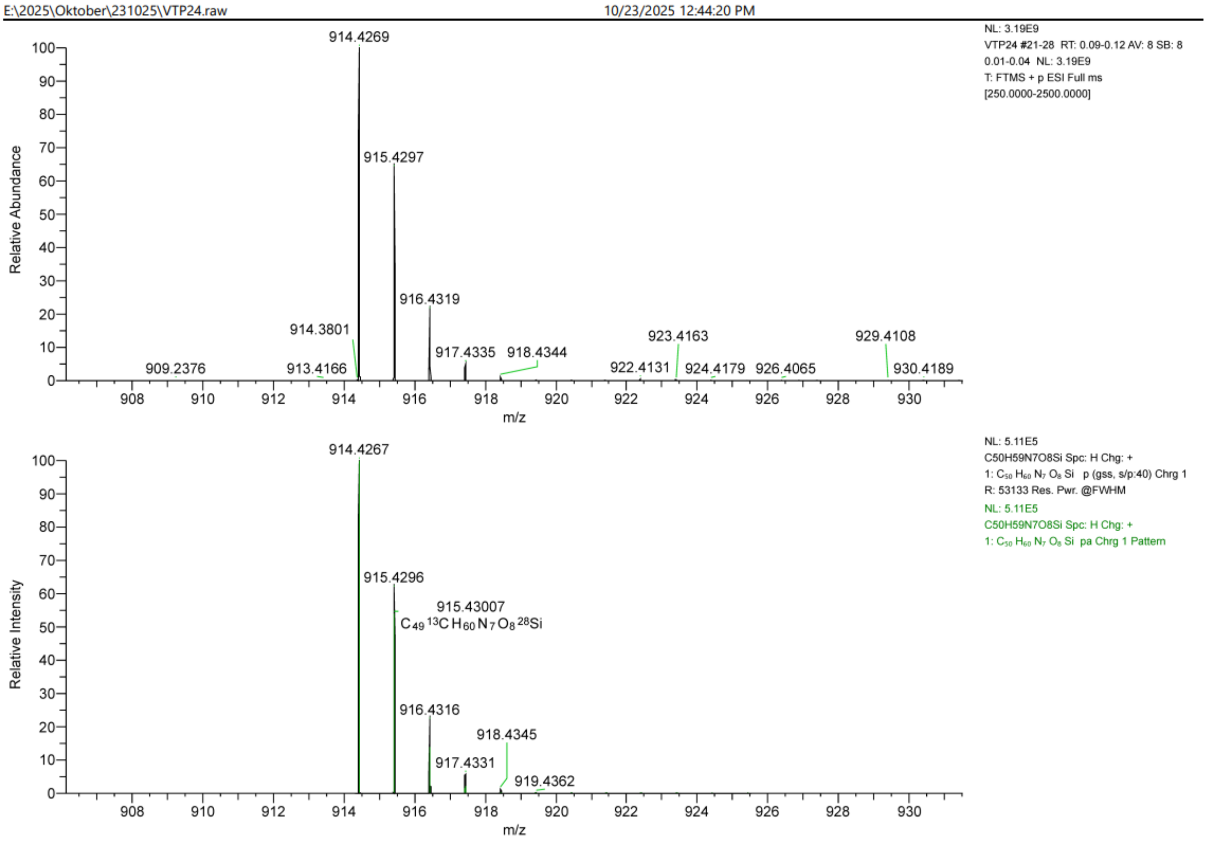
E:\2025\Oktober\231025\VTP24.raw
10/23/2025 12:44:20 PM
NL: 3.19E9
VTP24 #21-28 RT: 0.09-0.12 AV: 8 SB: 8 0.01-0.04 NL: 3.19E9
T: FTMS + p ESI Full ms [250.0000-2500.0000]
Relative Abundance
914.4269
915.4297
916.4319
914.3801
917.4335
918.4344
923.4163
929.4108
909.2376
913.4166
922.4131
924.4179
926.4065
930.4189
m/z
NL: 5.11E5
C50H59N7O8Si Spc: H Chg: +
1: C50 H60 N7 O8 Si p (gss, s/p:40) Chrg 1
R: 53133 Res. Pwr. @FWHM
NL: 5.11E5
C50H59N7O8Si Spc: H Chg: +
1: C50 H60 N7 O8 Si pa Chrg 1 Pattern
Relative Intensity
914.4267
915.4296
915.43007
C49 13C H60 N7 O8 28Si
916.4316
918.4345
917.4331
919.4362
m/z

Probe **2**:

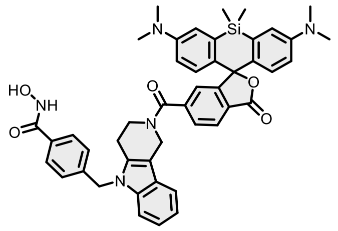

Reaction was conducted and purified following general method C. Fractions containing the product were collected, evaporated and lyophilized from acetonitrile: water mixture. The obtained solid was dissolved in 700 µL of $d_6$-DMSO. The samples from DMSO solution were diluted x100 in PBS +0.1%SDS and concentration was determined spectroscopically with Nanodrop. Determined stock concentration was 1.27 mM which constitutes 21% yield (0.69 mg).

$^1$H NMR (400 MHz, DMSO-$d_6$) δ 11.07 (s, 1H), 8.99 (s, 1H), 7.98 (s, 1H), 7.84 (dd, $J$ = 7.9, 1.6 Hz, 1H), 7.69 – 7.63 (m, 2H), 7.55 (s, 1H), 7.39 (d, $J$ = 8.1 Hz, 1H), 7.33 (dd, $J$ = 7.9, 0.8 Hz, 1H), 7.17 – 7.03 (m, 4H), 7.01 (d, $J$ = 2.7 Hz, 2H), 6.71 – 6.63 (m, 4H), 5.42 (s, 2H), 4.88 (s, 2H), 3.73 (s, 2H), 2.92 (s, 14H), 0.62 (s, 3H), 0.52 (s, 3H). HRMS (ESI) $m/z$ calculated 776.3268 for $C_{46}H_{46}N_5O_5Si$ [M+H]$^+$, found 776.3266.

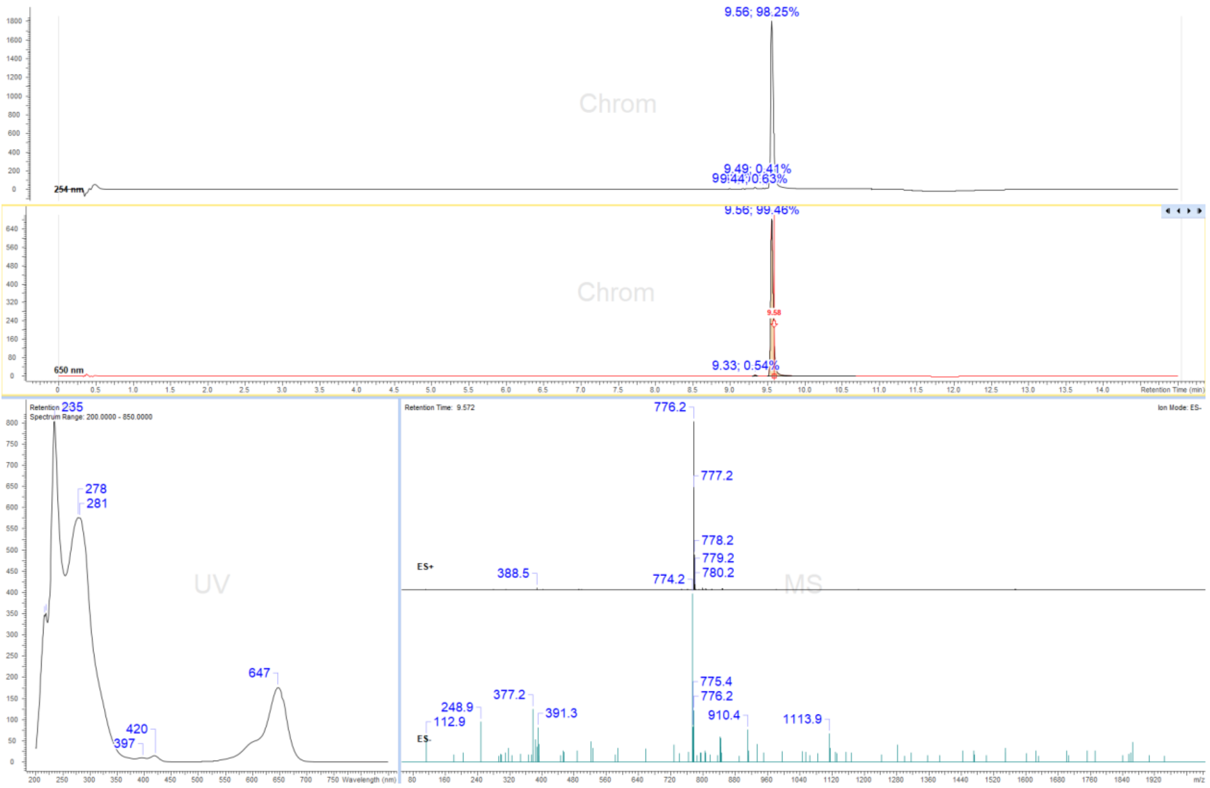

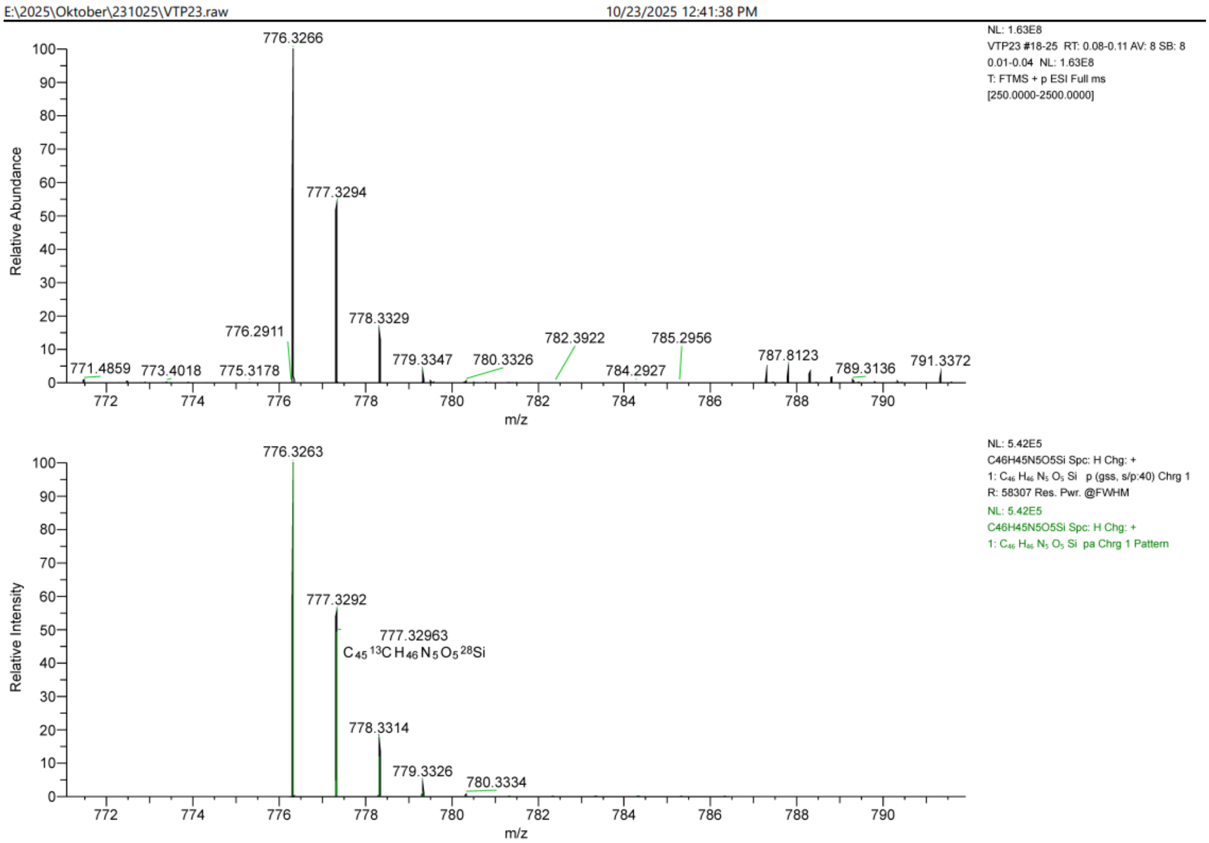
E:\2025\Oktober\231025\VTP23.raw
10/23/2025 12:41:38 PM
NL: 1.63E8
VTP23 #18-25 RT: 0.08-0.11 AV: 8 SB: 8 0.01-0.04 NL: 1.63E8
T: FTMS + p ESI Full ms [250.0000-2500.0000]
Relative Abundance
776.3266
777.3294
778.3329
776.2911
779.3347
780.3326
782.3922
785.2956
784.2927
787.8123
789.3136
791.3372
771.4859
773.4018
775.3178
m/z
NL: 5.42E5
C46H45N5O5Si Spc: H Chg: +
1: C46 H46 N5 O5 Si p (gss, s/p:40) Chrg 1
R: 58307 Res. Pwr. @FWHM
NL: 5.42E5
C46H45N5O5Si Spc: H Chg: +
1: C46 H46 N5 O5 Si pa Chrg 1 Pattern
Relative Intensity
776.3263
777.3292
777.32963
C45 13C H46 N5 O5 28Si
778.3314
779.3326
780.3334
m/z

Probe **3:**

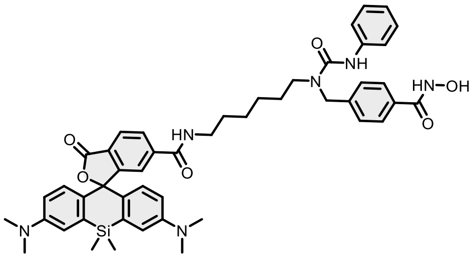

Reaction was conducted and purified following general method C. Fractions containing the product were collected, evaporated and lyophilized from acetonitrile: water mixture. The obtained solid was dissolved in 700 µL of $d_6$-DMSO. The samples from DMSO solution were diluted x100 in PBS +0.1%SDS and concentration was determined spectroscopically with Nanodrop. Determined stock concentration was 1.93 mM which constitutes 32% yield (1.13 mg).

$^1$H NMR (400 MHz, DMSO-$d_6$) δ 11.14 (s, 1H), 8.98 (s, 1H), 8.32 (s, 1H), 8.10 – 7.96 (m, 2H), 7.70 (d, $J$ = 8.2 Hz, 2H), 7.62 (s, 1H), 7.43 (d, $J$ = 7.8 Hz, 2H), 7.29 (d, $J$ = 8.2 Hz, 2H), 7.24 – 7.16 (m, 3H), 7.00 (d, $J$ = 2.4 Hz, 2H), 6.93 (d, $J$ = 7.3 Hz, 1H), 6.67 – 6.59 (m, 4H), 4.58 (s, 2H), 3.17 (t, $J$ = 5.5 Hz, 2H), 2.90 (s, 12H), 2.66 (p, $J$ = 1.9 Hz, 4H), 2.32 (p, $J$ = 1.9 Hz, 4H), 0.62 (s, 3H), 0.51 (s, 3H). HRMS (ESI) $m/z$ calculated 839.3952 for $C_{48}H_{55}N_6O_6Si$ $[M+H]^+$, found 839.3953.

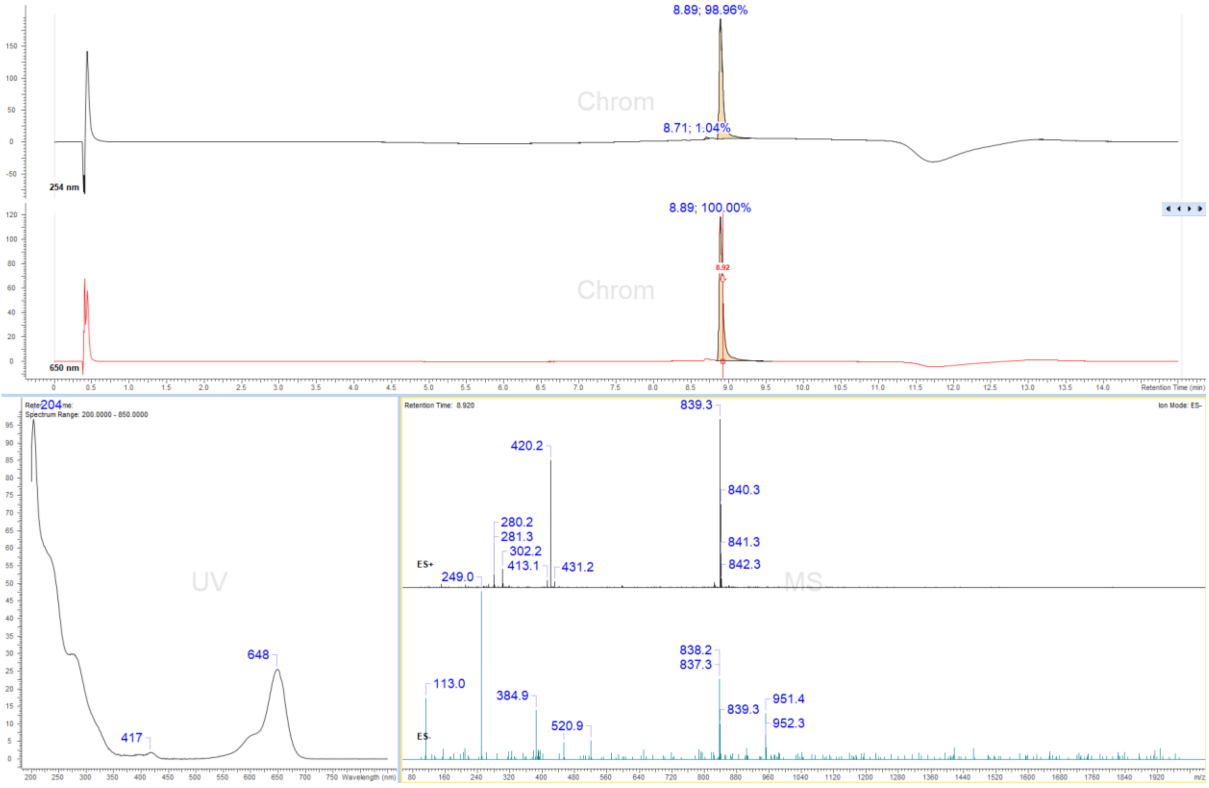

#### Display Report

**Analysis Info**

| | | | |
|---|---|---|---|
| Analysis Name | Z:\Data\2024\2409\sam030924\VTP16_16_01_127389.d | Acquisition Date | 03.09.2024 19:10:17 |
| Method | hystar_pl.m | Operator | BDAL@DE |
| Sample Name | VTP16 | Instrument / Ser# | micrOTOF 10237 |
| Comment | | | |

**Acquisition Parameter**

| | | | | | |
|---|---|---|---|---|---|
| Source Type | ESI | Ion Polarity | Positive | Set Nebulizer | 1.2 Bar |
| Focus | Not active | | | Set Dry Heater | 180 °C |
| Scan Begin | 50 m/z | Set Capillary | 4500 V | Set Dry Gas | 4.0 l/min |
| Scan End | 1600 m/z | Set End Plate Offset | -500 V | Set Divert Valve | Source |

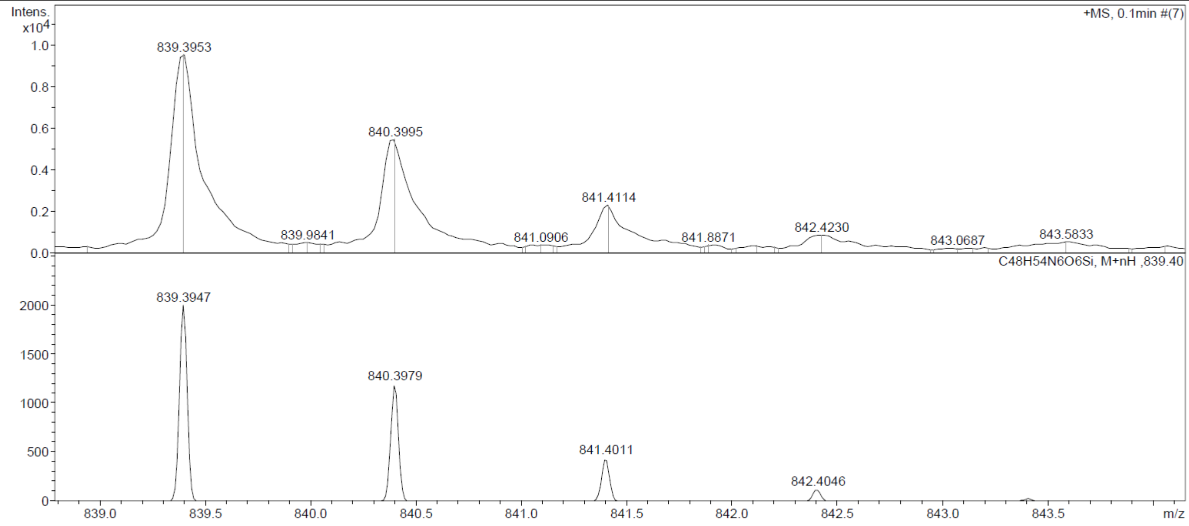

Probe **4**:

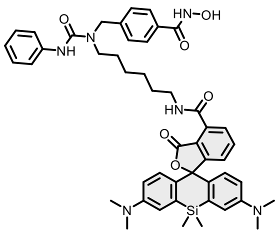

$^{1}$H NMR (400 MHz, DMSO-$d_6$) δ 11.15 (s, 1H), 9.02 – 8.95 (m, 2H), 8.35 (s, 1H), 7.78 – 7.67 (m, 4H), 7.50 – 7.42 (m, 2H), 7.31 (d, $J$ = 8.2 Hz, 2H), 7.28 – 7.16 (m, 3H), 6.99 (d, $J$ = 2.8 Hz, 2H), 6.95 – 6.89 (m, 1H), 6.69 (d, $J$ = 8.9 Hz, 2H), 6.63 (dd, $J$ = 9.0, 2.8 Hz, 2H), 4.61 (s, 2H), 3.29 (d, $J$ = 0.8 Hz, 4H), 2.91 (s, 12H), 1.54 (q, $J$ = 7.3 Hz, 4H), 1.41 – 1.27 (m, 4H), 0.61 (s, 3H), 0.51 (s, 3H). HRMS (ESI) $m/z$ calculated 839.3952 for $C_{48}H_{55}N_6O_6Si$ $[M+H]^+$, found 839.3946.

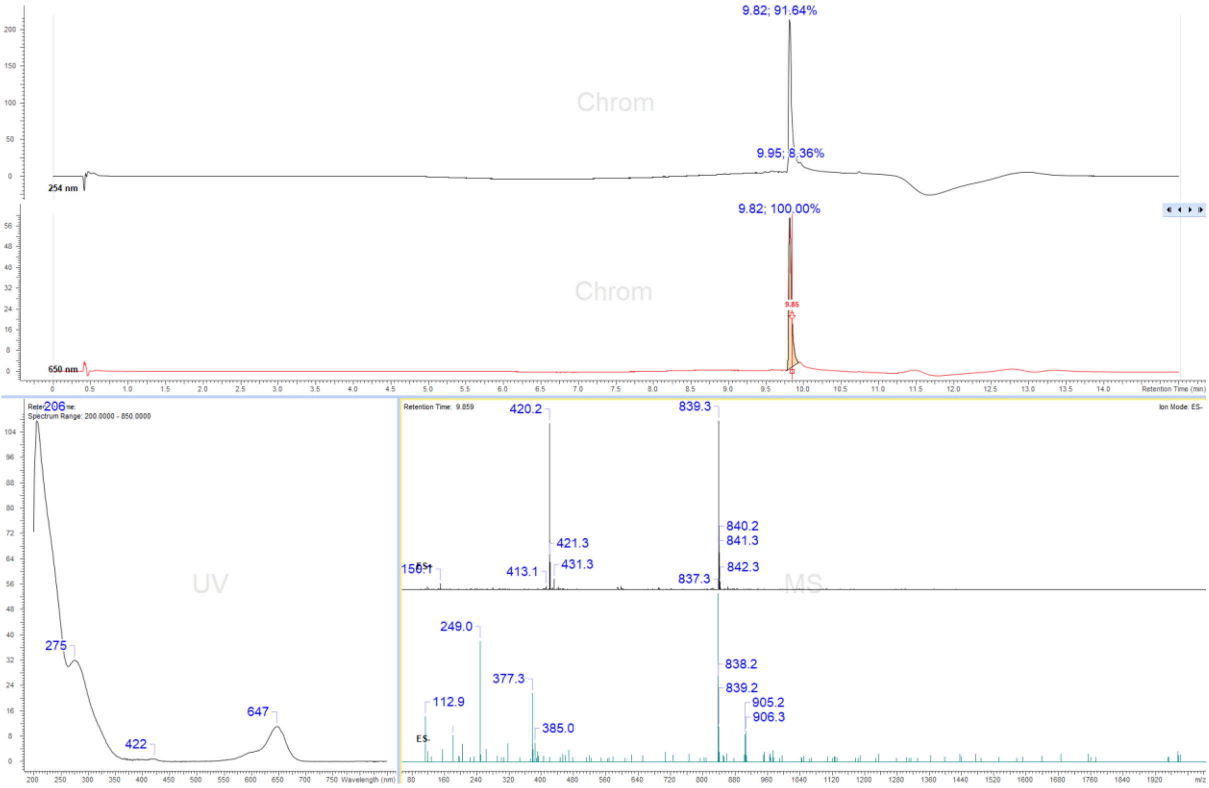

### Display Report

**Analysis Info**

| | | | |
|---|---|---|---|
| Analysis Name | Z:\Data\2024\2409\sam040924\VTP17_9_01_127413.d | Acquisition Date | 04.09.2024 18:30:16 |
| Method | hystar_pl.m | Operator | BDAL@DE |
| Sample Name | VTP17 | Instrument / Ser# | micrOTOF 10237 |
| Comment | | | |

**Acquisition Parameter**

| | | | | | |
|---|---|---|---|---|---|
| Source Type | ESI | Ion Polarity | Positive | Set Nebulizer | 1.2 Bar |
| Focus | Not active | | | Set Dry Heater | 180 °C |
| Scan Begin | 50 m/z | Set Capillary | 4500 V | Set Dry Gas | 4.0 l/min |
| Scan End | 1600 m/z | Set End Plate Offset | -500 V | Set Divert Valve | Source |

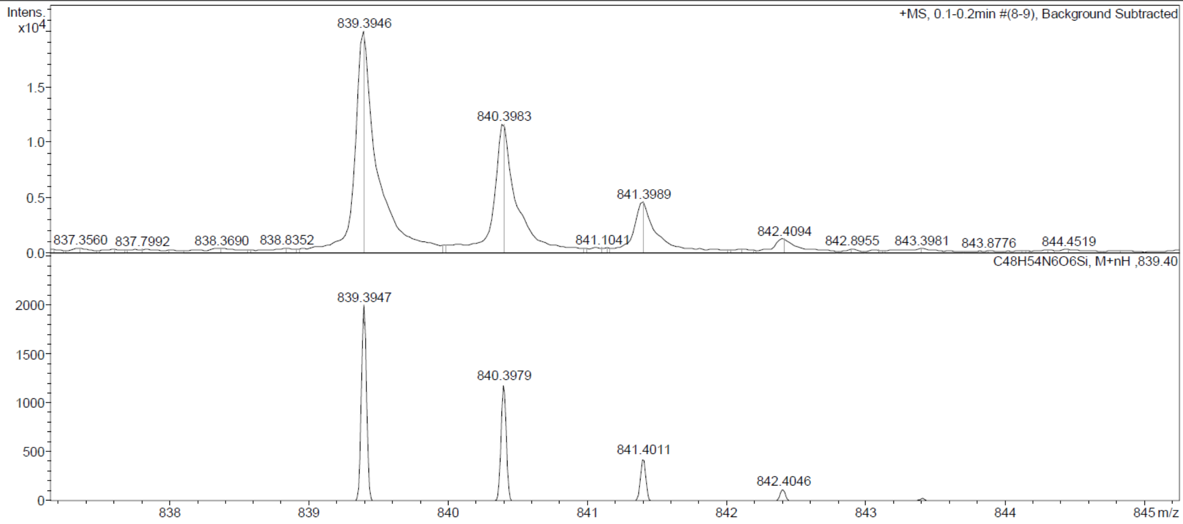

Probe **5:**

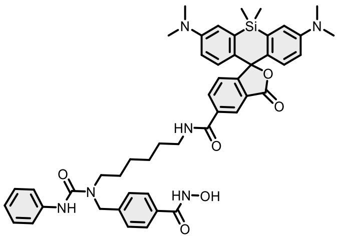

Reaction was conducted and purified following general method C. Fractions containing the product were collected, evaporated and lyophilized from acetonitrile: water mixture. The obtained solid was dissolved in 700 µL of *d*$_6$-DMSO. The samples from DMSO solution were diluted x100 in PBS +0.1%SDS and concentration was determined spectroscopically with Nanodrop. Determined stock concentration was 1.15 mM which constitutes to 19% yield (0.67 mg).

$^1$H NMR (400 MHz, DMSO-*d*$_6$) δ 11.15 (s, 1H), 8.99 (s, 1H), 8.73 (t, $J$ = 5.5 Hz, 1H), 8.42 – 8.32 (m, 2H), 8.20 (dd, $J$ = 8.1, 1.6 Hz, 1H), 7.76 – 7.68 (m, 2H), 7.50 – 7.42 (m, 2H), 7.37 – 7.27 (m, 3H), 7.25 – 7.16 (m, 2H), 7.01 (dd, $J$ = 2.3, 1.0 Hz, 2H), 6.92 (tt, $J$ = 7.4, 1.2 Hz, 1H), 6.61 (d, $J$ = 2.2 Hz, 4H), 4.61 (s, 2H), 3.29 (s, 4H), 2.91 (s, 12H), 1.52 (t, $J$ = 7.2 Hz, 4H), 1.30 (s, 4H), 0.62 (s, 3H), 0.51 (s, 3H). HRMS (ESI) *m*/*z* calculated 839.3952 for $C_{48}H_{55}N_6O_6Si$ $[M+H]^+$, found 839.3951.

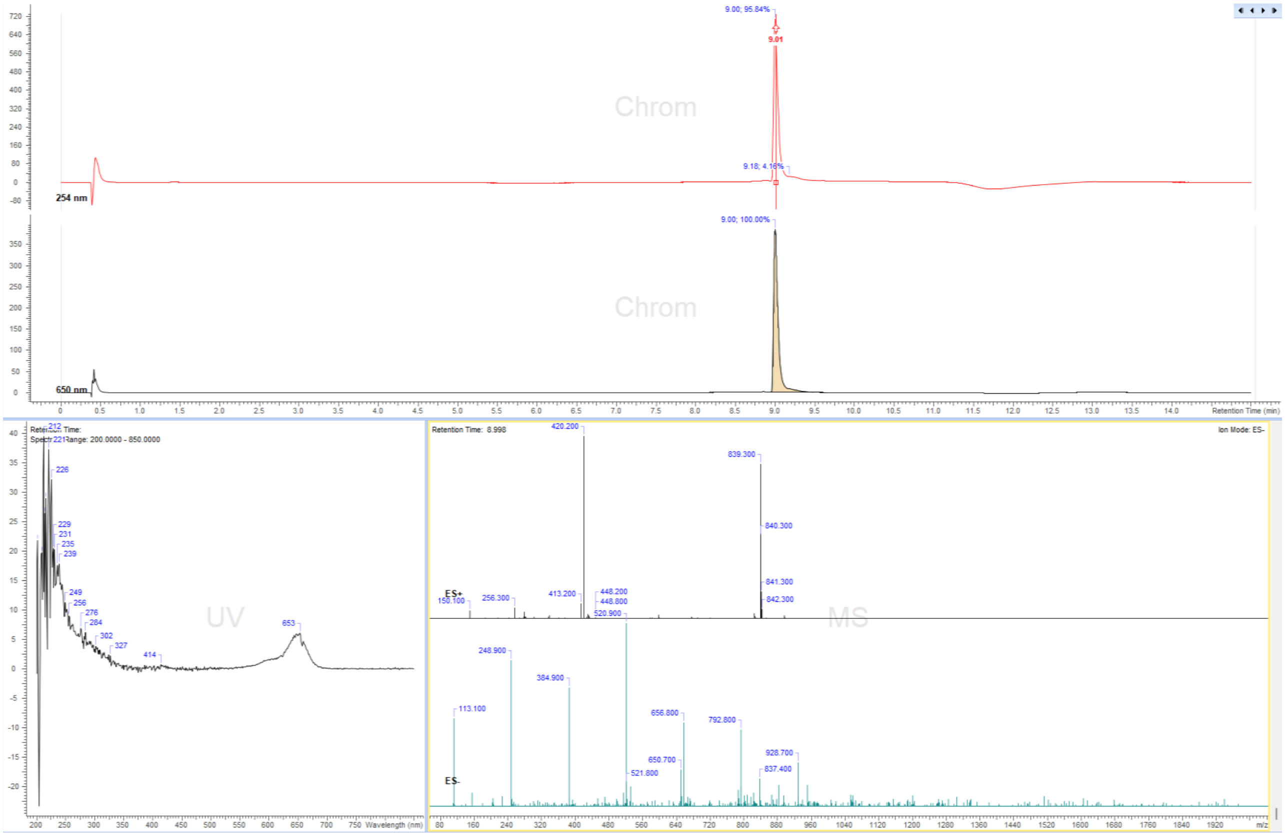

### Display Report

**Analysis Info**

| | | | |
|---|---|---|---|
| Analysis Name | Z:\Data\2024\2409\sam040924\VTP14_12_01_127416.d | Acquisition Date | 04.09.2024 18:44:15 |
| Method | hystar_pl.m | Operator | BDAL@DE |
| Sample Name | VTP14 | Instrument / Ser# | micrOTOF 10237 |
| Comment | | | |

**Acquisition Parameter**

| | | | | | |
|---|---|---|---|---|---|
| Source Type | ESI | Ion Polarity | Positive | Set Nebulizer | 1.2 Bar |
| Focus | Not active | | | Set Dry Heater | 180 °C |
| Scan Begin | 50 m/z | Set Capillary | 4500 V | Set Dry Gas | 4.0 l/min |
| Scan End | 1600 m/z | Set End Plate Offset | -500 V | Set Divert Valve | Source |

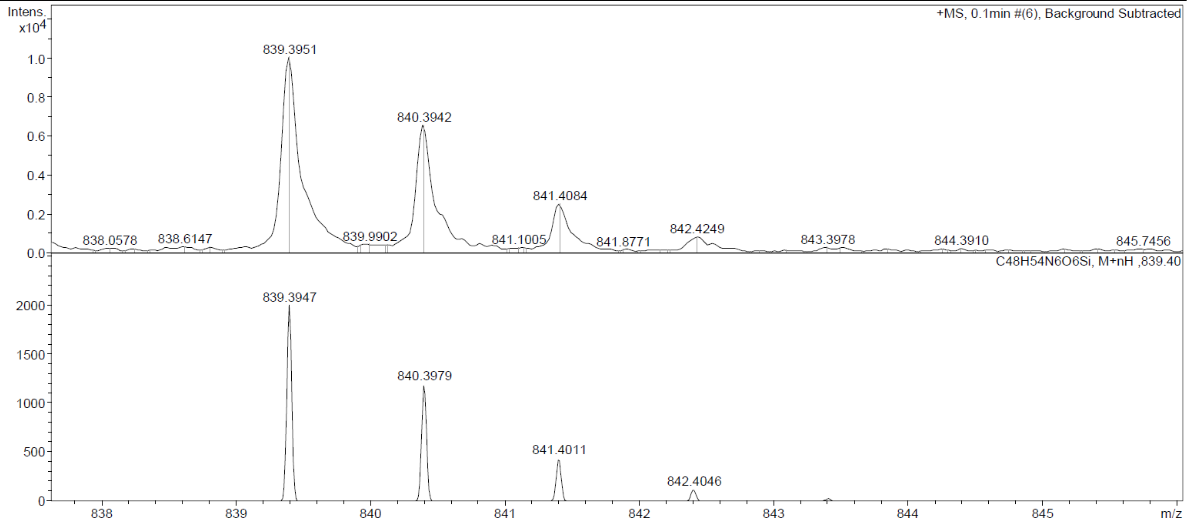

Probe **6:**

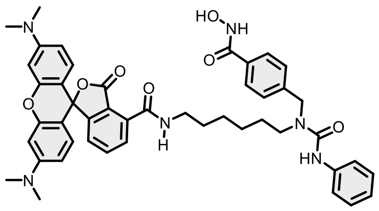

Reaction was conducted and purified following general method C. Fractions containing the product were collected, evaporated and lyophilized from acetonitrile: water mixture. The obtained solid was dissolved in 700 µL of $d_6$-DMSO. The samples from DMSO solution were diluted x100 in PBS +0.1%SDS and concentration was determined spectroscopically with Nanodrop. Determined stock concentration was 1.72 mM which constitutes to 26% yield (0.96 mg).

$^1$H NMR (400 MHz, DMSO-$d_6$) δ 11.15 (s, 1H), 9.01 (d, $J$ = 19.5 Hz, 2H), 8.35 (s, 1H), 7.88 – 7.65 (m, 5H), 7.47 – 7.44 (m, 2H), 7.31 (d, $J$ = 8.1 Hz, 2H), 7.22 (td, $J$ = 7.8, 5.5 Hz, 3H), 6.92 (t, $J$ = 7.2 Hz, 2H), 6.60 (d, $J$ = 8.7 Hz, 1H), 6.48 (s, 3H), 4.62 (s, 2H), 2.94 (s, 12H), 1.53 (s, 4H), 1.34 (d, $J$ = 37.9 Hz, 4H). HRMS (ESI) $m/z$ calculated 797.3662 for $C_{46}H_{49}N_6O_7$ $[M+H]^+$, found 797.3652.

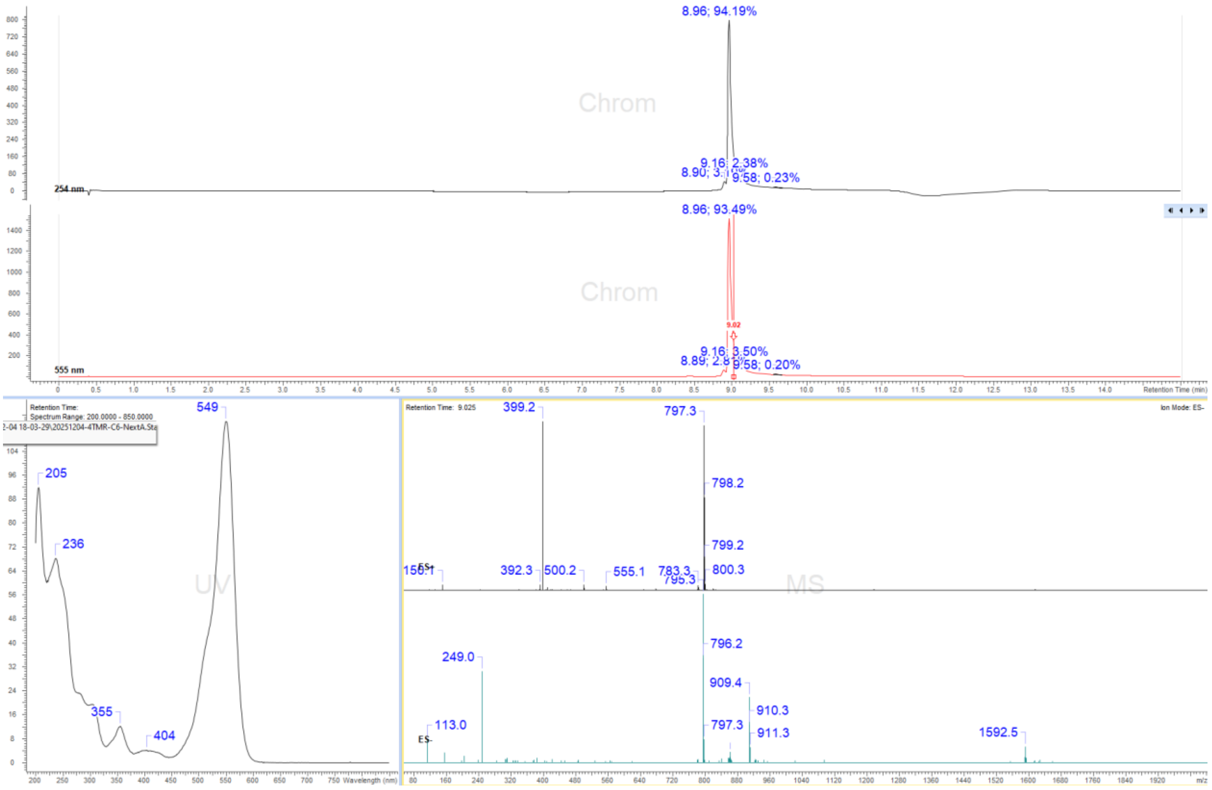

### Display Report

**Analysis Info**

| | | | |
|---|---|---|---|
| Analysis Name | Z:\Data\2024\2409\sam040924\VTP19_7_01_127411.d | Acquisition Date | 04.09.2024 18:20:58 |
| Method | hystar_pl.m | Operator | BDAL@DE |
| Sample Name | VTP19 | Instrument / Ser# | micrOTOF 10237 |
| Comment | | | |

**Acquisition Parameter**

| | | | | | |
|---|---|---|---|---|---|
| Source Type | ESI | Ion Polarity | Positive | Set Nebulizer | 1.2 Bar |
| Focus | Not active | | | Set Dry Heater | 180 °C |
| Scan Begin | 50 m/z | Set Capillary | 4500 V | Set Dry Gas | 4.0 l/min |
| Scan End | 1600 m/z | Set End Plate Offset | -500 V | Set Divert Valve | Source |

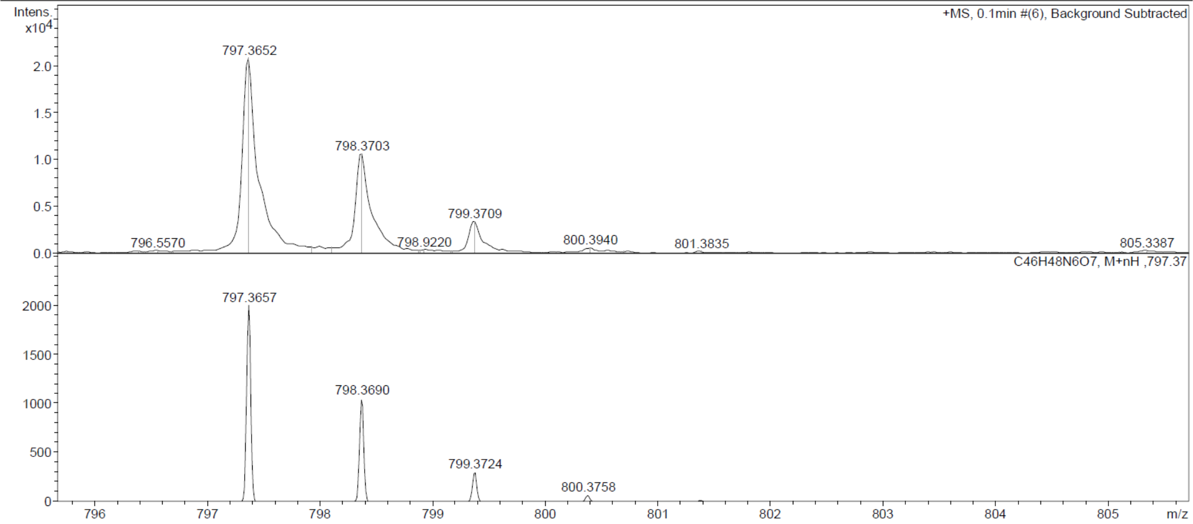

Probe **7:**

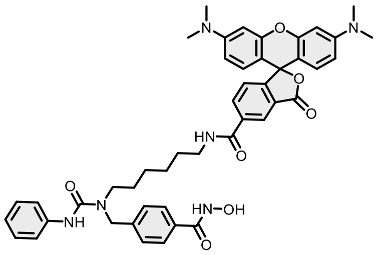

Reaction was conducted and purified following general method C. Fractions containing the product were collected, evaporated and lyophilized from acetonitrile: water mixture. The obtained solid was dissolved in 700 µL of *d*$_6$-DMSO. The samples from DMSO solution were diluted x100 in PBS +0.1%SDS and concentration was determined spectroscopically with Nanodrop. Determined stock concentration was 2.72 mM which constitutes to 41% yield (1.52 mg).

$^{1}$H NMR (400 MHz, DMSO-*d*$_6$) δ 11.15 (s, 1H), 8.98 (s, 1H), 8.77 (t, *J* = 5.5 Hz, 1H), 8.43 (dd, *J* = 1.7, 0.8 Hz, 1H), 8.36 (s, 1H), 8.21 (dd, *J* = 8.0, 1.6 Hz, 1H), 7.75 – 7.69 (m, 2H), 7.50 – 7.42 (m, 2H), 7.35 – 7.27 (m, 3H), 7.25 – 7.17 (m, 2H), 6.96 – 6.89 (m, 1H), 6.54 – 6.43 (m, 6H), 4.62 (s, 2H), 3.28 (d, *J* = 5.8 Hz, 4H), 2.93 (s, 12H), 1.57 – 1.48 (m, 4H), 1.31 (s, 4H). HRMS (ESI) *m*/*z* calculated 797.3662 for $C_{46}H_{49}N_6O_7$ [M+H]$^+$, found 797.3653.

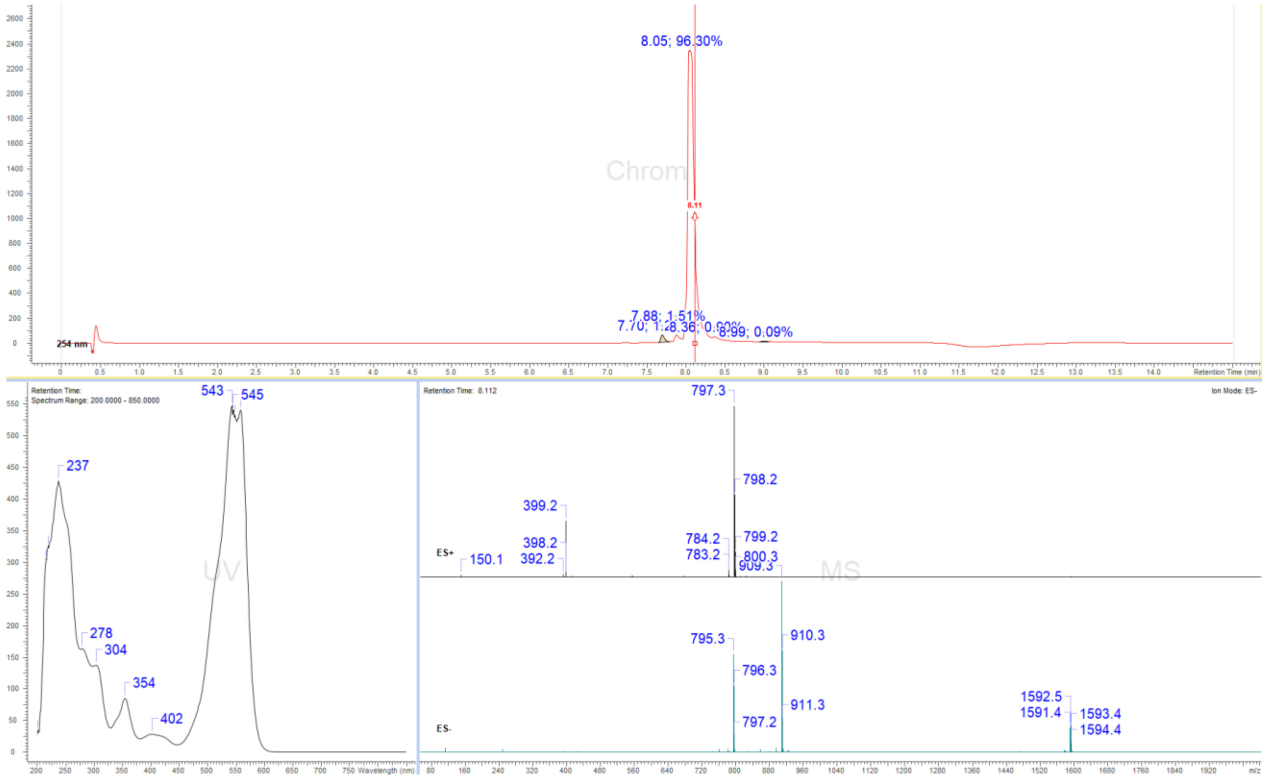

### Display Report

**Analysis Info**

| | | | |
|---|---|---|---|
| Analysis Name | Z:\Data\2024\2409\sam040924\VTP3_2_01_127406.d | Acquisition Date | 04.09.2024 17:57:46 |
| Method | hystar_pl.m | Operator | BDAL@DE |
| Sample Name | VTP3 | Instrument / Ser# | micrOTOF 10237 |
| Comment | | | |

**Acquisition Parameter**

| | | | | | |
|---|---|---|---|---|---|
| Source Type | ESI | Ion Polarity | Positive | Set Nebulizer | 1.2 Bar |
| Focus | Not active | | | Set Dry Heater | 180 °C |
| Scan Begin | 50 m/z | Set Capillary | 4500 V | Set Dry Gas | 4.0 l/min |
| Scan End | 1600 m/z | Set End Plate Offset | -500 V | Set Divert Valve | Source |

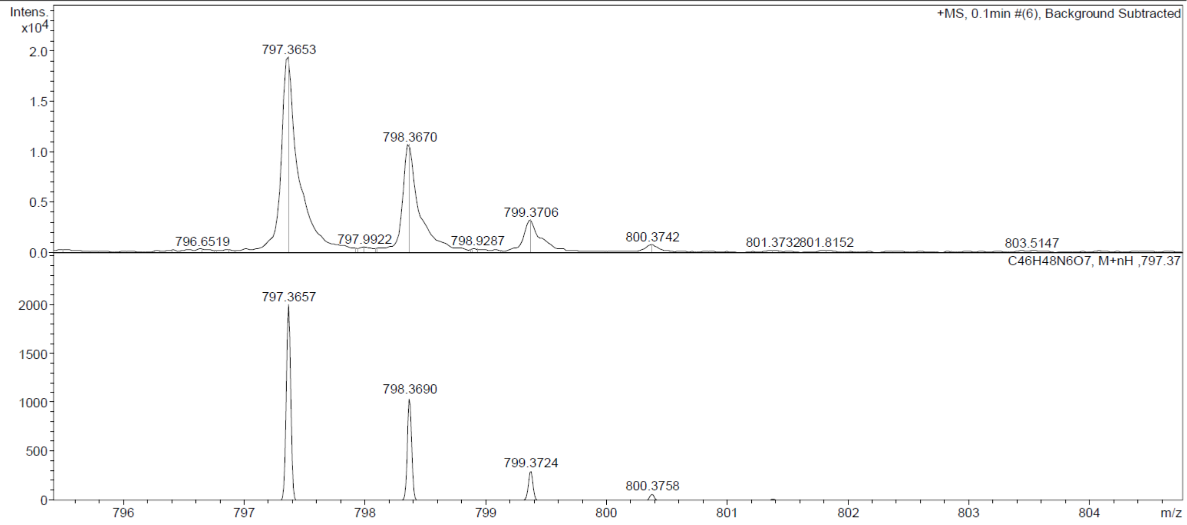

Probe **8:**

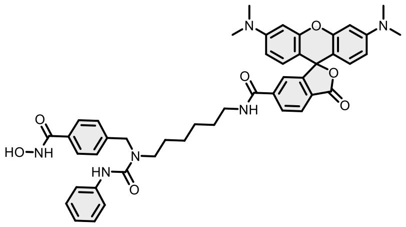

Reaction was conducted and purified following general method C. Fractions containing the product were collected, evaporated and lyophilized from acetonitrile: water mixture. The obtained solid was dissolved in 700 µL of d6-DMSO. The samples from DMSO solution were diluted x100 in PBS +0.1%SDS and concentration was determined spectroscopically with Nanodrop. Determined stock concentration was 1.66 mM which constitutes 25% yield (0.92 mg).

$^1$H NMR (400 MHz, DMSO-$d_6$) δ 11.14 (s, 1H), 8.99 (s, 1H), 8.64 (t, $J$ = 5.6 Hz, 1H), 8.32 (s, 1H), 8.14 (dd, $J$ = 8.0, 1.4 Hz, 1H), 8.04 (dd, $J$ = 8.0, 0.7 Hz, 1H), 7.74 – 7.65 (m, 2H), 7.61 (t, $J$ = 1.1 Hz, 1H), 7.46 – 7.39 (m, 2H), 7.33 – 7.26 (m, 2H), 7.24 – 7.15 (m, 2H), 6.98 – 6.87 (m, 1H), 6.53 – 6.46 (m, 6H), 4.57 (s, 2H), 3.24 (d, $J$ = 7.7 Hz, 2H), 3.16 (q, $J$ = 6.7 Hz, 2H), 2.93 (s, 12H), 1.50 – 1.40 (m, 4H), 1.23 (s, 4H). HRMS (ESI) *m*/*z* calculated 797.3662 for $C_{46}H_{49}N_6O_7$ $[M+H]^+$, found 797.3660.

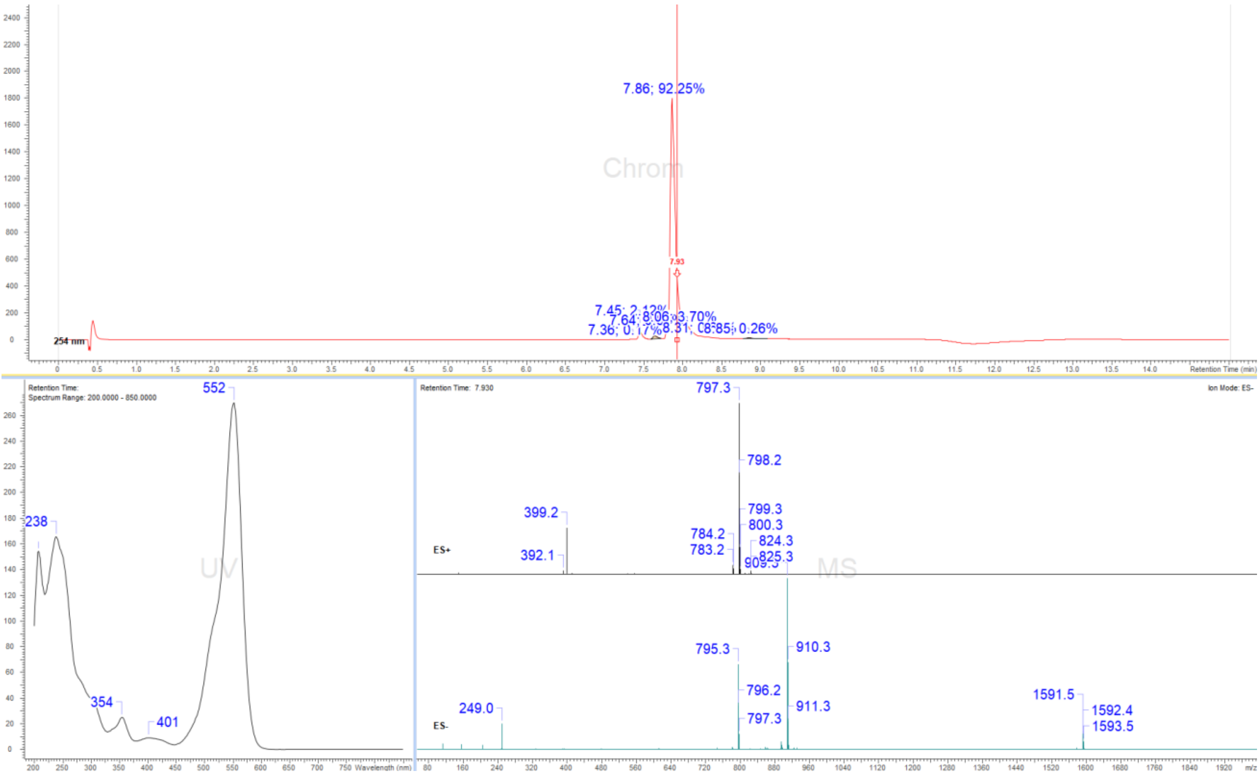

#### Display Report

**Analysis Info**

| | | | |
|---|---|---|---|
| Analysis Name | Z:\Data\2024\2409\sam040924\VTP2_11_01_127415.d | Acquisition Date | 04.09.2024 18:39:37 |
| Method | hystar_pl.m | Operator | BDAL@DE |
| Sample Name | VTP2 | Instrument / Ser# | micrOTOF 10237 |
| Comment | | | |

**Acquisition Parameter**

| | | | | | |
|---|---|---|---|---|---|
| Source Type | ESI | Ion Polarity | Positive | Set Nebulizer | 1.2 Bar |
| Focus | Not active | | | Set Dry Heater | 180 °C |
| Scan Begin | 50 m/z | Set Capillary | 4500 V | Set Dry Gas | 4.0 l/min |
| Scan End | 1600 m/z | Set End Plate Offset | -500 V | Set Divert Valve | Source |

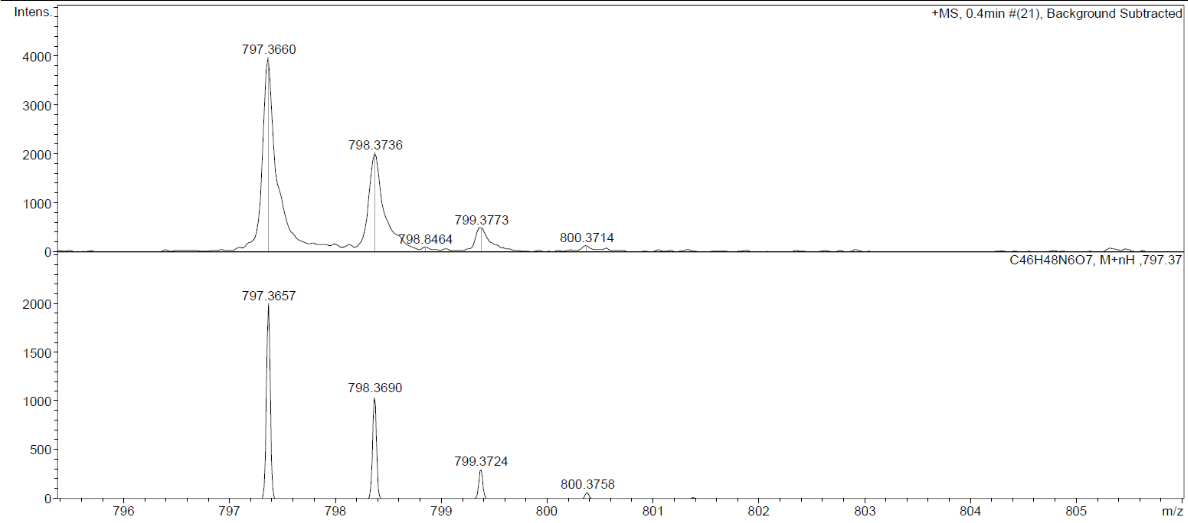

Probe **9**:

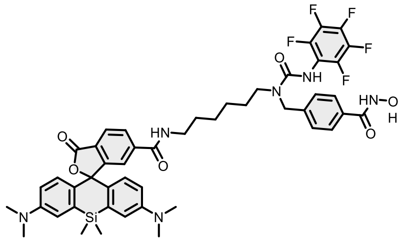

Reaction was conducted and purified following general method C. Fractions containing the product were collected, evaporated and lyophilized from acetonitrile: water mixture. The obtained solid was dissolved in 700 µL of d6-DMSO. The samples from DMSO solution were diluted x100 in PBS +0.1%SDS and concentration was determined spectroscopically with Nanodrop. Determined stock concentration was 1.63 mM which constitutes 27% yield (1.06 mg).

$^{1}$H NMR (400 MHz, DMSO-$d_6$) δ 8.69 (t, $J$ = 5.6 Hz, 1H), 8.57 (s, 1H), 8.29 (s, 1H), 8.07 – 7.98 (m, 2H), 7.74 – 7.70 (m, 2H), 7.63 (t, $J$ = 1.1 Hz, 1H), 7.32 – 7.27 (m, 2H), 7.00 (dd, $J$ = 2.5, 0.9 Hz, 2H), 6.66 – 6.59 (m, 4H), 4.56 (s, 2H), 3.17 (d, $J$ = 10.1 Hz, 4H), 2.91 (s, 12H), 1.45 (d, $J$ = 9.5 Hz, 4H), 1.23 (s, 4H), 0.62 (s, 3H), 0.51 (s, 3H). HRMS (ESI) $m/z$ calculated 929.3481 for $C_{48}H_{50}F_5N_6O_6Si$ $[M+H]^+$, found 929.3482.

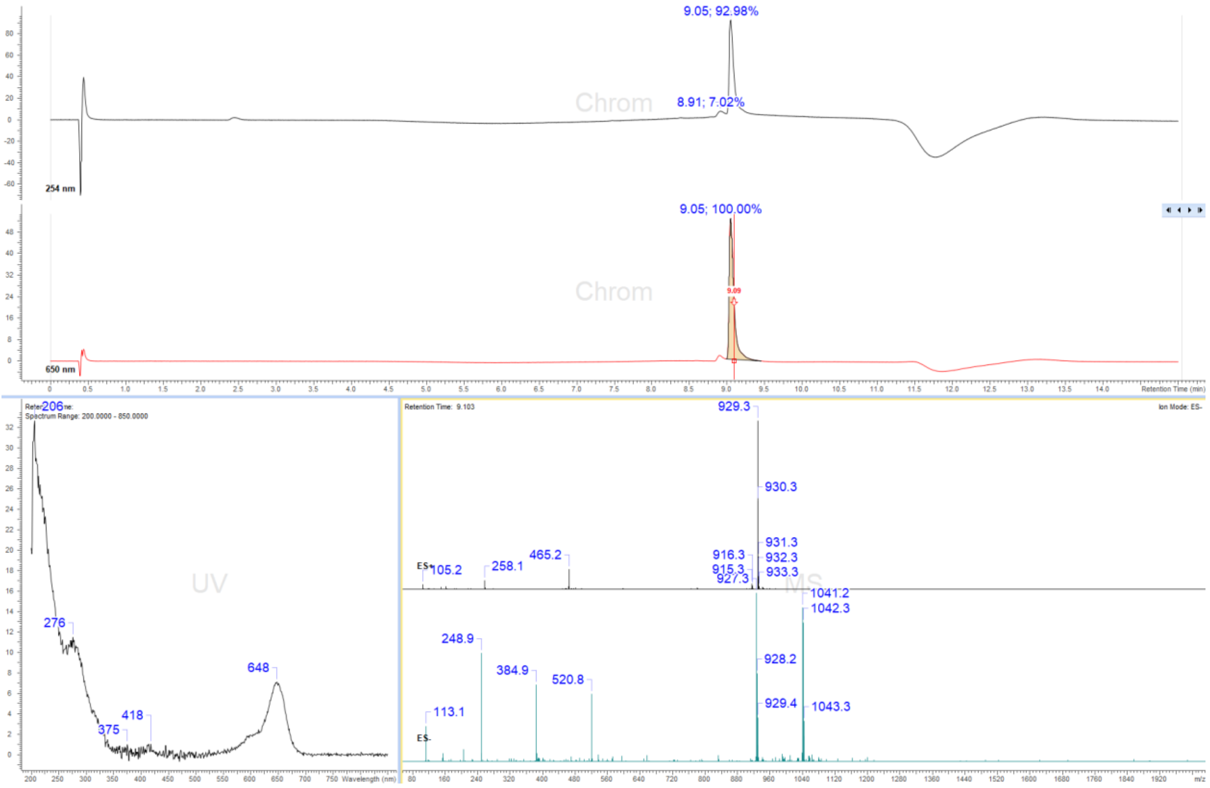

### Display Report

**Analysis Info**

| | | | |
|---|---|---|---|
| Analysis Name | Z:\Data\2024\2409\sam300924\VTP22_14_01_127691.d | Acquisition Date | 30.09.2024 13:55:47 |
| Method | hystar_pl.m | Operator | BDAL@DE |
| Sample Name | VTP22 | Instrument / Ser# | micrOTOF 10237 |
| Comment | | | |

**Acquisition Parameter**

| | | | | | |
|---|---|---|---|---|---|
| Source Type | ESI | Ion Polarity | Positive | Set Nebulizer | 1.2 Bar |
| Focus | Not active | | | Set Dry Heater | 180 °C |
| Scan Begin | 50 m/z | Set Capillary | 4500 V | Set Dry Gas | 4.0 l/min |
| Scan End | 1600 m/z | Set End Plate Offset | -500 V | Set Divert Valve | Source |

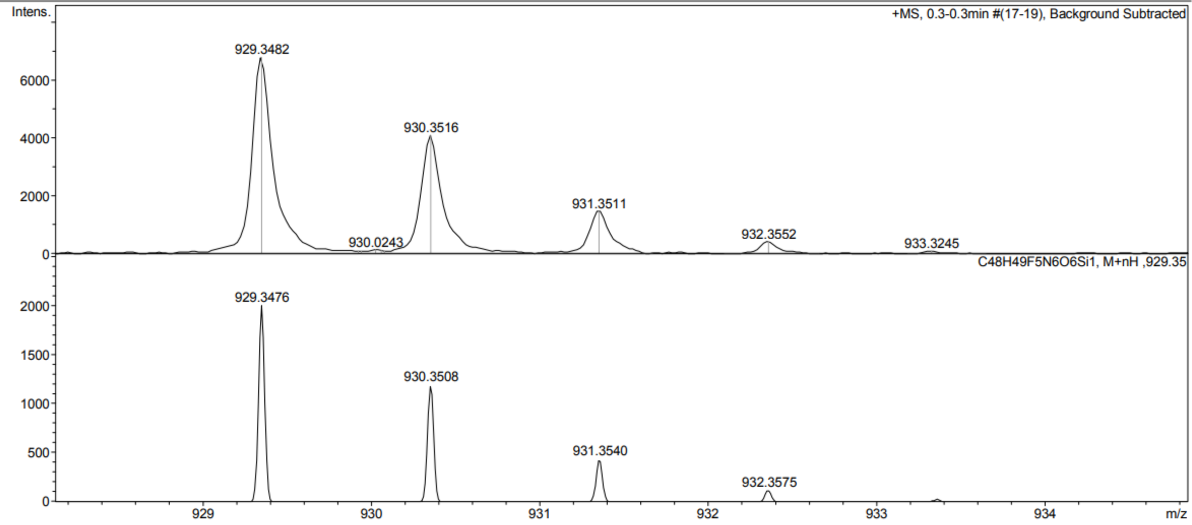

Probe **10:**

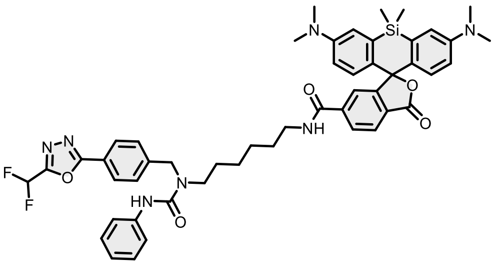

Reaction was conducted and purified following general method C. Fractions containing the product were collected, evaporated and lyophilized from acetonitrile: water mixture. The obtained solid was dissolved in 700 µL of $d_6$-DMSO. The samples from DMSO solution were diluted x100 in PBS +0.1%SDS and concentration was determined spectroscopically with Nanodrop. Determined stock concentration was 3.70 mM which constitutes 62% yield (2.35 mg).

$^1$H NMR (400 MHz, DMSO-$d_6$) δ 8.68 (s, 1H), 8.37 (s, 1H), 8.06 – 7.97 (m, 4H), 7.63 (d, $J$ = 1.2 Hz, 1H), 7.51 – 7.41 (m, 4H), 7.24 – 7.18 (m, 2H), 7.00 (dd, $J$ = 2.6, 0.8 Hz, 2H), 6.96 – 6.89 (m, 1H), 6.65 – 6.59 (m, 4H), 4.65 (s, 2H), 2.90 (s, 12H), 2.66 (p, $J$ = 1.8 Hz, 2H), 2.32 (p, $J$ = 1.9 Hz, 2H), 1.45 (s, 4H), 1.23 (s, 4H), 0.62 (s, 3H), 0.51 (s, 3H). HRMS (ESI) $m/z$ calculated 898.3923 for $C_{50}H_{54}F_2N_7O_5Si$ [M+H]$^+$, found 898.3911.

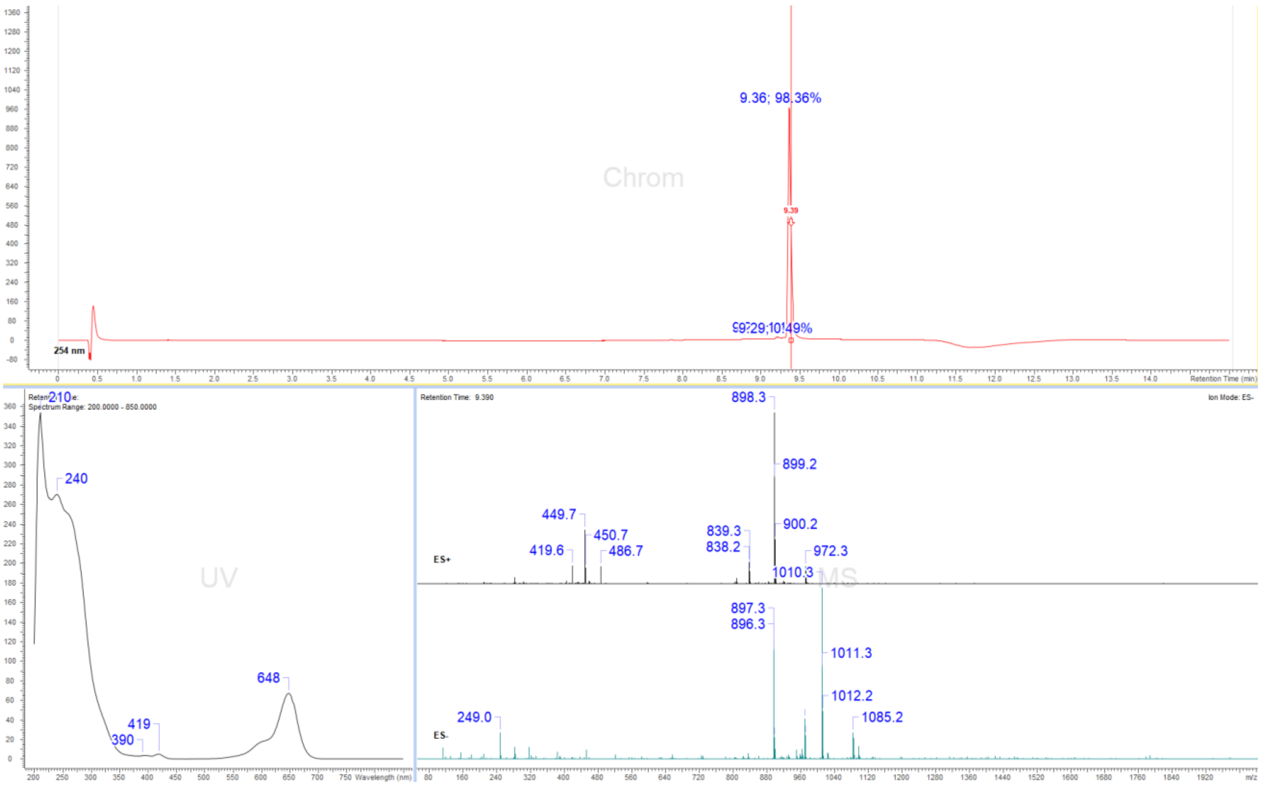

### Display Report

**Analysis Info**

| | | | |
|---|---|---|---|
| Analysis Name | Z:\Data\2024\2409\sam040924\VTP18_8_01_127412.d | Acquisition Date | 04.09.2024 18:25:36 |
| Method | hystar_pl.m | Operator | BDAL@DE |
| Sample Name | VTP18 | Instrument / Ser# | micrOTOF 10237 |
| Comment | | | |

**Acquisition Parameter**

| | | | | | |
|---|---|---|---|---|---|
| Source Type | ESI | Ion Polarity | Positive | Set Nebulizer | 1.2 Bar |
| Focus | Not active | | | Set Dry Heater | 180 °C |
| Scan Begin | 50 m/z | Set Capillary | 4500 V | Set Dry Gas | 4.0 l/min |
| Scan End | 1600 m/z | Set End Plate Offset | -500 V | Set Divert Valve | Source |

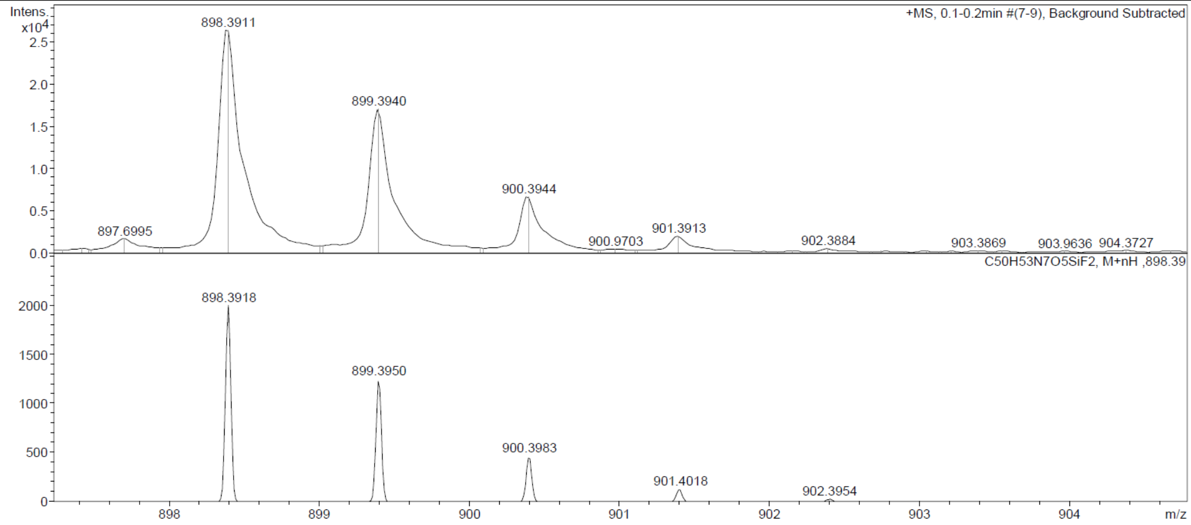

Probe **11:**

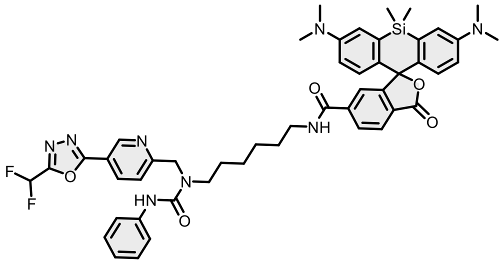

Reaction was conducted and purified following general method C. Fractions containing the product were collected, evaporated and lyophilized from acetonitrile: water mixture. The obtained solid was dissolved in 700 µL of $d_6$-DMSO. The samples from DMSO solution were diluted x100 in PBS +0.1%SDS and concentration was determined spectroscopically with Nanodrop. Determined stock concentration was 3.41 mM which constitutes to 52% yield (1.98 mg).

$^{1}$H NMR (400 MHz, DMSO-$d_6$) δ 8.98 (dd, $J$ = 2.3, 0.8 Hz, 1H), 8.68 (t, $J$ = 5.6 Hz, 1H), 8.57 (s, 1H), 8.20 (dd, $J$ = 8.2, 2.3 Hz, 1H), 8.08 – 7.97 (m, 2H), 7.63 (t, $J$ = 1.1 Hz, 1H), 7.43 (dd, $J$ = 8.6, 1.2 Hz, 3H), 7.20 (dd, $J$ = 8.5, 7.3 Hz, 2H), 7.00 (dd, $J$ = 2.6, 0.8 Hz, 2H), 6.95 – 6.89 (m, 1H), 6.68 – 6.59 (m, 4H), 6.50 (d, $J$ = 53.1 Hz, 1H), 4.68 (s, 2H), 3.29 (s, 2H), 3.18 (q, $J$ = 7.6 Hz, 2H), 2.90 (s, 12H), 1.46 (dd, $J$ = 16.3, 9.6 Hz, 4H), 1.23 (d, $J$ = 5.7 Hz, 4H), 0.62 (s, 3H), 0.51 (s, 3H). HRMS (ESI) $m/z$ calculated 899.3876 for $C_{49}H_{53}F_2N_8O_5Si$ $[M+H]^+$, found 899.3865.

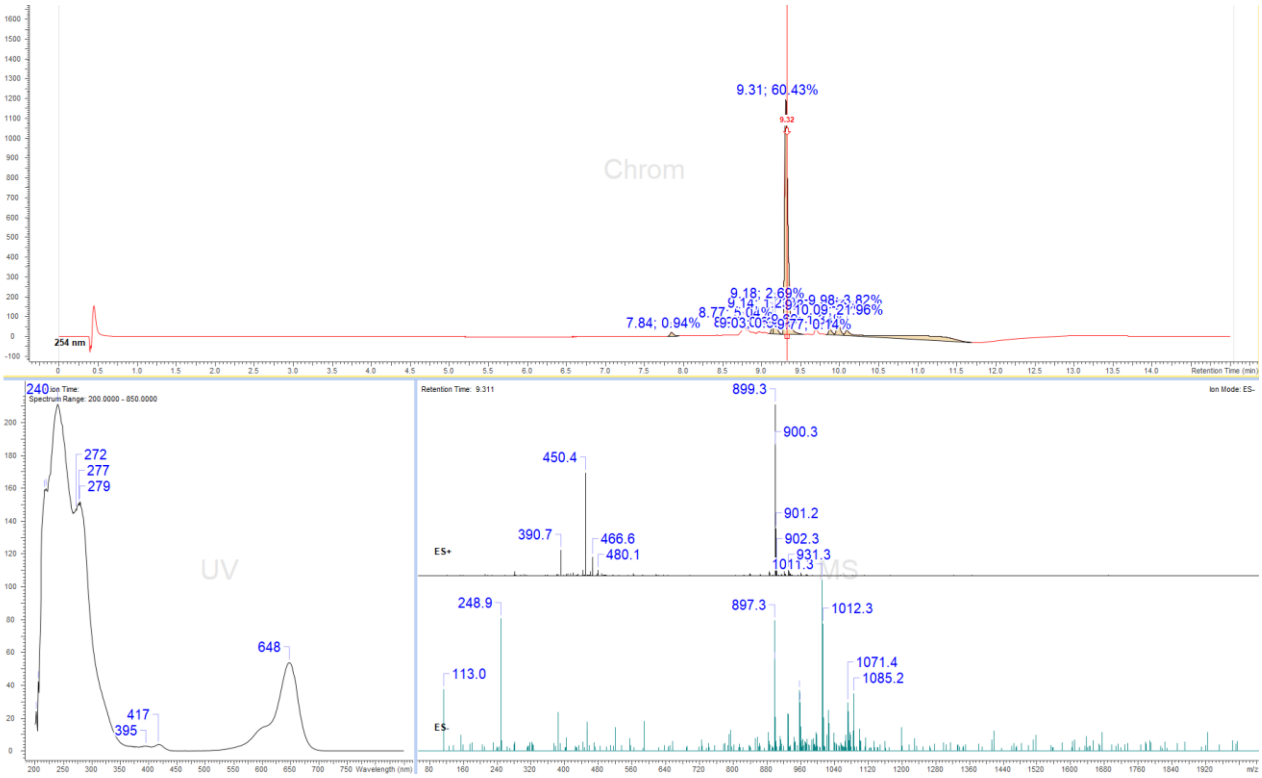

#### Display Report

| Analysis Info | | Acquisition Date | 04.09.2024 18:16:20 | |
|---|---|---|---|---|
| Analysis Name | Z:\Data\2024\2409\sam040924\VTP21_6_01_127410.d | | | |
| Method | hystar_pl.m | Operator | BDAL@DE | |
| Sample Name | VTP21 | Instrument / Ser# | micrOTOF | 10237 |
| Comment | | | | |

**Acquisition Parameter**

| | | | | | |
|---|---|---|---|---|---|
| Source Type | ESI | Ion Polarity | Positive | Set Nebulizer | 1.2 Bar |
| Focus | Not active | | | Set Dry Heater | 180 °C |
| Scan Begin | 50 m/z | Set Capillary | 4500 V | Set Dry Gas | 4.0 l/min |
| Scan End | 1600 m/z | Set End Plate Offset | -500 V | Set Divert Valve | Source |

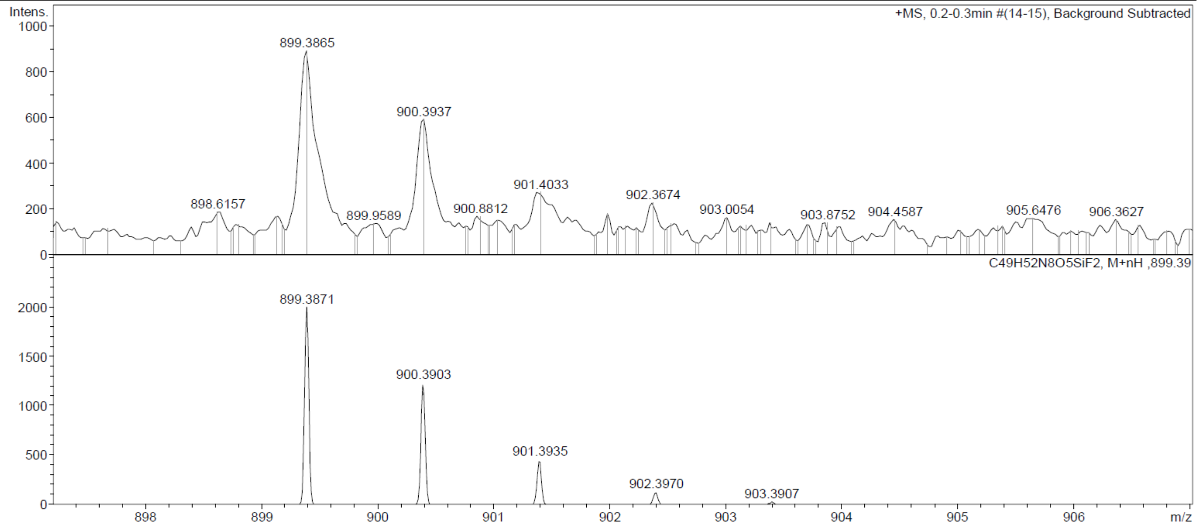

Probe **12:**

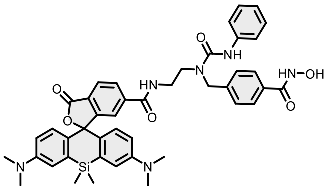

Reaction was conducted and purified following general method C. Fractions containing the product were collected, evaporated and lyophilized from acetonitrile: water mixture. The obtained solid was dissolved in 700 µL of d6-DMSO. The samples from DMSO solution were diluted x100 in PBS +0.1%SDS and concentration was determined spectroscopically with Nanodrop. Determined stock concentration was 3.5 mM which constitutes 49% yield (1.6 mg).

$^{1}$H NMR (400 MHz, DMSO-$d_6$) δ 11.15 (s, 1H), 9.00 (s, 1H), 8.58 (s, 1H), 8.10 – 7.97 (m, 2H), 7.79 – 7.62 (m, 3H), 7.52 – 7.43 (m, 2H), 7.28 (d, $J$ = 8.1 Hz, 2H), 7.19 (dd, $J$ = 8.6, 7.3 Hz, 2H), 7.01 (t, $J$ = 1.7 Hz, 2H), 6.97 – 6.88 (m, 1H), 6.61 (d, $J$ = 1.6 Hz, 4H), 4.62 (s, 2H), 3.29 (s, 4H), 2.90 (s, 12H), 0.65 (s, 3H), 0.52 (s, 3H). HRMS (ESI) $m/z$ calculated 783.3326 for $C_{44}H_{47}N_6O_6Si$, found 783.3324 ($[M+H]^+$).

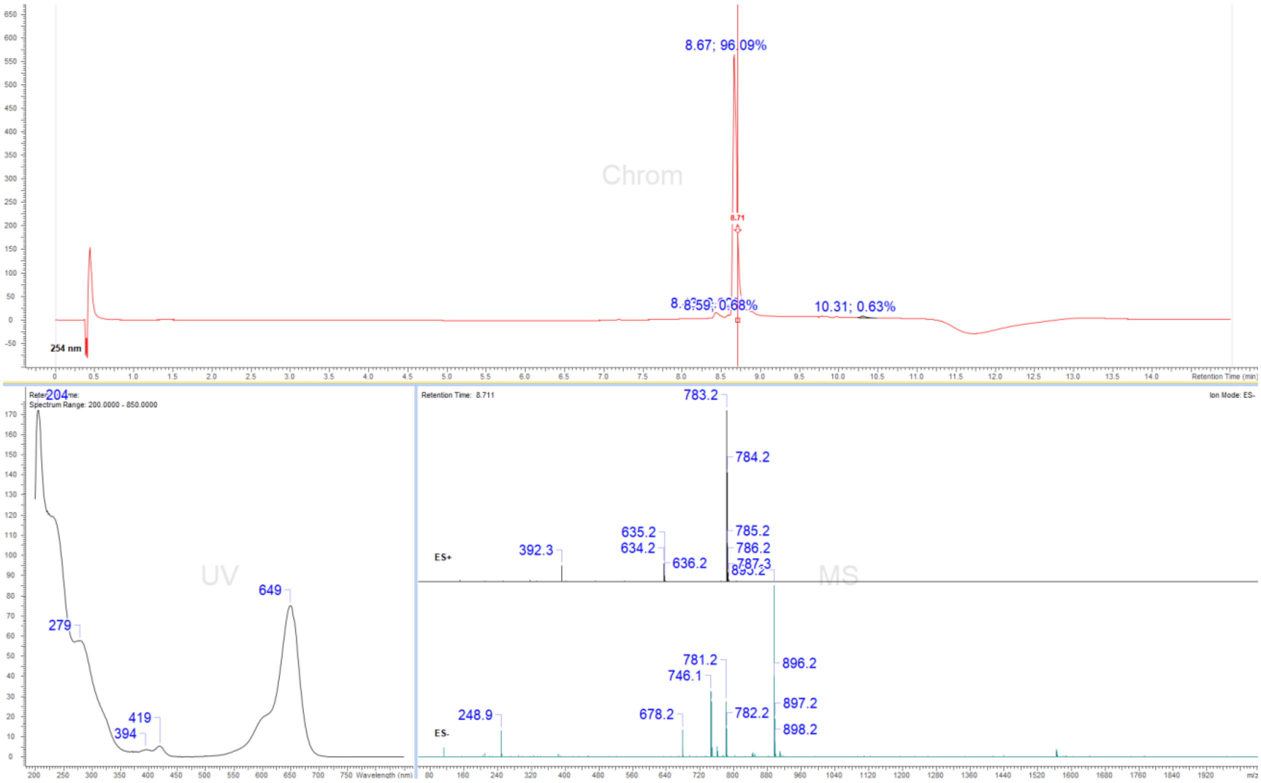

### Display Report

**Analysis Info**

| | | | |
|---|---|---|---|
| Analysis Name | Z:\Data\2024\2409\sam030924\VTP15_12_01_127385.d | Acquisition Date | 03.09.2024 18:51:44 |
| Method | hystar_pl.m | Operator | BDAL@DE |
| Sample Name | VTP15 | Instrument / Ser# | micrOTOF 10237 |
| Comment | | | |

**Acquisition Parameter**

| | | | | | |
|---|---|---|---|---|---|
| Source Type | ESI | Ion Polarity | Positive | Set Nebulizer | 1.2 Bar |
| Focus | Not active | | | Set Dry Heater | 180 °C |
| Scan Begin | 50 m/z | Set Capillary | 4500 V | Set Dry Gas | 4.0 l/min |
| Scan End | 1600 m/z | Set End Plate Offset | -500 V | Set Divert Valve | Source |

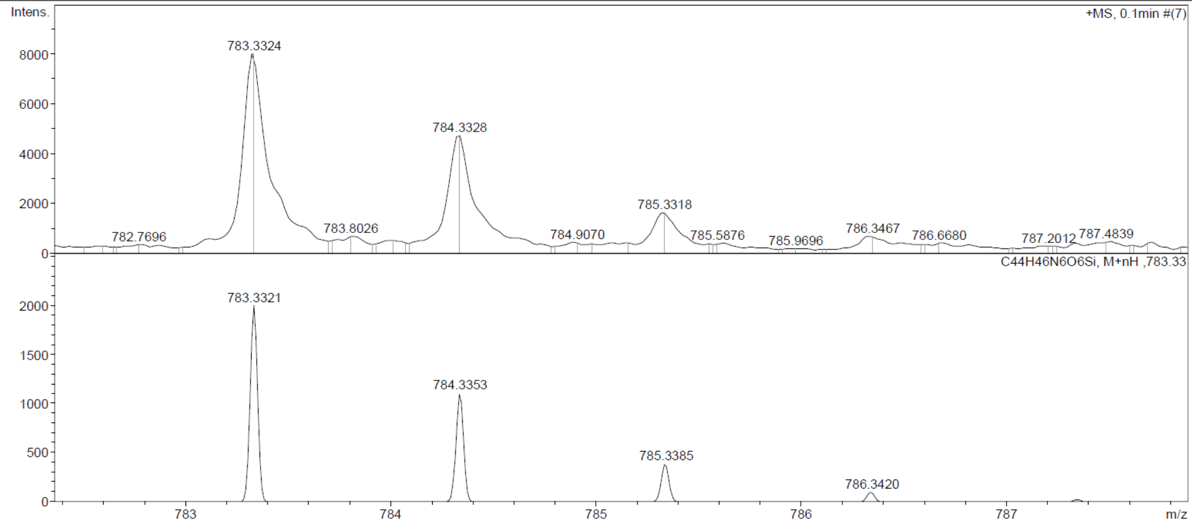

Probe **13:**

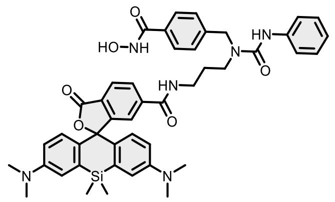

Reaction was conducted and purified following general method C. Fractions containing the product were collected, evaporated and lyophilized from acetonitrile: water mixture. The obtained solid was dissolved in 700 µL of d6-DMSO. The samples from DMSO solution were diluted x100 in PBS +0.1%SDS and concentration was determined spectroscopically with Nanodrop. Determined stock concentration was 1.57 mM which constitutes to 24% yield (0.82 mg).

$^{1}$H NMR (400 MHz, DMSO-$d_6$) δ 11.14 (s, 1H), 8.97 (d, $J$ = 1.8 Hz, 1H), 8.71 (t, $J$ = 5.6 Hz, 1H), 8.38 (s, 1H), 8.10 – 7.96 (m, 2H), 7.71 – 7.64 (m, 3H), 7.44 – 7.40 (m, 2H), 7.27 (d, $J$ = 8.2 Hz, 2H), 7.21 – 7.16 (m, 2H), 7.01 (dd, $J$ = 2.2, 1.1 Hz, 2H), 6.95 – 6.89 (m, 1H), 6.67 – 6.59 (m, 4H), 4.60 (s, 2H), 3.30 – 3.20 (m, 4H), 2.90 (s, 12H), 1.82 – 1.70 (m, 2H), 0.62 (s, 3H), 0.51 (s, 3H). HRMS (ESI) $m/z$ calculated 797.3482 for $C_{45}H_{49}N_6O_6Si$ $[M+H]^+$, found 797.3478.

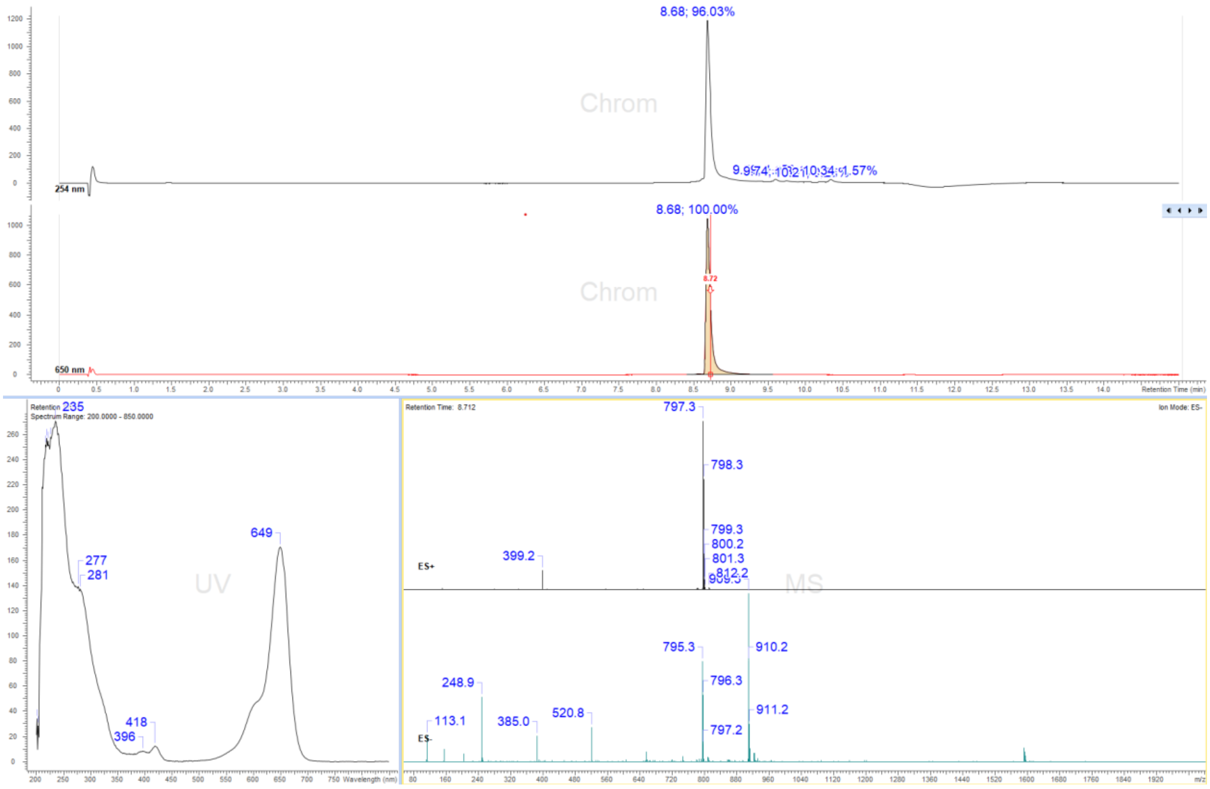

### Display Report

**Analysis Info**

| | | | |
|---|---|---|---|
| Analysis Name | Z:\Data\2024\2409\sam030924\VTP20_13_01_127386.d | Acquisition Date | 03.09.2024 18:56:23 |
| Method | hystar_pl.m | Operator | BDAL@DE |
| Sample Name | VTP20 | Instrument / Ser# | micrOTOF 10237 |
| Comment | | | |

**Acquisition Parameter**

| | | | | | |
|---|---|---|---|---|---|
| Source Type | ESI | Ion Polarity | Positive | Set Nebulizer | 1.2 Bar |
| Focus | Not active | | | Set Dry Heater | 180 °C |
| Scan Begin | 50 m/z | Set Capillary | 4500 V | Set Dry Gas | 4.0 l/min |
| Scan End | 1600 m/z | Set End Plate Offset | -500 V | Set Divert Valve | Source |

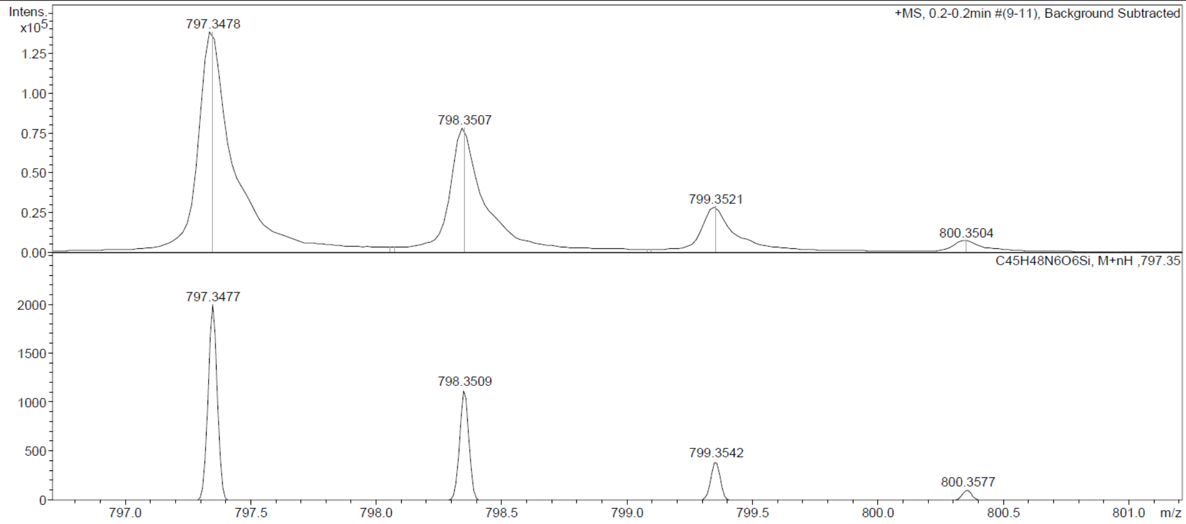

Probe **14:**

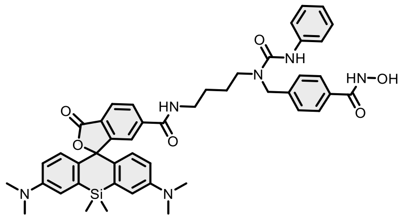

Reaction was conducted and purified following general method C. Fractions containing the product were collected, evaporated and lyophilized from acetonitrile: water mixture. The obtained solid was dissolved in 700 µL of d6-DMSO. The samples from DMSO solution were diluted x100 in PBS +0.1%SDS and concentration was determined spectroscopically with Nanodrop. Determined stock concentration was 1.93 mM which constitutes to 33% yield (1.10 mg).

$^{1}$H NMR (400 MHz, DMSO-$d_6$) δ 11.14 (s, 1H), 8.97 (s, 1H), 8.75 (s, 1H), 8.32 (s, 1H), 8.08 – 7.98 (m, 2H), 7.69 (d, $J$ = 8.2 Hz, 2H), 7.64 (t, $J$ = 1.1 Hz, 1H), 7.45 – 7.39 (m, 2H), 7.28 (d, $J$ = 8.2 Hz, 2H), 7.20 – 7.14 (m, 2H), 7.02 – 6.98 (m, 2H), 6.91 (t, $J$ = 7.3 Hz, 1H), 6.67 – 6.58 (m, 4H), 4.58 (s, 2H), 3.28 – 3.16 (m, 4H), 2.90 (s, 12H), 1.48 (s, 4H), 0.62 (s, 3H), 0.51 (s, 3H). HRMS (ESI) $m/z$ calculated 811.3639 for $C_{46}H_{51}N_6O_6Si$, found 811.3644 ($[M+H]^+$).

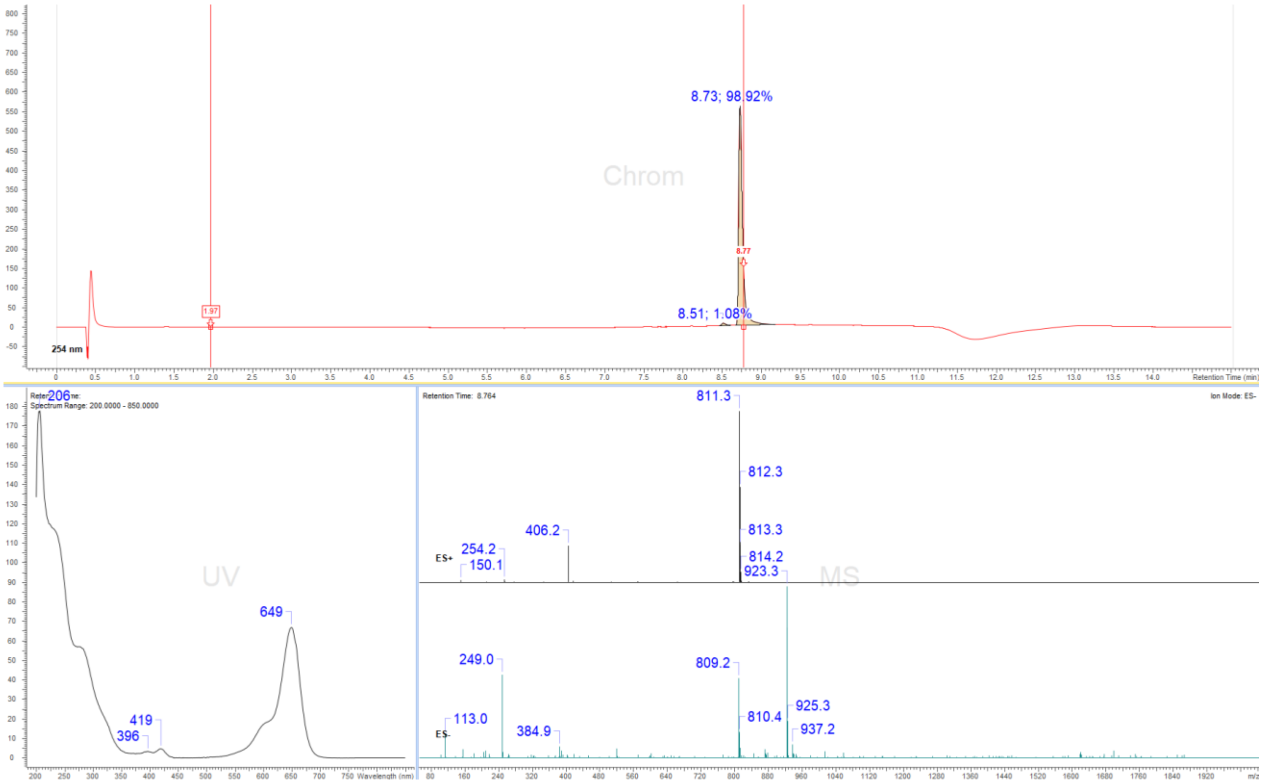

### Display Report

**Analysis Info**

| | | | |
|---|---|---|---|
| Analysis Name | Z:\Data\2024\2409\sam030924\VTP9_14_01_127387.d | Acquisition Date | 03.09.2024 19:01:01 |
| Method | hystar_pl.m | Operator | BDAL@DE |
| Sample Name | VTP9 | Instrument / Ser# | micrOTOF 10237 |
| Comment | | | |

**Acquisition Parameter**

| | | | | | |
|---|---|---|---|---|---|
| Source Type | ESI | Ion Polarity | Positive | Set Nebulizer | 1.2 Bar |
| Focus | Not active | | | Set Dry Heater | 180 °C |
| Scan Begin | 50 m/z | Set Capillary | 4500 V | Set Dry Gas | 4.0 l/min |
| Scan End | 1600 m/z | Set End Plate Offset | -500 V | Set Divert Valve | Source |

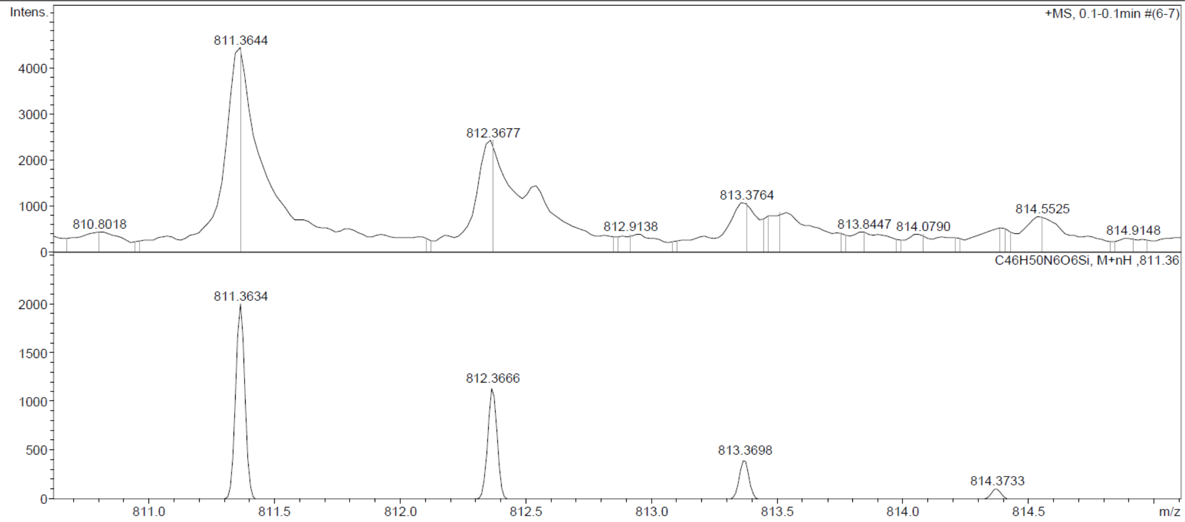

Probe **15:**

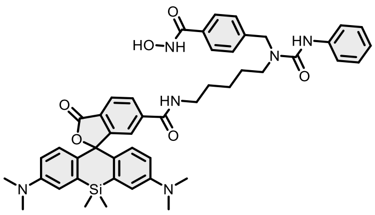

Reaction was conducted and purified following general method C. Fractions containing the product were collected, evaporated and lyophilized from acetonitrile: water mixture. The obtained solid was dissolved in 700 µL of $d_6$-DMSO. The samples from DMSO solution were diluted x100 in PBS +0.1%SDS and concentration was determined spectroscopically with Nanodrop. Determined stock concentration was 1.1 mM which constitutes to 19% yield (0.66 mg).

$^{1}$H NMR (400 MHz, DMSO-$d_6$) δ 11.14 (s, 1H), 8.97 (s, 1H), 8.70 (s, 1H), 8.32 (s, 1H), 8.09 – 7.96 (m, 2H), 7.69 (d, $J$ = 8.3 Hz, 2H), 7.63 (s, 1H), 7.43 (d, $J$ = 7.6 Hz, 2H), 7.28 (d, $J$ = 8.1 Hz, 2H), 7.24 – 7.13 (m, 3H), 7.00 (d, $J$ = 2.4 Hz, 2H), 6.92 (t, $J$ = 7.4 Hz, 1H), 6.64 – 6.61 (m, 3H), 4.58 (s, 2H), 3.25 – 3.14 (m, 4H), 2.90 (s, 12H), 1.46 (d, $J$ = 7.4 Hz, 6H), 0.62 (s, 3H), 0.51 (s, 3H). HRMS (ESI) $m/z$ calculated 825.3795 for $C_{47}H_{53}N_6O_6Si$ $[M+H]^+$, found 825.3790.

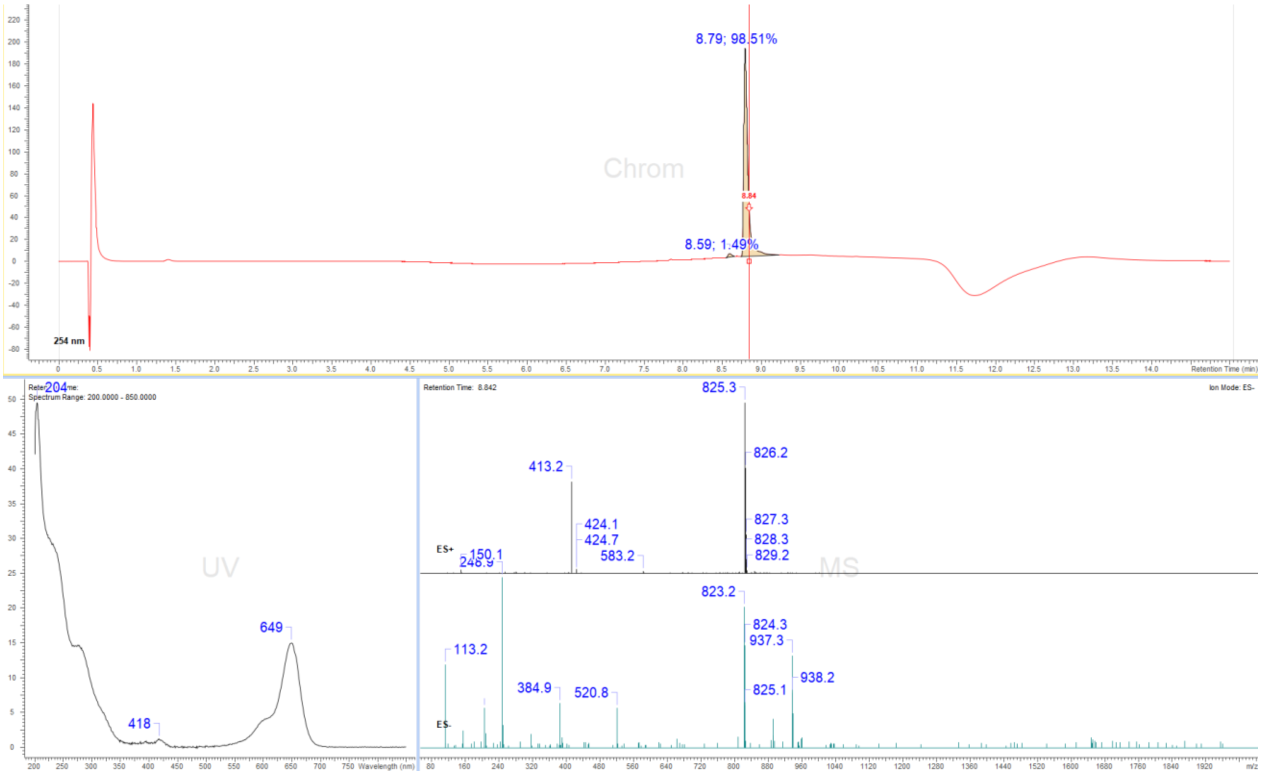

### Display Report

**Analysis Info**

| | | | |
|---|---|---|---|
| Analysis Name | Z:\Data\2024\2409\sam030924\VTP10_15_01_127388.d | Acquisition Date | 03.09.2024 19:05:39 |
| Method | hystar_pl.m | Operator | BDAL@DE |
| Sample Name | VTP10 | Instrument / Ser# | micrOTOF 10237 |
| Comment | | | |

**Acquisition Parameter**

| | | | | | |
|---|---|---|---|---|---|
| Source Type | ESI | Ion Polarity | Positive | Set Nebulizer | 1.2 Bar |
| Focus | Not active | | | Set Dry Heater | 180 °C |
| Scan Begin | 50 m/z | Set Capillary | 4500 V | Set Dry Gas | 4.0 l/min |
| Scan End | 1600 m/z | Set End Plate Offset | -500 V | Set Divert Valve | Source |

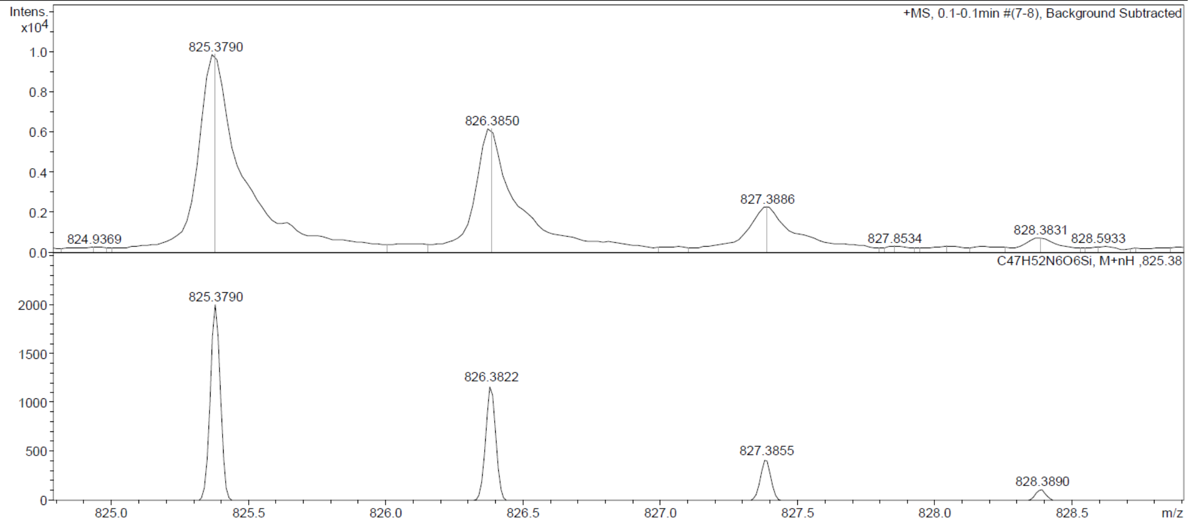

Probe **16:**

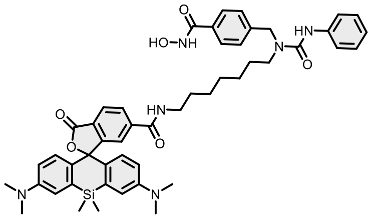

Reaction was conducted and purified following general method C. Fractions containing the product were collected, evaporated and lyophilized from acetonitrile: water mixture. The obtained solid was dissolved in 700 µL of $d_6$-DMSO. The samples from DMSO solution were diluted x100 in PBS +0.1%SDS and concentration was determined spectroscopically with Nanodrop. Determined stock concentration was 2.11 mM which constitutes 35% yield (1.26 mg).

$^{1}$H NMR (400 MHz, DMSO-$d_6$) δ 11.15 (s, 1H), 8.98 (s, 1H), 8.68 (s, 1H), 8.32 (s, 1H), 8.08 – 7.98 (m, 2H), 7.70 (d, $J$ = 8.3 Hz, 2H), 7.63 (s, 1H), 7.47 – 7.41 (m, 2H), 7.29 (d, $J$ = 8.2 Hz, 2H), 7.22 – 7.18 (m, 2H), 7.00 (d, $J$ = 2.5 Hz, 2H), 6.92 (t, $J$ = 7.3 Hz, 1H), 6.64 – 6.61 (m, 4H), 4.58 (s, 2H), 3.17 (t, $J$ = 5.4 Hz, 4H), 2.90 (s, 12H), 1.48 – 1.43 (m, 4H), 1.24 – 1.22 (m, 6H), 0.62 (s, 3H), 0.51 (s, 3H). HRMS (ESI) $m/z$ calculated 853.4108 for $C_{49}H_{57}N_6O_6Si$, found 853.4108 ([M+H]$^+$).

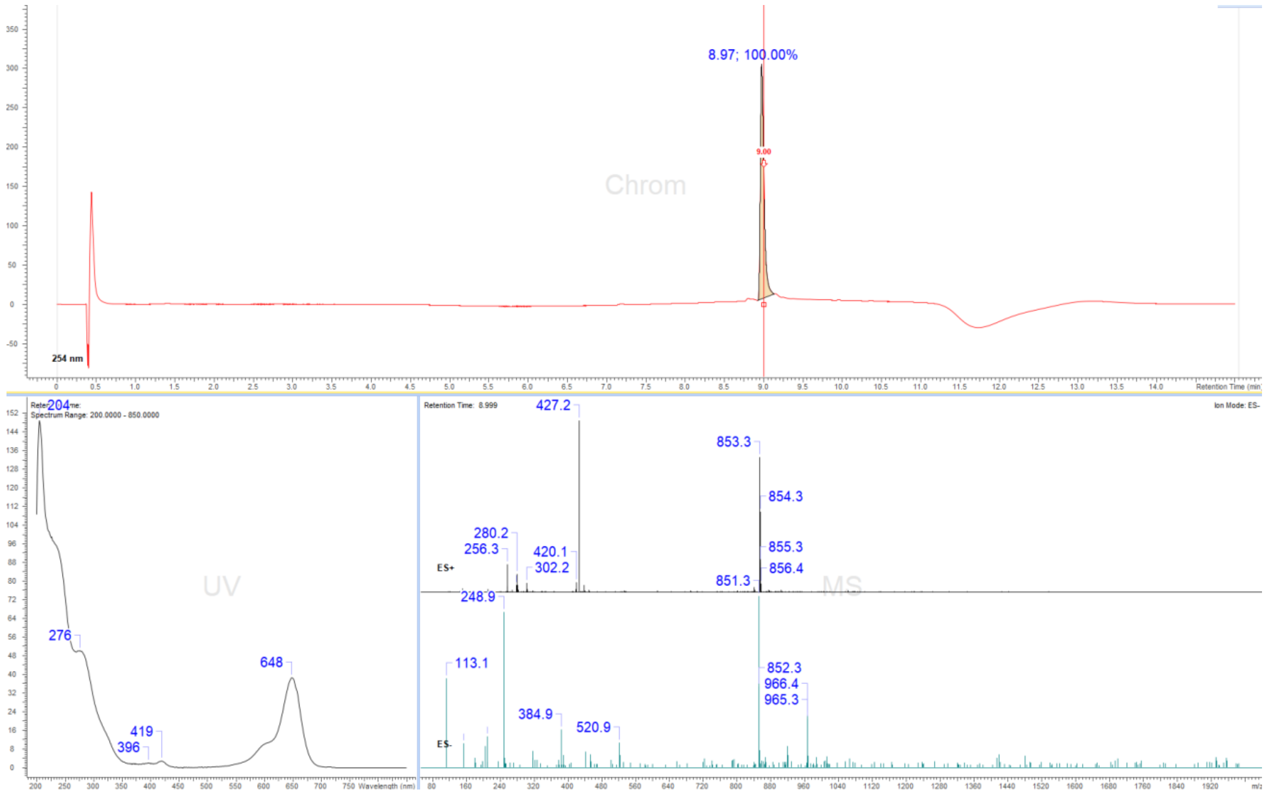

### Display Report

**Analysis Info**

| | | | |
|---|---|---|---|
| Analysis Name | Z:\Data\2024\2409\sam030924\VTP11_5_01_127378.d | Acquisition Date | 03.09.2024 18:19:09 |
| Method | hystar_pl.m | Operator | BDAL@DE |
| Sample Name | VTP11 | Instrument / Ser# | micrOTOF 10237 |
| Comment | | | |

**Acquisition Parameter**

| | | | | | |
|---|---|---|---|---|---|
| Source Type | ESI | Ion Polarity | Positive | Set Nebulizer | 1.2 Bar |
| Focus | Not active | | | Set Dry Heater | 180 °C |
| Scan Begin | 50 m/z | Set Capillary | 4500 V | Set Dry Gas | 4.0 l/min |
| Scan End | 1600 m/z | Set End Plate Offset | -500 V | Set Divert Valve | Source |

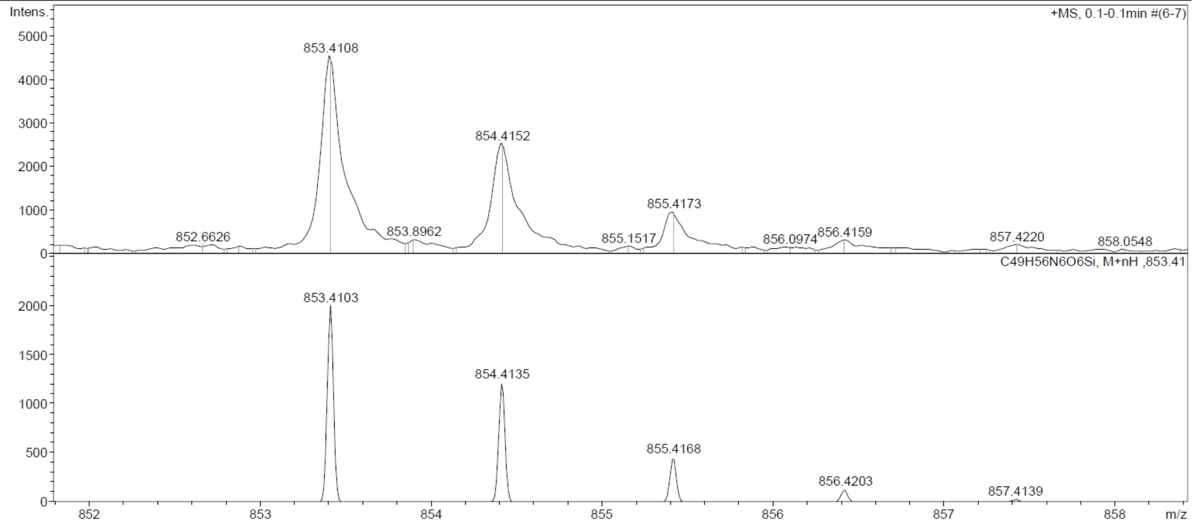

Probe **17:**

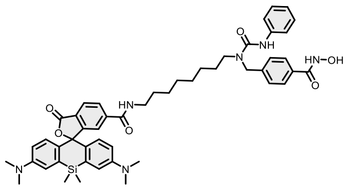

Reaction was conducted and purified following general method C. Fractions containing the product were collected, evaporated and lyophilized from acetonitrile: water mixture. The obtained solid was dissolved in 700 µL of $d_6$-DMSO. The samples from DMSO solution were diluted x100 in PBS +0.1%SDS and concentration was determined spectroscopically with Nanodrop. Determined stock concentration was 1.63 mM which constitutes to 27% yield (0.99 mg).

$^1$H NMR (400 MHz, DMSO-$d_6$) δ 11.15 (s, 1H), 8.98 (s, 1H), 8.68 (s, 1H), 8.33 (s, 1H), 8.07 – 7.99 (m, 2H), 7.70 (d, $J$ = 8.3 Hz, 2H), 7.63 (s, 1H), 7.44 (dd, $J$ = 8.7, 1.2 Hz, 2H), 7.30 (d, $J$ = 8.2 Hz, 2H), 7.22 – 7.18 (m, 2H), 7.00 (d, $J$ = 2.5 Hz, 2H), 6.92 (t, $J$ = 7.3 Hz, 1H), 6.64 – 6.61 (m, 4H), 4.58 (s, 2H), 3.17 (t, $J$ = 5.7 Hz, 4H), 2.90 (s, 12H), 1.45 (s, 4H), 1.22 (d, $J$ = 7.6 Hz, 8H), 0.63 (s, 3H), 0.51 (s, 3H). HRMS (ESI) $m/z$ calculated 867.4265 for $C_{50}H_{59}N_6O_6Si$, found 867.4267 ($[M+H]^+$).

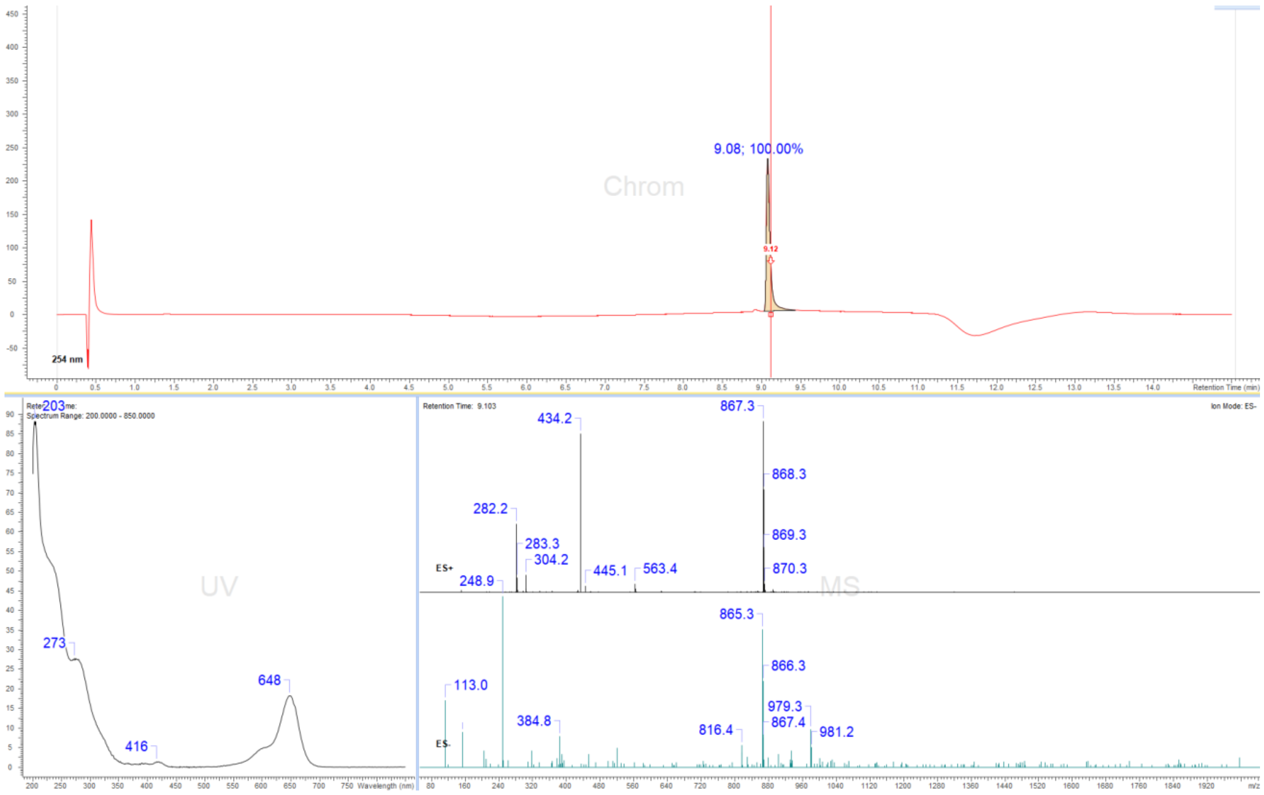

#### Display Report

**Analysis Info**

| | | | |
|---|---|---|---|
| Analysis Name | Z:\Data\2024\2409\sam030924\VTP12_6_01_127379.d | Acquisition Date | 03.09.2024 18:23:46 |
| Method | hystar_pl.m | Operator | BDAL@DE |
| Sample Name | VTP12 | Instrument / Ser# | micrOTOF 10237 |
| Comment | | | |

**Acquisition Parameter**

| | | | | | |
|---|---|---|---|---|---|
| Source Type | ESI | Ion Polarity | Positive | Set Nebulizer | 1.2 Bar |
| Focus | Not active | | | Set Dry Heater | 180 °C |
| Scan Begin | 50 m/z | Set Capillary | 4500 V | Set Dry Gas | 4.0 l/min |
| Scan End | 1600 m/z | Set End Plate Offset | -500 V | Set Divert Valve | Source |

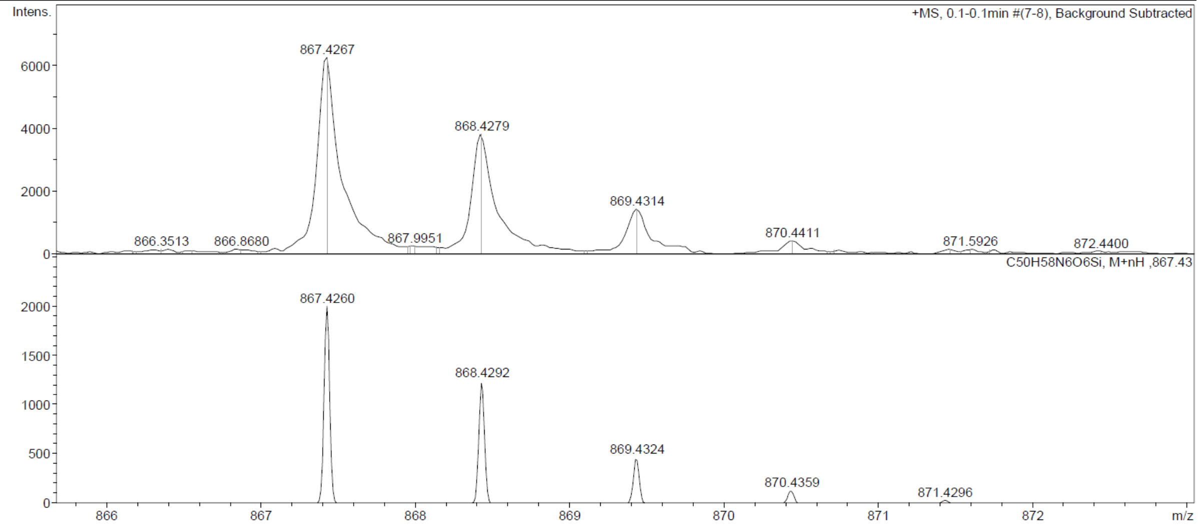

Probe **18:**

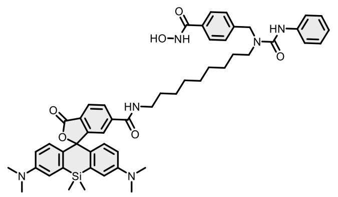

Reaction was conducted and purified following general method C. Fractions containing the product were collected, evaporated and lyophilized from acetonitrile: water mixture. The obtained solid was dissolved in 700 µL of $d_6$-DMSO. The samples from DMSO solution were diluted x100 in PBS +0.1%SDS and concentration was determined spectroscopically with Nanodrop. Determined stock concentration was 1.87 mM which constitutes 31% yield (1.15 mg).

$^1$H NMR (400 MHz, DMSO-$d_6$) δ 11.15 (s, 1H), 8.98 (s, 1H), 8.68 (t, $J$ = 5.6 Hz, 1H), 8.33 (s, 1H), 8.06 – 7.99 (m, 2H), 7.73 – 7.69 (m, 2H), 7.63 (dd, $J$ = 1.4, 0.8 Hz, 1H), 7.46 – 7.42 (m, 2H), 7.30 (d, $J$ = 8.2 Hz, 2H), 7.23 – 7.18 (m, 2H), 7.01 (dd, $J$ = 2.6, 0.8 Hz, 2H), 6.94 – 6.89 (m, 1H), 6.65 – 6.60 (m, 4H), 4.59 (s, 2H), 3.17 (t, $J$ = 5.9 Hz, 4H), 2.90 (s, 12H), 1.44 (s, 4H), 1.21 (d, $J$ = 11.6 Hz, 10H), 0.63 (s, 3H), 0.51 (s, 3H). HRMS (ESI) $m/z$ calculated 881.4421 for $C_{51}H_{61}N_6O_6Si$ [M+H]$^+$, found 881.4410.

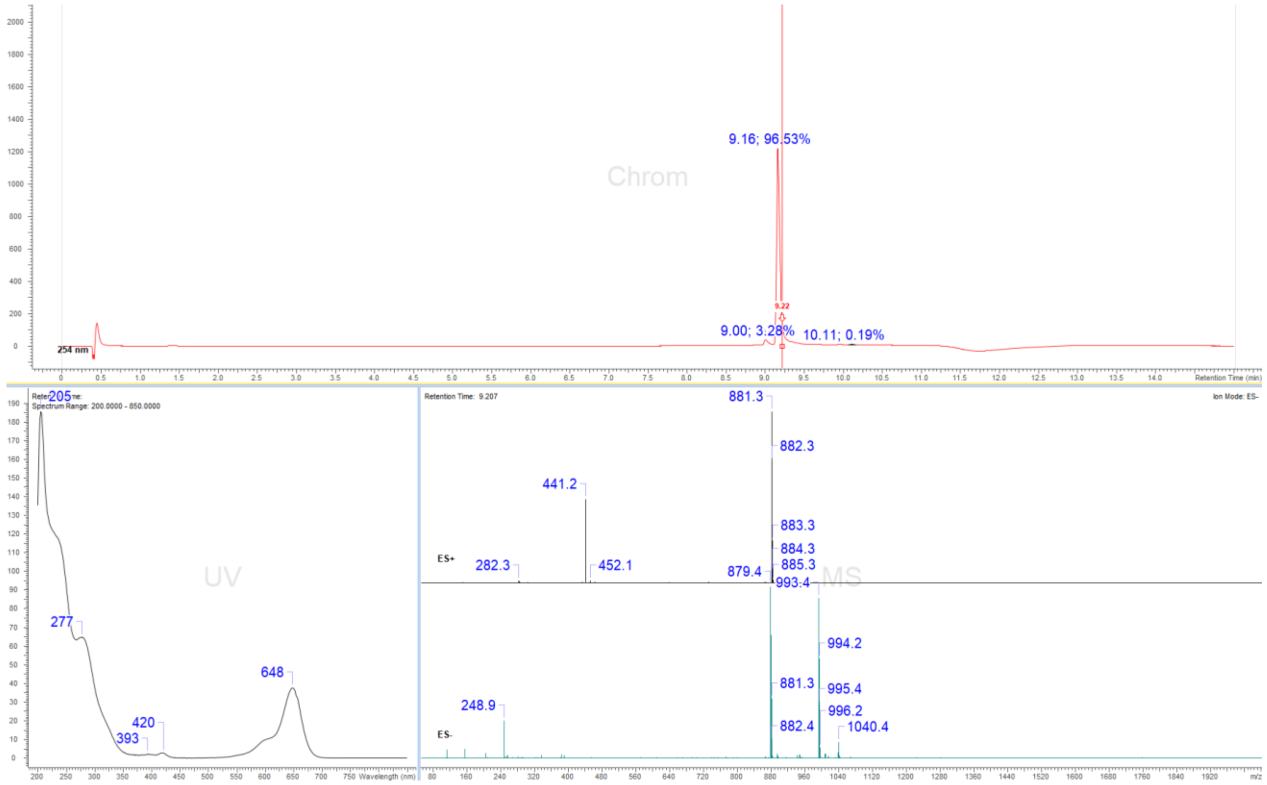

### Display Report

**Analysis Info**

| | | | |
|---|---|---|---|
| Analysis Name | Z:\Data\2024\2409\sam030924\VTP1_7_01_127380.d | Acquisition Date | 03.09.2024 18:28:25 |
| Method | hystar_pl.m | Operator | BDAL@DE |
| Sample Name | VTP1 | Instrument / Ser# | micrOTOF 10237 |
| Comment | | | |

**Acquisition Parameter**

| | | | | | |
|---|---|---|---|---|---|
| Source Type | ESI | Ion Polarity | Positive | Set Nebulizer | 1.2 Bar |
| Focus | Not active | | | Set Dry Heater | 180 °C |
| Scan Begin | 50 m/z | Set Capillary | 4500 V | Set Dry Gas | 4.0 l/min |
| Scan End | 1600 m/z | Set End Plate Offset | -500 V | Set Divert Valve | Source |

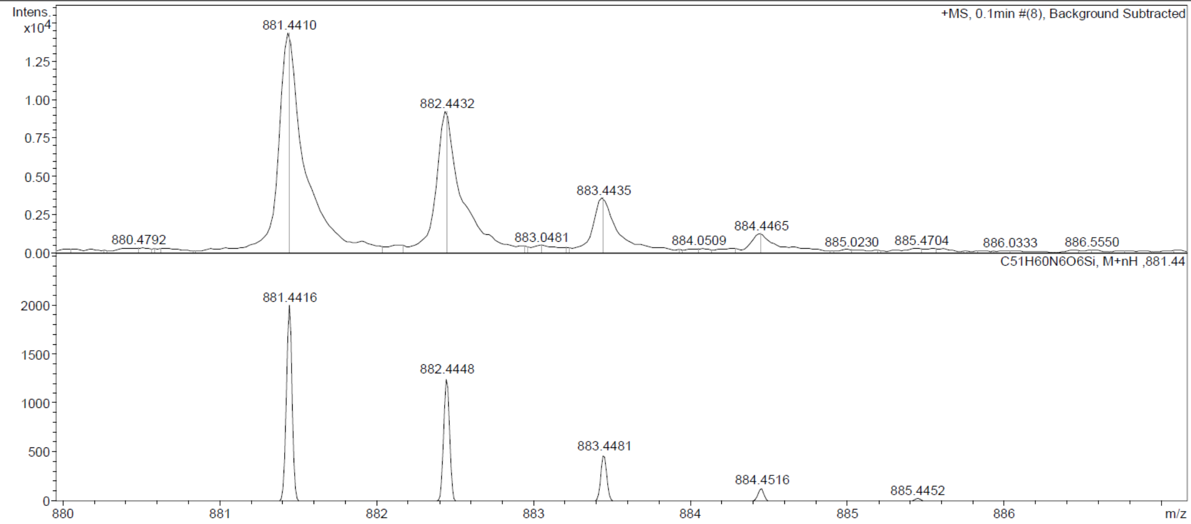

Probe **19:**

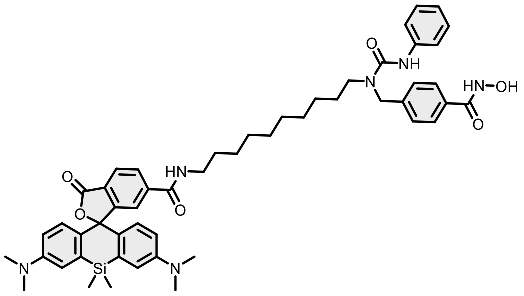

Reaction was conducted and purified following general method C. Fractions containing the product were collected, evaporated and lyophilized from acetonitrile: water mixture. The obtained solid was dissolved in 700 µL of $d_6$-DMSO. The samples from DMSO solution were diluted x100 in PBS +0.1%SDS and concentration was determined spectroscopically with Nanodrop. Determined stock concentration was 0.96 mM which constitutes to 16% yield (0.61 mg).

$^1$H NMR (400 MHz, DMSO-$d_6$) δ 11.15 (s, 1H), 8.98 (s, 1H), 8.68 (t, $J$ = 5.6 Hz, 1H), 8.33 (s, 1H), 8.07 – 7.99 (m, 2H), 7.71 (d, $J$ = 8.3 Hz, 2H), 7.63 (t, $J$ = 1.1 Hz, 1H), 7.46 – 7.42 (m, 2H), 7.30 (d, $J$ = 8.2 Hz, 2H), 7.23 – 7.18 (m, 2H), 7.02 – 6.99 (m, 2H), 6.94 – 6.89 (m, 1H), 6.64 – 6.60 (m, 4H), 4.59 (s, 2H), 3.26 – 3.14 (m, 4H), 2.90 (s, 12H), 1.45 (s, 4H), 1.21 (d, $J$ = 12.9 Hz, 12H), 0.63 (s, 3H), 0.51 (s, 3H). HRMS (ESI) $m/z$ calculated 895.4578 for $C_{52}H_{63}N_6O_6Si$ $[M+H]^+$, found 895.4551.

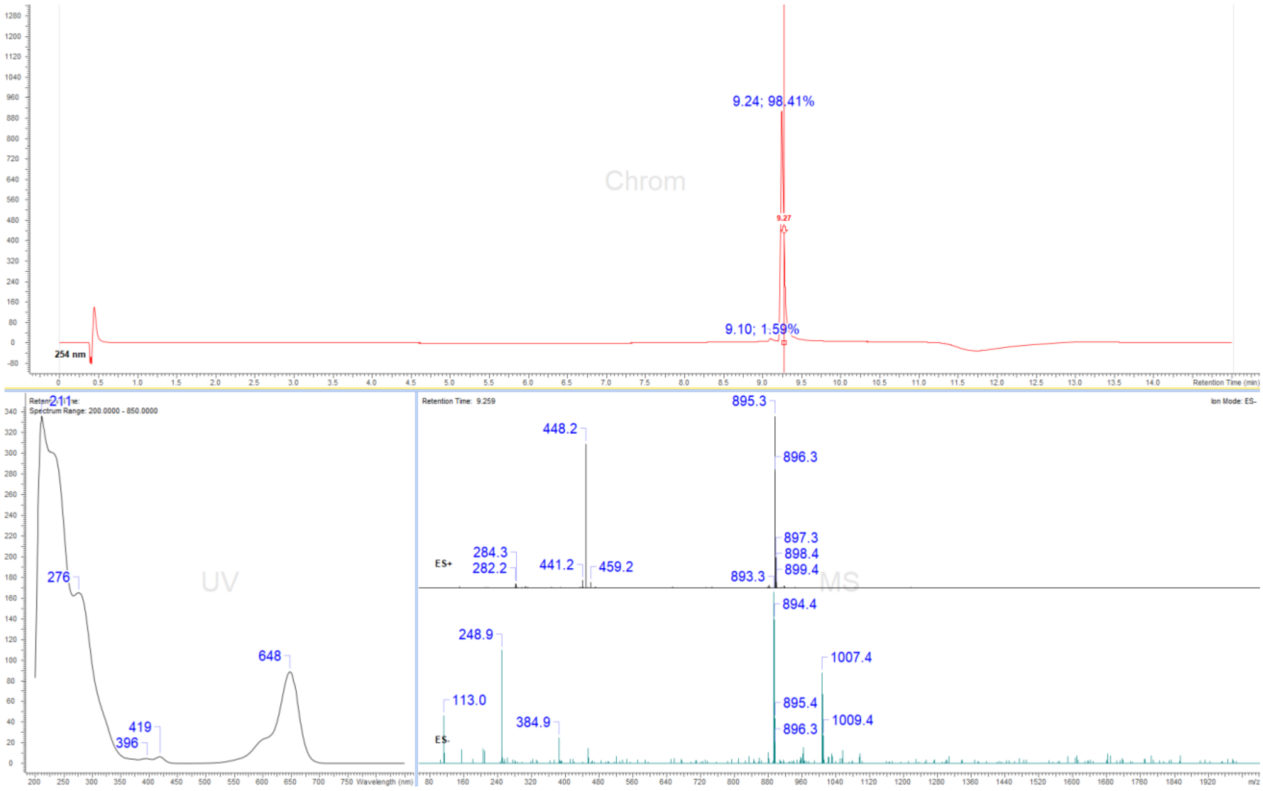

### Display Report

| Analysis Info | | | |
|---|---|---|---|
| Analysis Name | Z:\Data\2024\2409\sam030924\VTP13_8_01_127381.d | Acquisition Date | 03.09.2024 18:33:05 |
| Method | hystar_pl.m | Operator | BDAL@DE |
| Sample Name | VTP13 | Instrument / Ser# | micrOTOF 10237 |
| Comment | | | |

**Acquisition Parameter**

| | | | | | |
|---|---|---|---|---|---|
| Source Type | ESI | Ion Polarity | Positive | Set Nebulizer | 1.2 Bar |
| Focus | Not active | | | Set Dry Heater | 180 °C |
| Scan Begin | 50 m/z | Set Capillary | 4500 V | Set Dry Gas | 4.0 l/min |
| Scan End | 1600 m/z | Set End Plate Offset | -500 V | Set Divert Valve | Source |

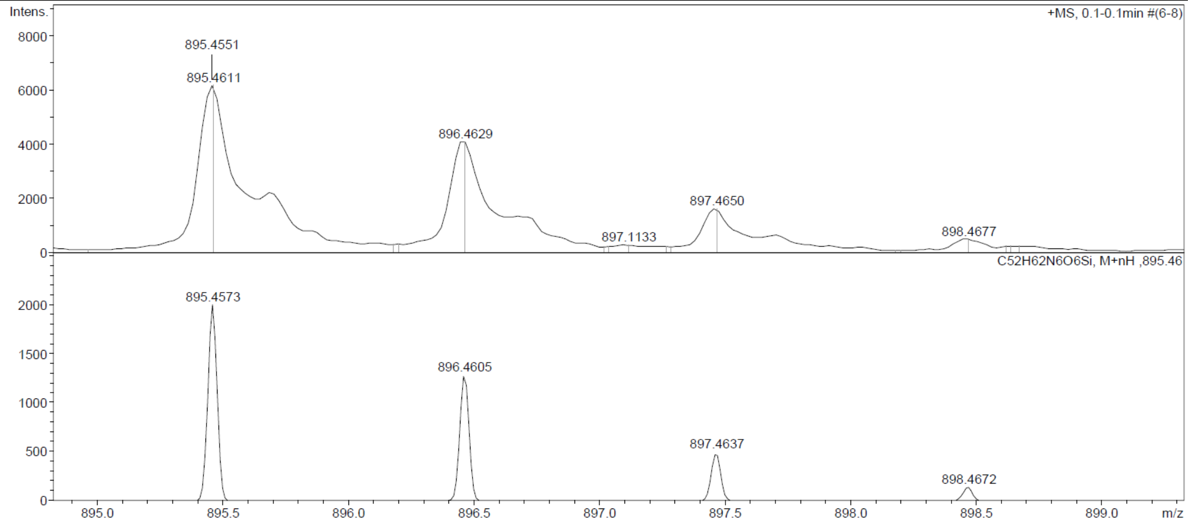

Probe **20:**

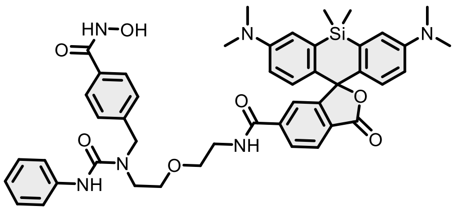

Reaction was conducted and purified following general method C. Fractions containing the product were collected, evaporated and lyophilized from acetonitrile: water mixture. The obtained solid was dissolved in 700 µL of $d_6$-DMSO. The samples from DMSO solution were diluted x100 in PBS +0.1%SDS and concentration was determined spectroscopically with Nanodrop. Determined stock concentration was 1.69 mM which constitutes 28% yield (0.98 mg).

$^1$H NMR (400 MHz, DMSO-$d_6$) δ 11.14 (s, 1H), 8.98 (s, 1H), 8.80 (t, $J$ = 5.5 Hz, 1H), 8.36 (s, 1H), 8.08 (dd, $J$ = 8.0, 1.4 Hz, 1H), 8.00 (dd, $J$ = 8.0, 0.7 Hz, 1H), 7.72 – 7.66 (m, 3H), 7.39 – 7.34 (m, 2H), 7.25 (d, $J$ = 8.2 Hz, 2H), 7.15 (dd, $J$ = 8.6, 7.4 Hz, 2H), 7.00 (t, $J$ = 1.7 Hz, 2H), 6.92 – 6.86 (m, 1H), 6.61 – 6.57 (m, 4H), 4.59 (s, 2H), 3.54 (dt, $J$ = 8.9, 5.4 Hz, 4H), 3.46 – 3.39 (m, 4H), 2.89 (s, 12H), 0.62 (s, 3H), 0.50 (s, 3H). HRMS (ESI) $m/z$ calculated 827.3588 for $C_{46}H_{51}N_6O_7Si$ $[M+H]^+$, found 827.3597.

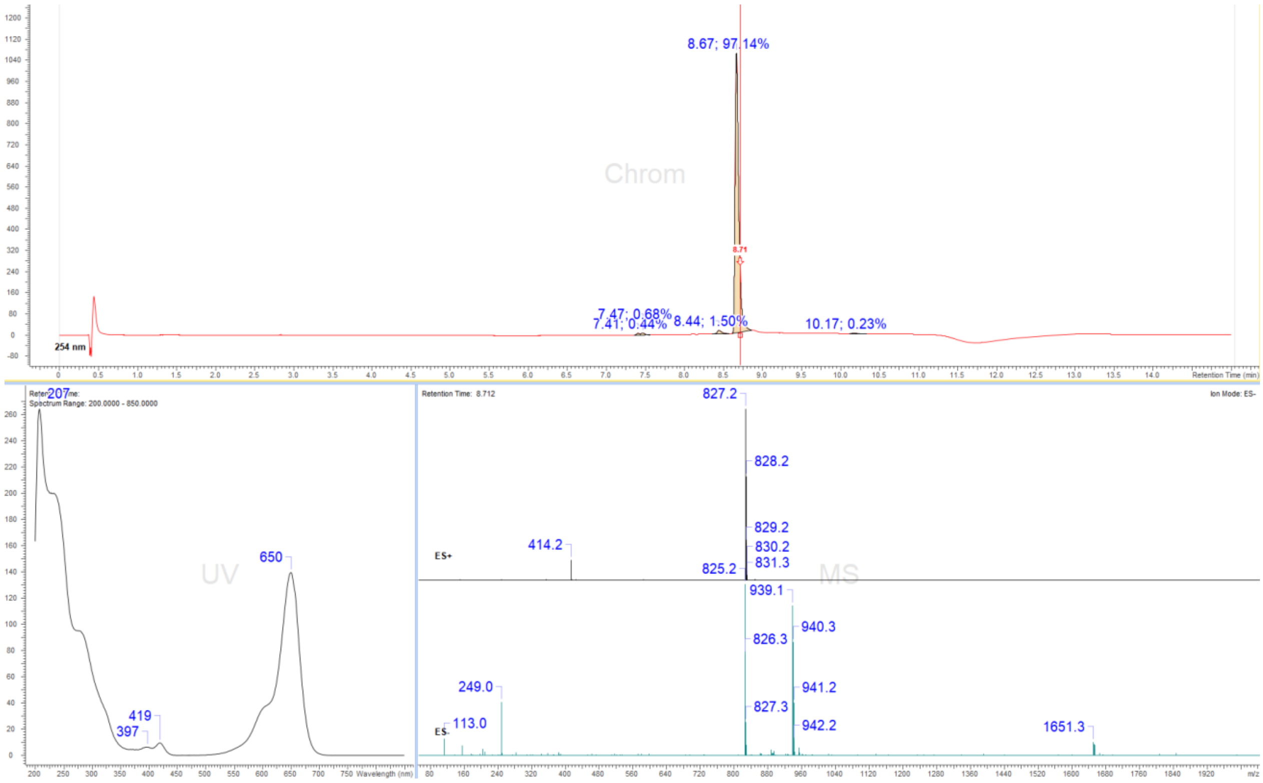

### Display Report

**Analysis Info**

| | | | |
|---|---|---|---|
| Analysis Name | Z:\Data\2024\2409\sam030924\VTP5_9_01_127382.d | Acquisition Date | 03.09.2024 18:37:44 |
| Method | hystar_pl.m | Operator | BDAL@DE |
| Sample Name | VTP5 | Instrument / Ser# | micrOTOF 10237 |
| Comment | | | |

**Acquisition Parameter**

| | | | | | |
|---|---|---|---|---|---|
| Source Type | ESI | Ion Polarity | Positive | Set Nebulizer | 1.2 Bar |
| Focus | Not active | | | Set Dry Heater | 180 °C |
| Scan Begin | 50 m/z | Set Capillary | 4500 V | Set Dry Gas | 4.0 l/min |
| Scan End | 1600 m/z | Set End Plate Offset | -500 V | Set Divert Valve | Source |

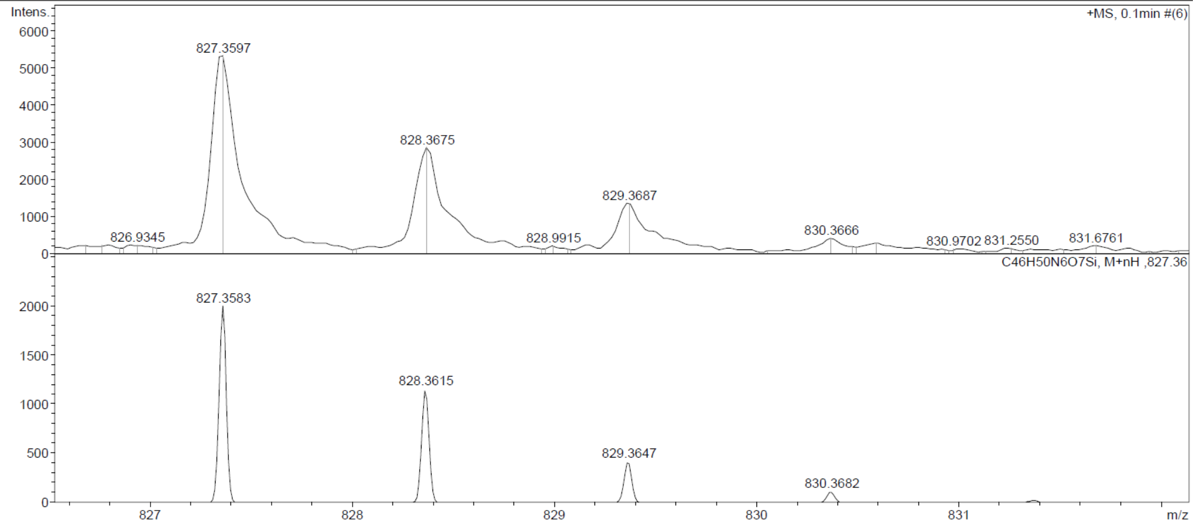

Probe **21:**

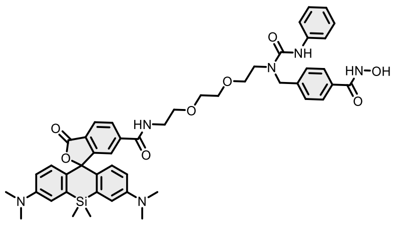

Reaction was conducted and purified following general method C. Fractions containing the product were collected, evaporated and lyophilized from acetonitrile: water mixture. The obtained solid was dissolved in 700 µL of $d_6$-DMSO. The samples from DMSO solution were diluted x100 in PBS +0.1%SDS and concentration was determined spectroscopically with Nanodrop. Determined stock concentration was 1.21 mM which constitutes to 20% yield (0.74 mg).

$^1$H NMR (400 MHz, DMSO-$d_6$) δ 11.15 (s, 1H), 8.79 (t, $J$ = 5.6 Hz, 1H), 8.42 (s, 1H), 8.10 – 7.99 (m, 2H), 7.74 – 7.68 (m, 2H), 7.66 (s, 1H), 7.40 – 7.36 (m, 2H), 7.30 (d, $J$ = 8.2 Hz, 2H), 7.22 – 7.17 (m, 2H), 7.02 (s, 2H), 6.89 (tt, $J$ = 7.4, 1.2 Hz, 1H), 6.67 – 6.58 (m, 4H), 4.59 (s, 2H), 3.51 (d, $J$ = 4.7 Hz, 8H), 3.44 (d, $J$ = 5.0 Hz, 4H), 2.91 (s, 12H), 0.63 (s, 3H), 0.51 (s, 3H). HRMS (ESI) $m/z$ calculated 870.3772 for $C_{48}H_{54}N_6O_8Si$ $[M+H]^+$, found 871.3847.

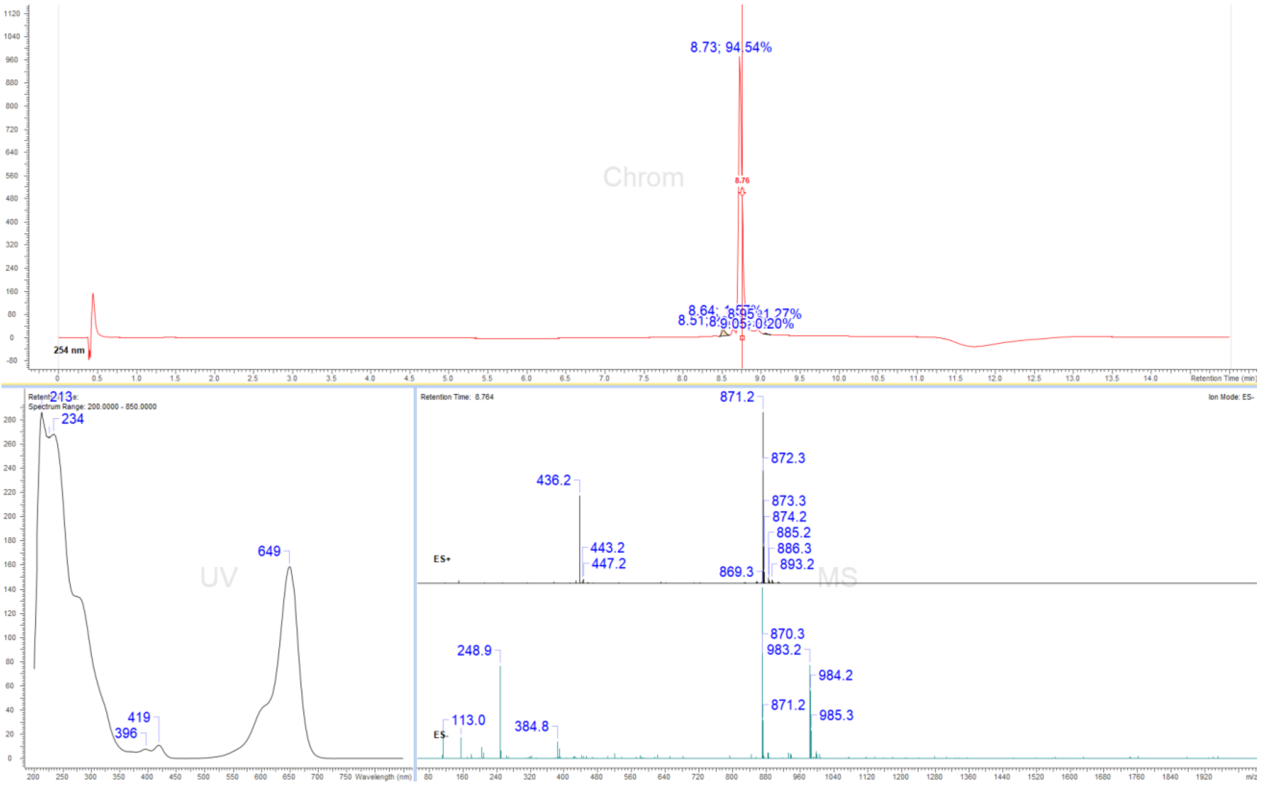

#### Display Report

| Analysis Info | | Acquisition Date | 03.09.2024 18:42:25 | |
|---|---|---|---|---|
| Analysis Name | Z:\Data\2024\2409\sam030924\VTP6_10_01_127383.d | | | |
| Method | hystar_pl.m | Operator | BDAL@DE | |
| Sample Name | VTP6 | Instrument / Ser# | micrOTOF | 10237 |
| Comment | | | | |

**Acquisition Parameter**

| | | | | | |
|---|---|---|---|---|---|
| Source Type | ESI | Ion Polarity | Positive | Set Nebulizer | 1.2 Bar |
| Focus | Not active | | | Set Dry Heater | 180 °C |
| Scan Begin | 50 m/z | Set Capillary | 4500 V | Set Dry Gas | 4.0 l/min |
| Scan End | 1600 m/z | Set End Plate Offset | -500 V | Set Divert Valve | Source |

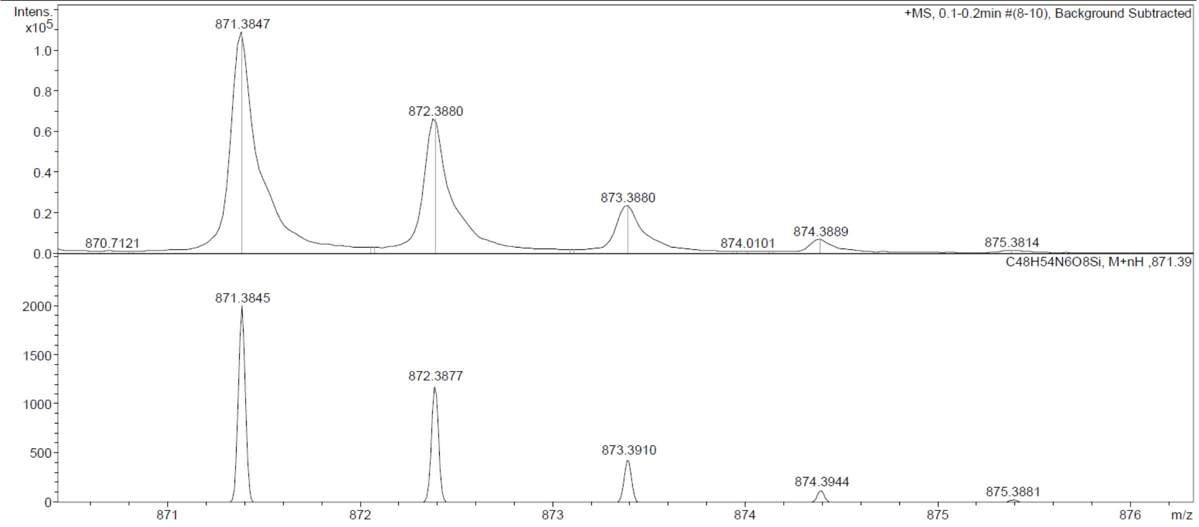

Probe **22:**

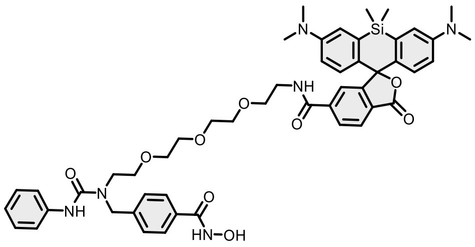

Reaction was conducted and purified following general method C. Fractions containing the product were collected, evaporated and lyophilized from acetonitrile: water mixture. The obtained solid was dissolved in 700 µL of $d_6$-DMSO. The samples from DMSO solution were diluted x100 in PBS +0.1%SDS and concentration was determined spectroscopically with Nanodrop. Determined stock concentration was 1.99 mM which constitutes to 33% yield (1.28 mg).

$^1$H NMR (400 MHz, DMSO-$d_6$) δ 11.15 (s, 1H), 8.98 (s, 1H), 8.77 (t, $J$ = 5.5 Hz, 1H), 8.43 (s, 1H), 8.07 (dd, $J$ = 8.0, 1.4 Hz, 1H), 8.01 (dd, $J$ = 8.0, 0.7 Hz, 1H), 7.71 (d, $J$ = 8.3 Hz, 2H), 7.65 (dd, $J$ = 1.4, 0.8 Hz, 1H), 7.41 – 7.37 (m, 2H), 7.31 (d, $J$ = 8.2 Hz, 2H), 7.19 (dd, $J$ = 8.6, 7.3 Hz, 2H), 7.02 – 6.99 (m, 2H), 6.91 – 6.86 (m, 1H), 6.63 – 6.58 (m, 4H), 4.60 (s, 2H), 3.46 (d, $J$ = 5.5 Hz, 12H), 3.34 (s, 4H), 2.90 (s, 12H), 0.63 (s, 3H), 0.51 (s, 3H). HRMS (ESI) $m/z$ calculated 915.4112 for $C_{50}H_{59}N_6O_9Si$ $[M+H]^+$, found 915.4107.

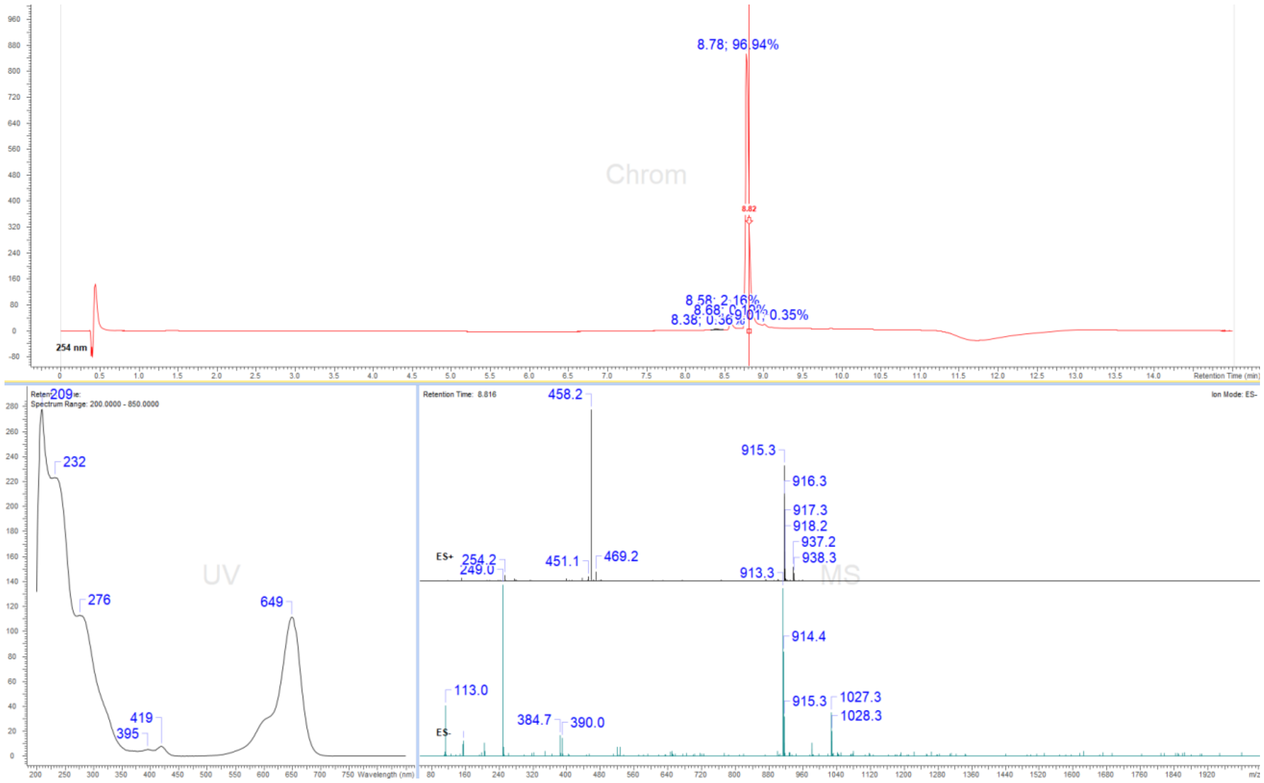

### Display Report

**Analysis Info**

| | | | |
|---|---|---|---|
| Analysis Name | Z:\Data\2024\2409\sam030924\VTP7_11_01_127384.d | Acquisition Date | 03.09.2024 18:47:05 |
| Method | hystar_pl.m | Operator | BDAL@DE |
| Sample Name | VTP7 | Instrument / Ser# | micrOTOF 10237 |
| Comment | | | |

**Acquisition Parameter**

| | | | | | |
|---|---|---|---|---|---|
| Source Type | ESI | Ion Polarity | Positive | Set Nebulizer | 1.2 Bar |
| Focus | Not active | | | Set Dry Heater | 180 °C |
| Scan Begin | 50 m/z | Set Capillary | 4500 V | Set Dry Gas | 4.0 l/min |
| Scan End | 1600 m/z | Set End Plate Offset | -500 V | Set Divert Valve | Source |

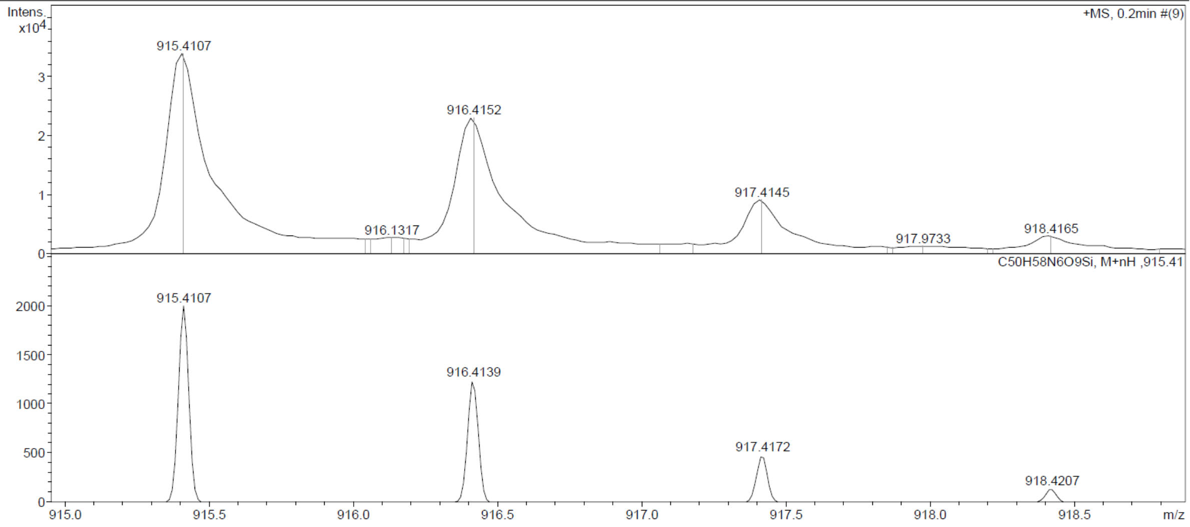

Probe **23:**

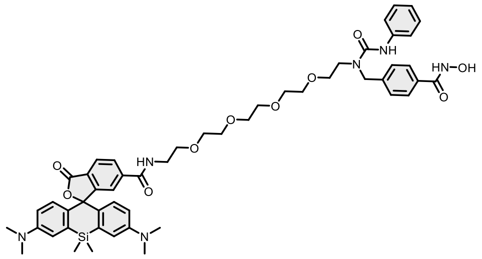

Reaction was conducted and purified following general method C. Fractions containing the product were collected, evaporated and lyophilized from acetonitrile: water mixture. The obtained solid was dissolved in 700 µL of $d_6$-DMSO. The samples from DMSO solution were diluted x100 in PBS +0.1%SDS and concentration was determined spectroscopically with Nanodrop. Determined stock concentration was 2.11 mM which constitutes 35% yield (1.42 mg).

$^1$H NMR (400 MHz, DMSO-$d_6$) δ 11.15 (s, 1H), 8.98 (s, 1H), 8.78 (t, $J$ = 5.5 Hz, 1H), 8.44 (s, 1H), 8.07 (dd, $J$ = 8.1, 1.4 Hz, 1H), 8.01 (dd, $J$ = 8.1, 0.7 Hz, 1H), 7.71 (d, $J$ = 8.2 Hz, 2H), 7.66 (t, $J$ = 1.1 Hz, 1H), 7.41 – 7.38 (m, 2H), 7.31 (d, $J$ = 8.2 Hz, 2H), 7.20 (dd, $J$ = 8.6, 7.3 Hz, 2H), 7.01 – 7.00 (m, 2H), 6.93 – 6.88 (m, 1H), 6.64 – 6.60 (m, 4H), 4.61 (s, 2H), 3.48 – 3.41 (m, 16H), 3.38 – 3.33 (m, 4H), 2.90 (s, 12H), 0.63 (s, 3H), 0.51 (s, 3H). HRMS (ESI) $m/z$ calculated 959.4374 for $C_{52}H_{63}N_6O_{10}Si$, found 959.4377 ([M+H]$^+$).

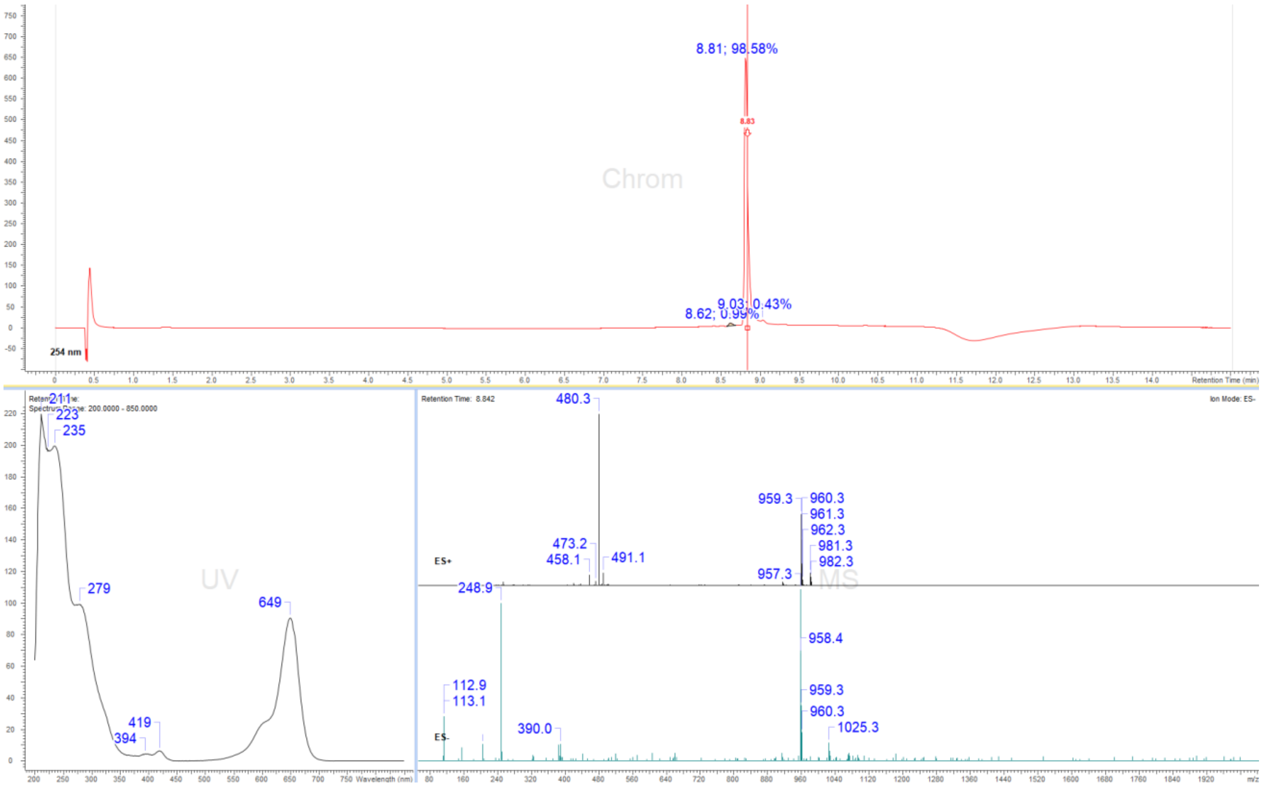

### Display Report

| | | | |
|---|---|---|---|
| **Analysis Info** | | Acquisition Date | 03.09.2024 18:14:30 |
| Analysis Name | Z:\Data\2024\2409\sam030924\VTP8_4_01_127377.d | | |
| Method | hystar_pl.m | Operator | BDAL@DE |
| Sample Name | VTP8 | Instrument / Ser# | micrOTOF 10237 |
| Comment | | | |

**Acquisition Parameter**

| | | | | | |
|---|---|---|---|---|---|
| Source Type | ESI | Ion Polarity | Positive | Set Nebulizer | 1.2 Bar |
| Focus | Not active | | | Set Dry Heater | 180 °C |
| Scan Begin | 50 m/z | Set Capillary | 4500 V | Set Dry Gas | 4.0 l/min |
| Scan End | 1600 m/z | Set End Plate Offset | -500 V | Set Divert Valve | Source |

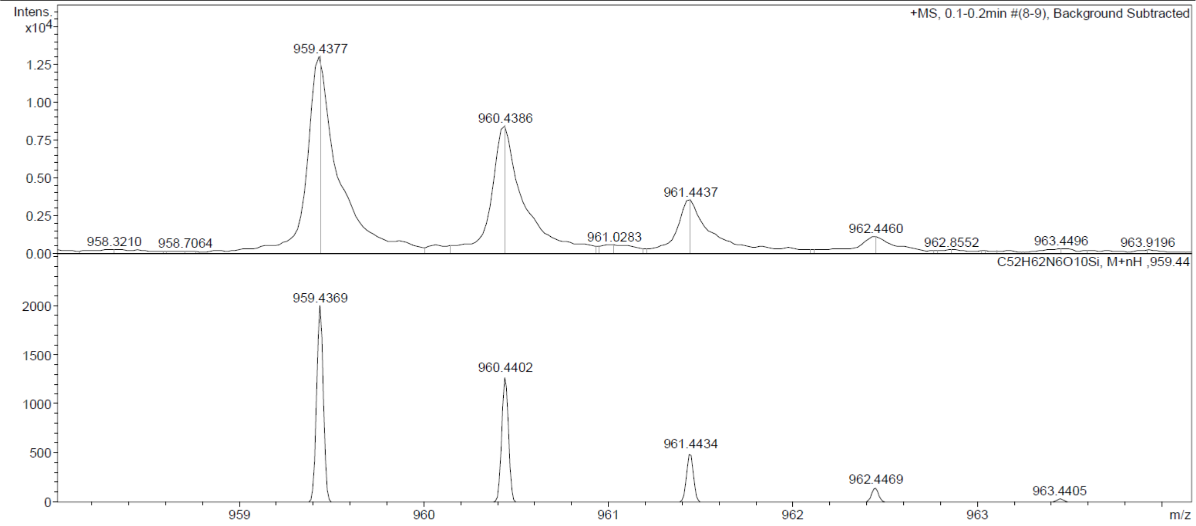

**Copies of NMR spectra**

Compound **1a**:

1H NMR spectrum

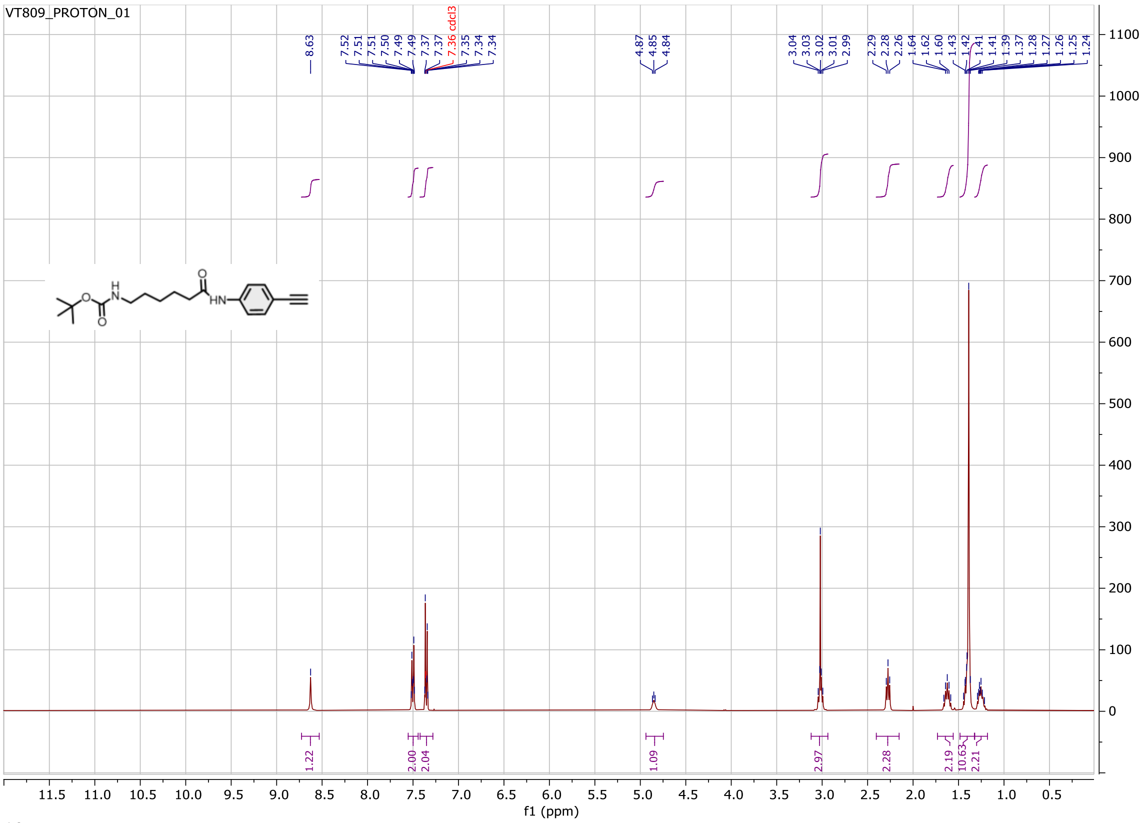

$^{13}C$ NMR spectrum

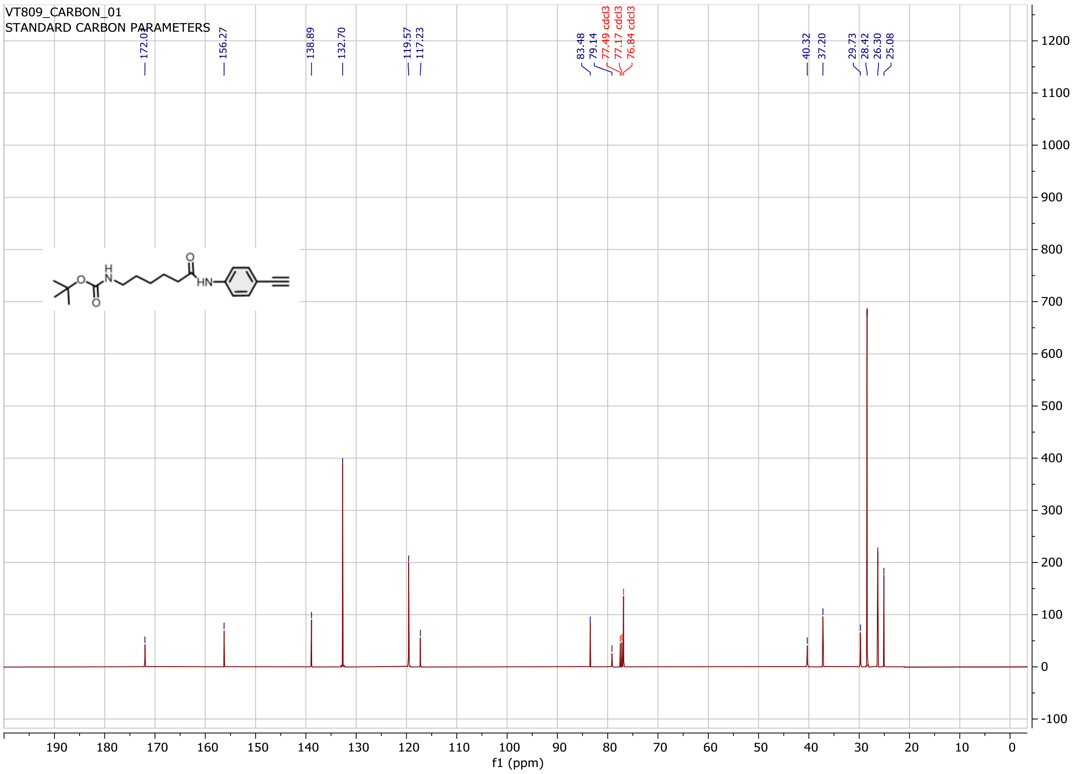

Compound **3a**:

$^{1}H$ NMR spectrum

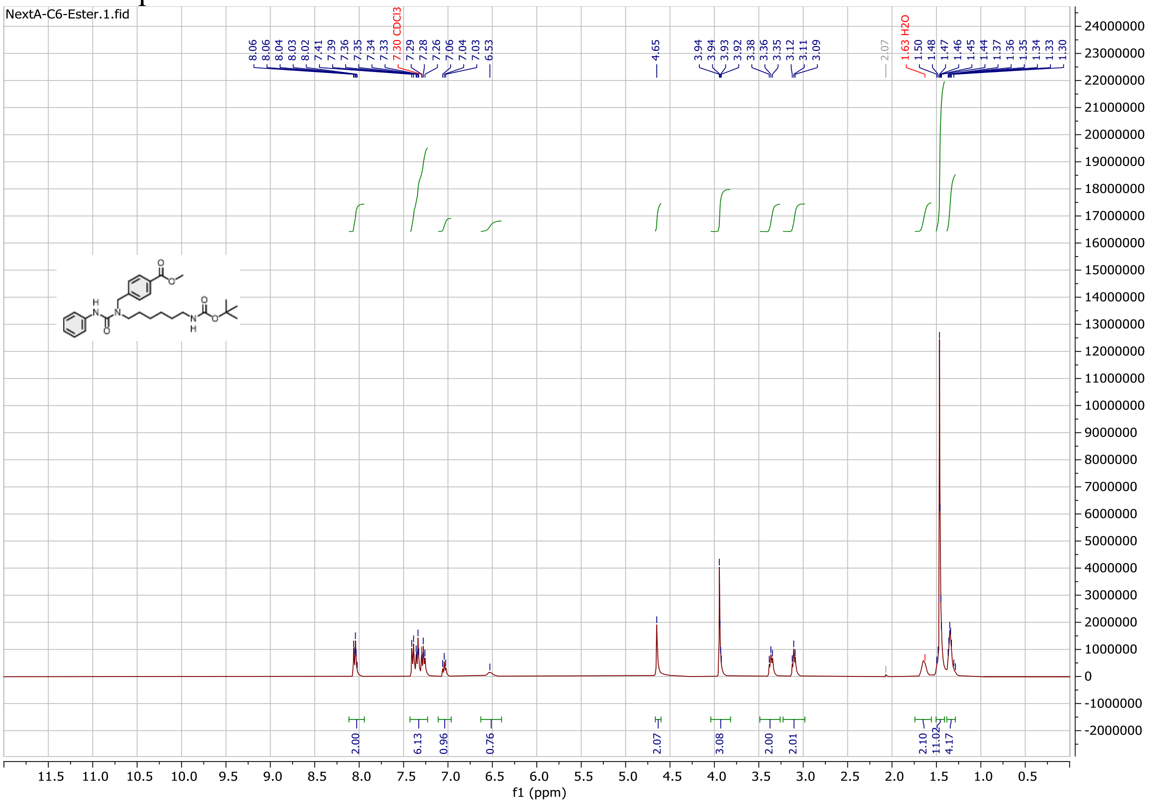

$^{13}C$ NMR spectrum

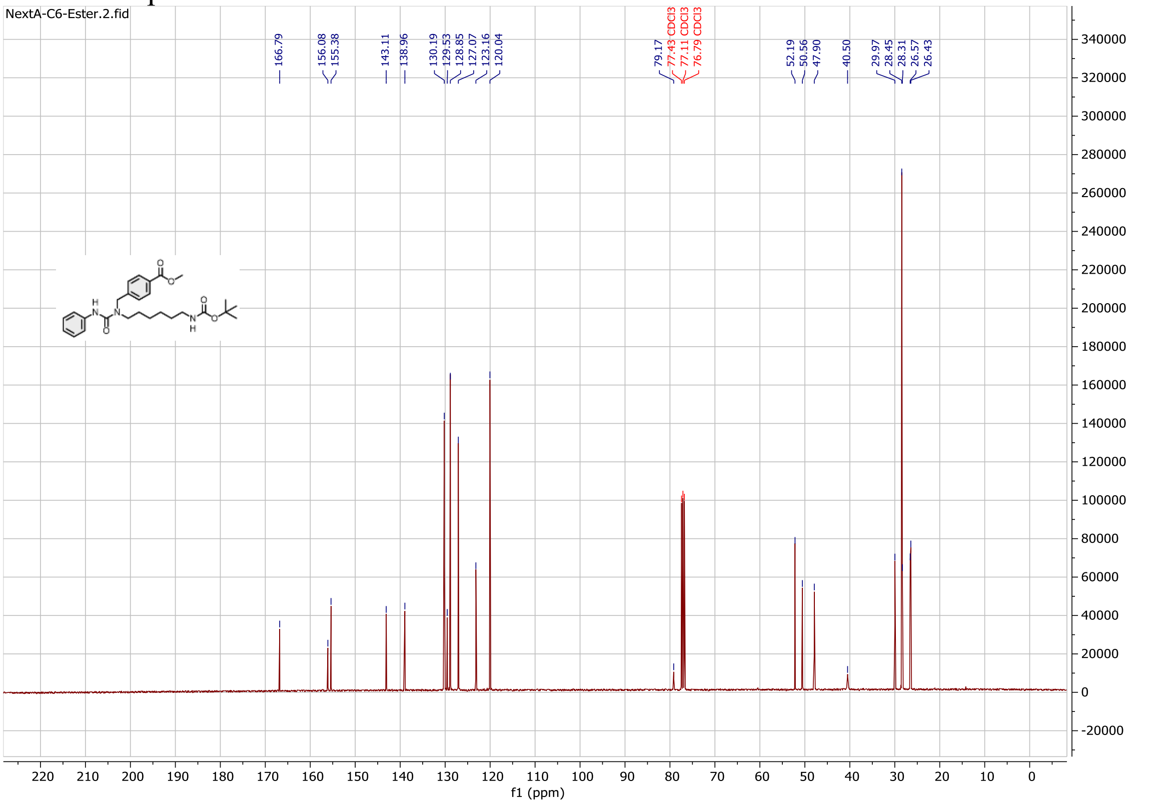

Compound **9a**

$^{1}$H NMR spectrum

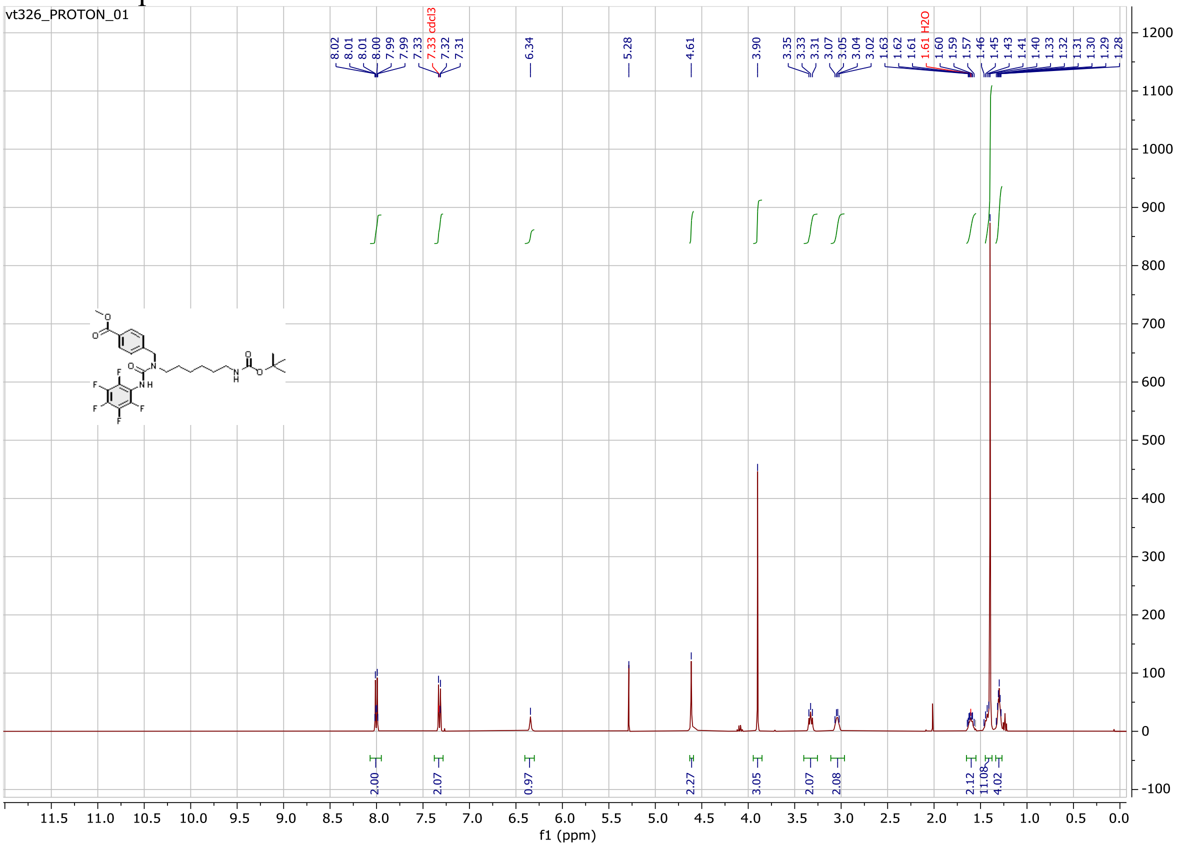

$^{13}$C NMR spectrum

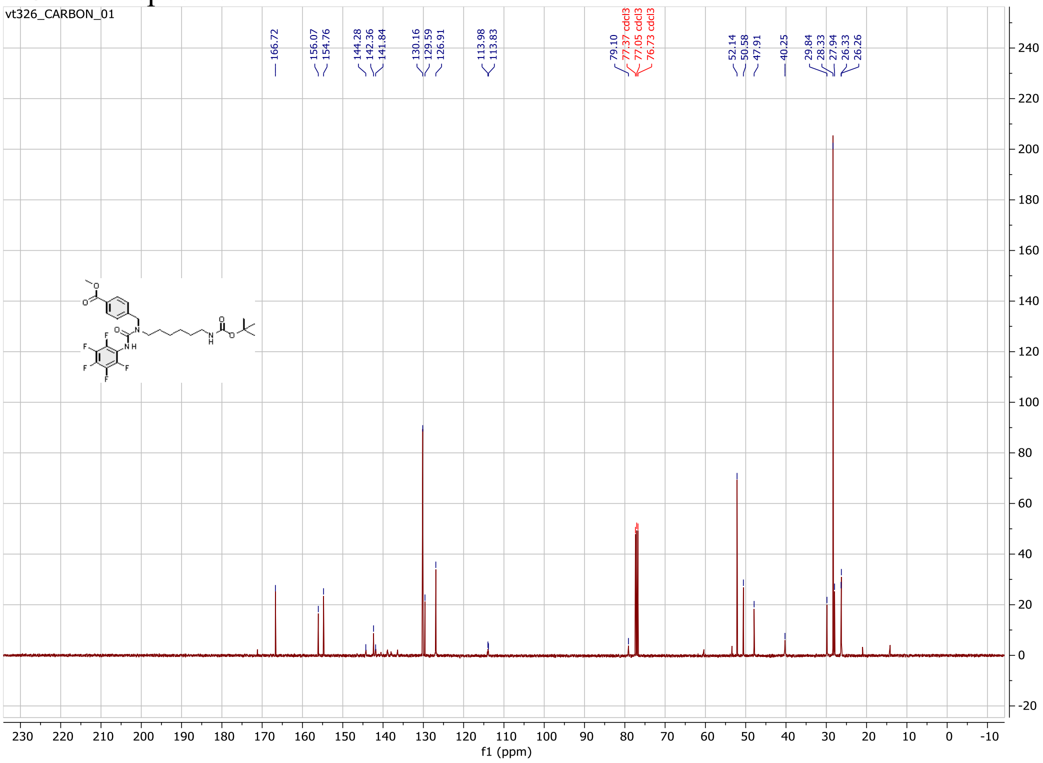

Compound **11a**:

$^{1}$H NMR spectrum

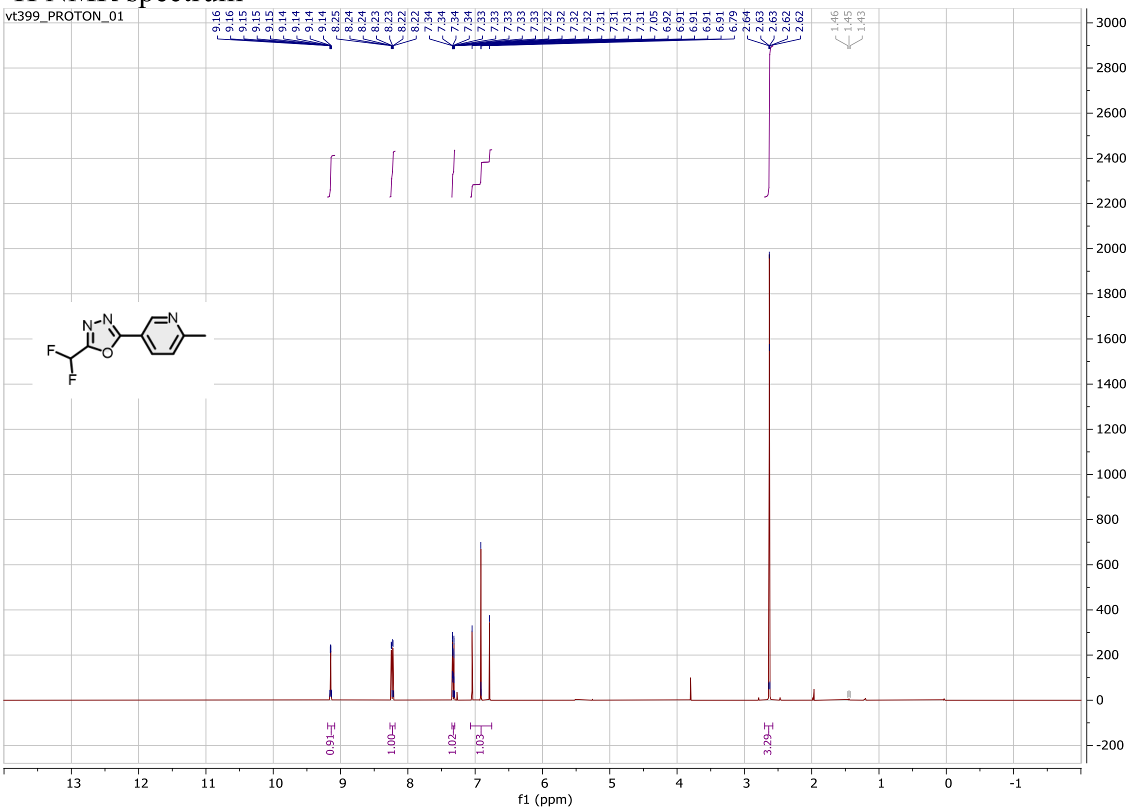

$^{13}$C NMR spectrum

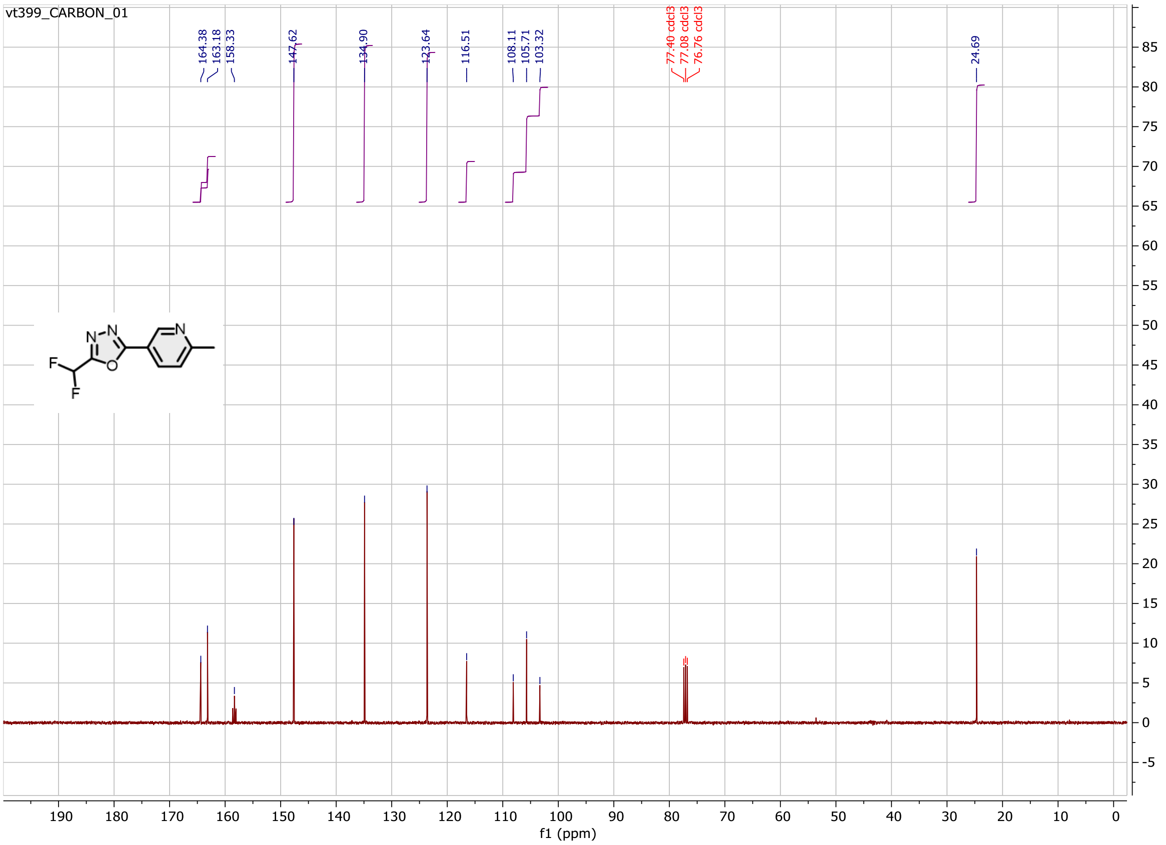

Compound **12a**:

$^{1}$H NMR spectrum

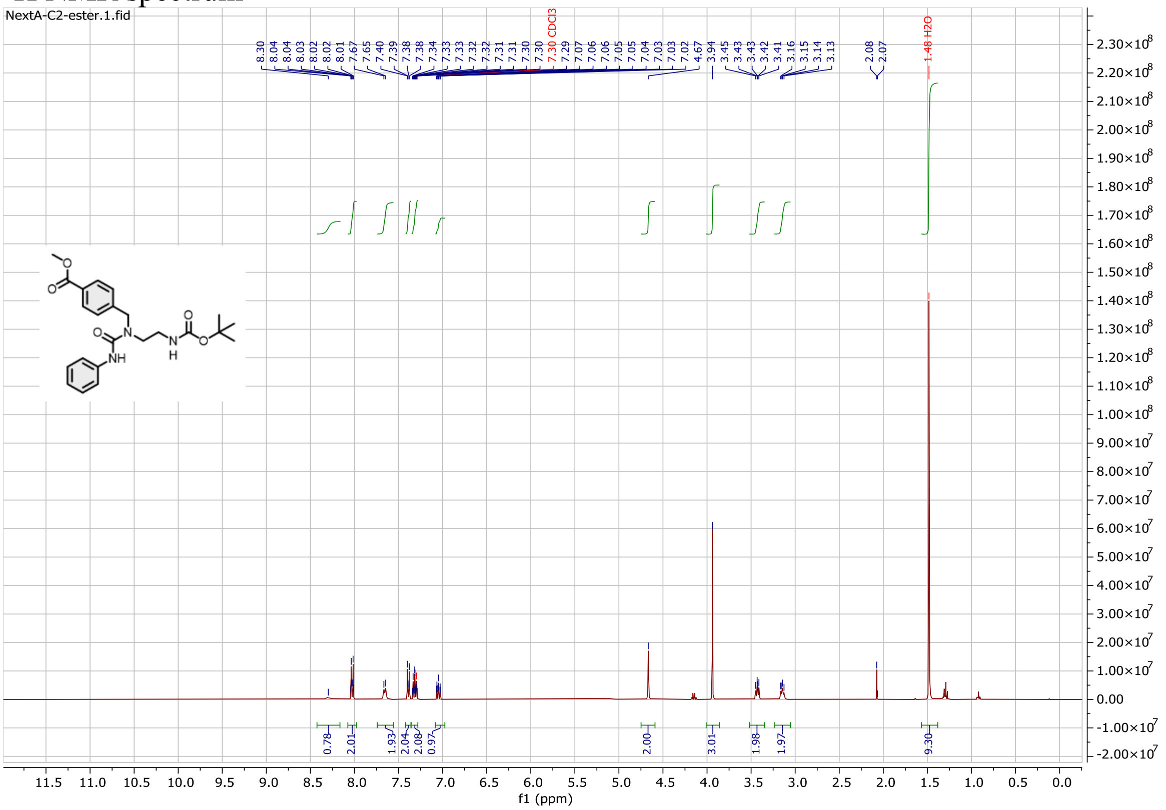

$^{13}$C NMR spectrum

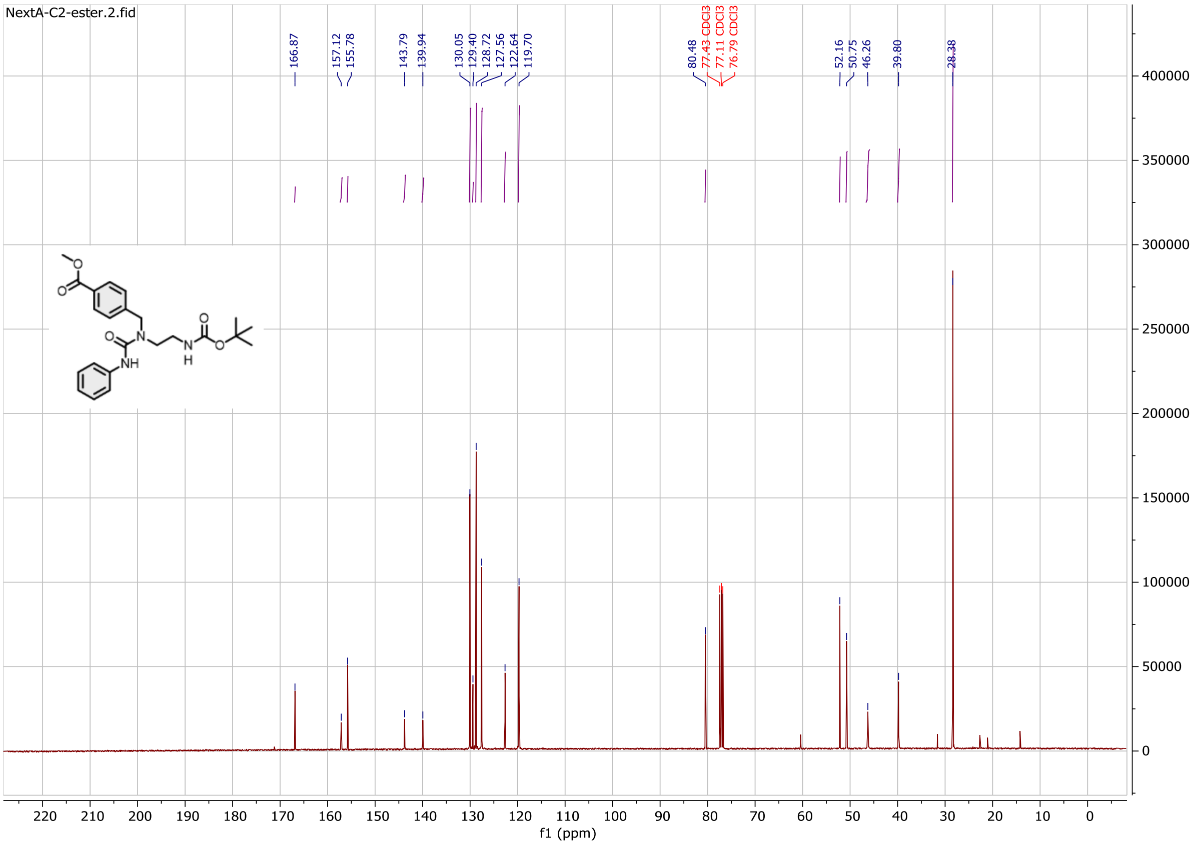

Compound **13a**:

$^{1}$H NMR spectrum

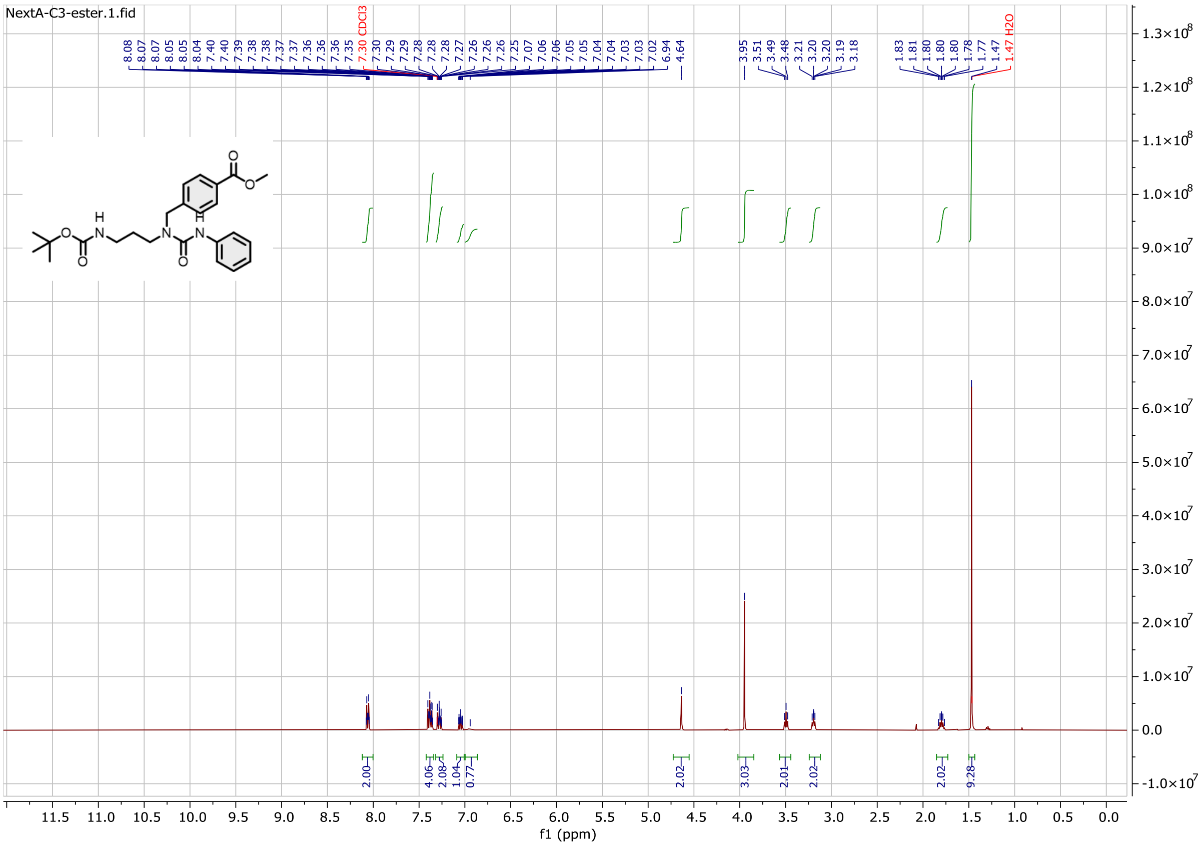

$^{13}$C NMR spectrum

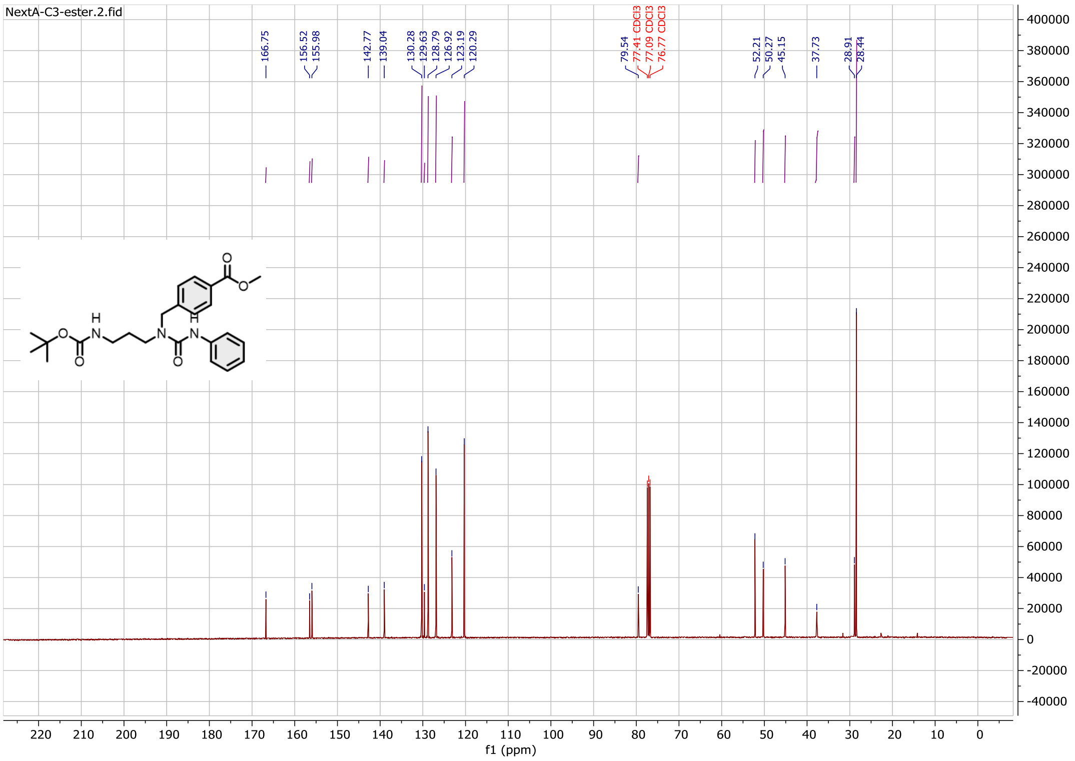

Compound **14a**:

$^{1}$H NMR spectrum

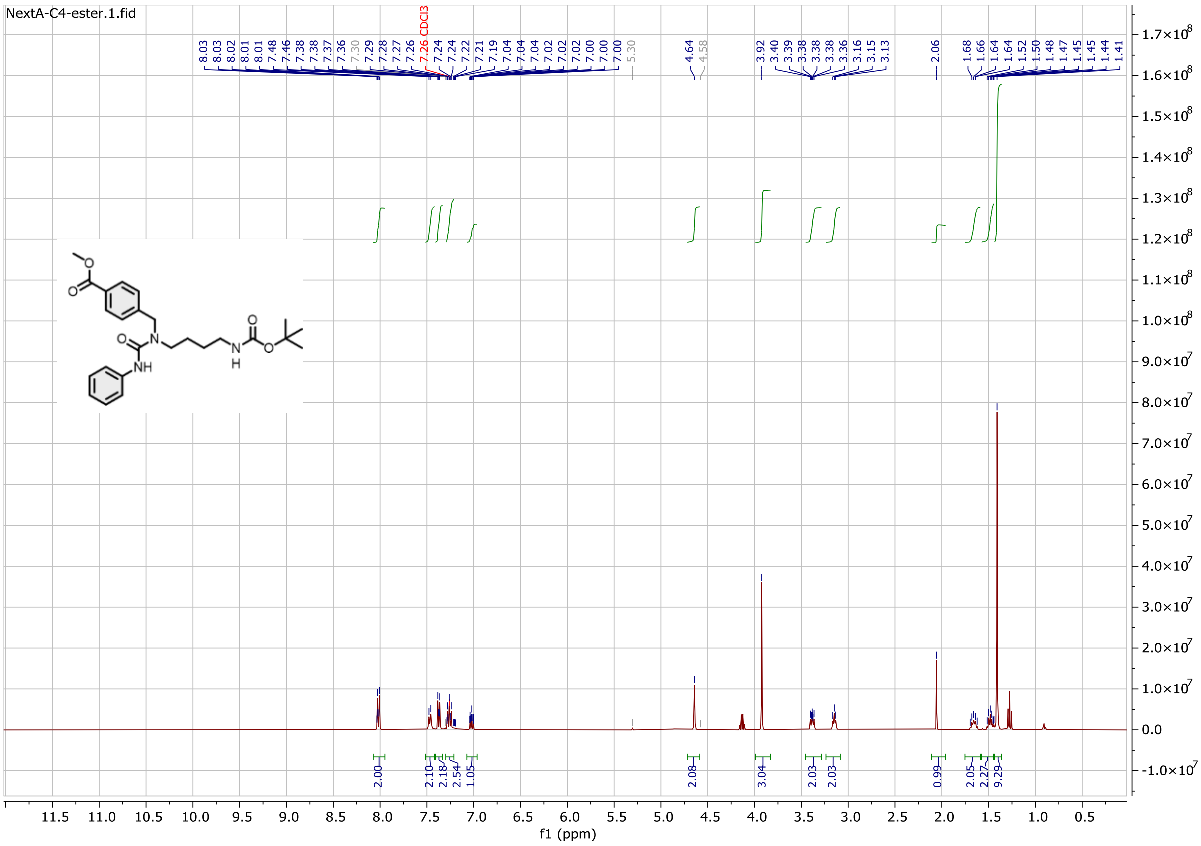

$^{13}$C NMR spectrum

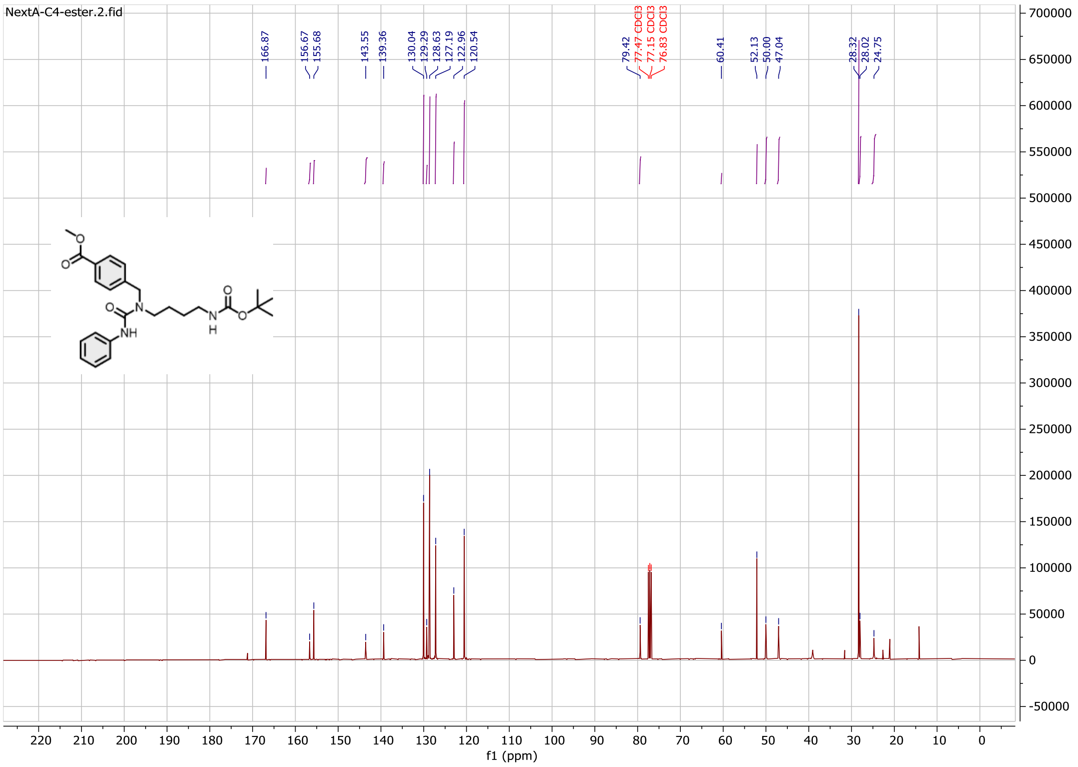

Compound **15a**:

$^{1}$H NMR spectrum

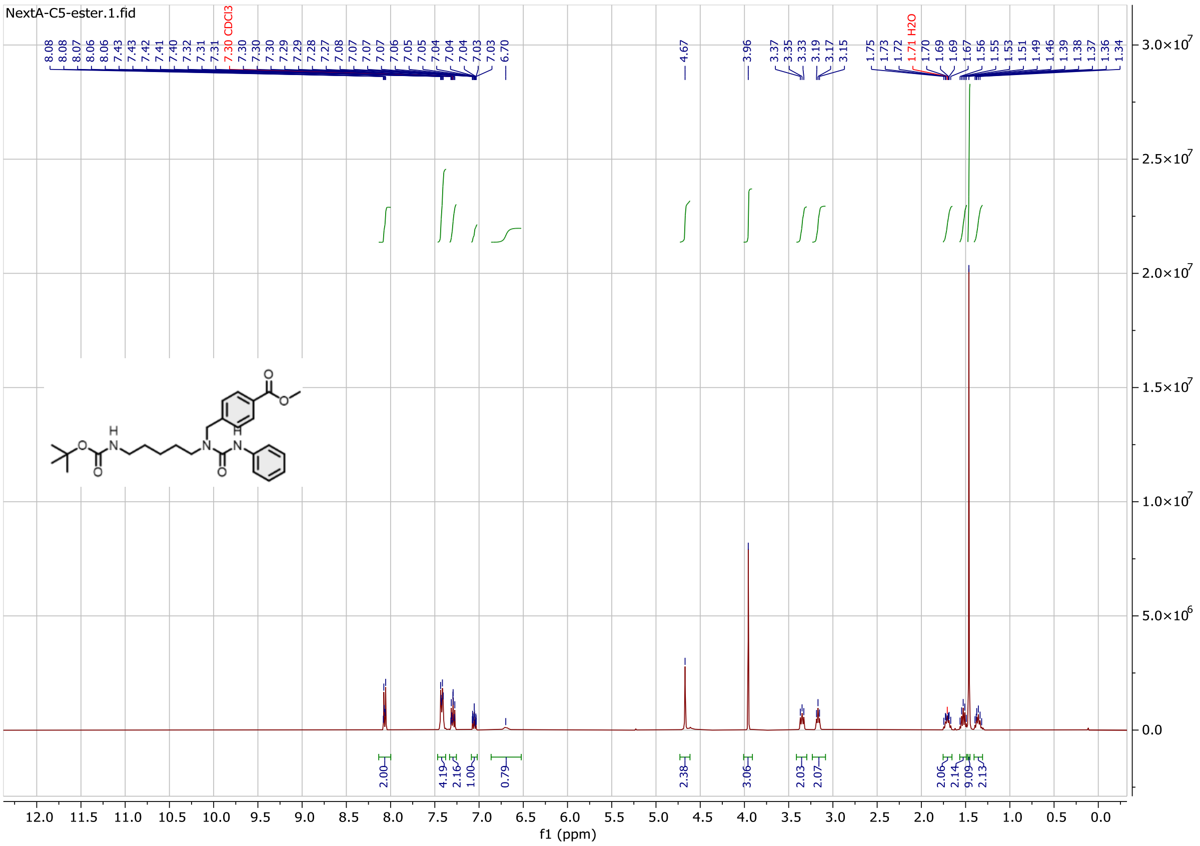

$^{13}$C NMR spectrum

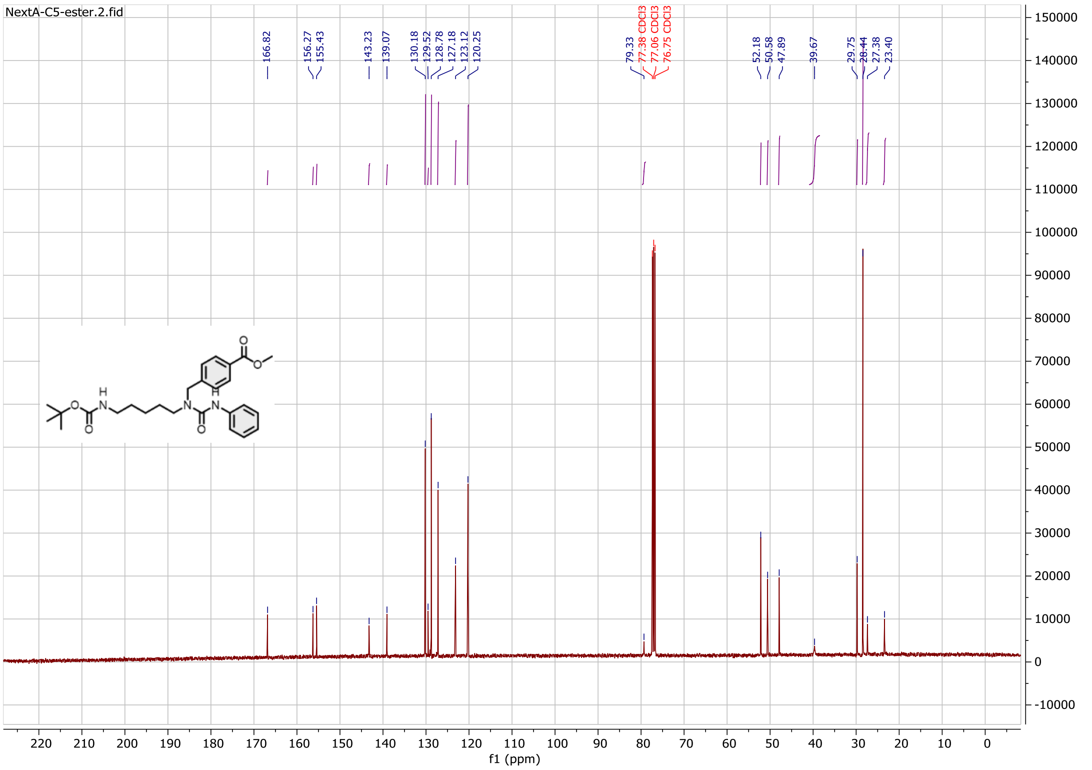

Compound **16a**:

$^{1}$H NMR spectrum

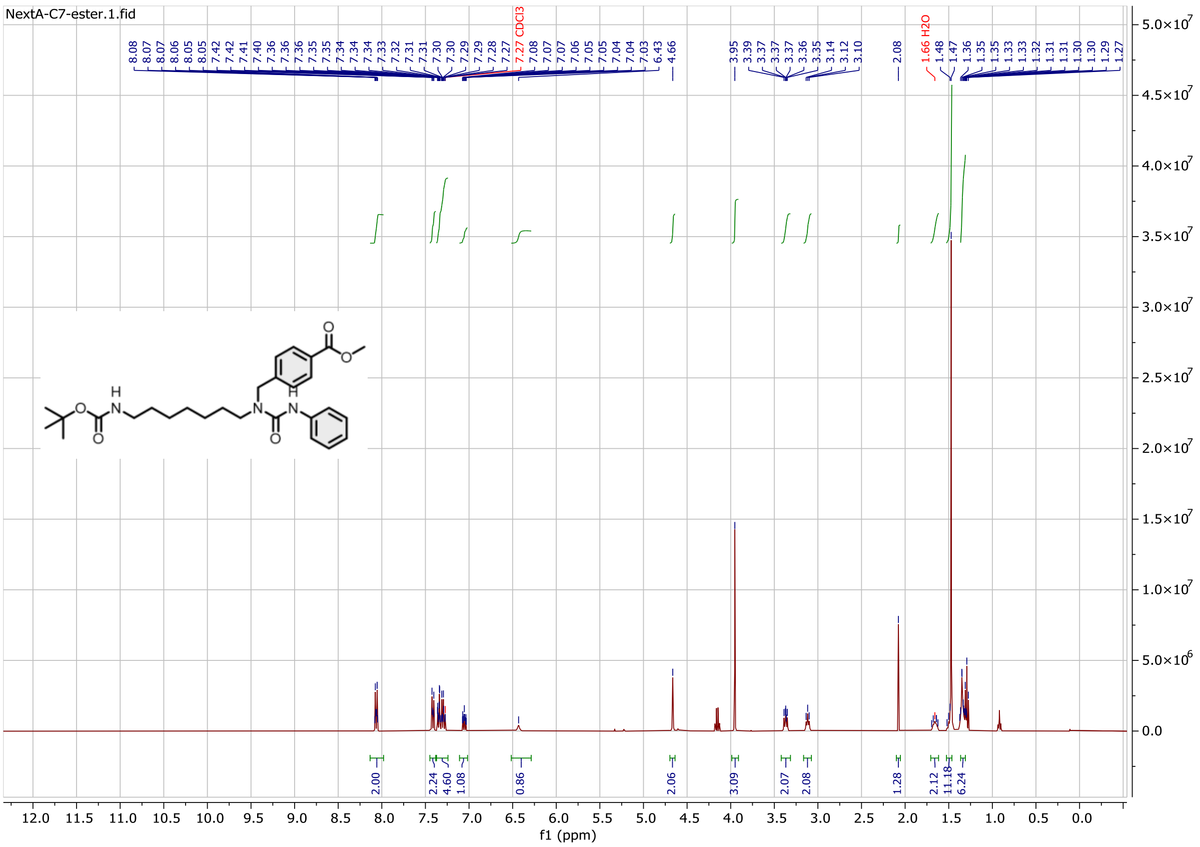

$^{13}$C NMR spectrum

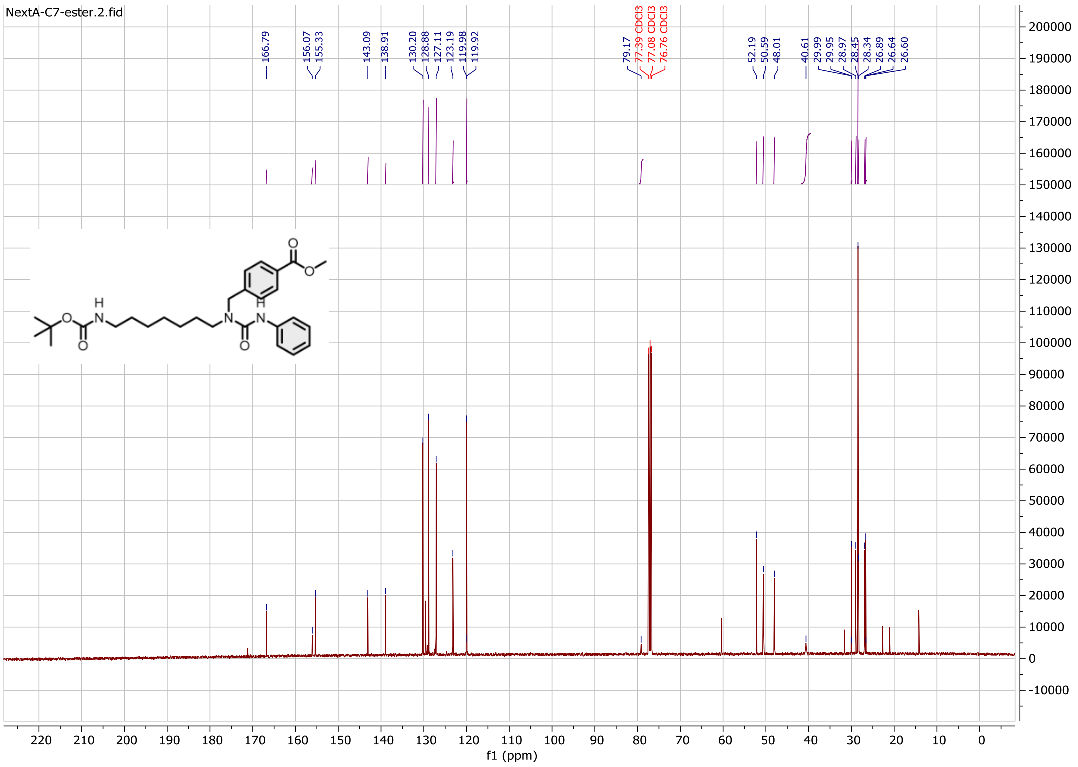

Compound **17a**:

$^{1}$H NMR spectrum

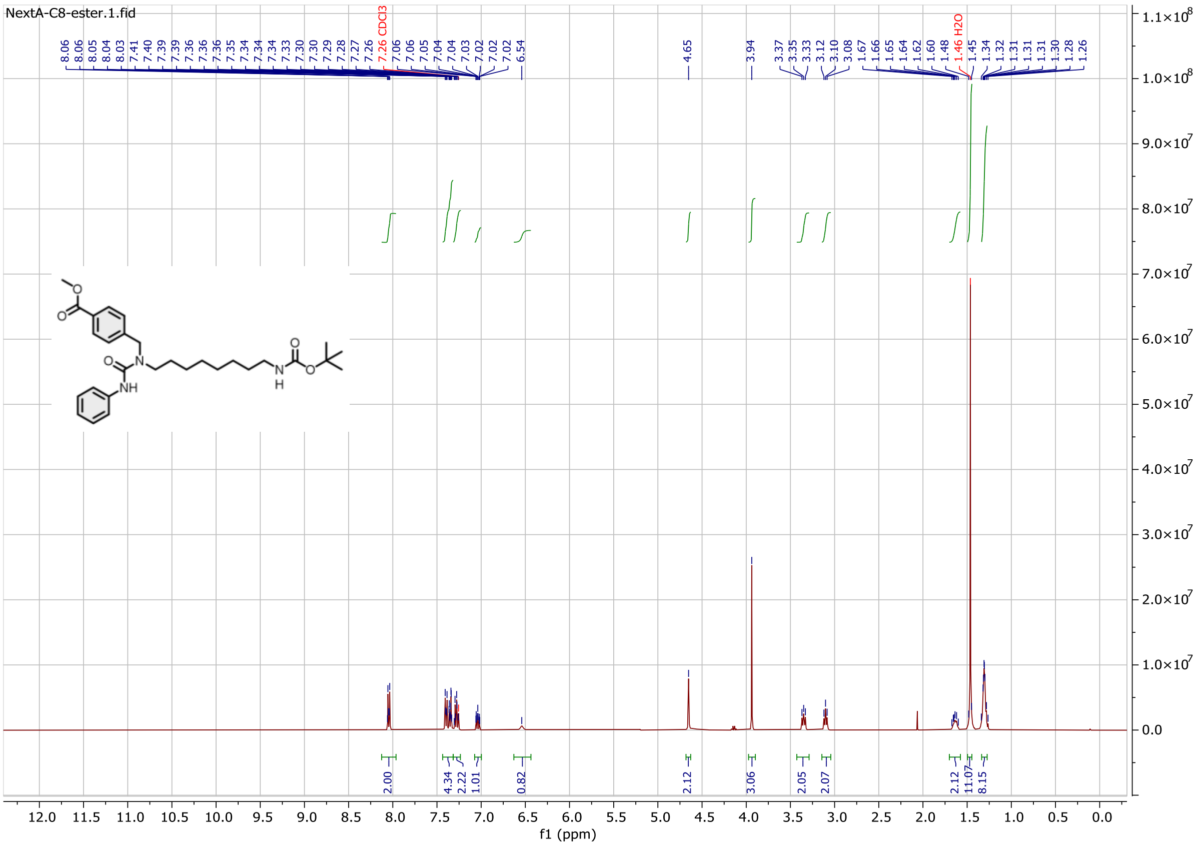

$^{13}$C NMR spectrum

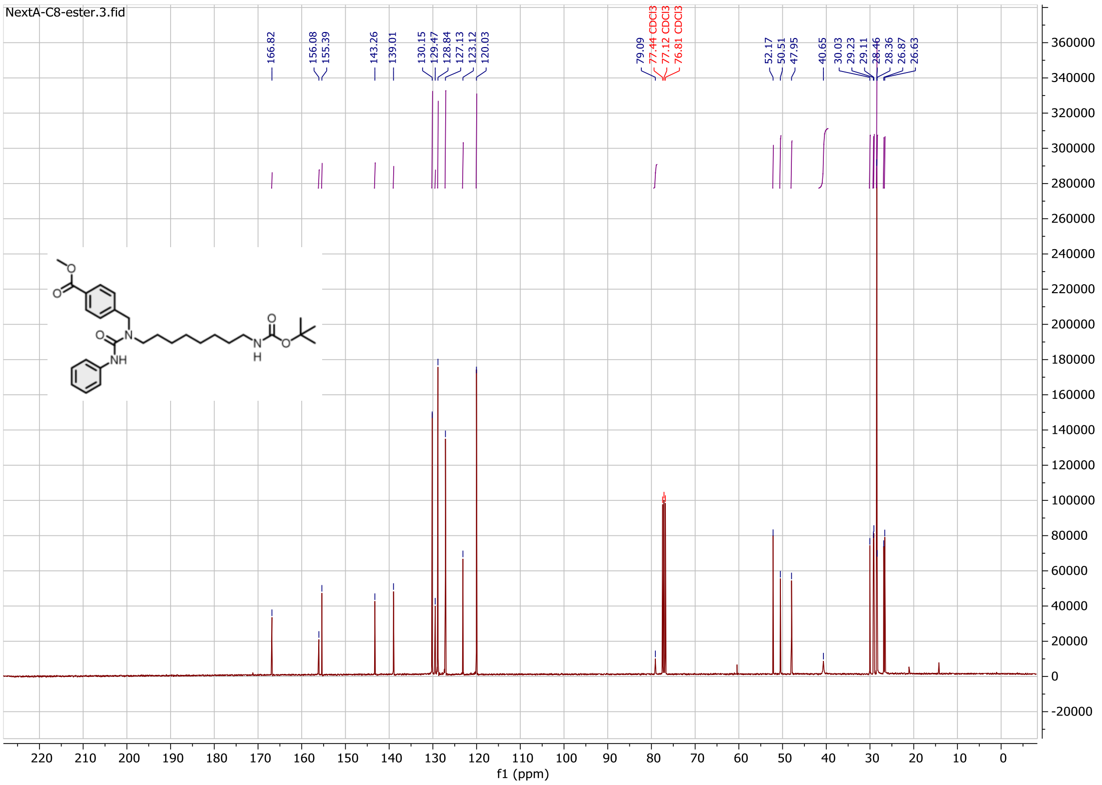

Compound **18a**:

$^{1}$H NMR spectrum

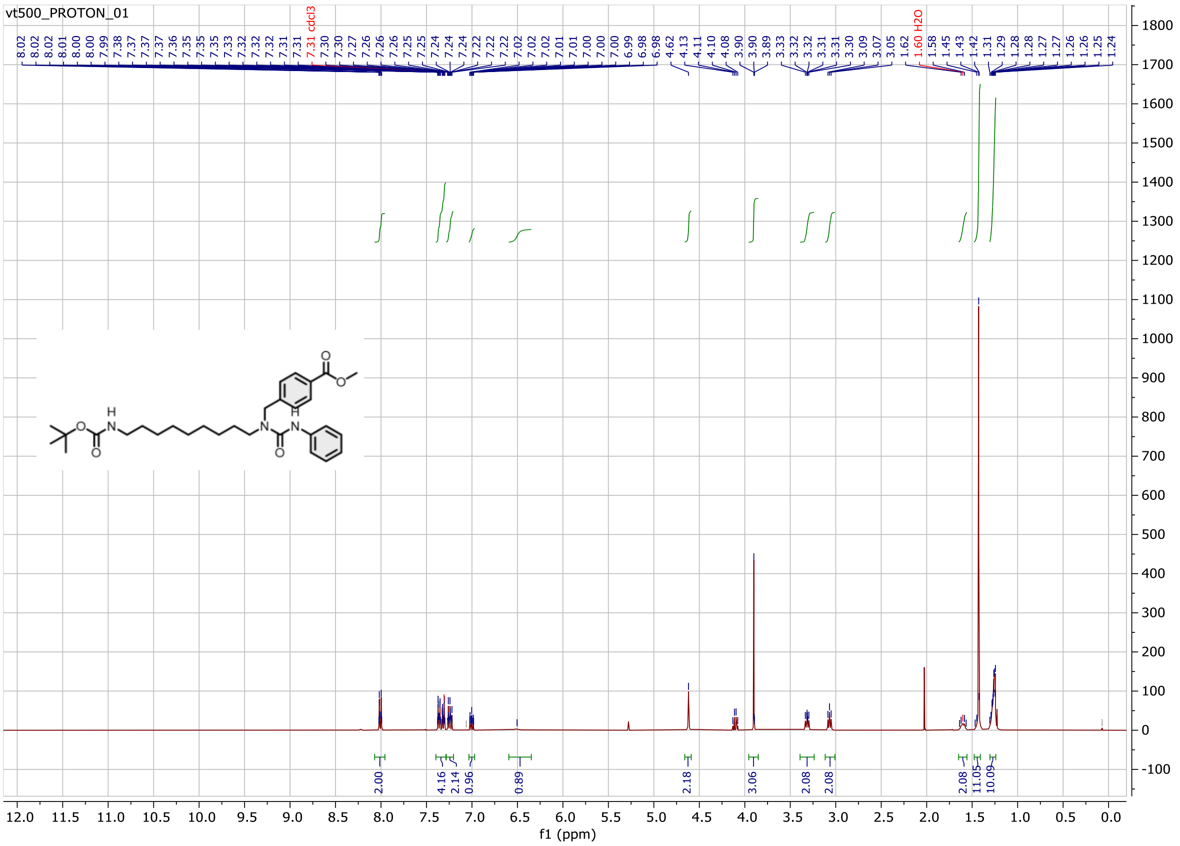

$^{13}$C NMR spectrum

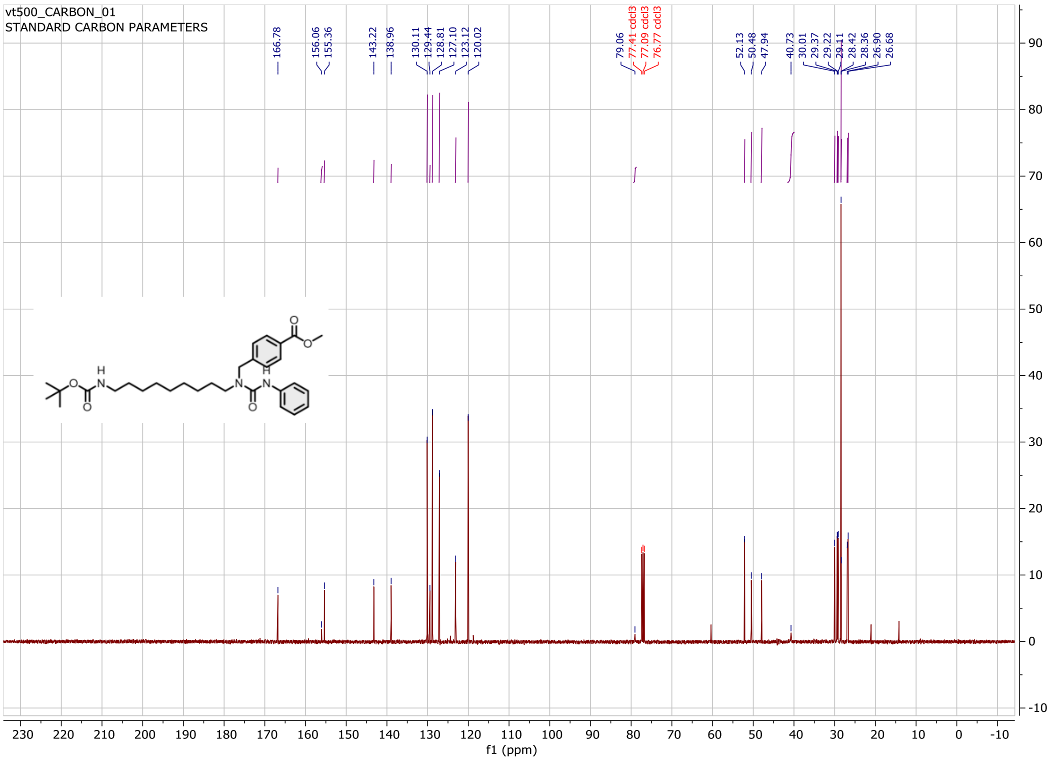

Compound **19a**:

$^{1}$H NMR spectrum

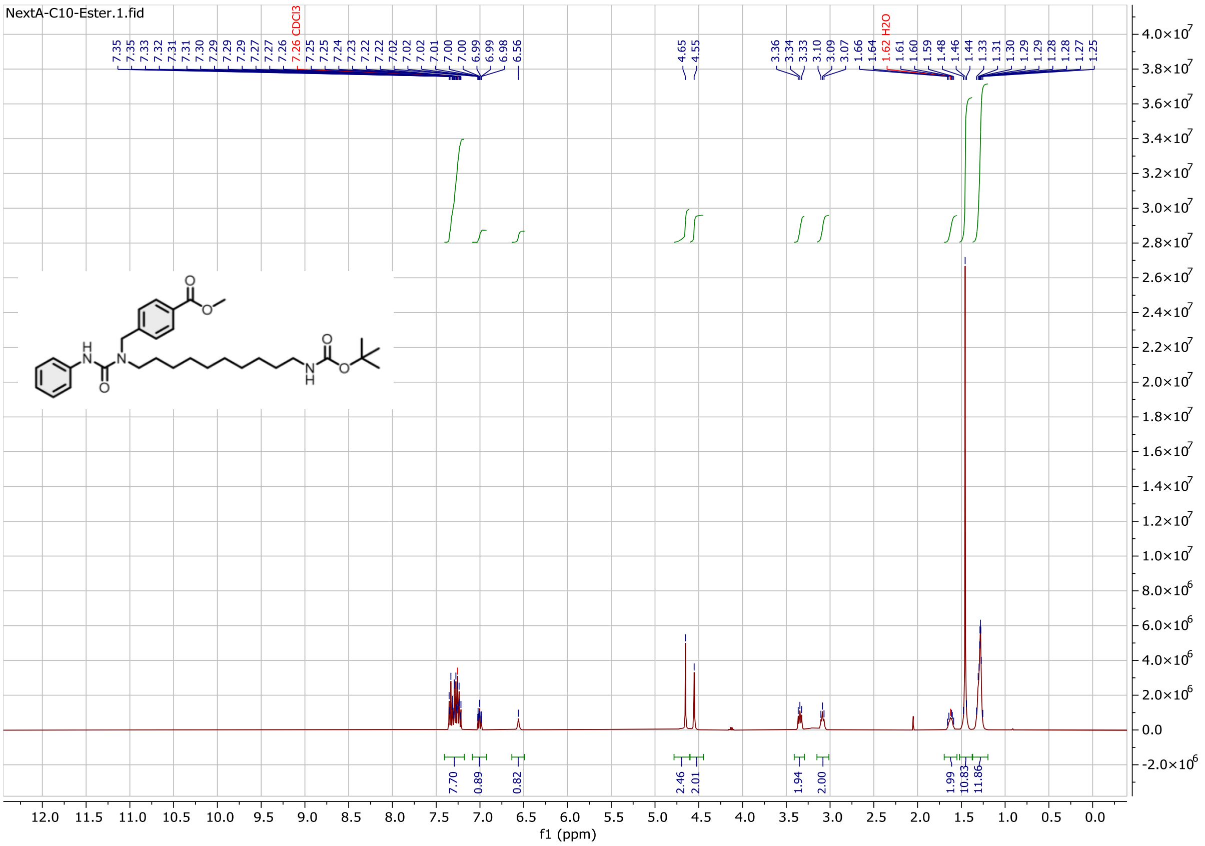

$^{13}$C NMR spectrum

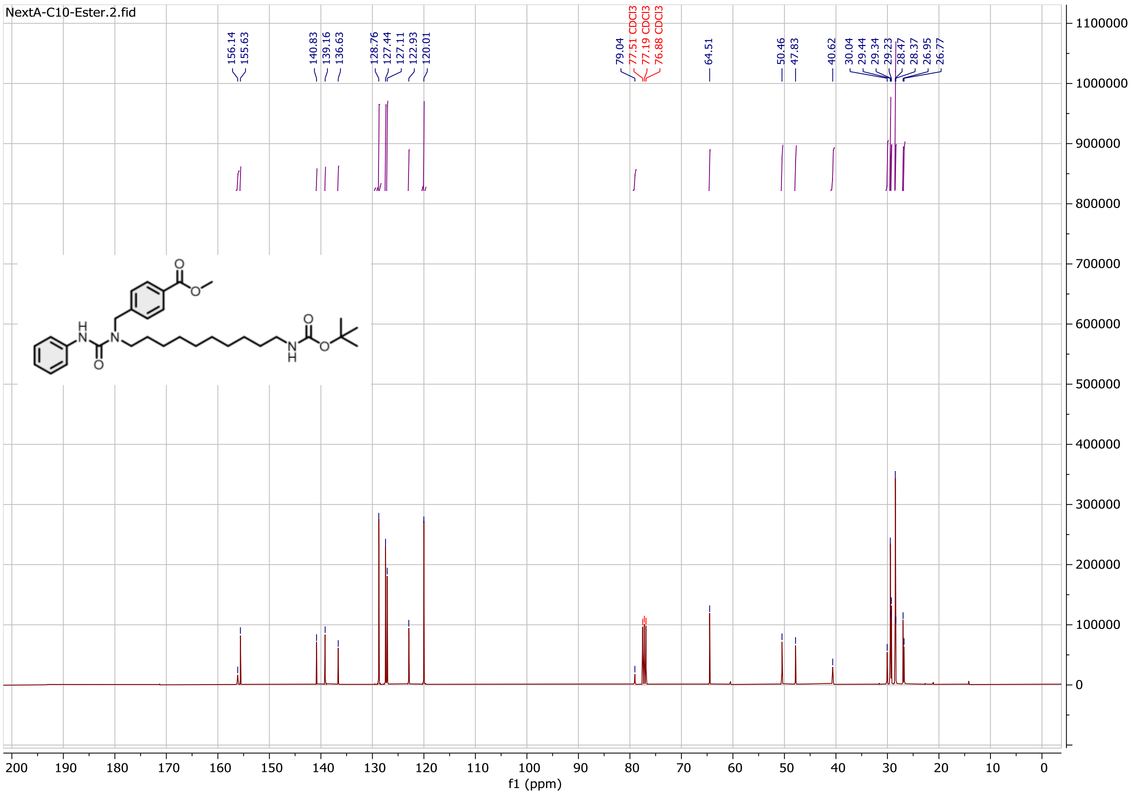

Compound **20a**:

$^{1}$H NMR spectrum

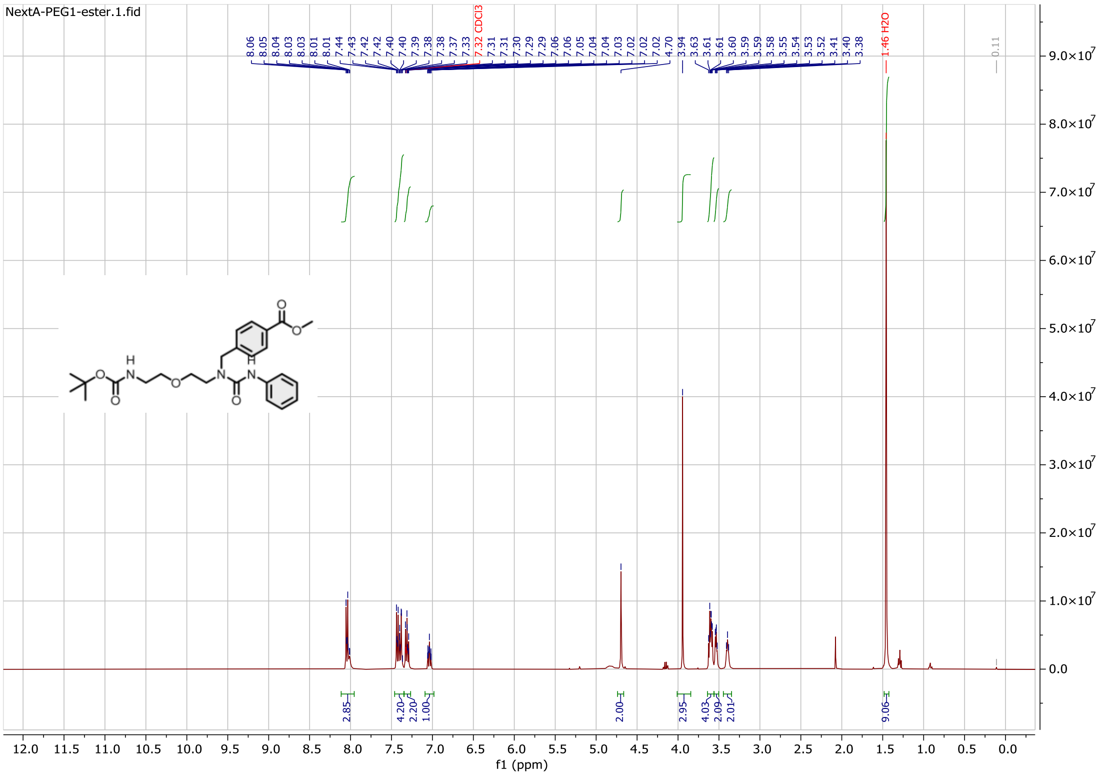

$^{13}$C NMR spectrum

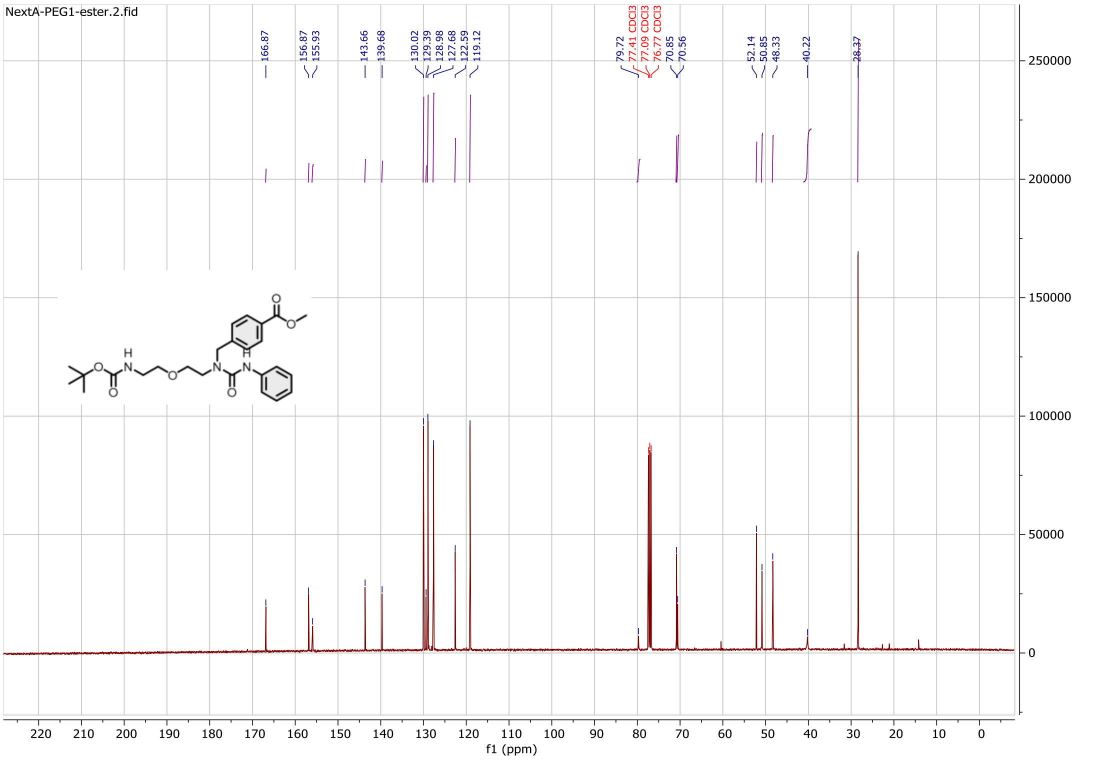

Compound **21a**:

$^{1}$H NMR spectrum

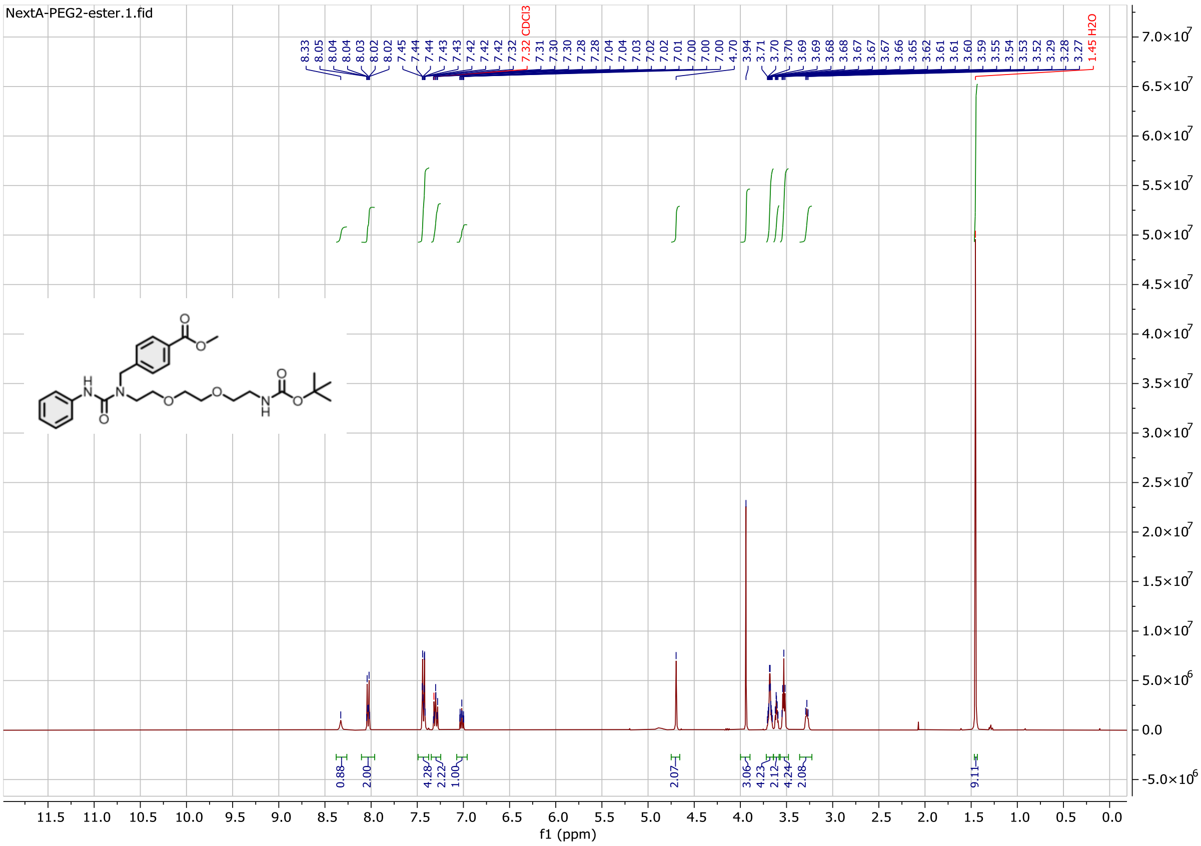

$^{13}$C NMR spectrum

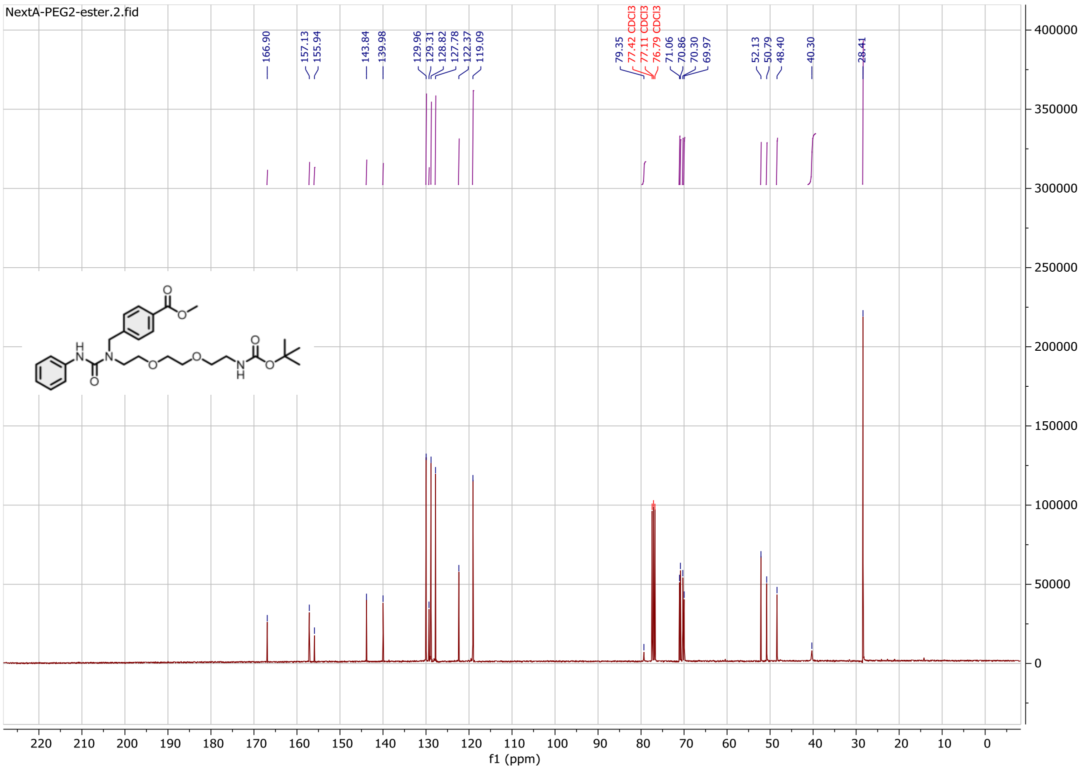

Compound **22a**:

$^{1}$H NMR spectrum

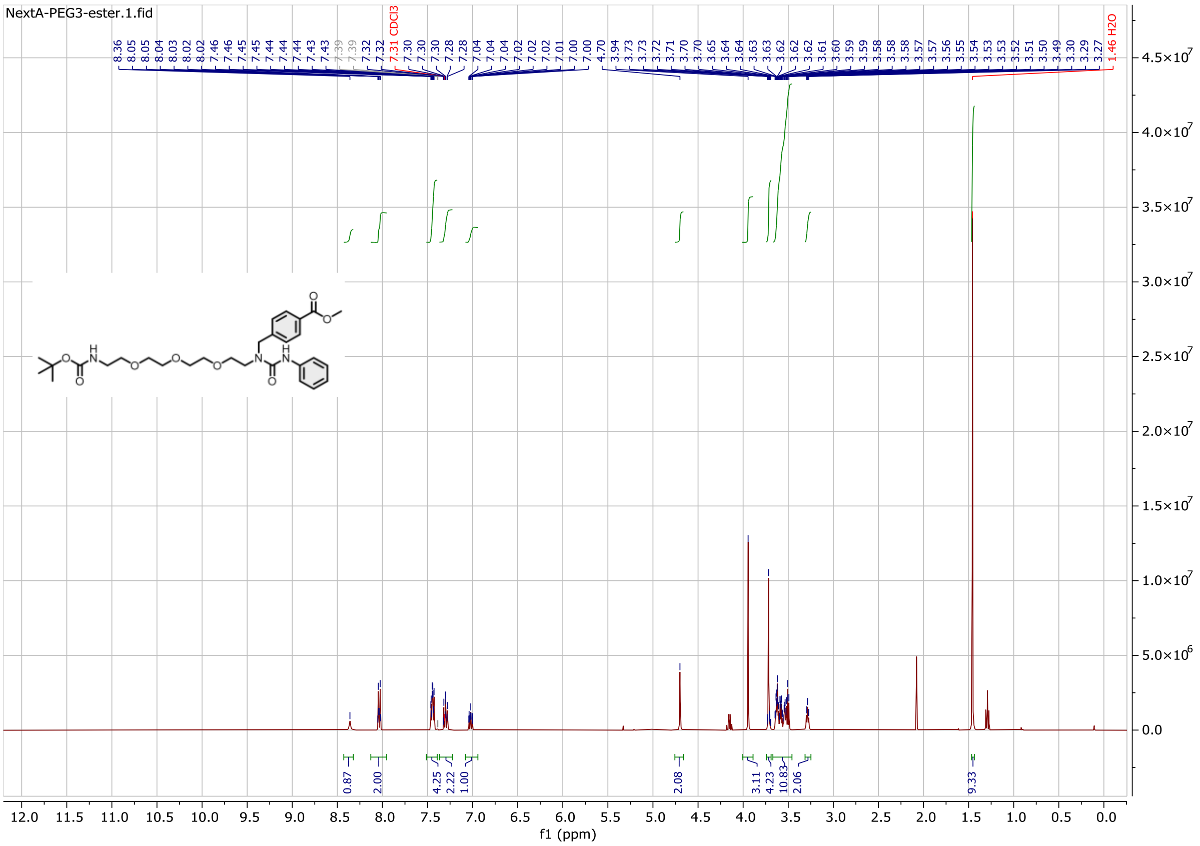

$^{13}$C NMR spectrum

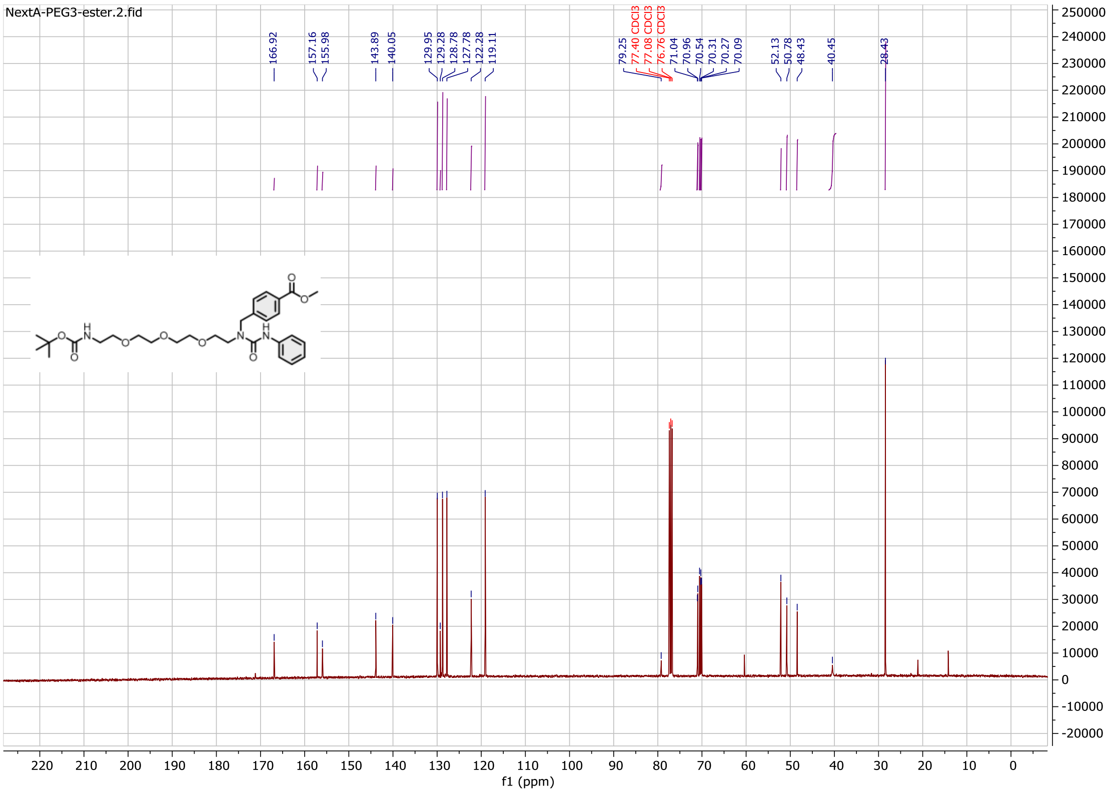

Compound **23a**:

$^{1}$H NMR spectrum

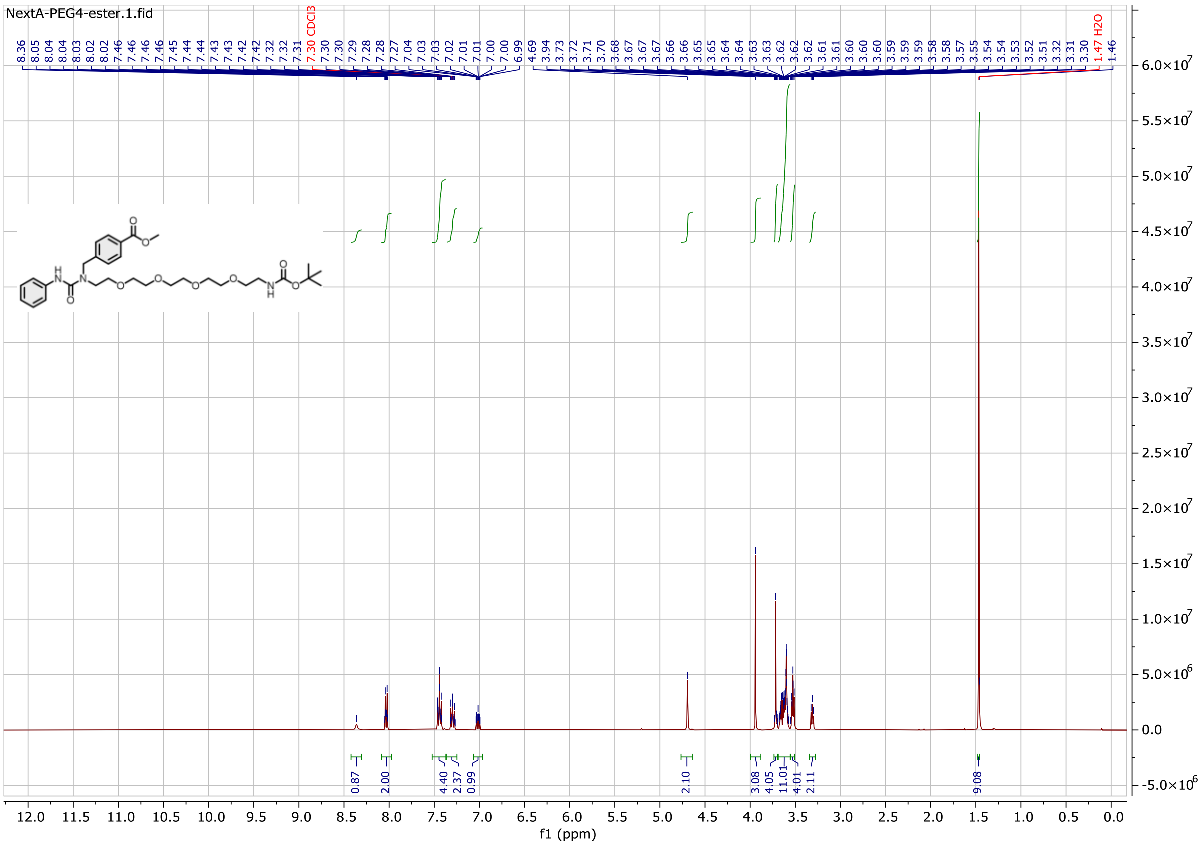

$^{13}$C NMR spectrum

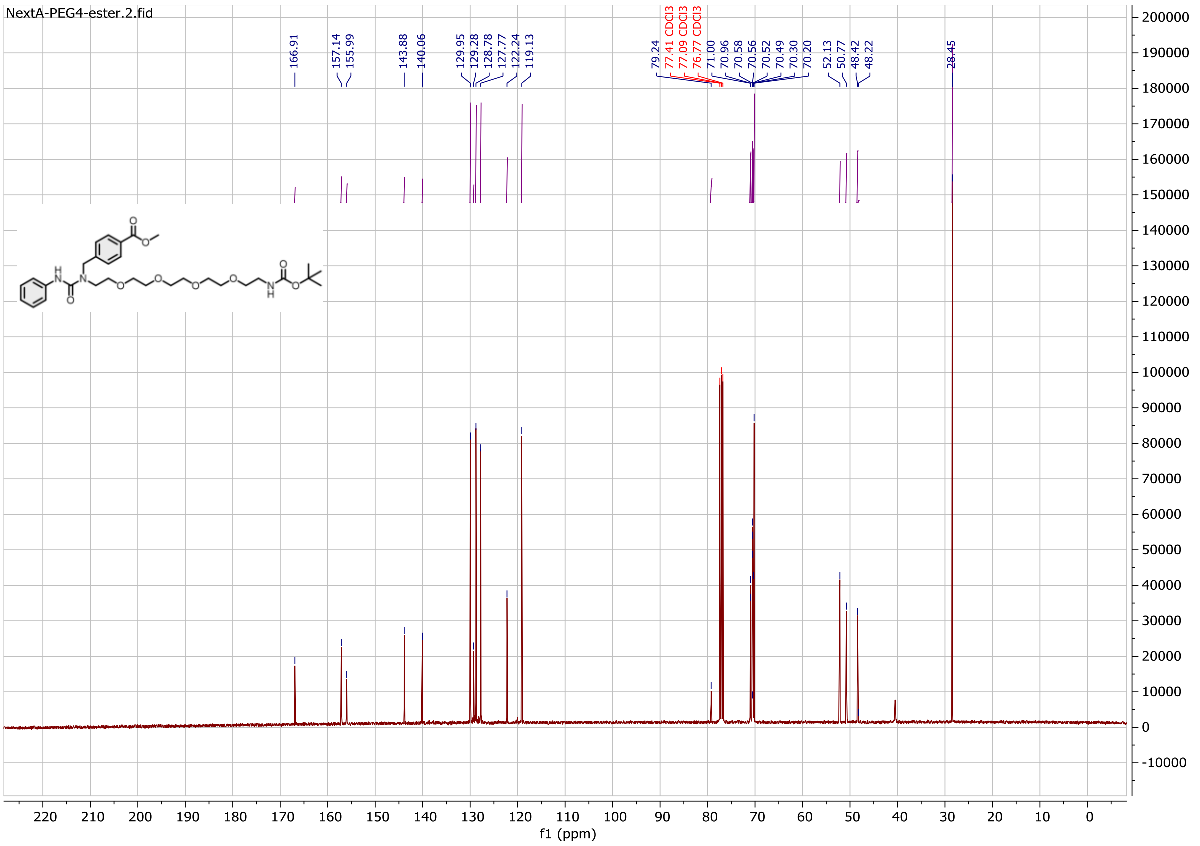

Compound **1b**:

$^{1}$H NMR spectrum

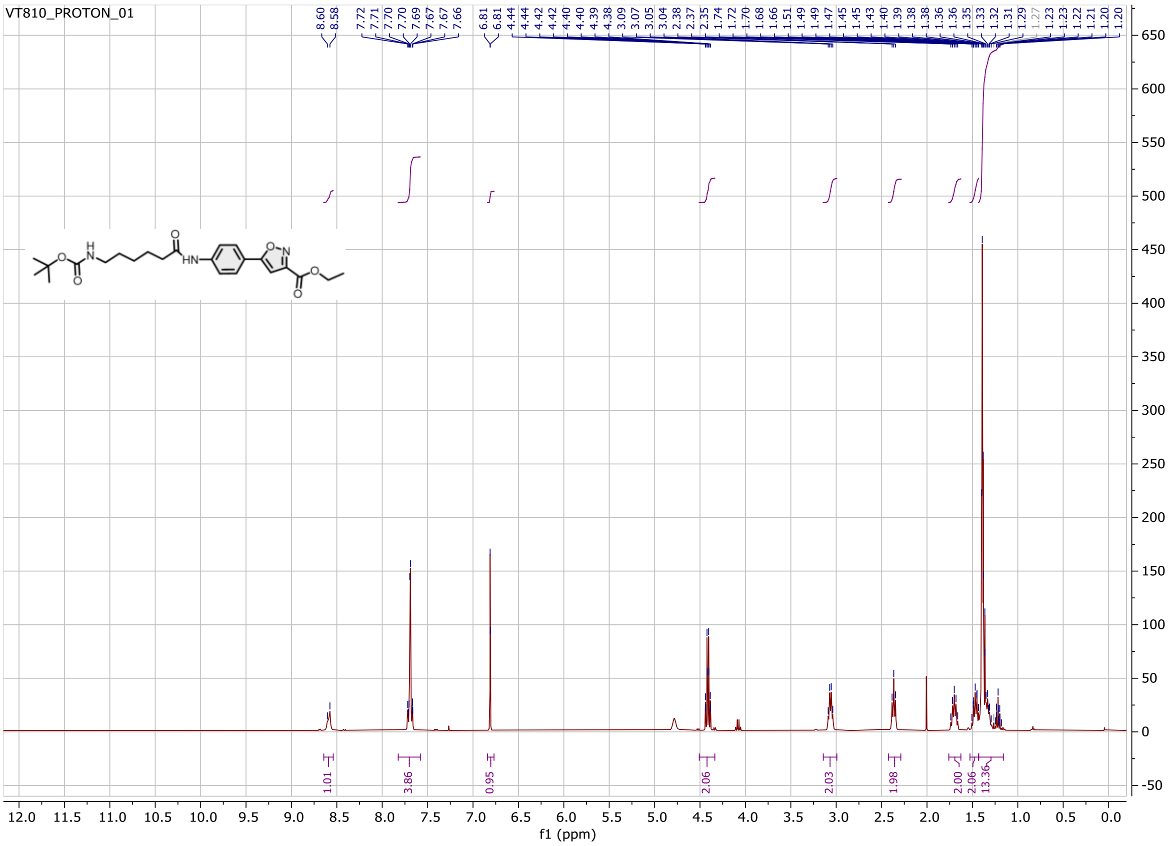

$^{13}$C NMR spectrum

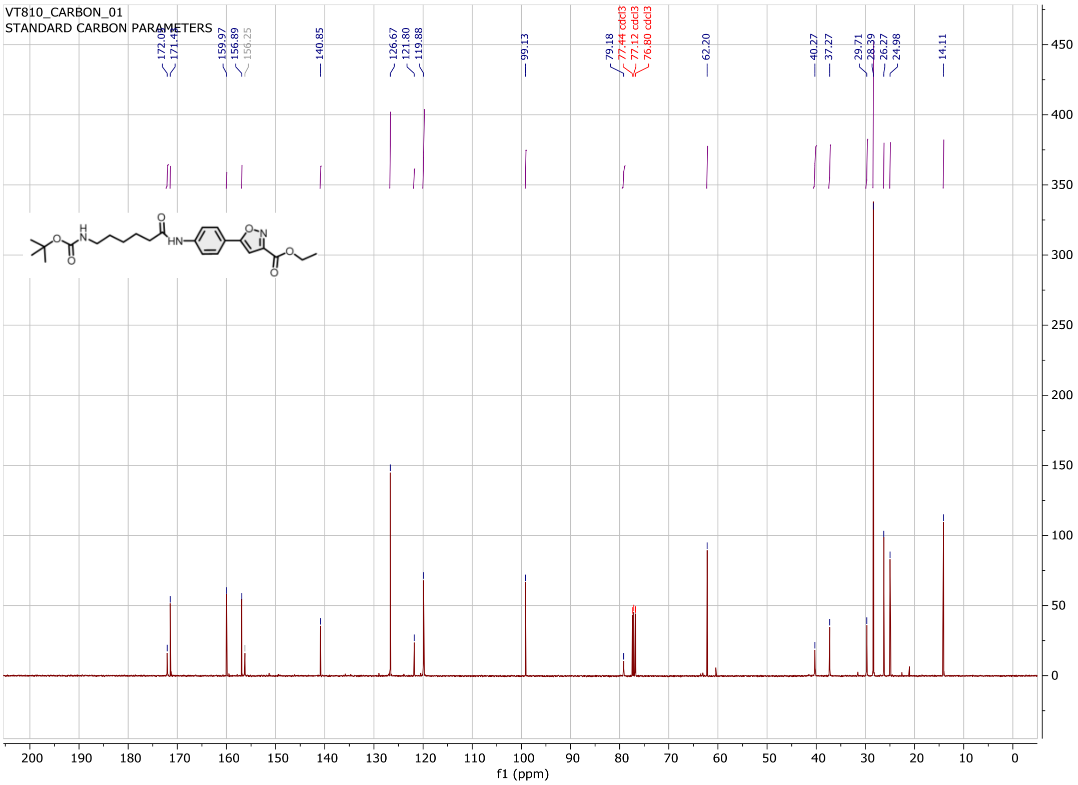

Compound **3b**:

$^{1}$H NMR spectrum

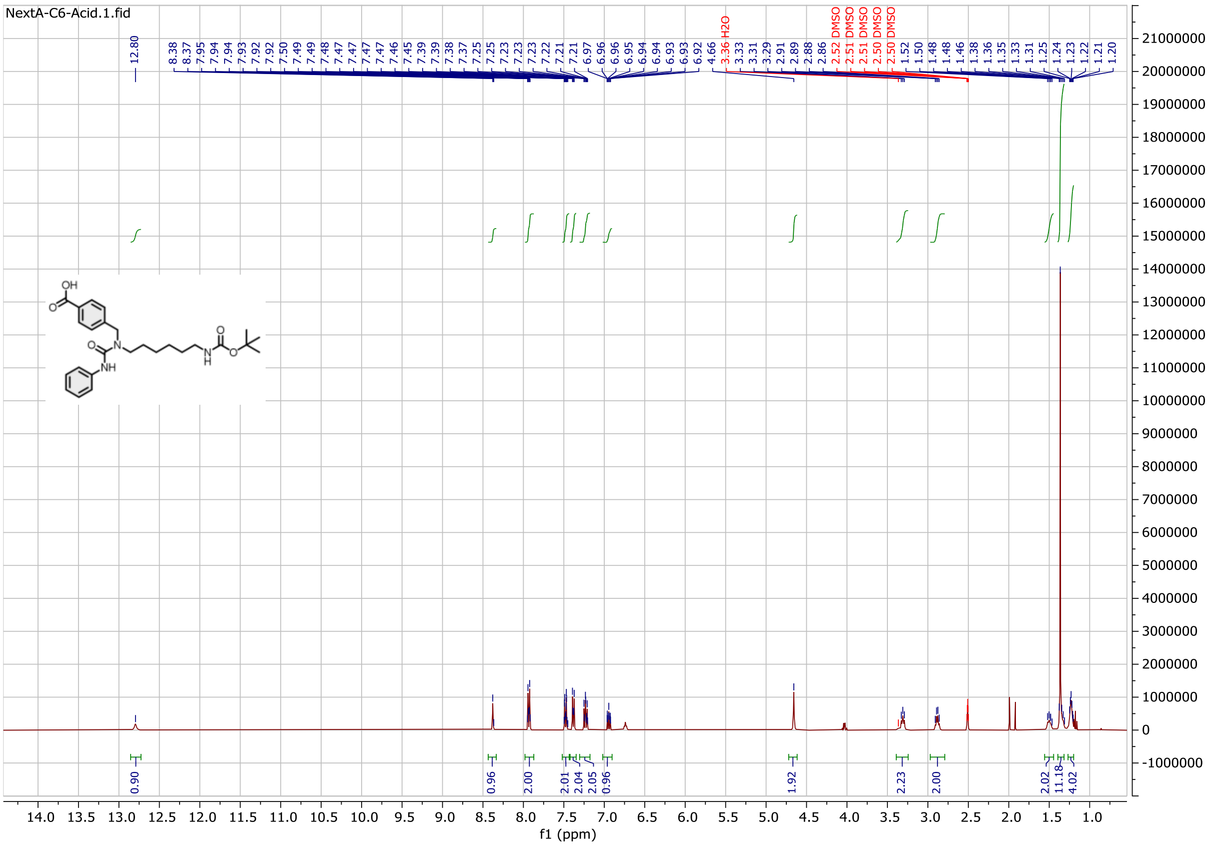

$^{13}$C NMR spectrum

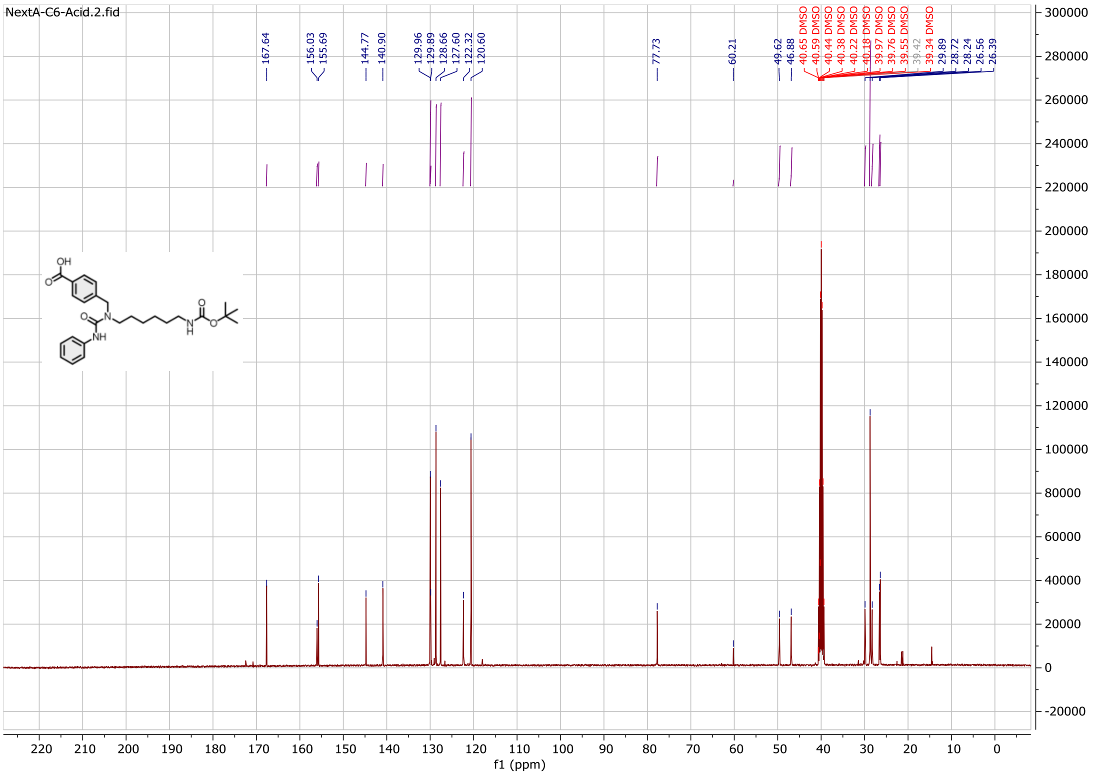

Compound **9b**:

$^{1}$H NMR spectrum

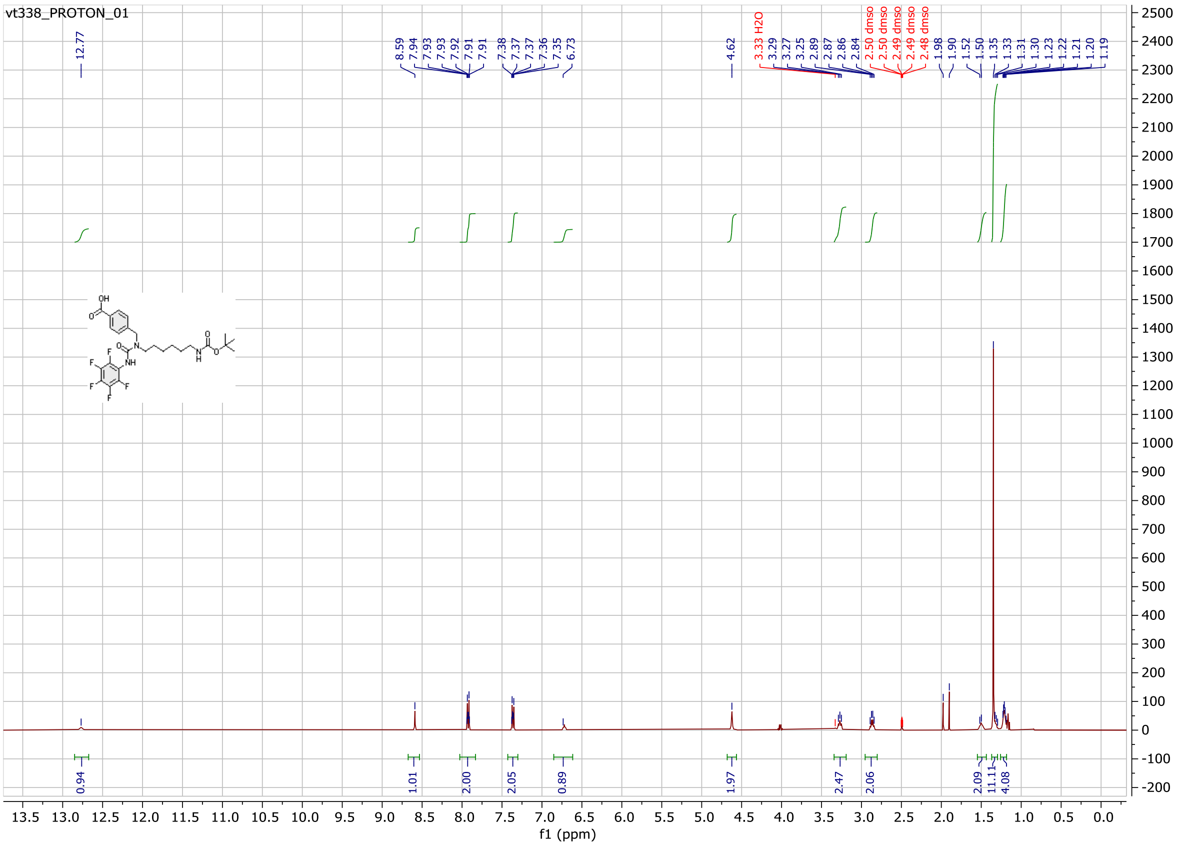

$^{13}$C NMR spectrum

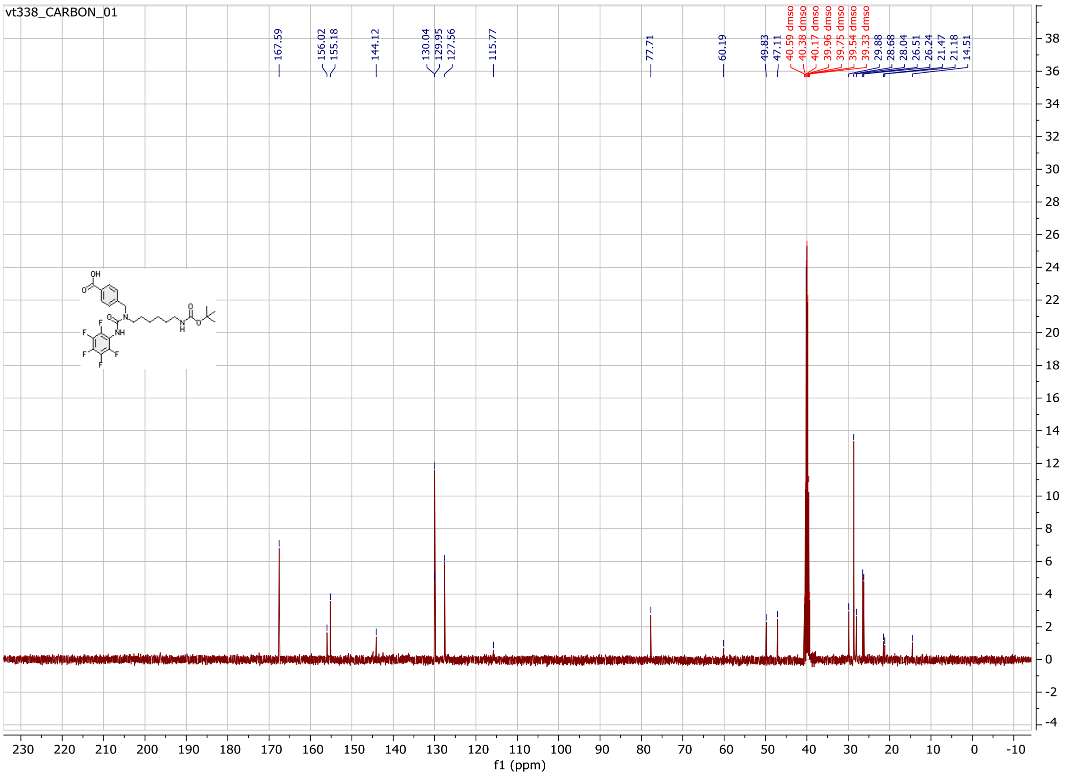

Compound **10b**:

$^{1}$H NMR spectrum

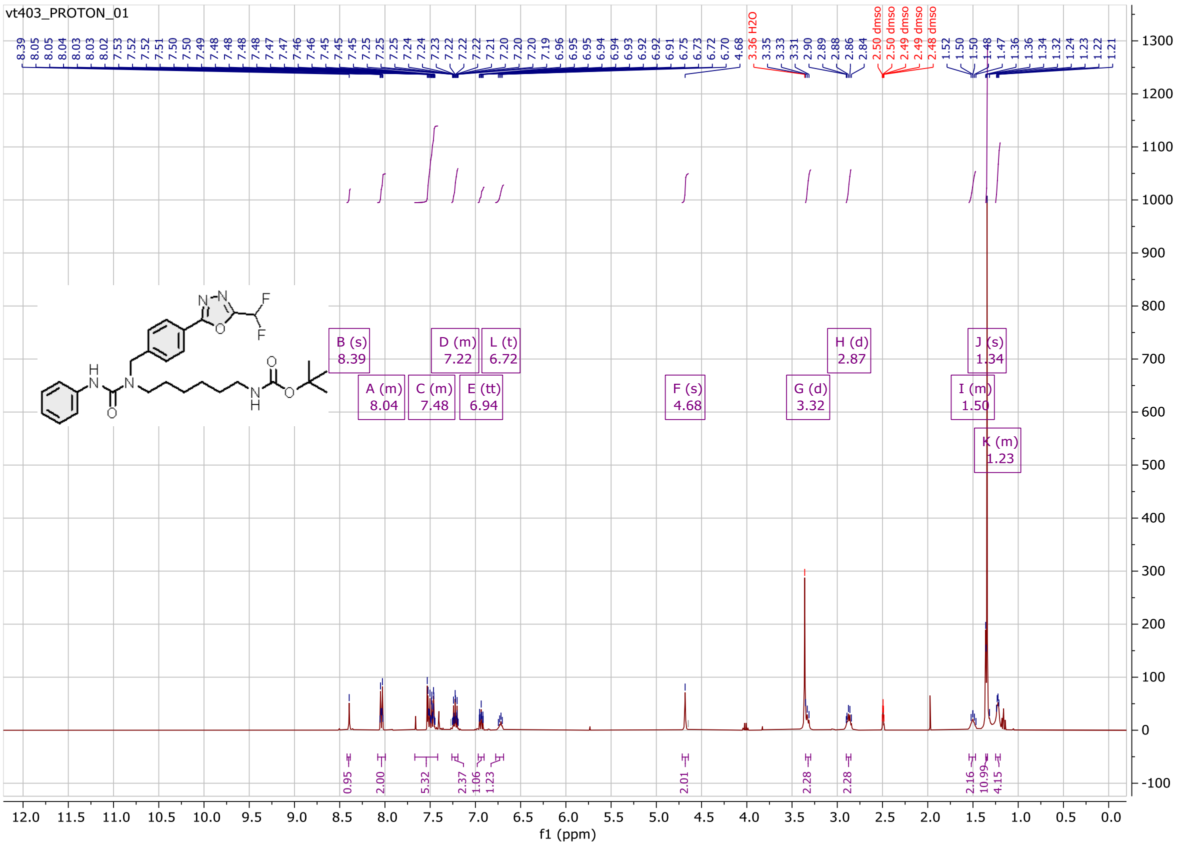

$^{13}$C NMR spectrum

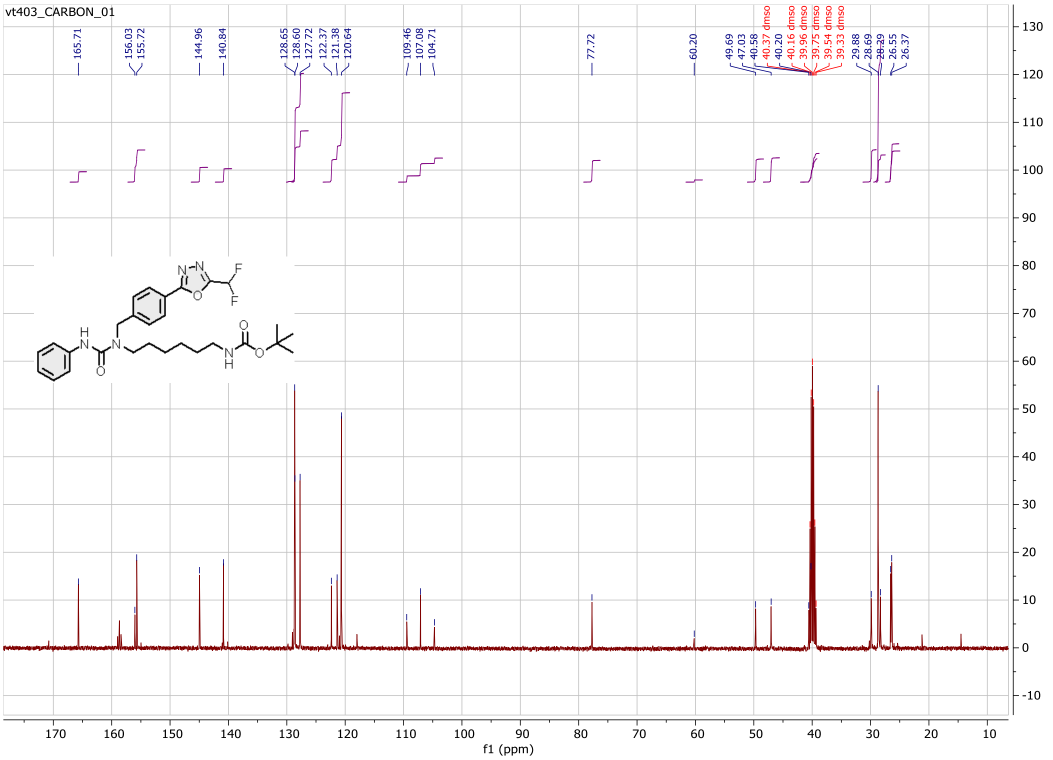

Compound **11b**:

$^{1}$H NMR spectrum

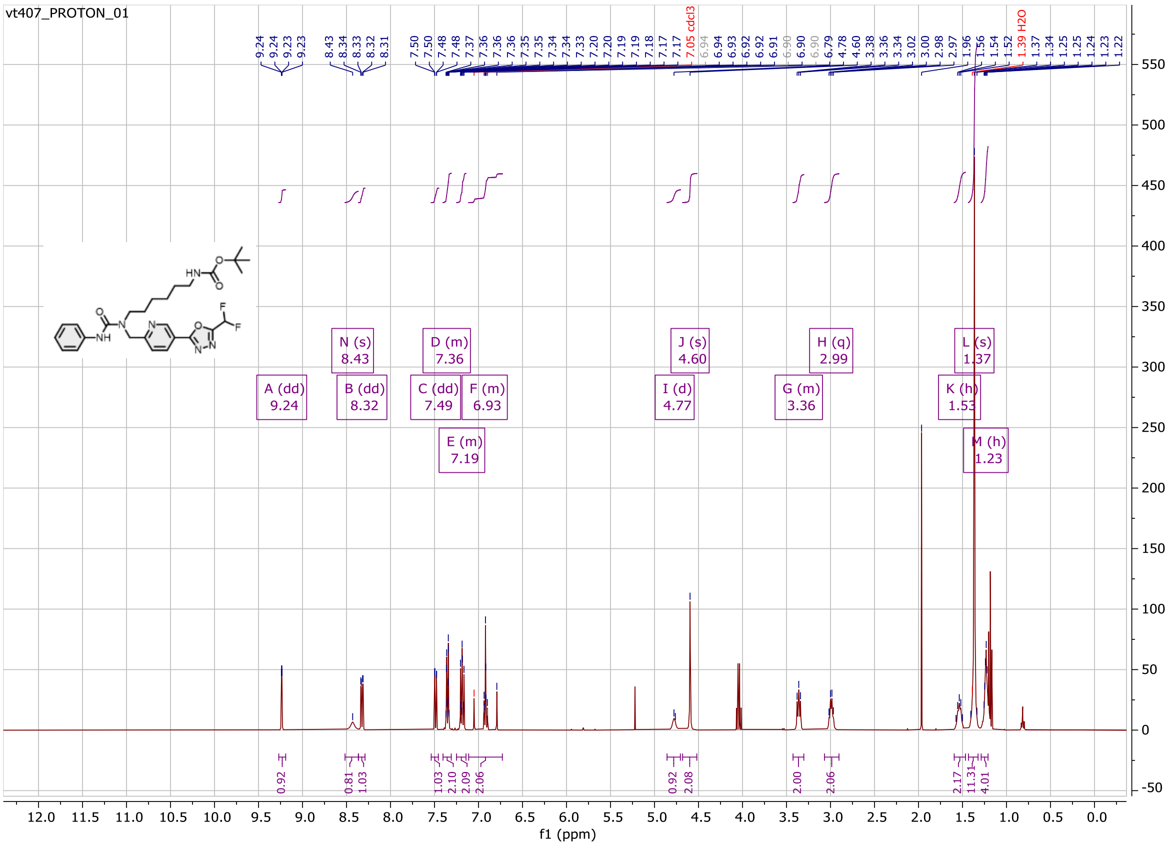

$^{13}$C NMR spectrum

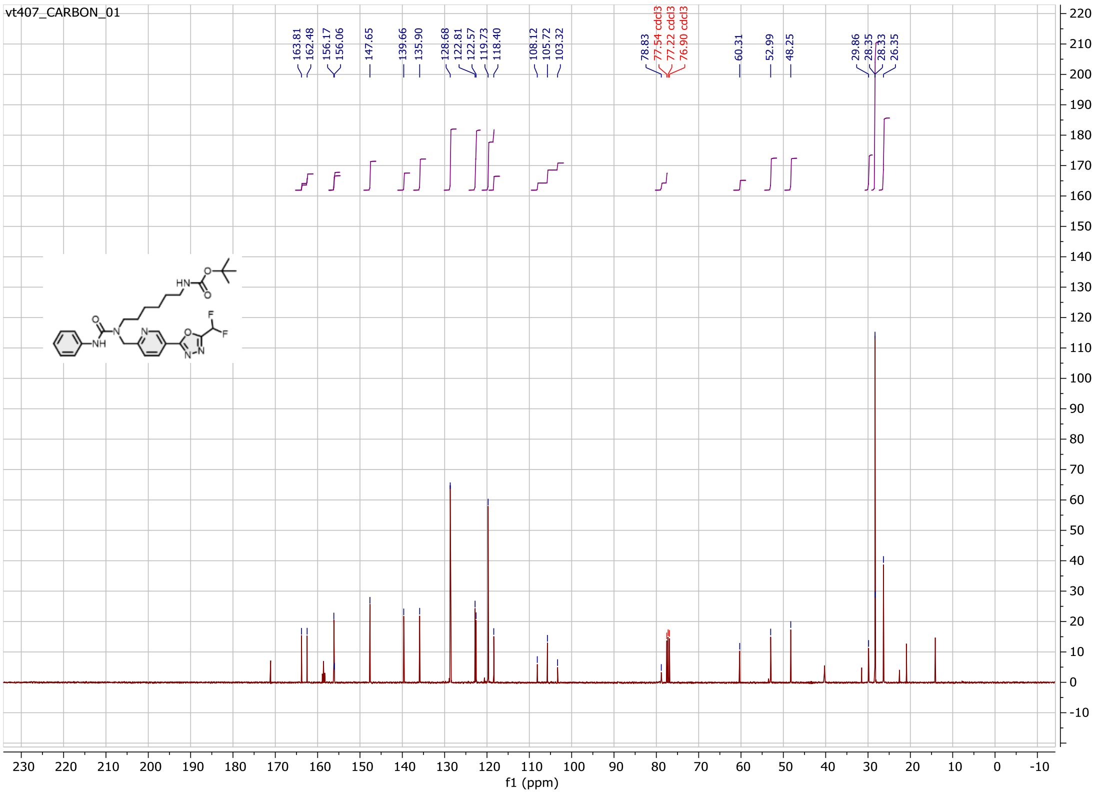

Compound **12b**:

$^{1}$H NMR spectrum

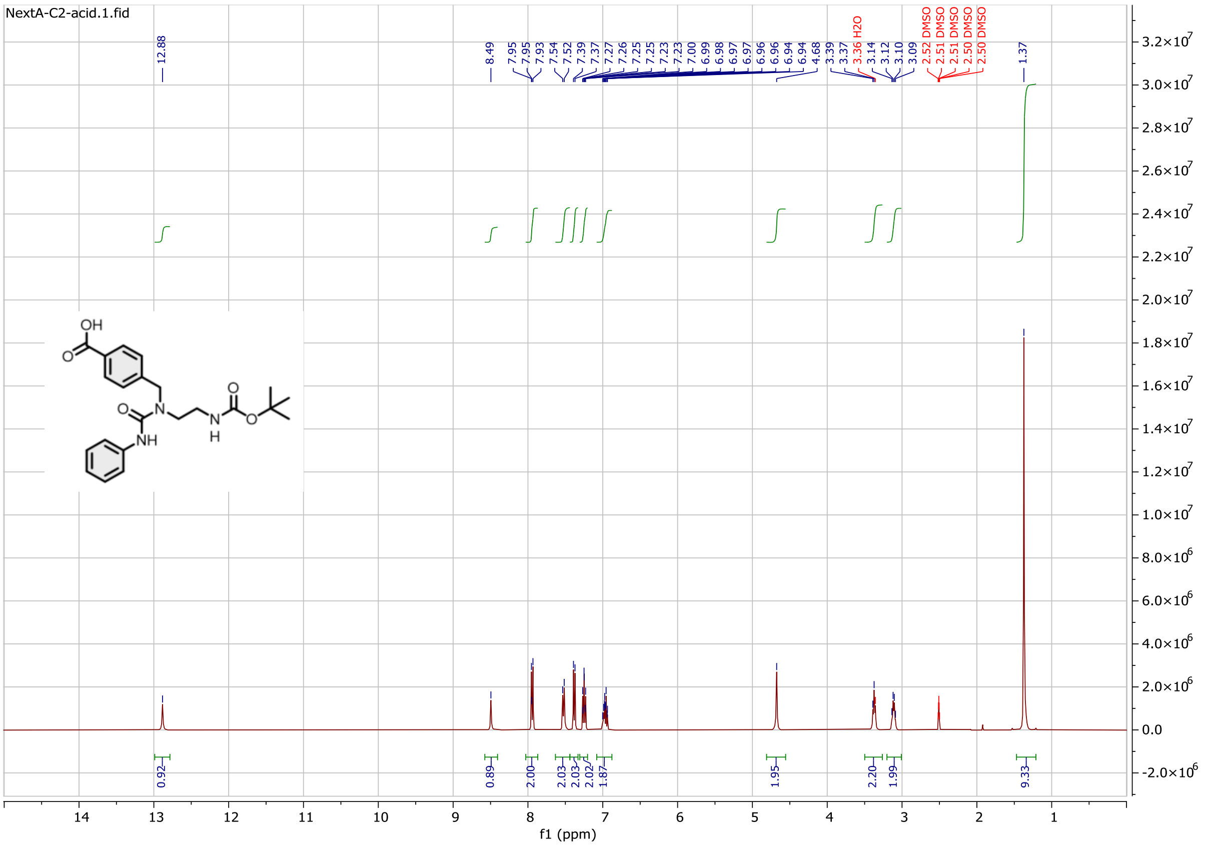

$^{13}$C NMR spectrum

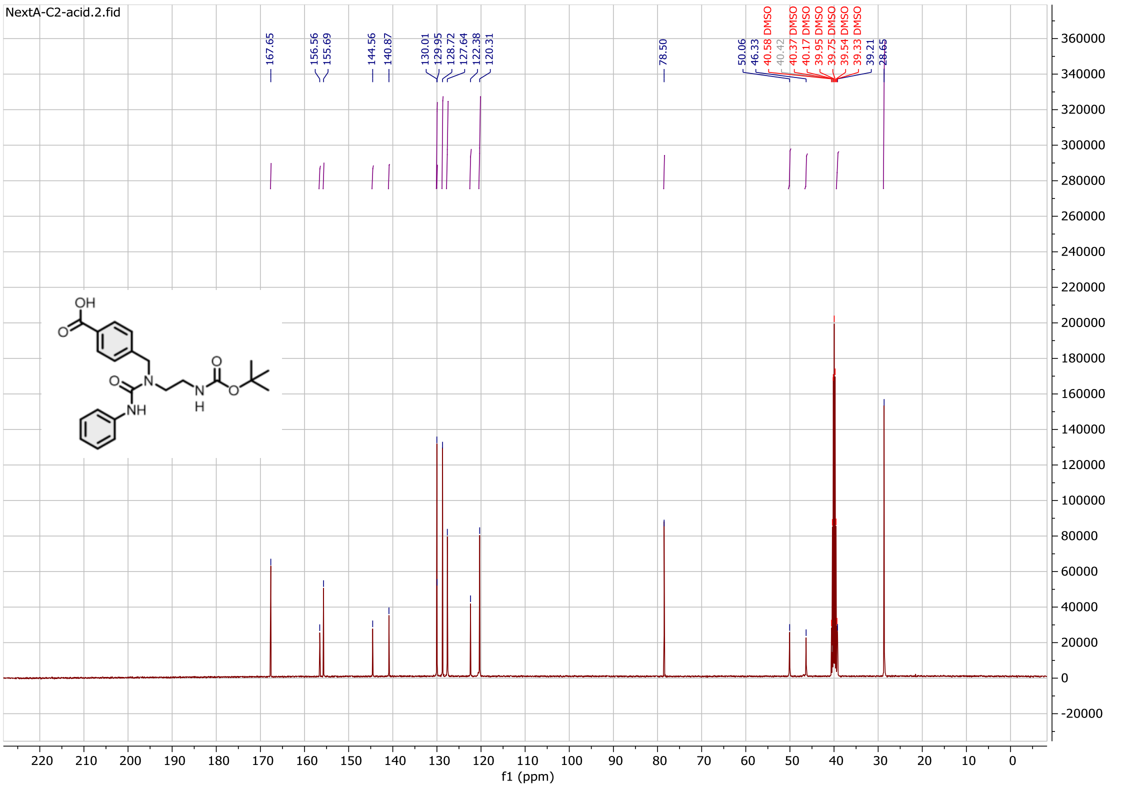

Compound **13b**:

$^{1}$H NMR spectrum

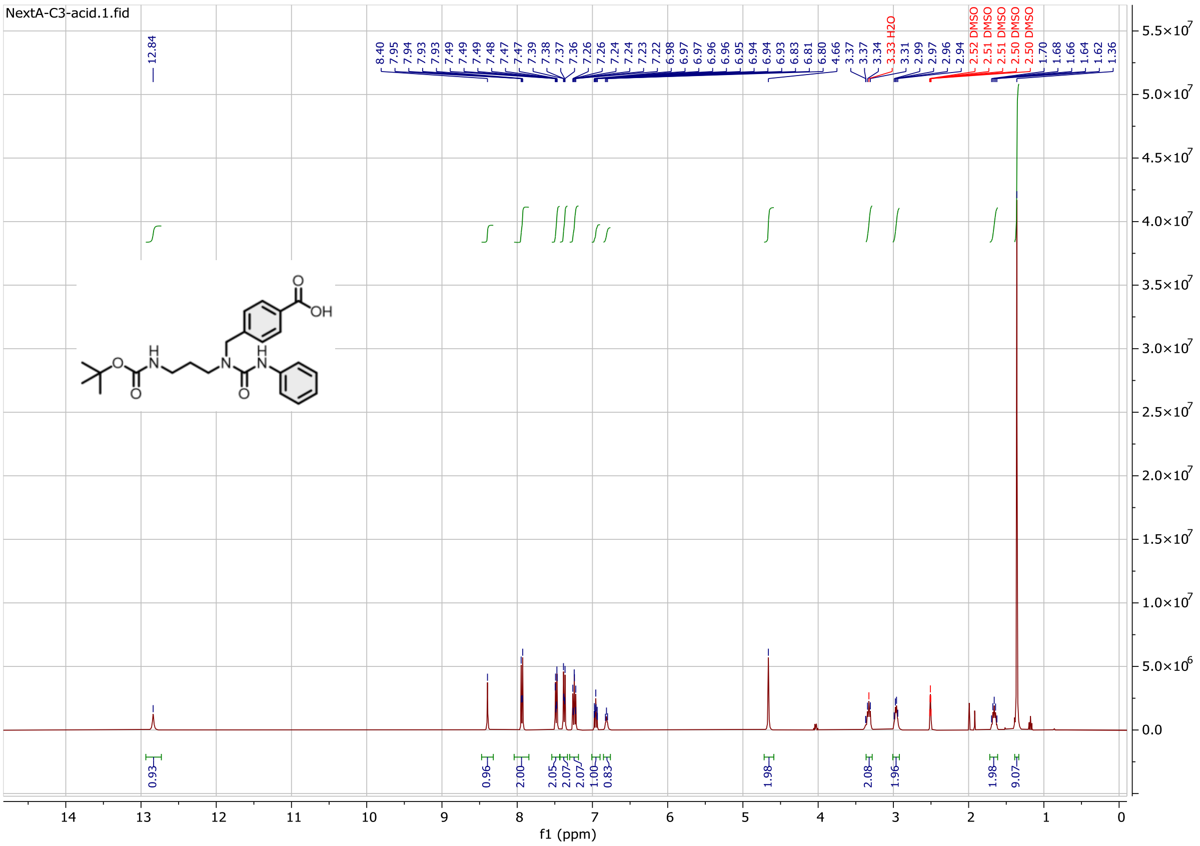

$^{13}$C NMR spectrum

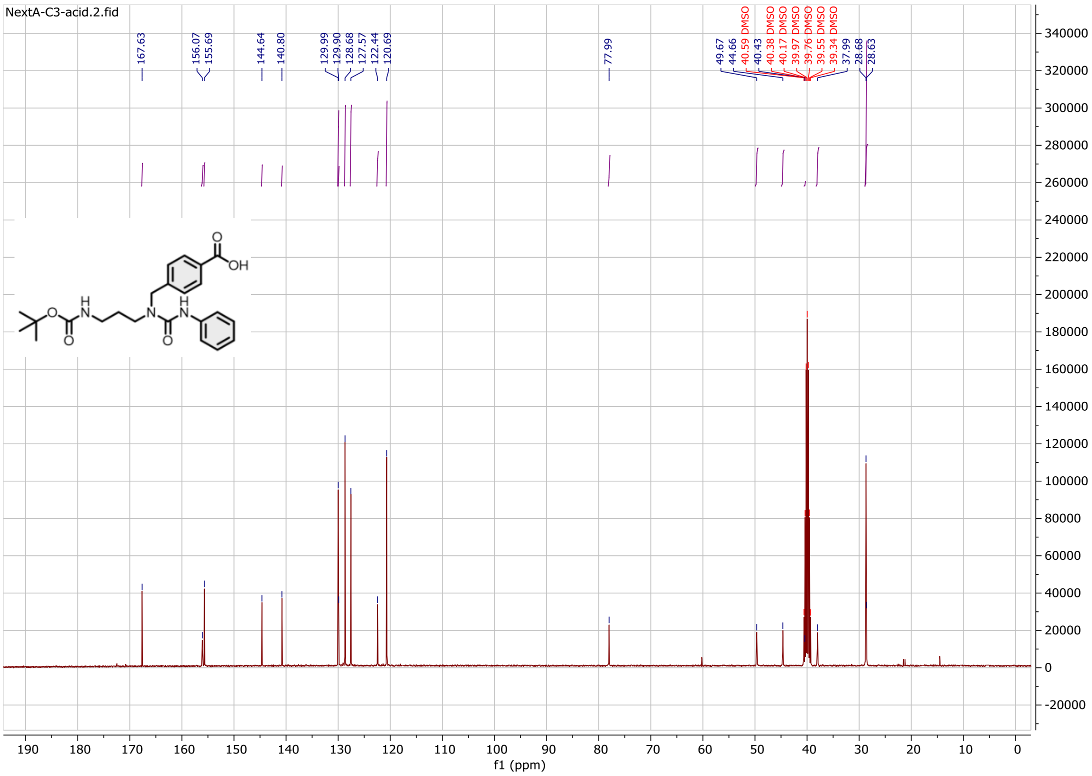

Compound **14b**:

$^{1}$H NMR spectrum

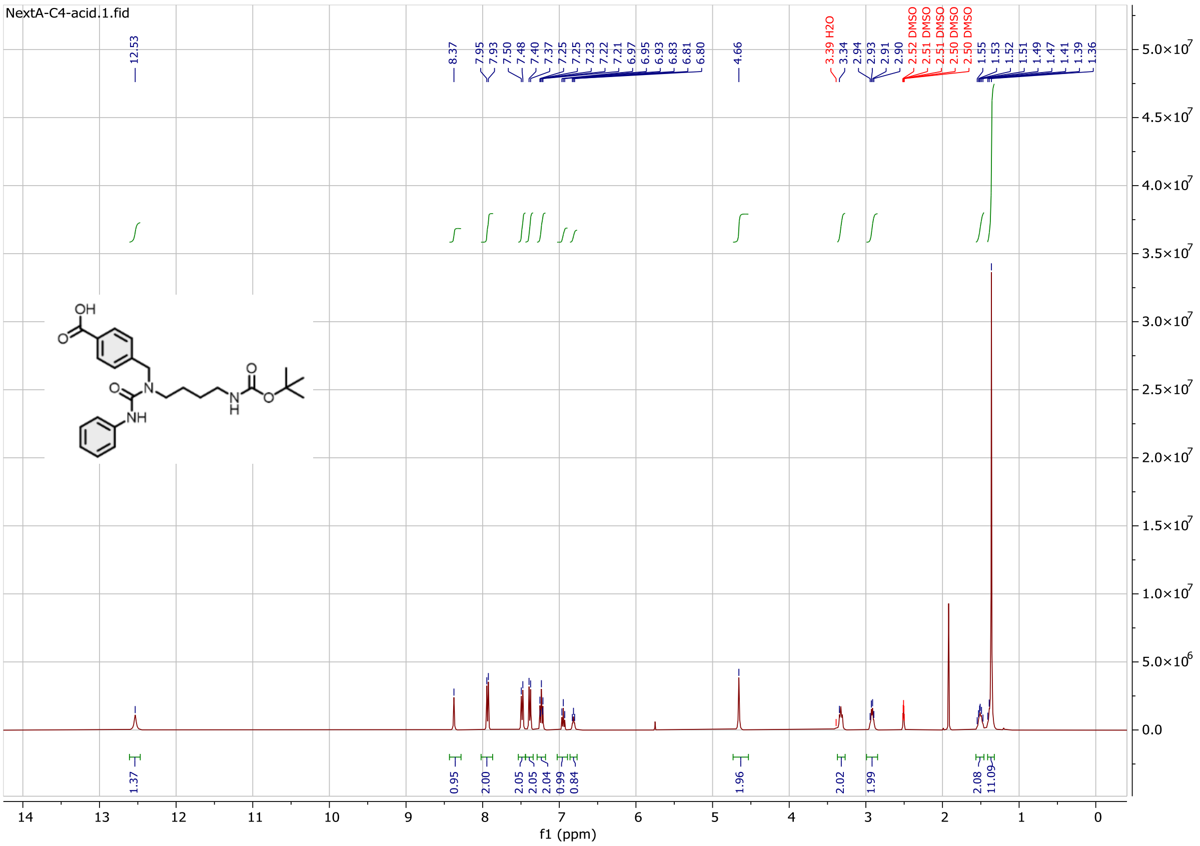

$^{13}$C NMR spectrum

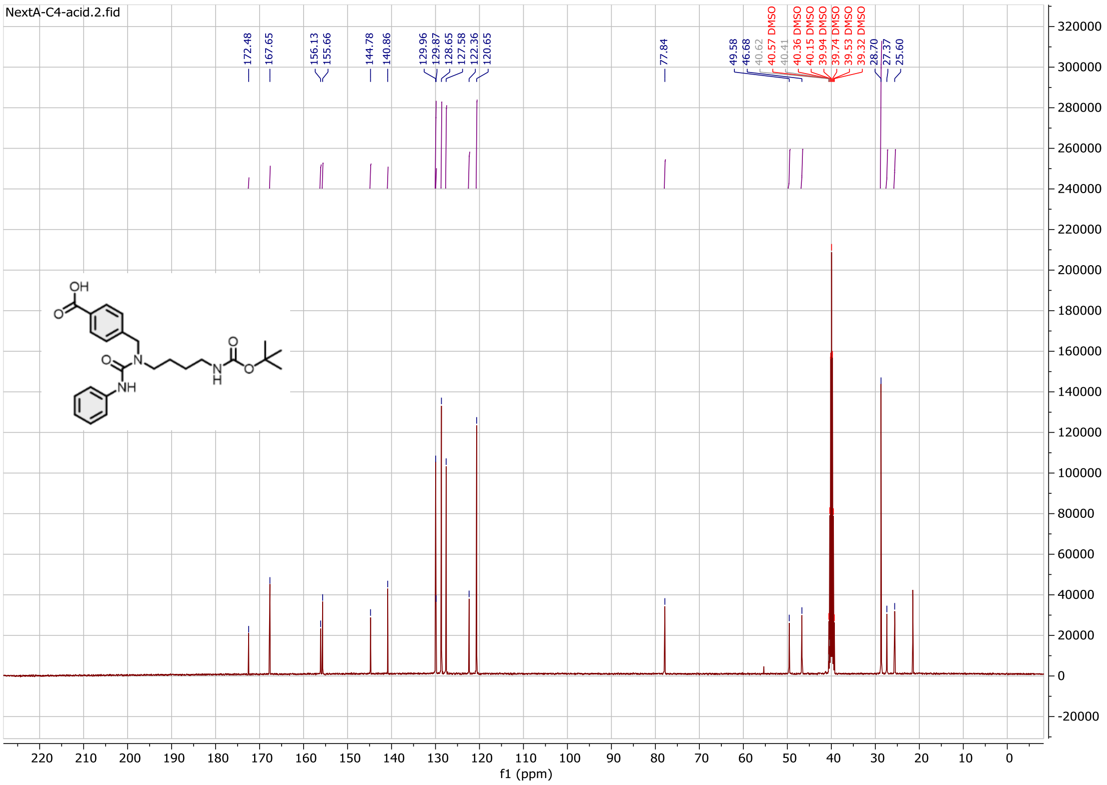

Compound **15b**:

$^{1}$H NMR spectrum

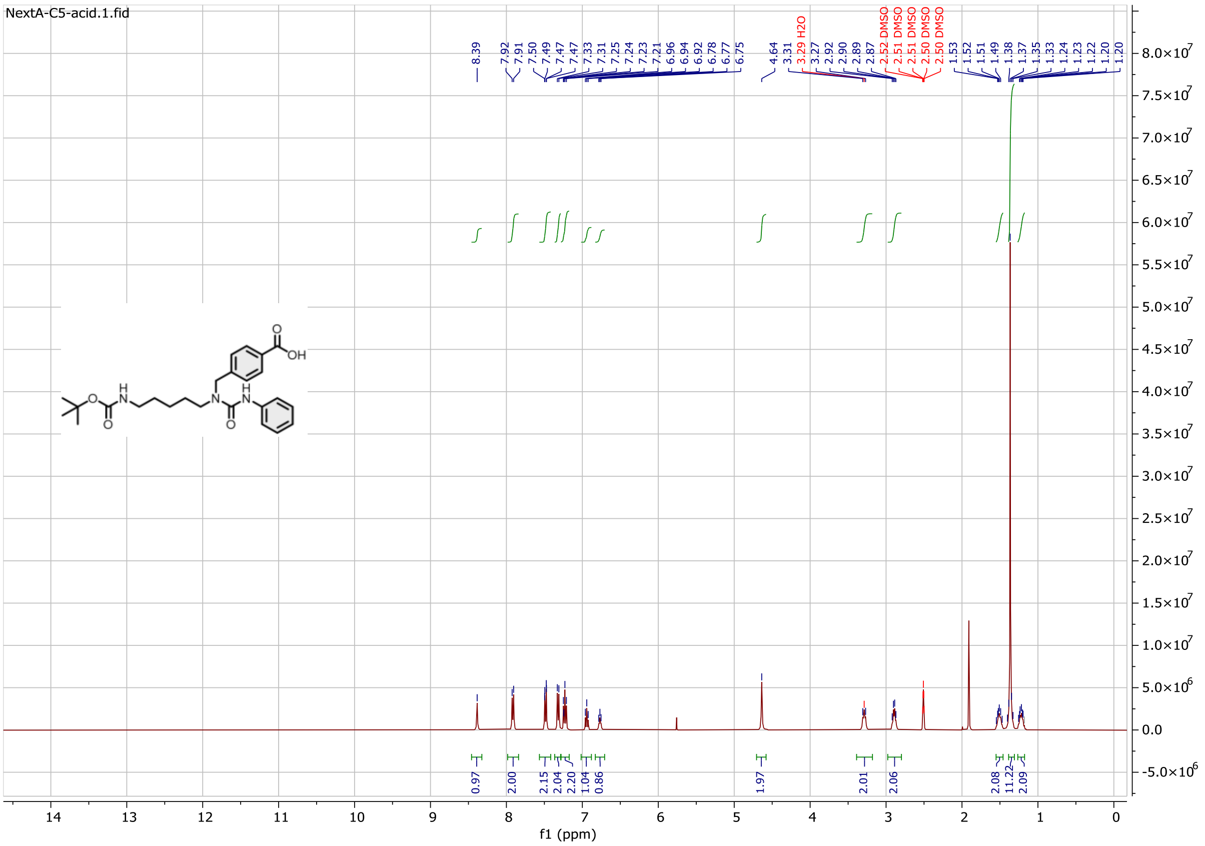

$^{13}$C NMR spectrum

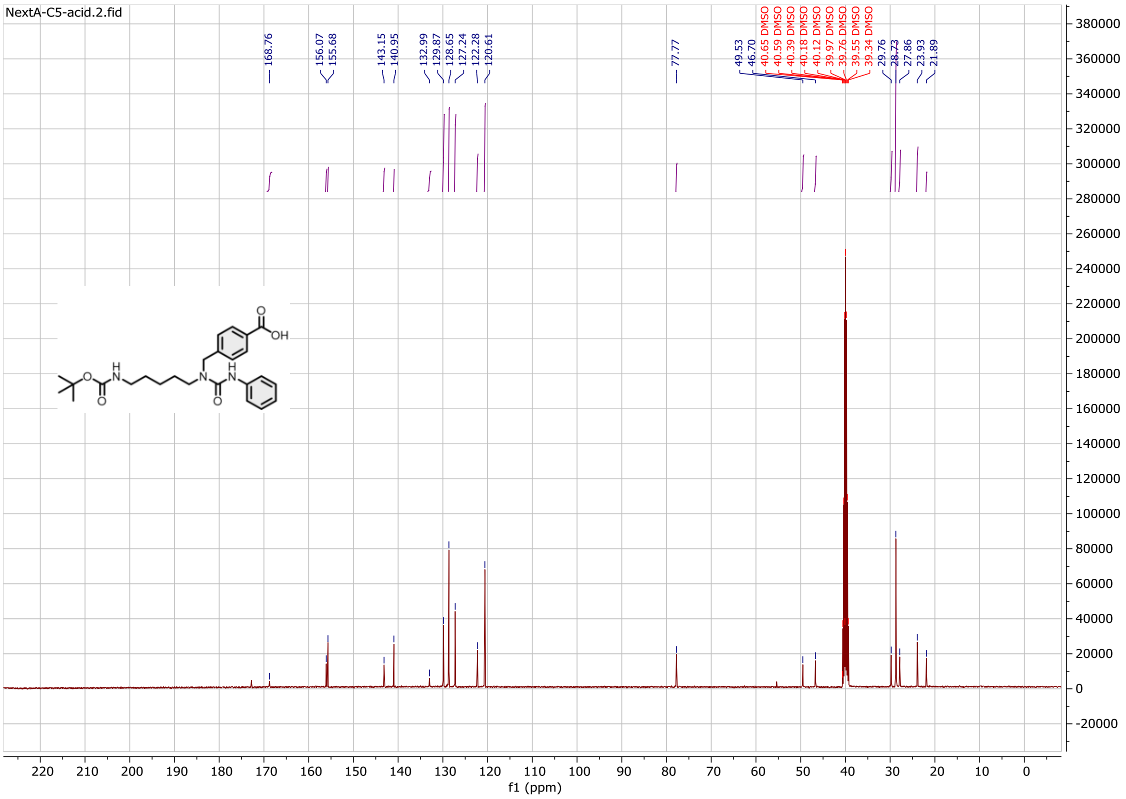

Compound **16b**:

$^{1}$H NMR spectrum

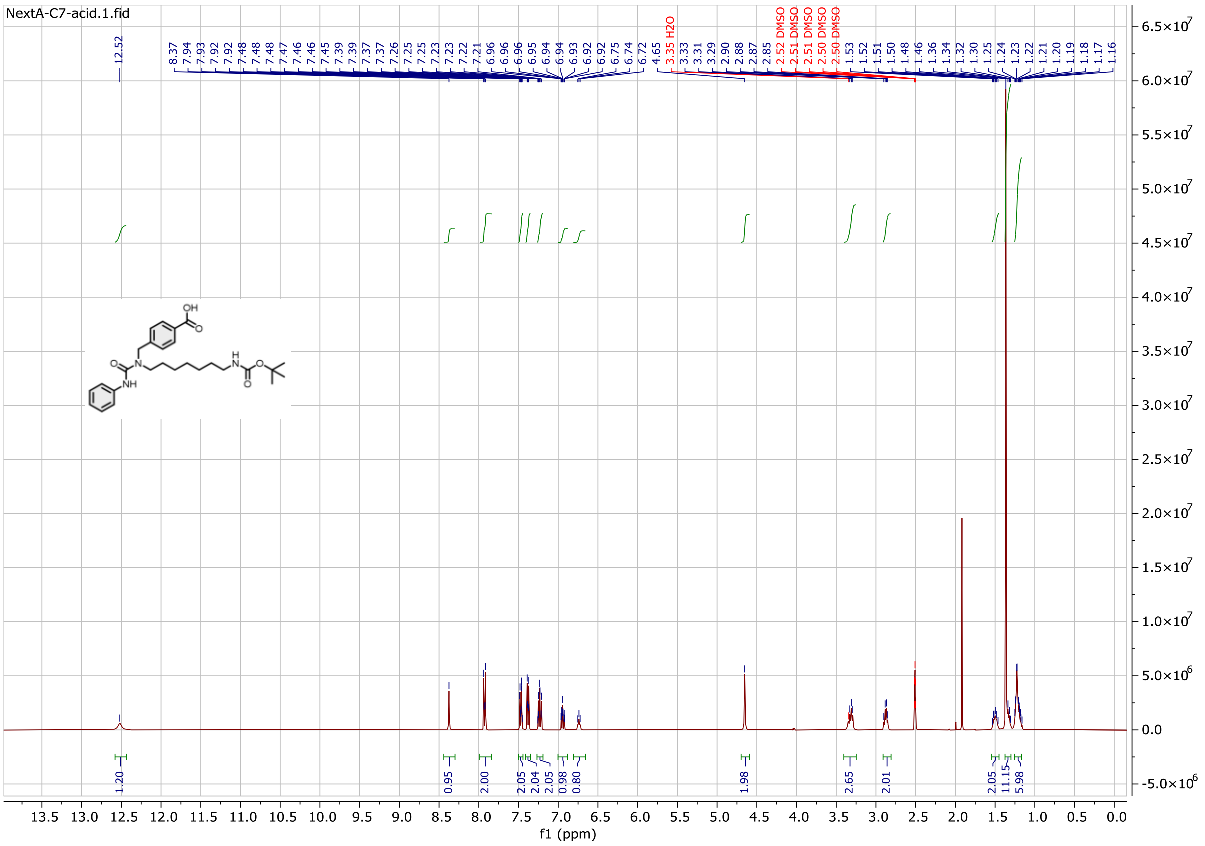

$^{13}$C NMR spectrum

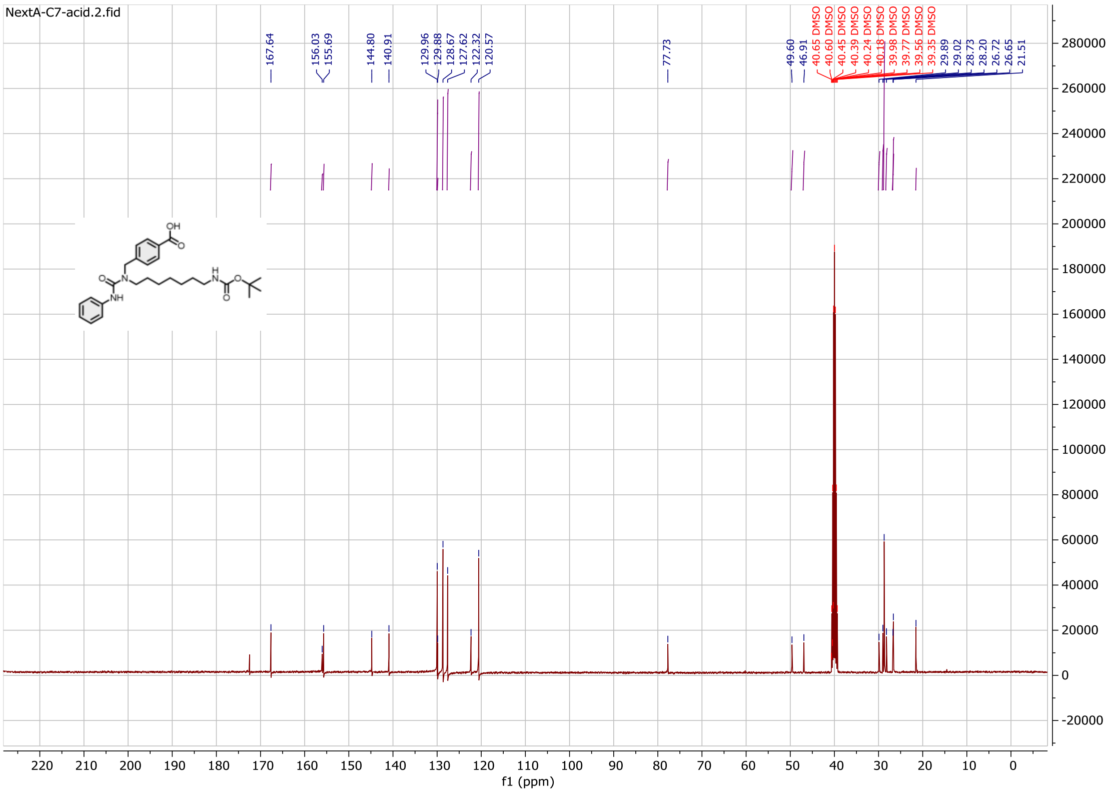

Compound **17b**:

$^{1}$H NMR spectrum

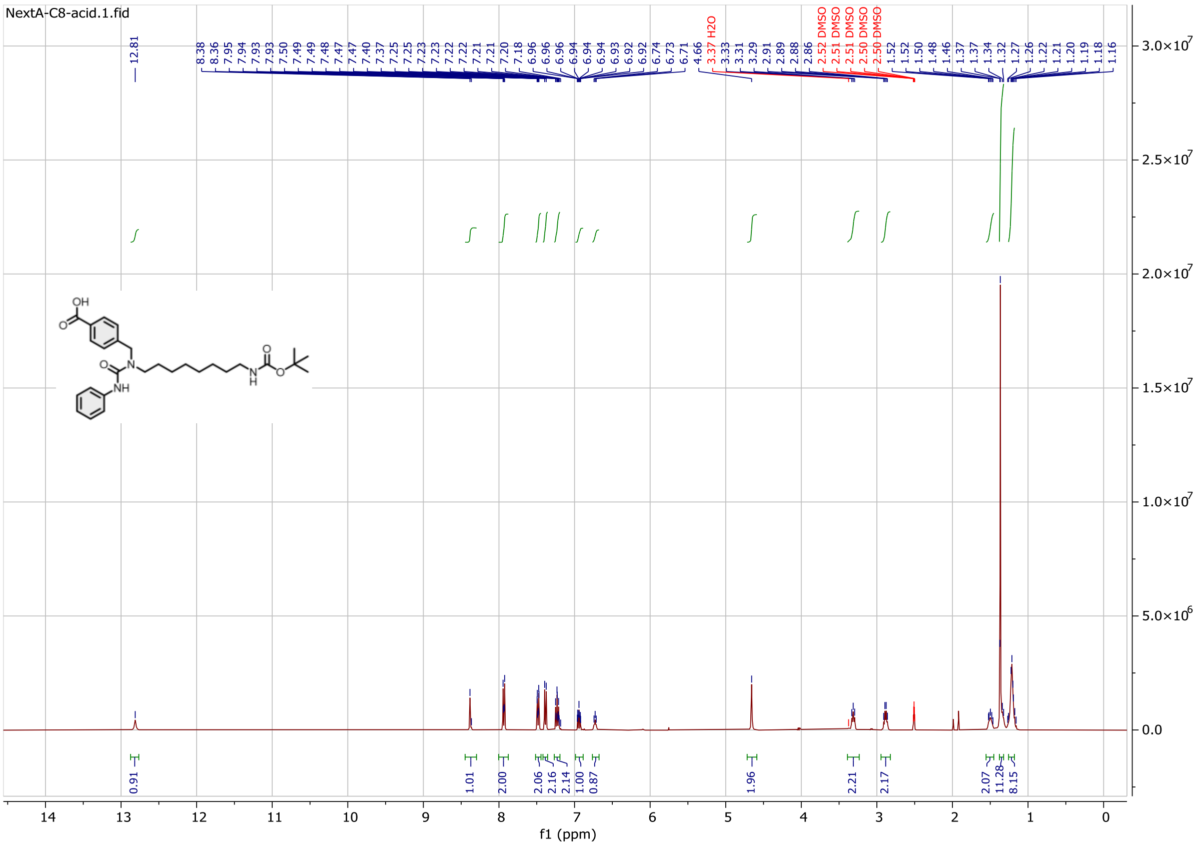

$^{13}$C NMR spectrum

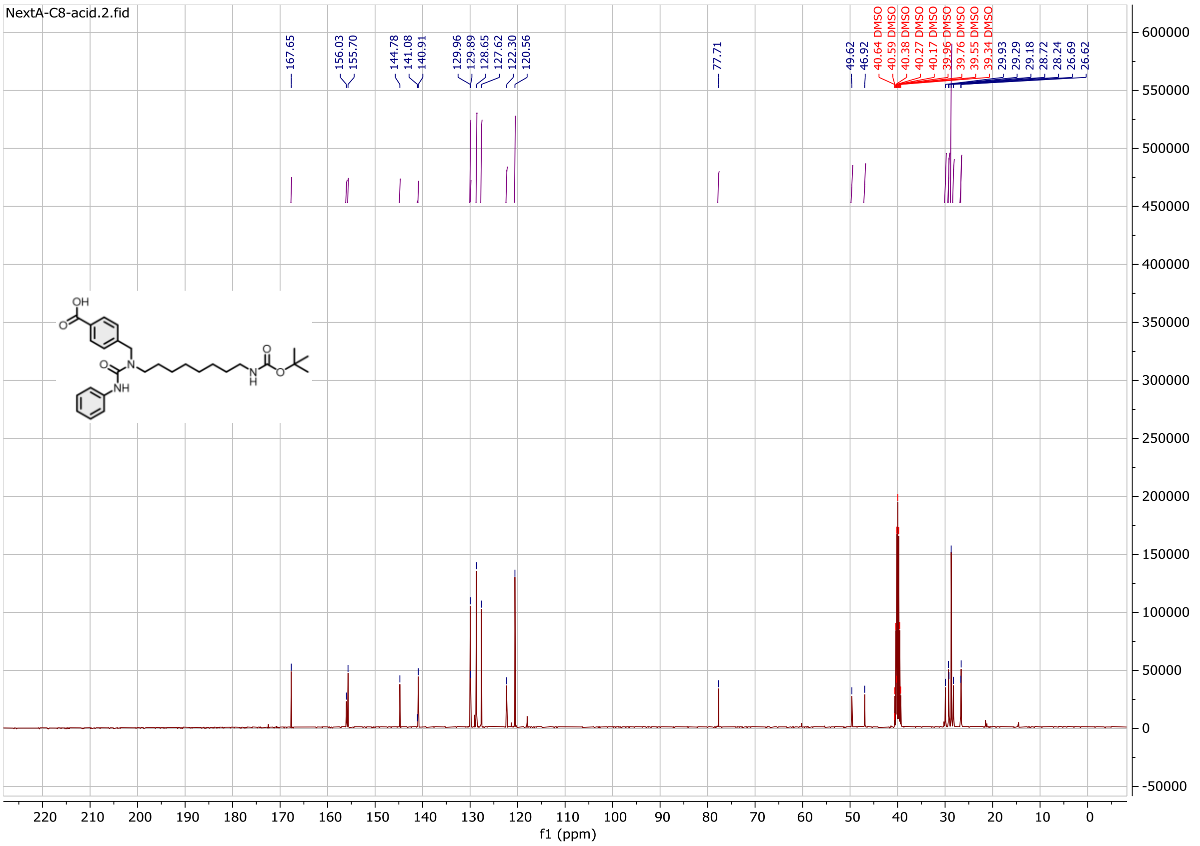

Compound **18b**:

$^{1}$H NMR spectrum

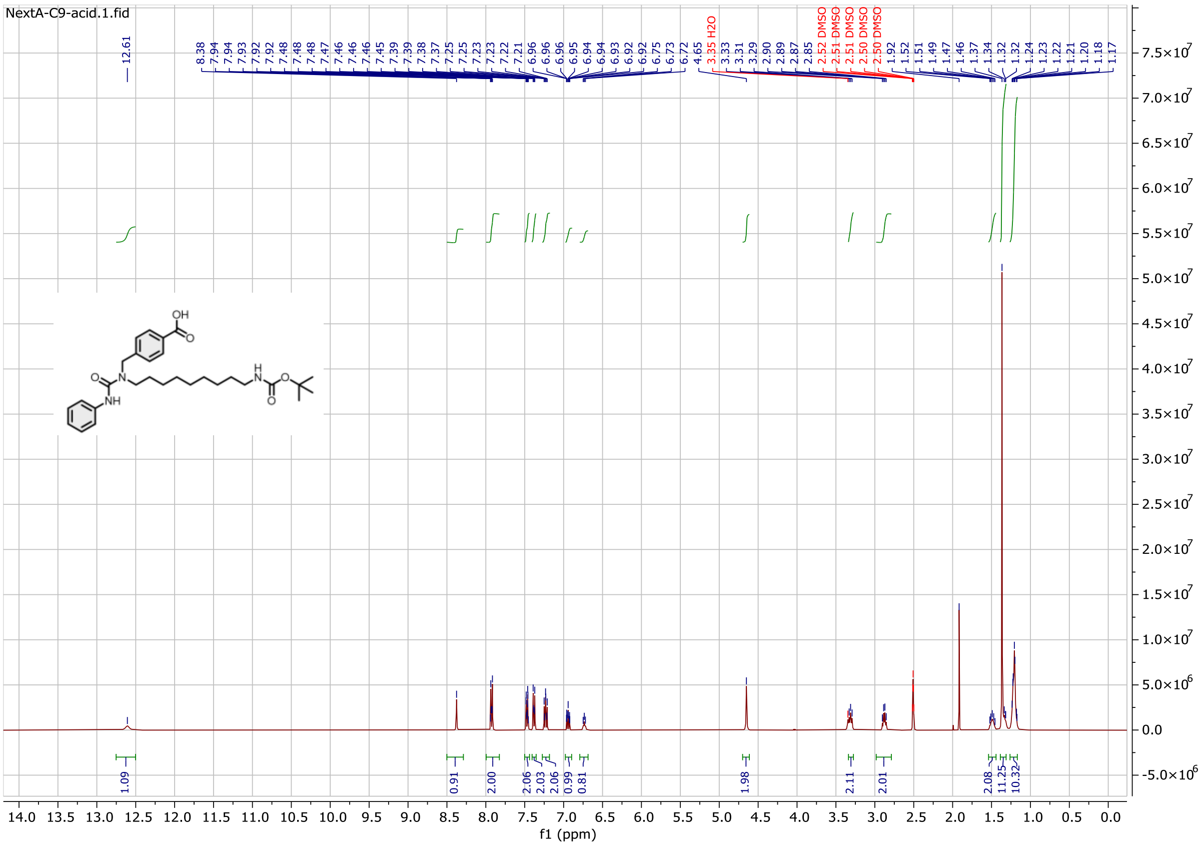

$^{13}$C NMR spectrum

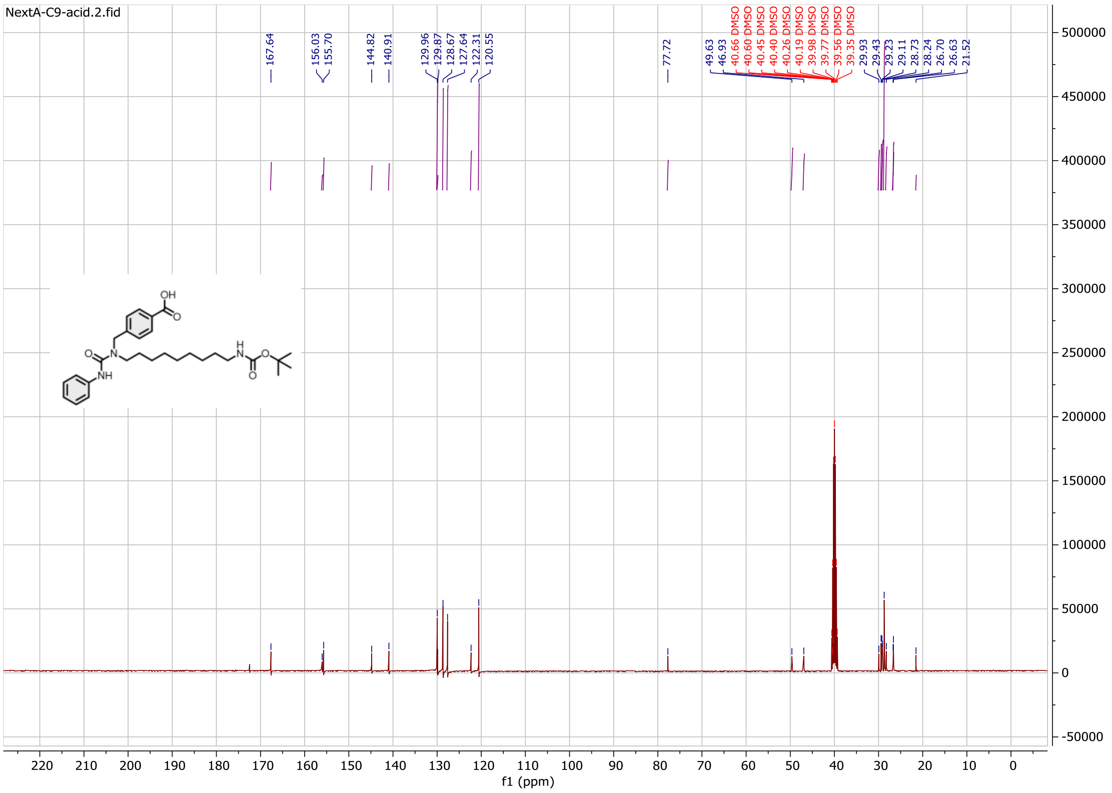

Compound **19b**:

$^{1}$H NMR spectrum

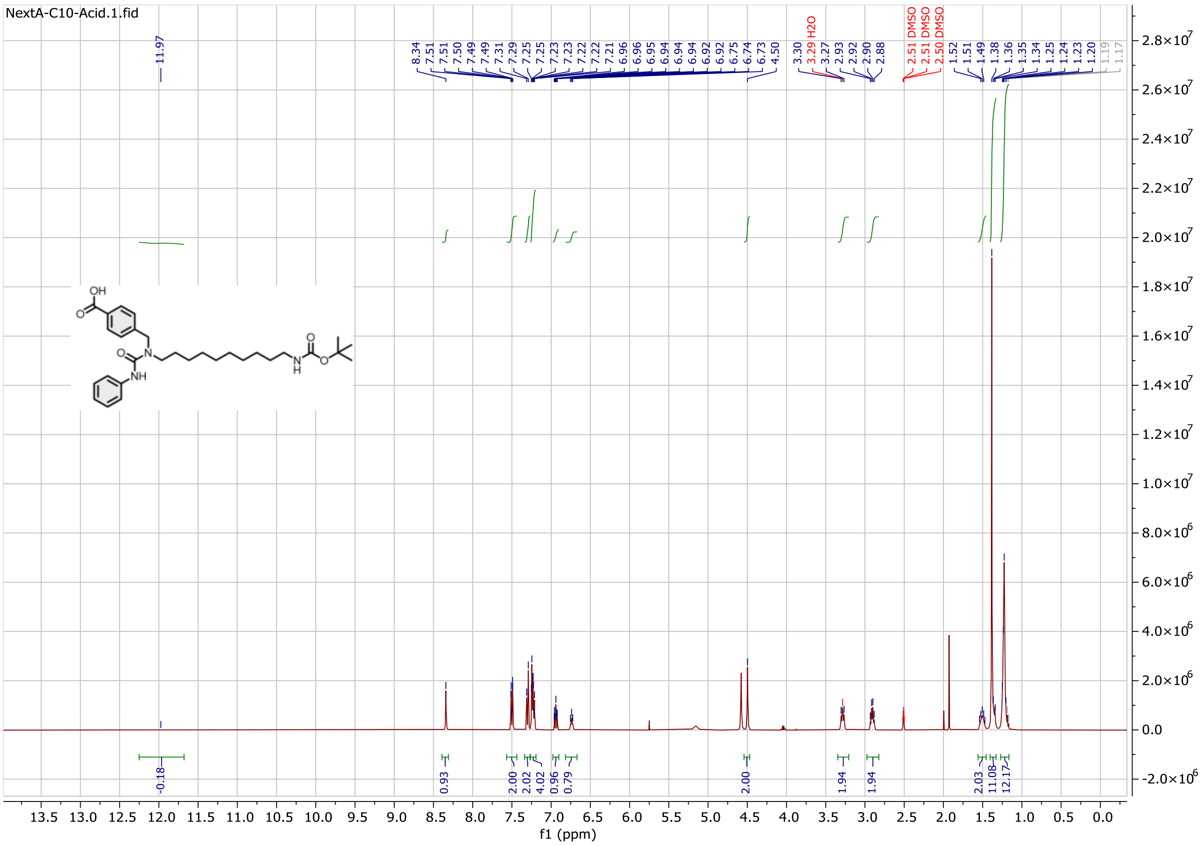

$^{13}$C NMR spectrum

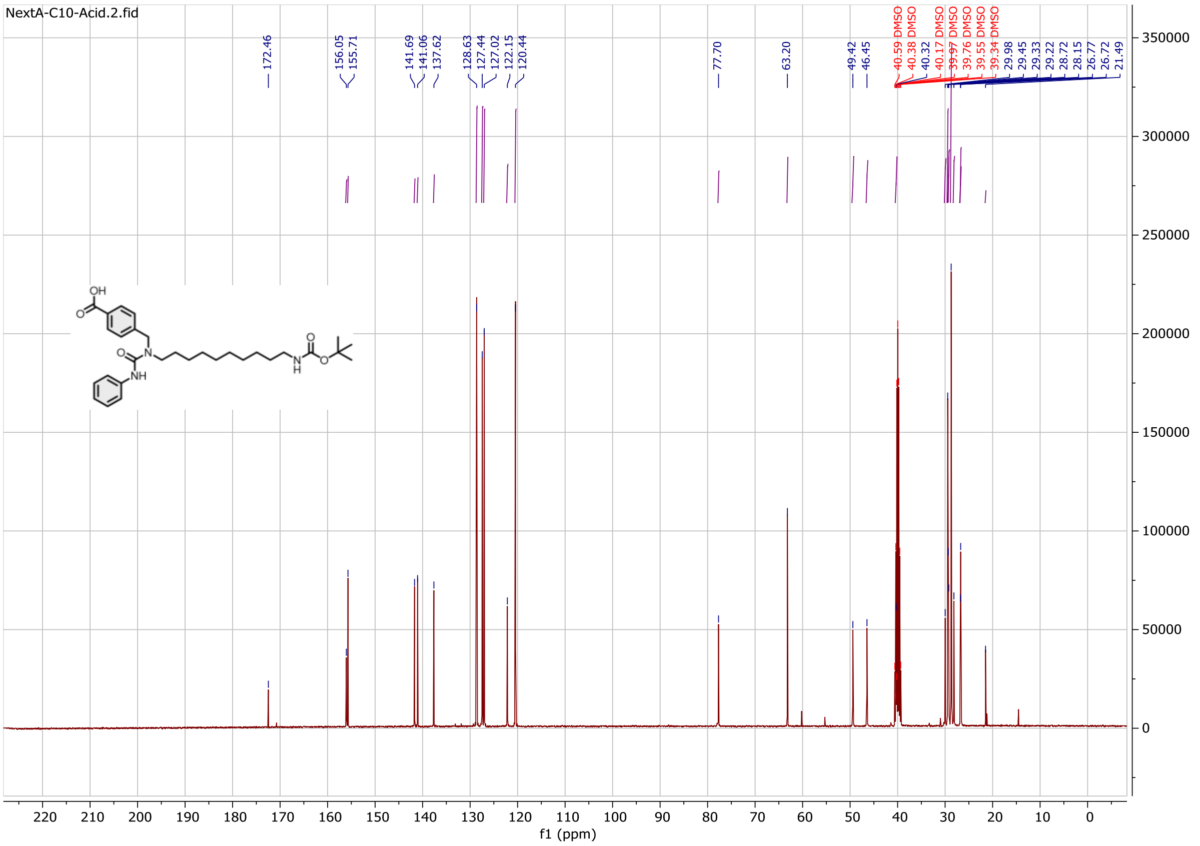

Compound **20b**:

$^{1}$H NMR spectrum

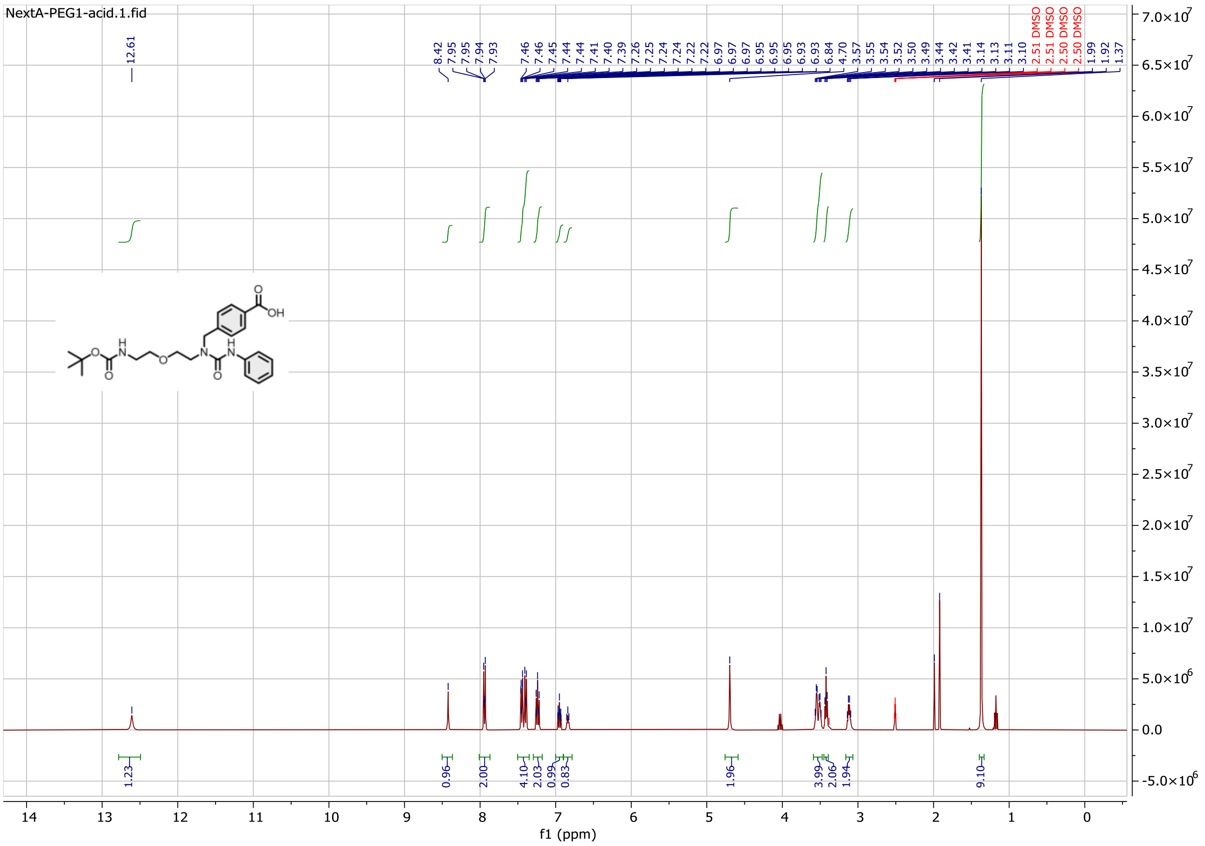

$^{13}$C NMR spectrum

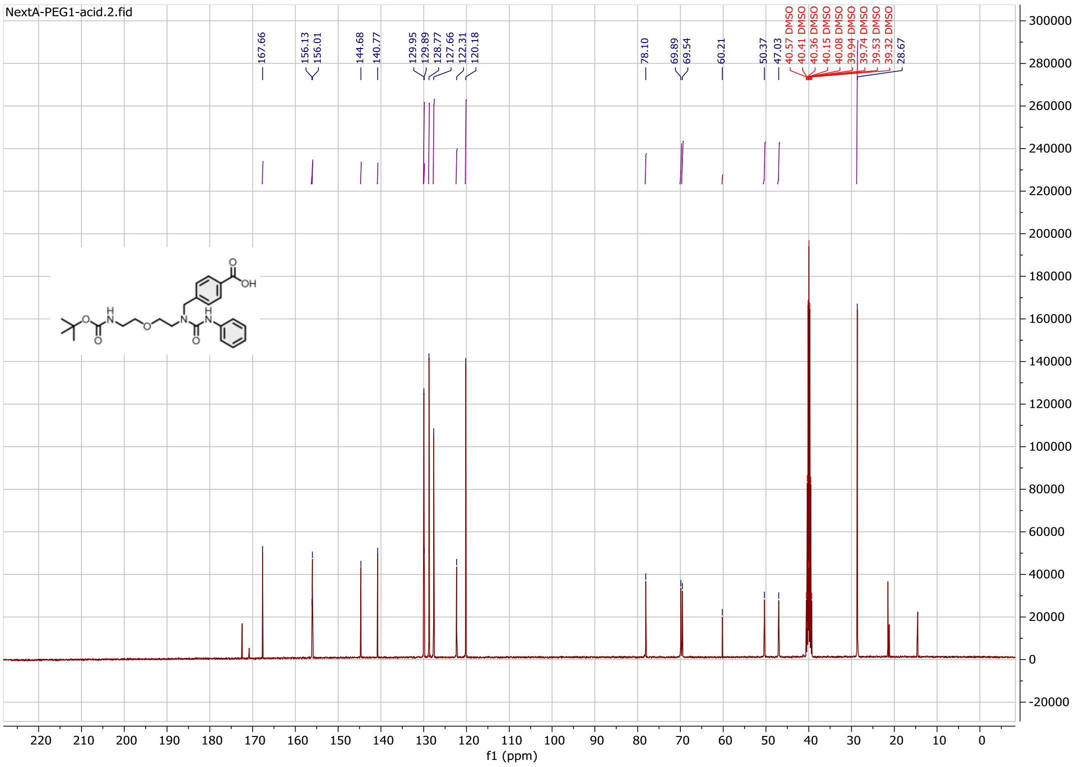

Compound **21b**:

$^{1}$H NMR spectrum

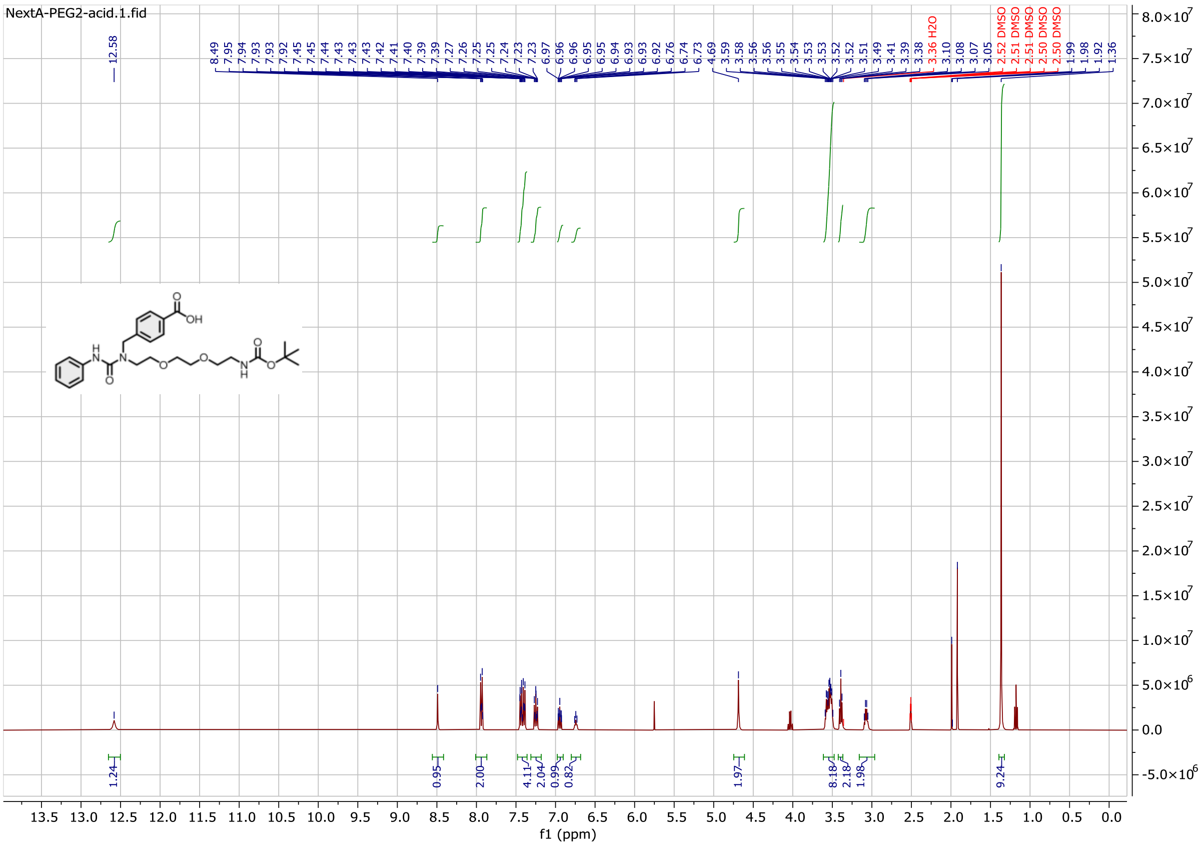

$^{13}$C NMR spectrum

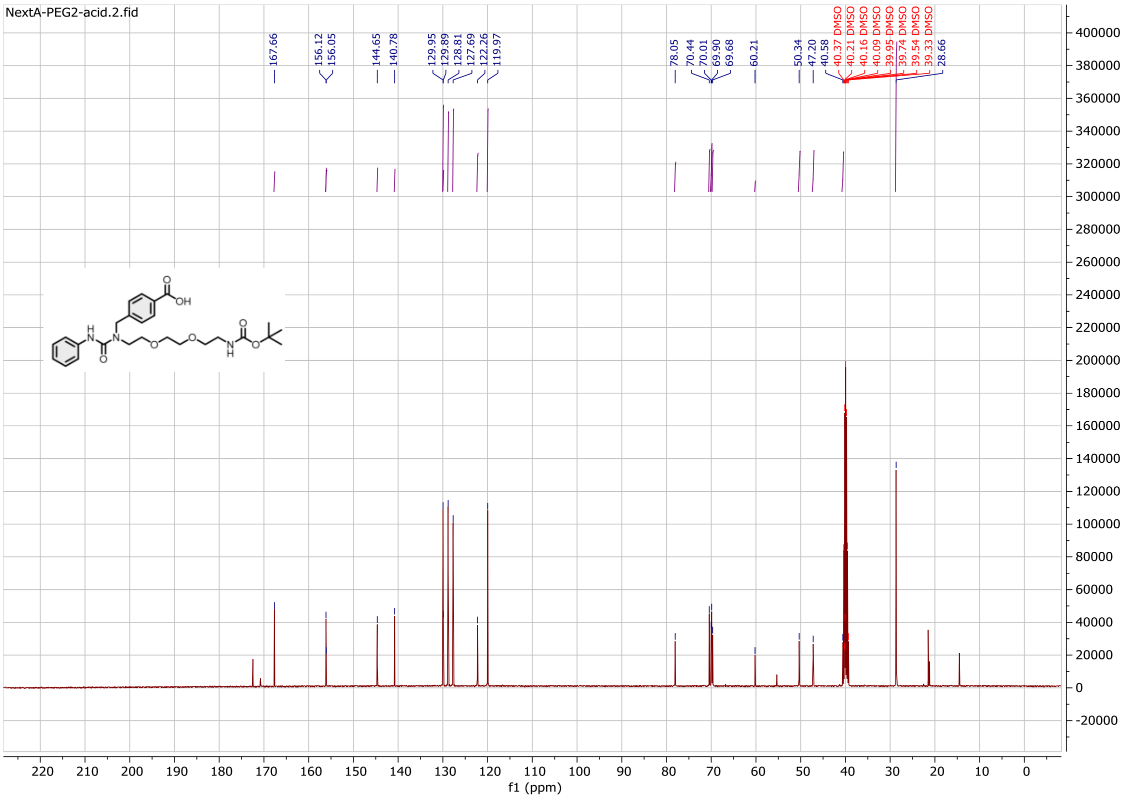

Compound **22b**:

$^{1}$H NMR spectrum

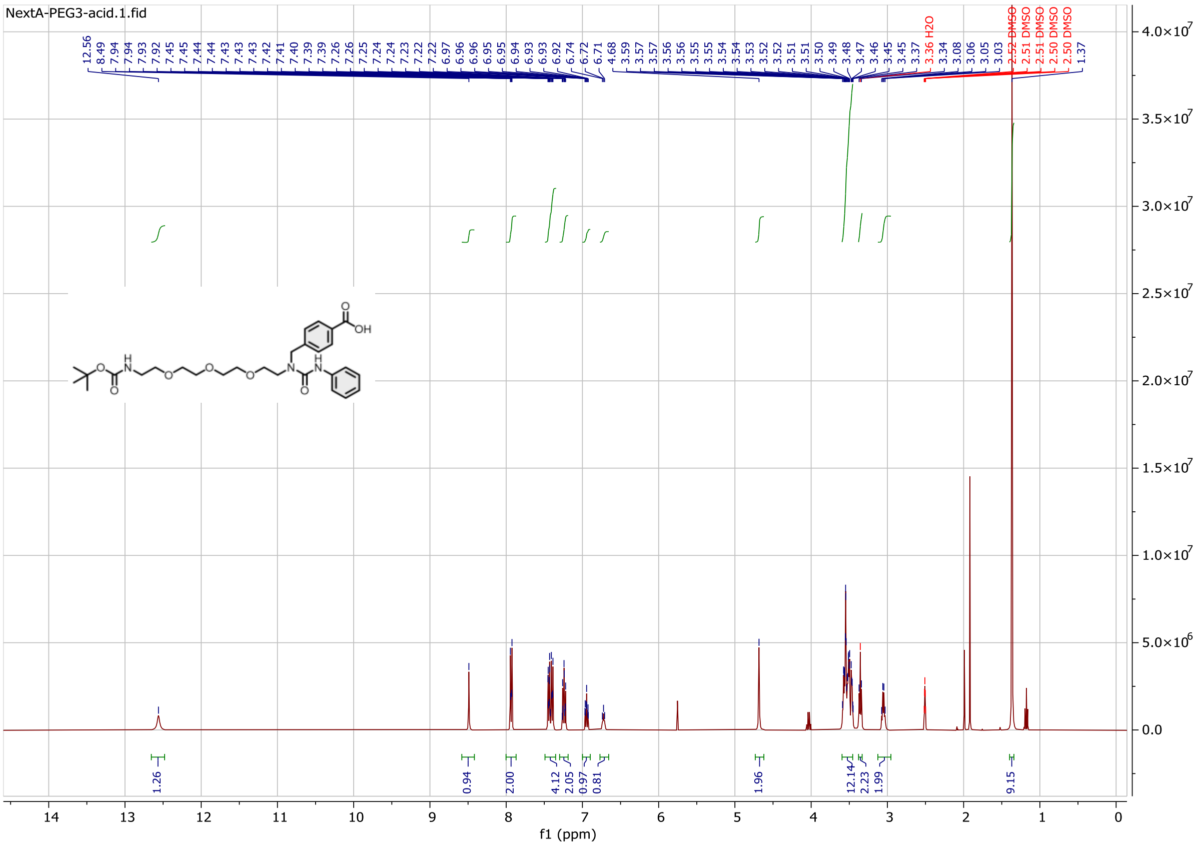

$^{13}$C NMR spectrum

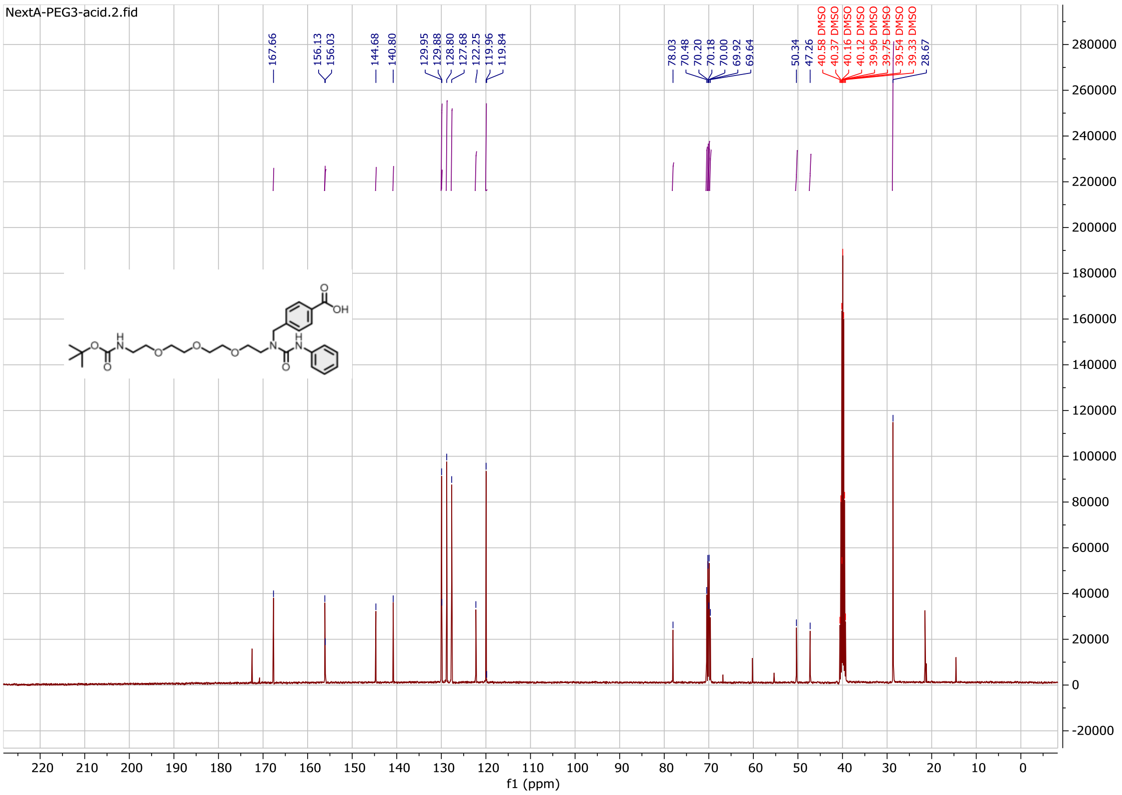

Compound **23b**:

$^{1}$H NMR spectrum

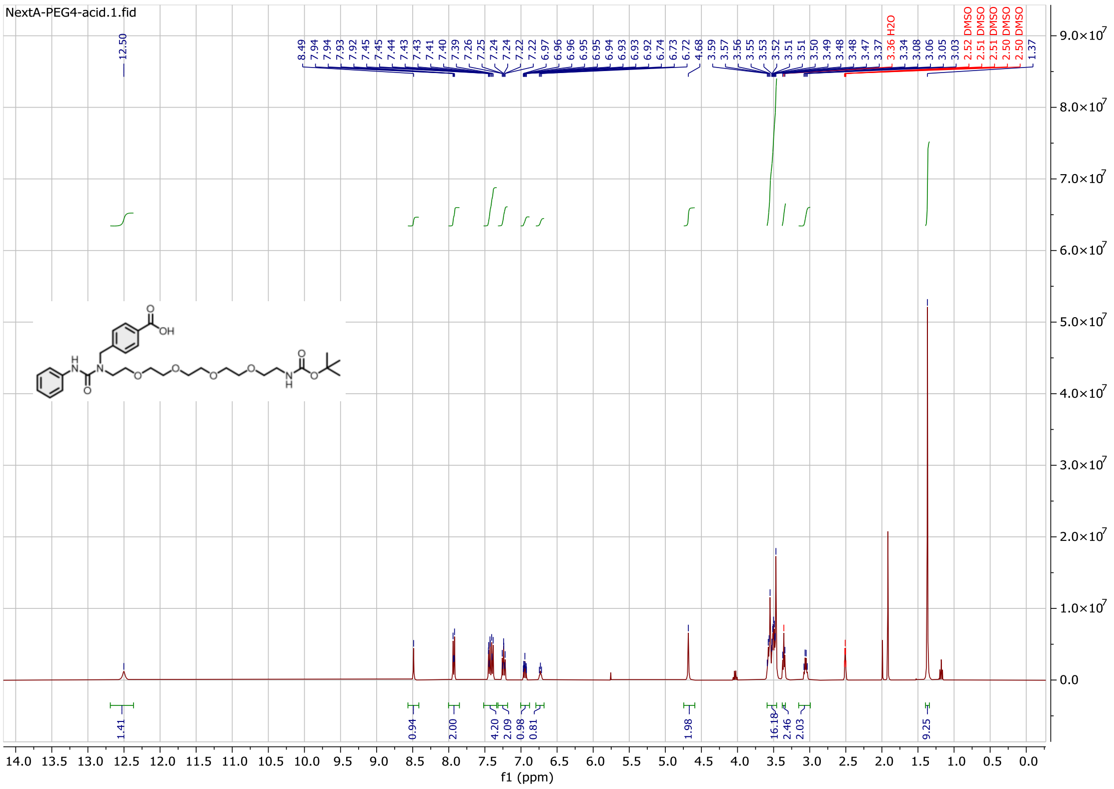

$^{13}$C NMR spectrum

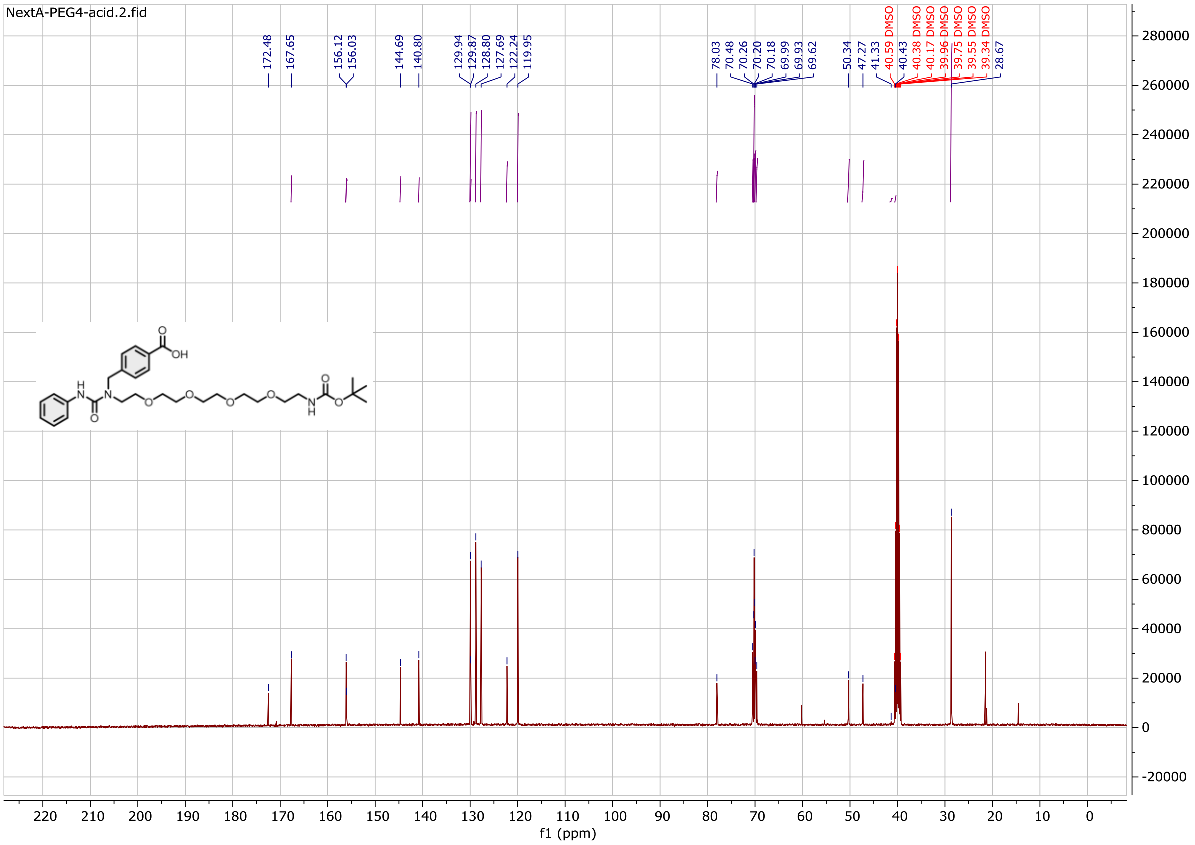

Compound **1c**:

$^{1}$H NMR spectrum

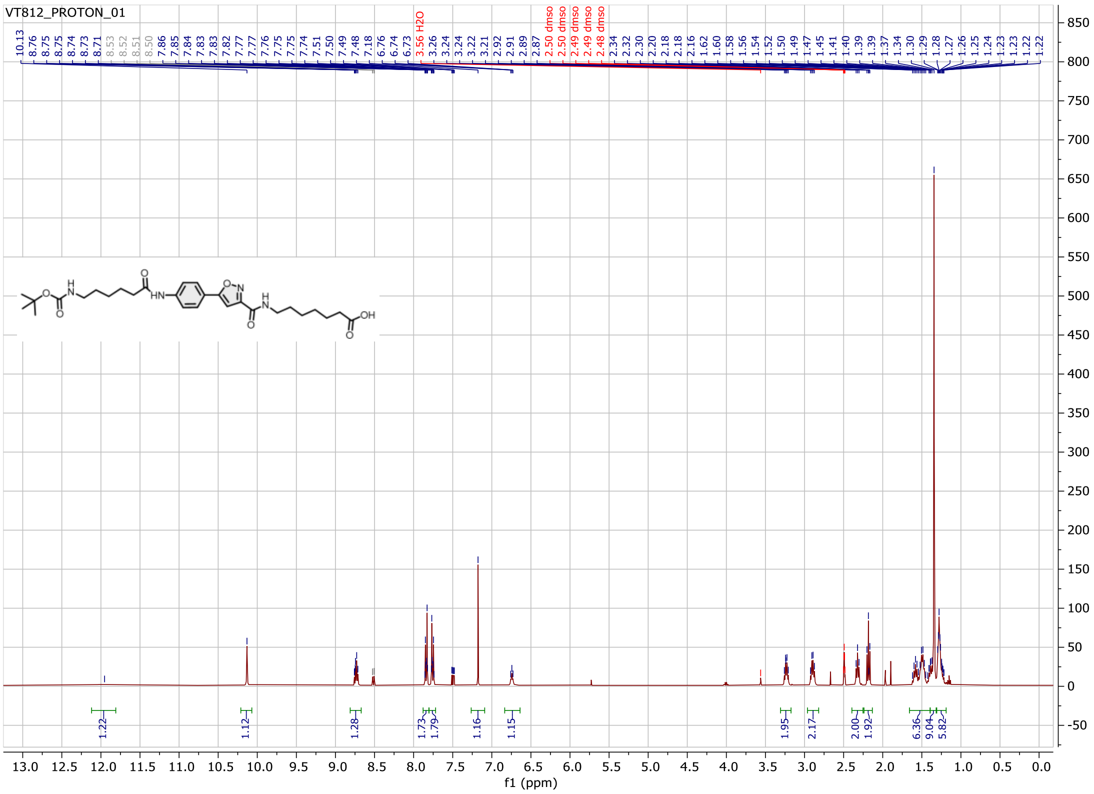

$^{13}$C NMR spectrum

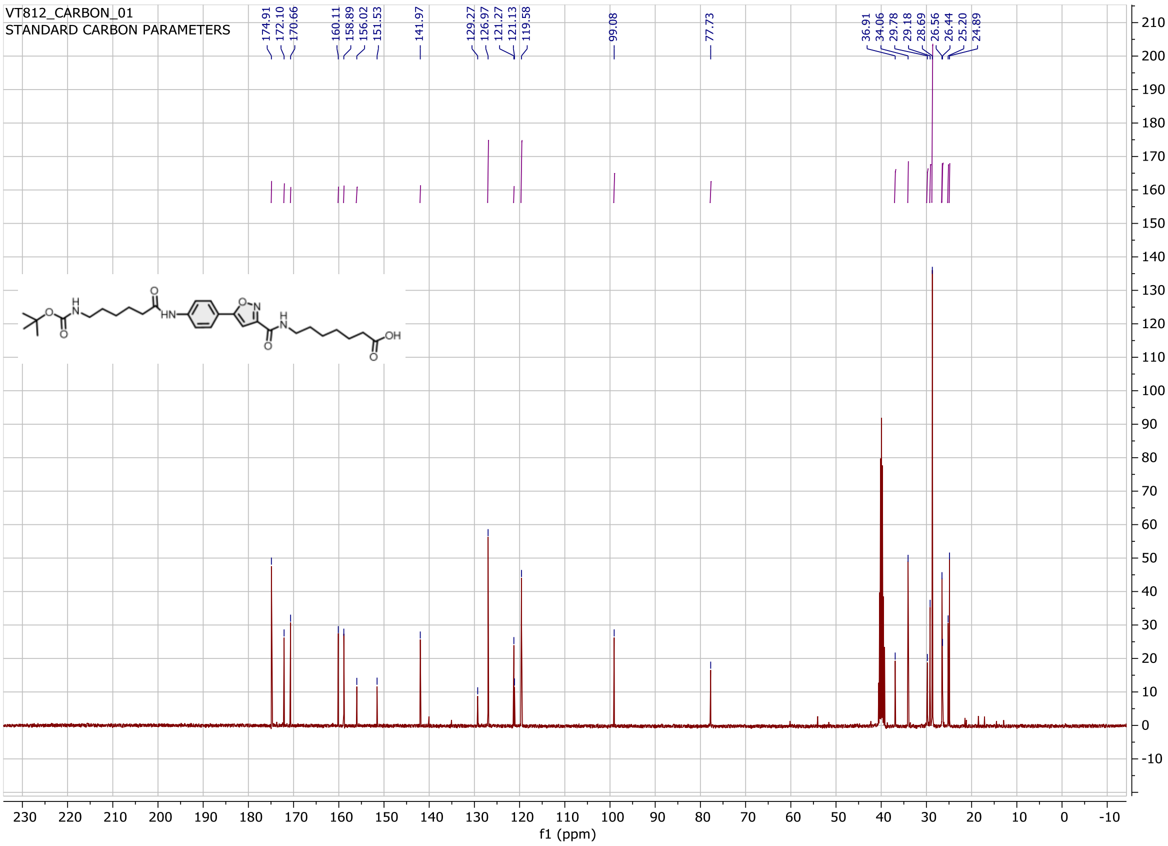

Probe **1**:

$^{1}$H NMR spectrum

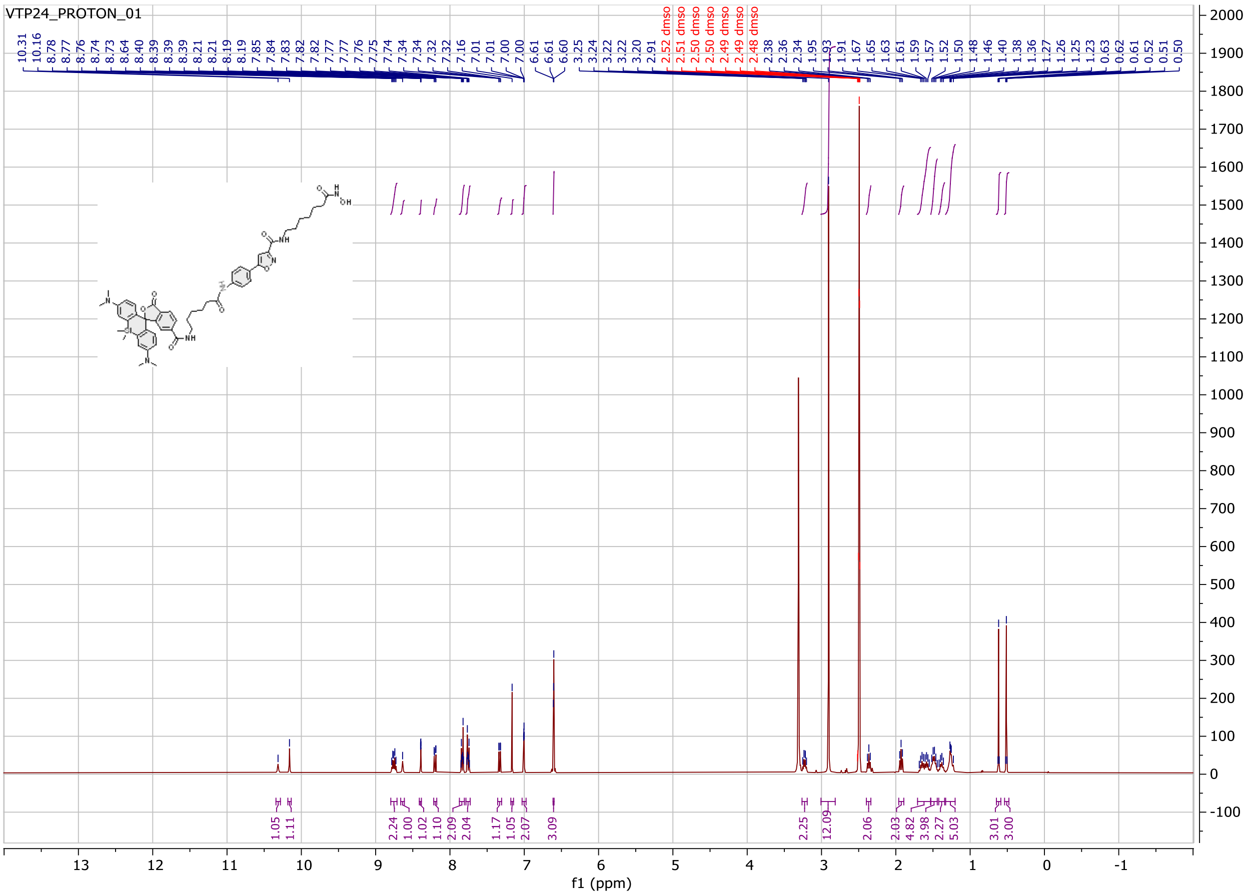

Probe **2**:

$^{1}$H NMR spectrum

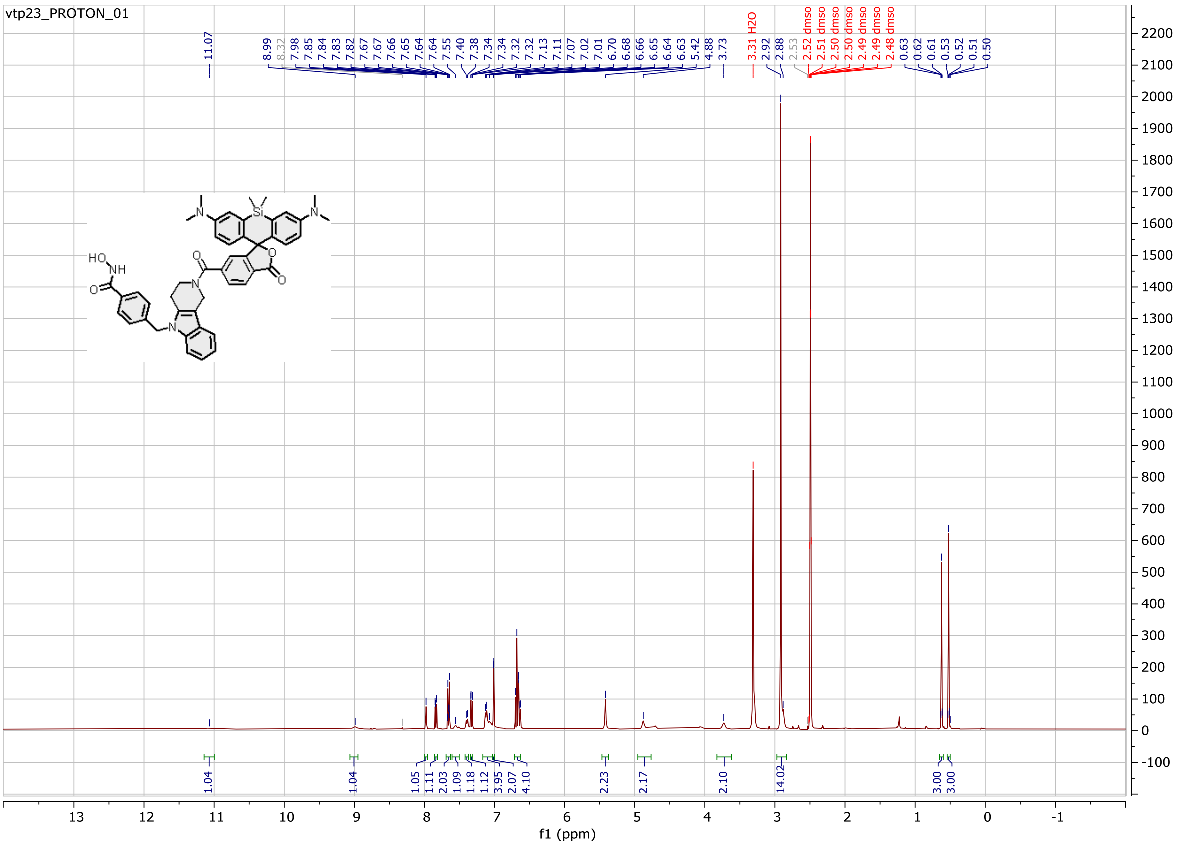

Probe **3**:

$^{1}$H NMR spectrum

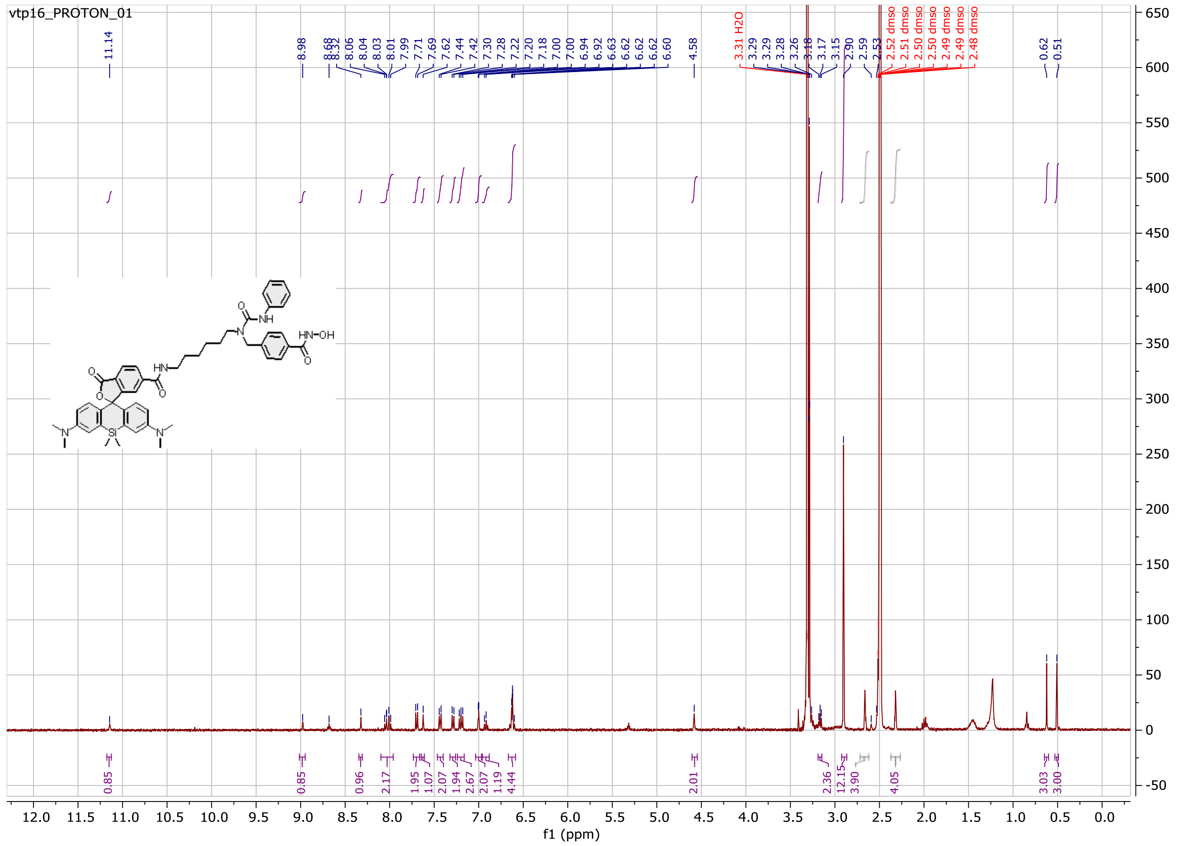

Probe **4**:

$^{1}$H NMR spectrum

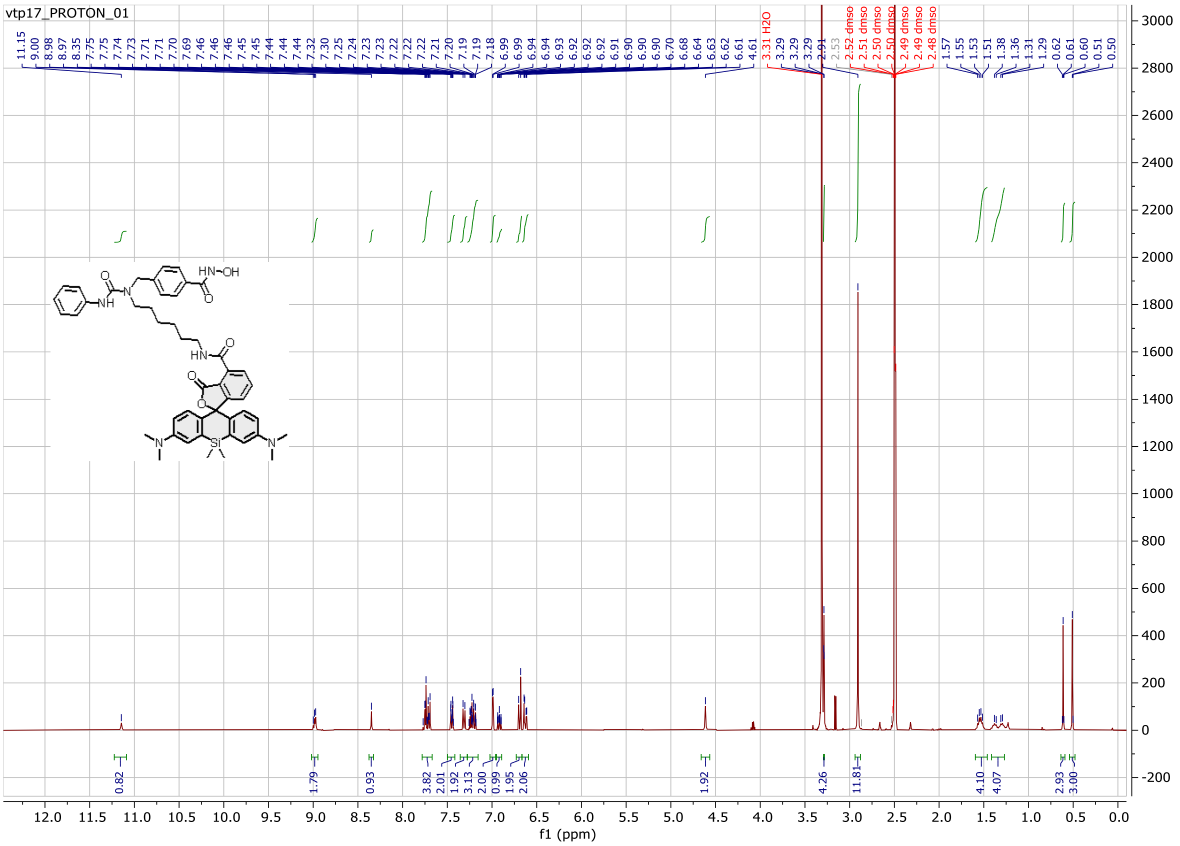

Probe **5**:

$^{1}$H NMR spectrum

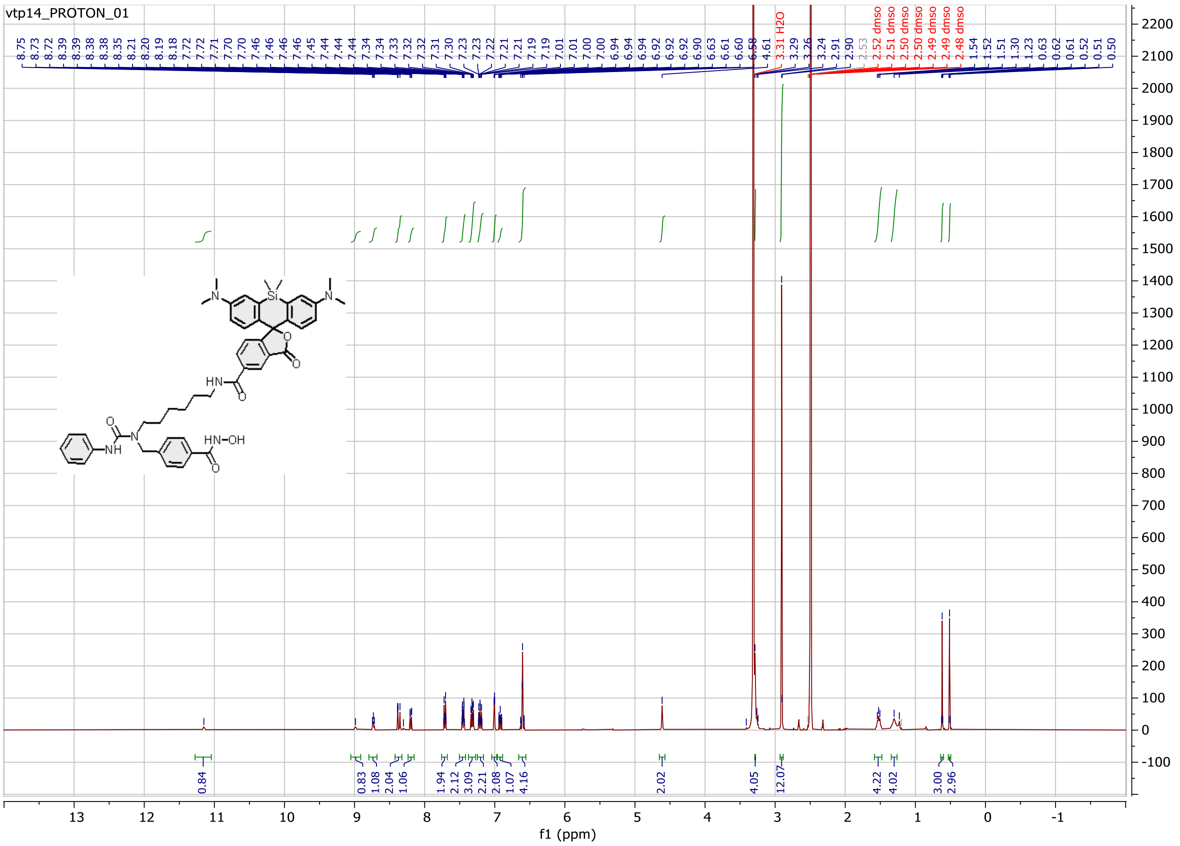

Probe **6**:

$^{1}$H NMR spectrum

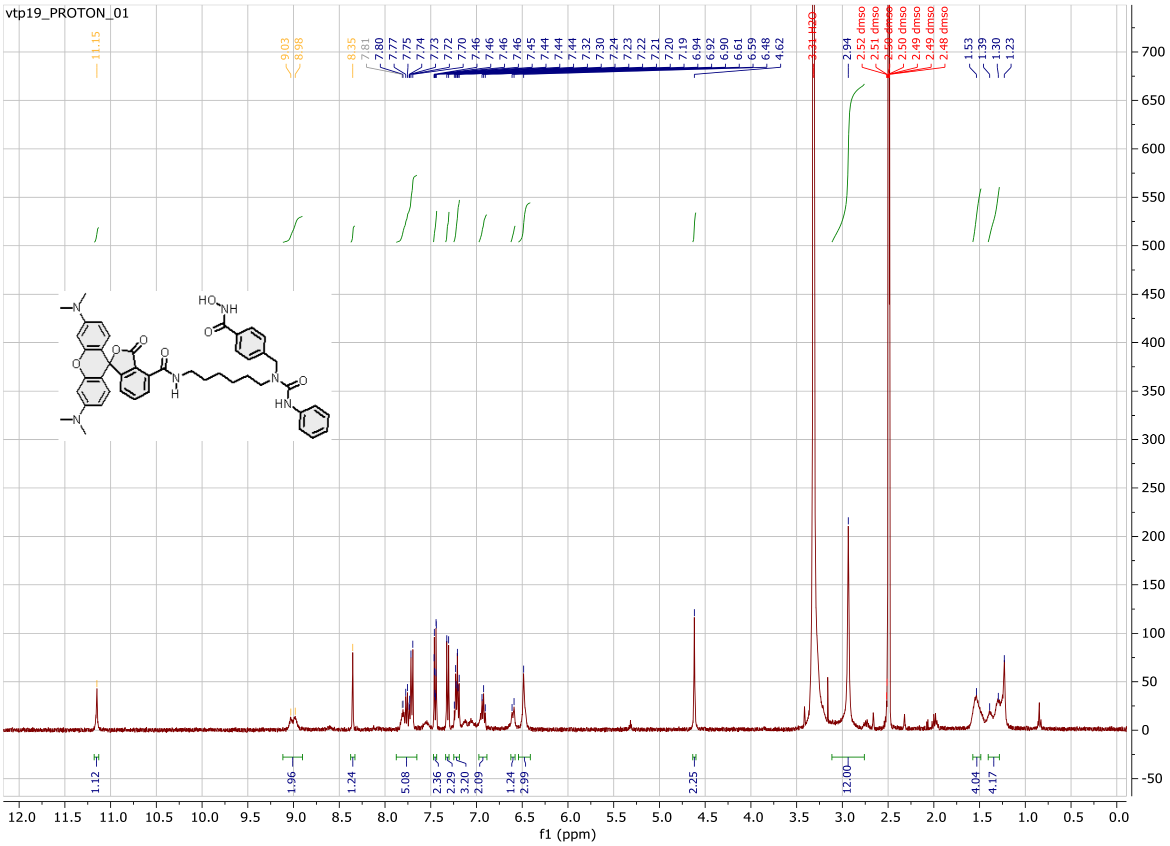

Probe **7**:

$^{1}$H NMR spectrum

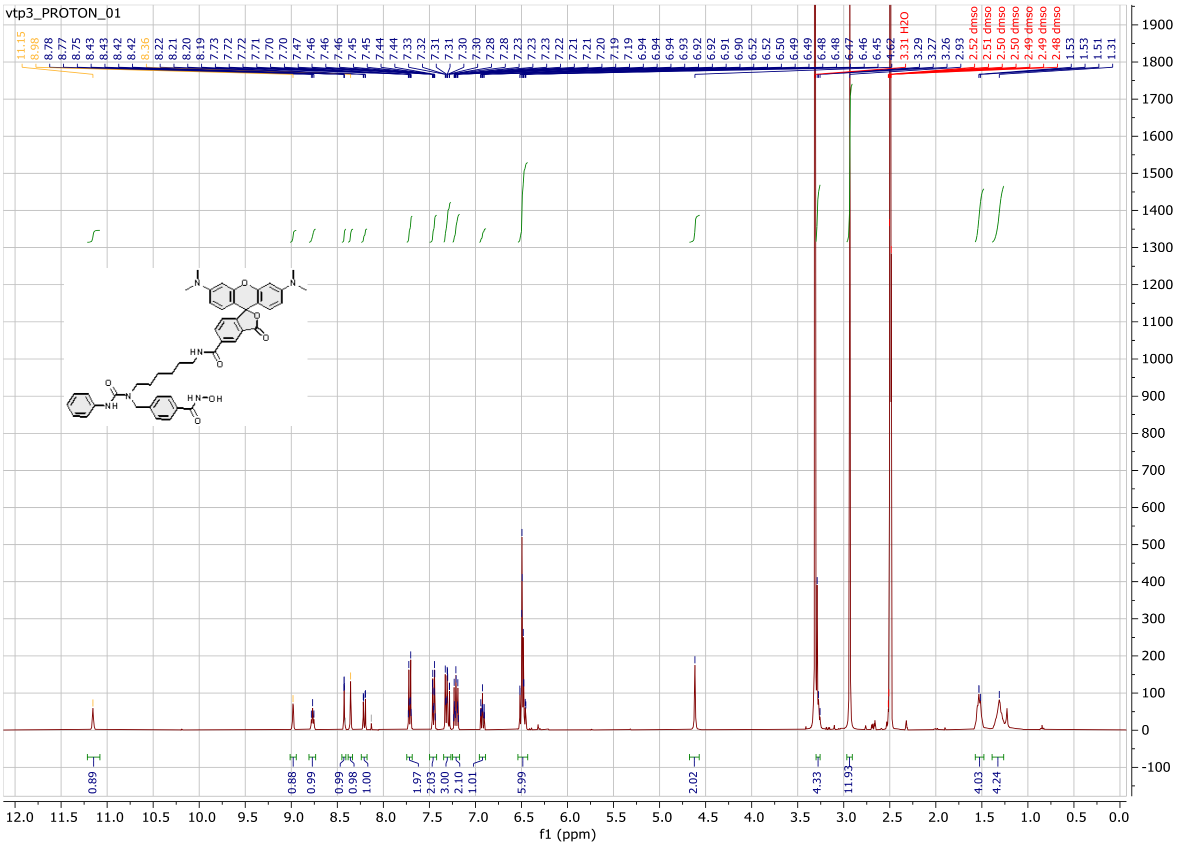

Probe **8**:

$^1$H NMR spectrum

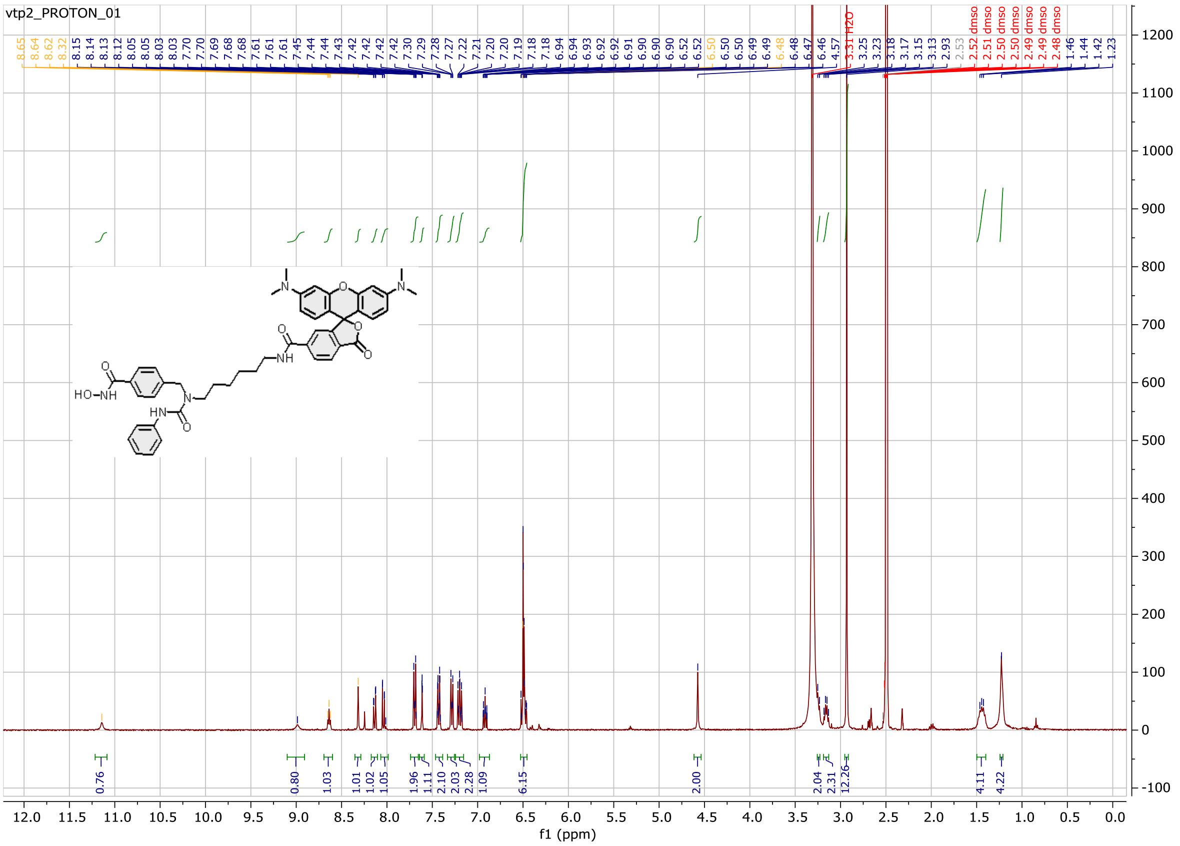

Probe **9**:

[1]H NMR spectrum

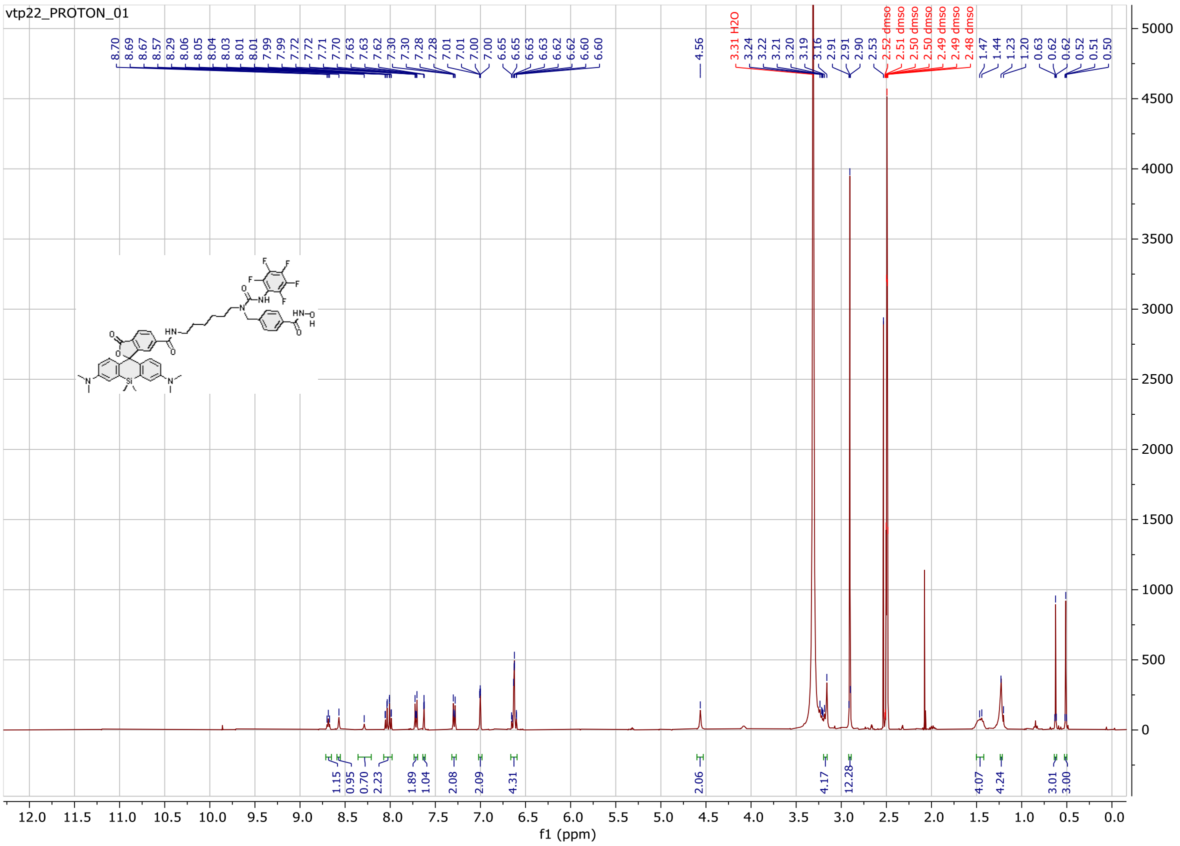

Probe **10**:

$^{1}H$ NMR spectrum

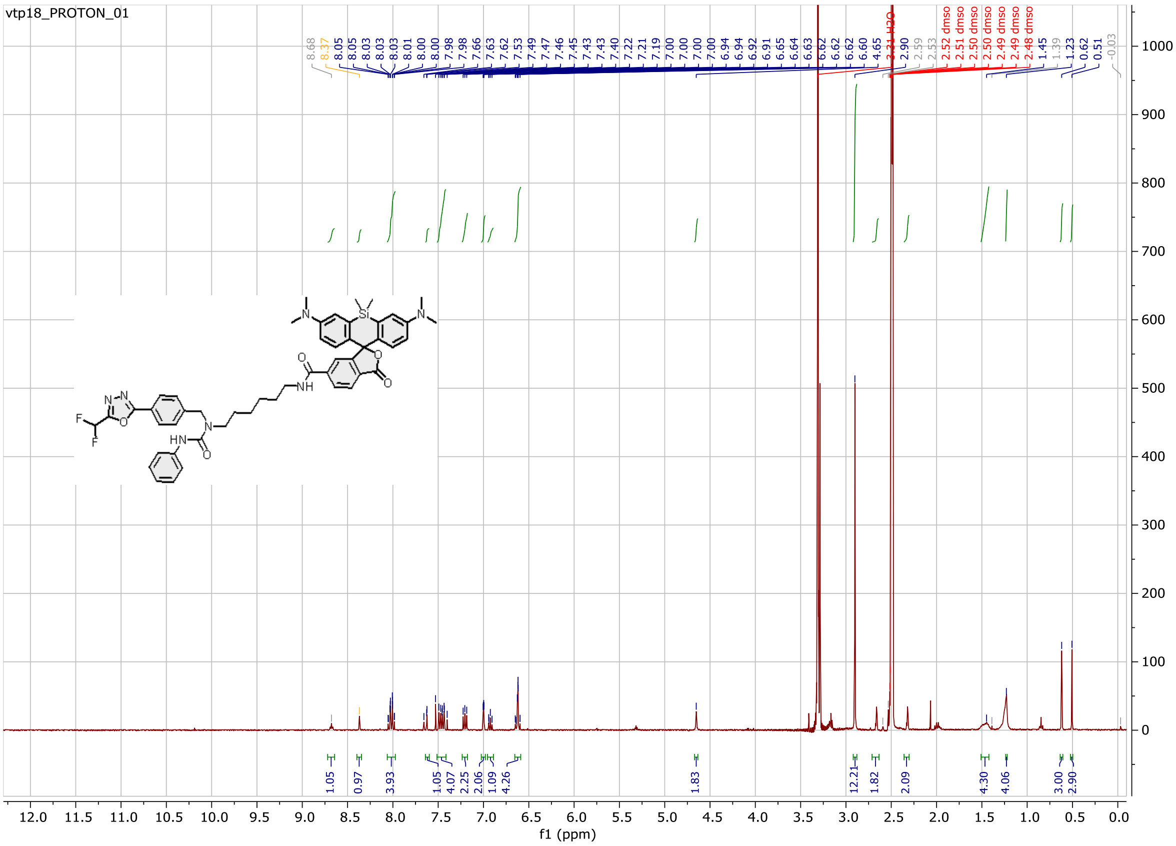

Probe **11**:

[1]H NMR spectrum

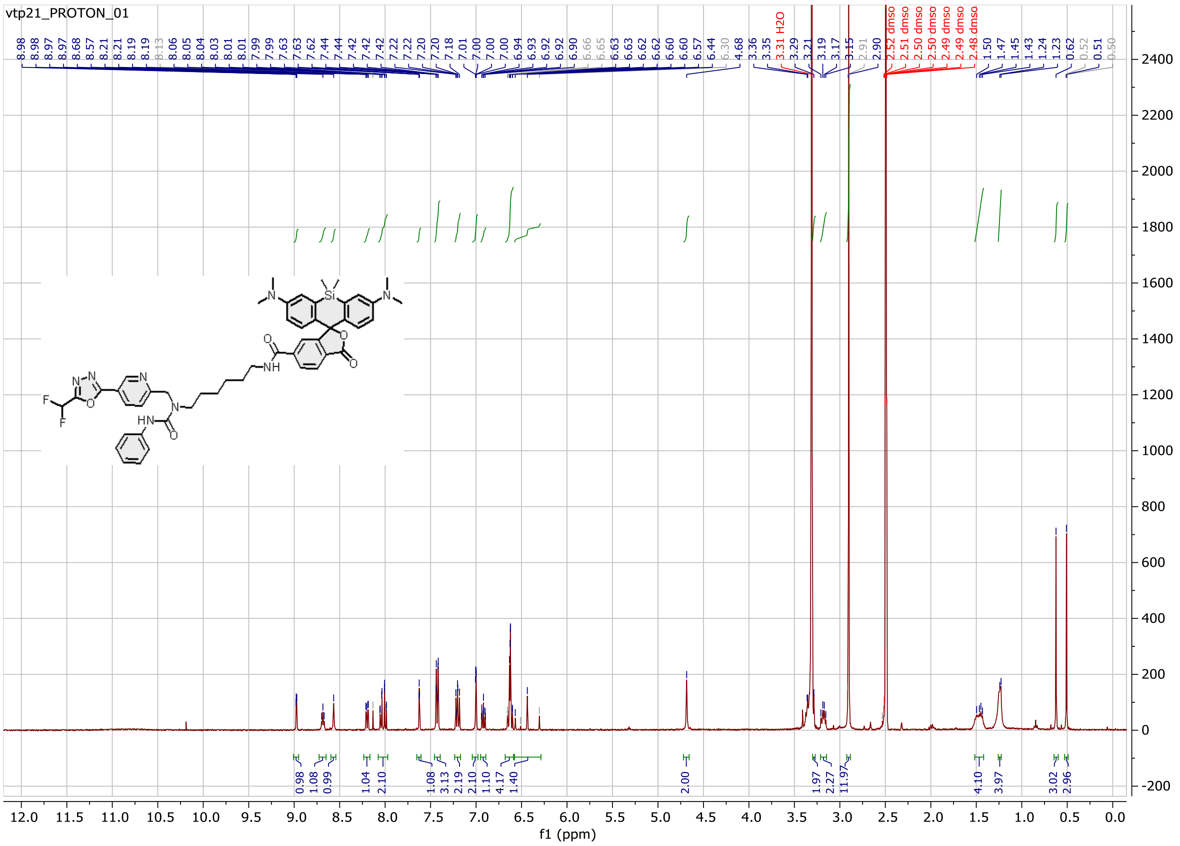

Probe **12**:

$^{1}$H NMR spectrum

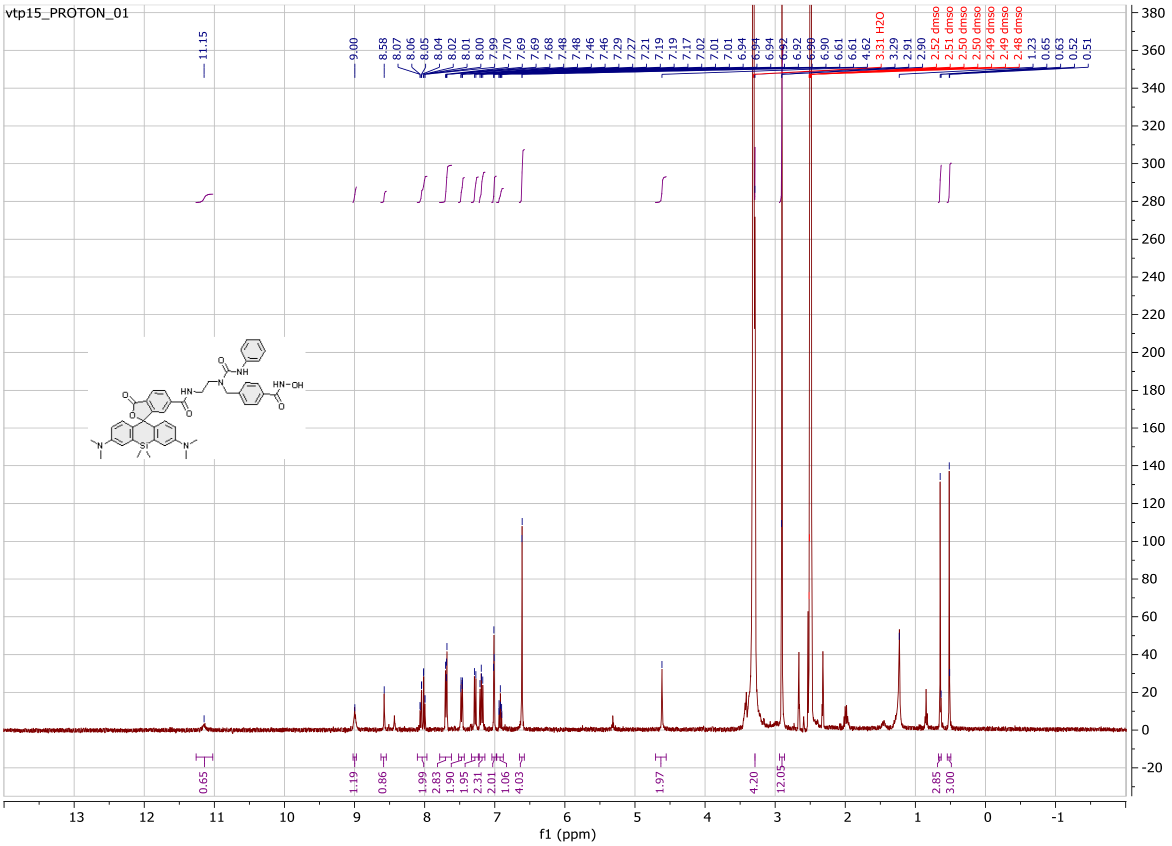

Probe **13**:

$^{1}$H NMR spectrum

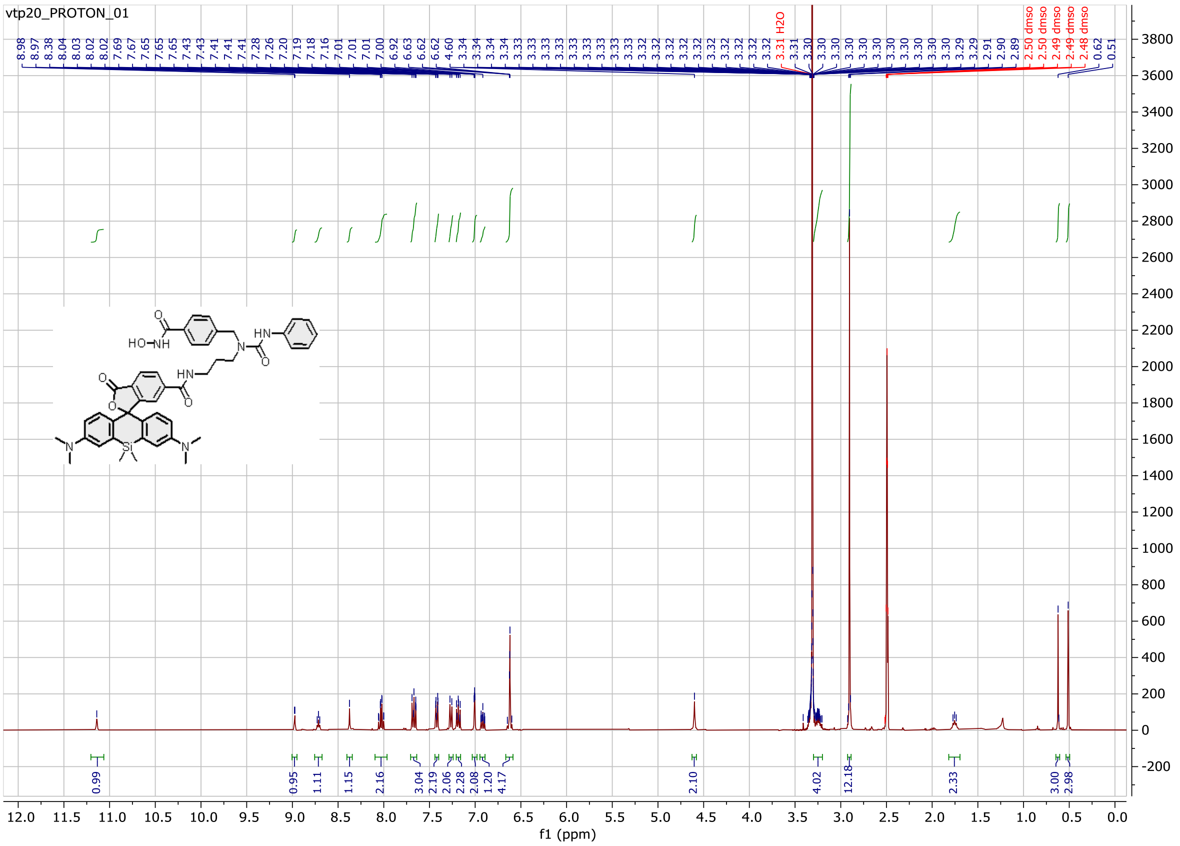

Probe **14**:

[1]H NMR spectrum

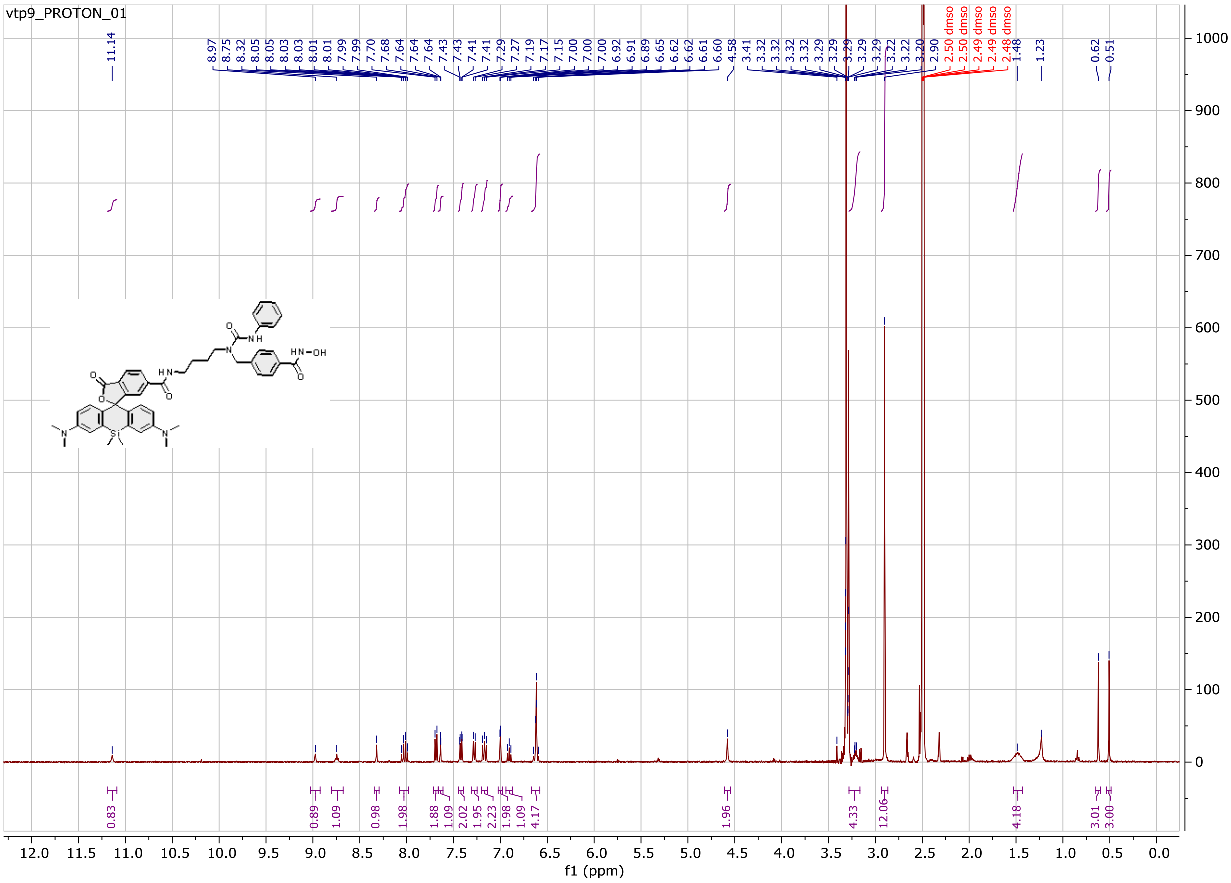

Probe **15**:

$^{1}H$ NMR spectrum

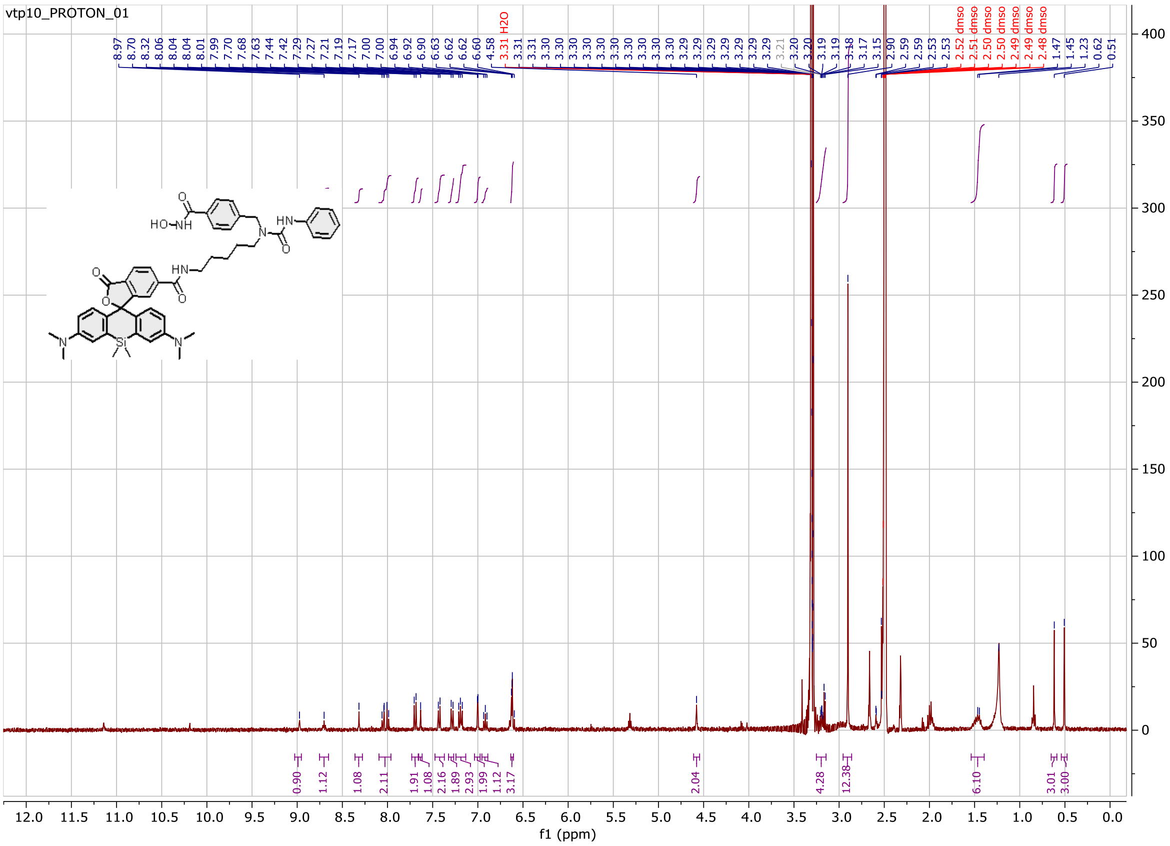

Probe **16**:

[1]H NMR spectrum

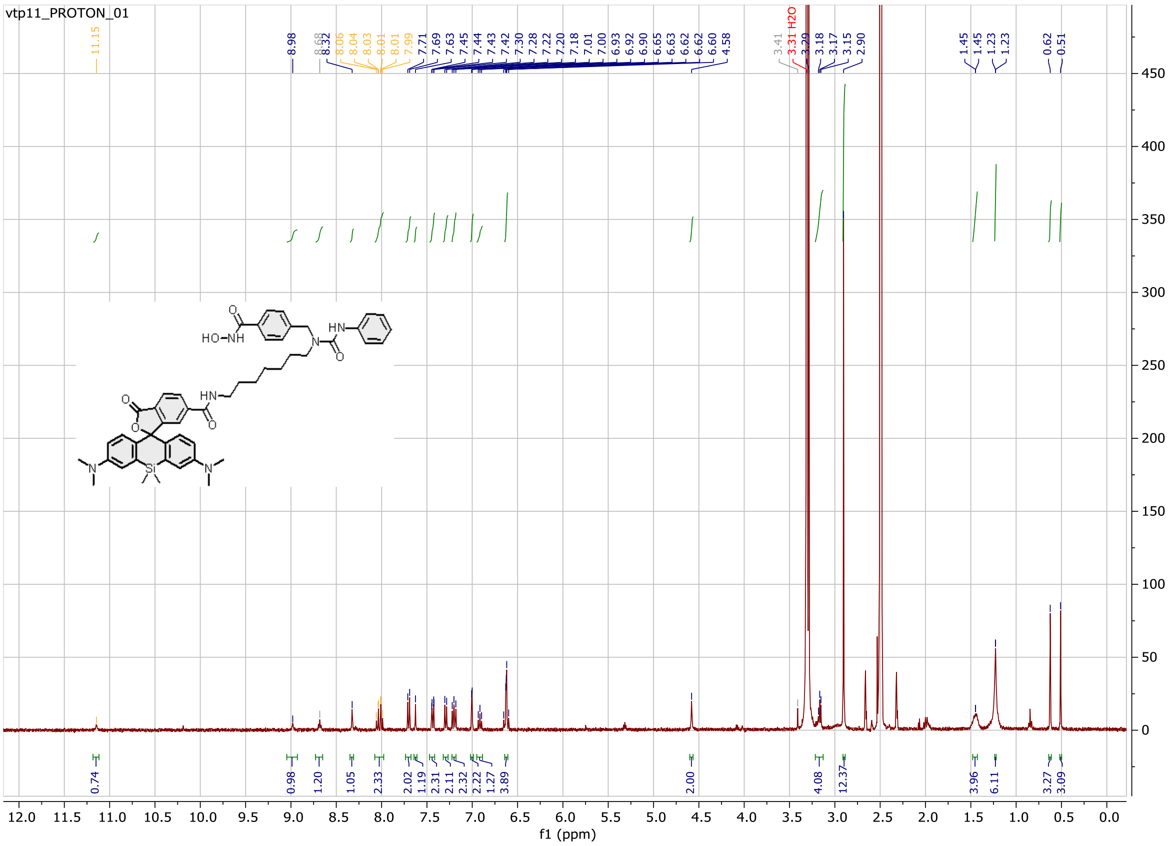

Probe **17**:

$^{1}$H NMR spectrum

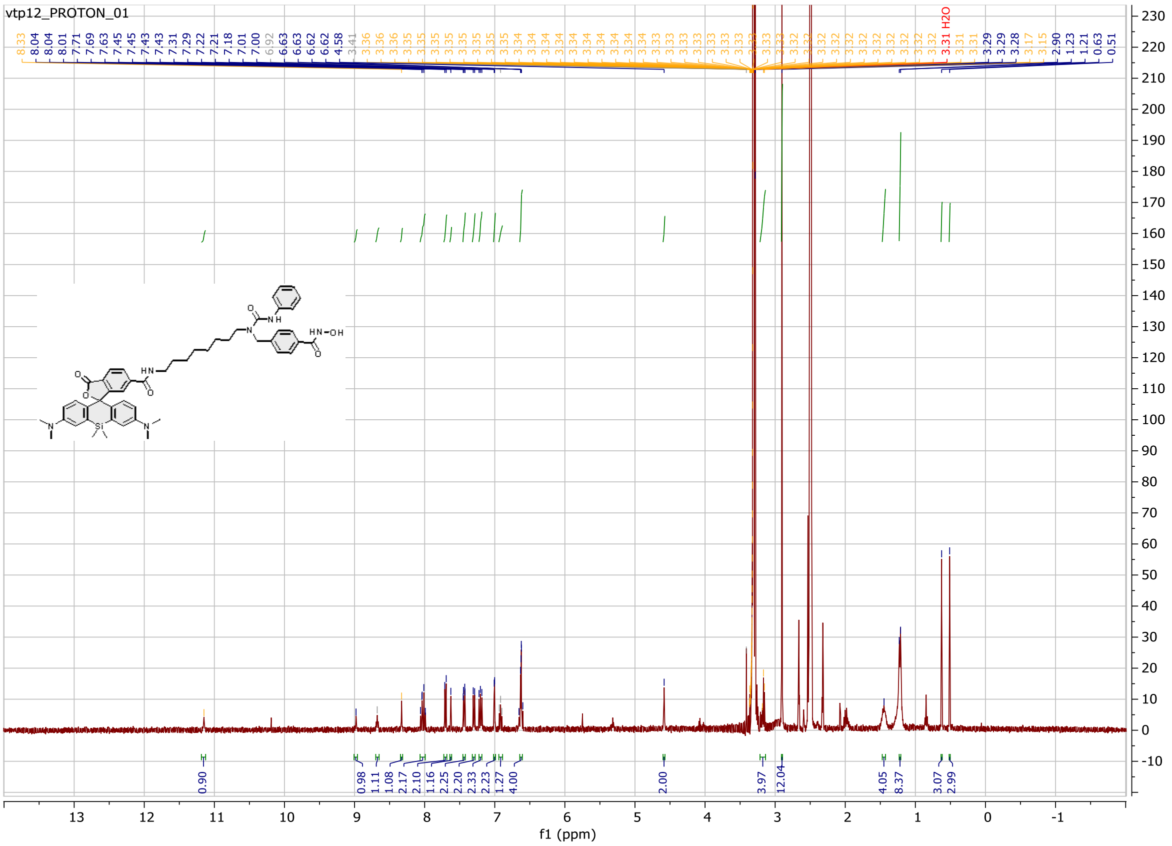

Probe **18**:

$^{1}$H NMR spectrum

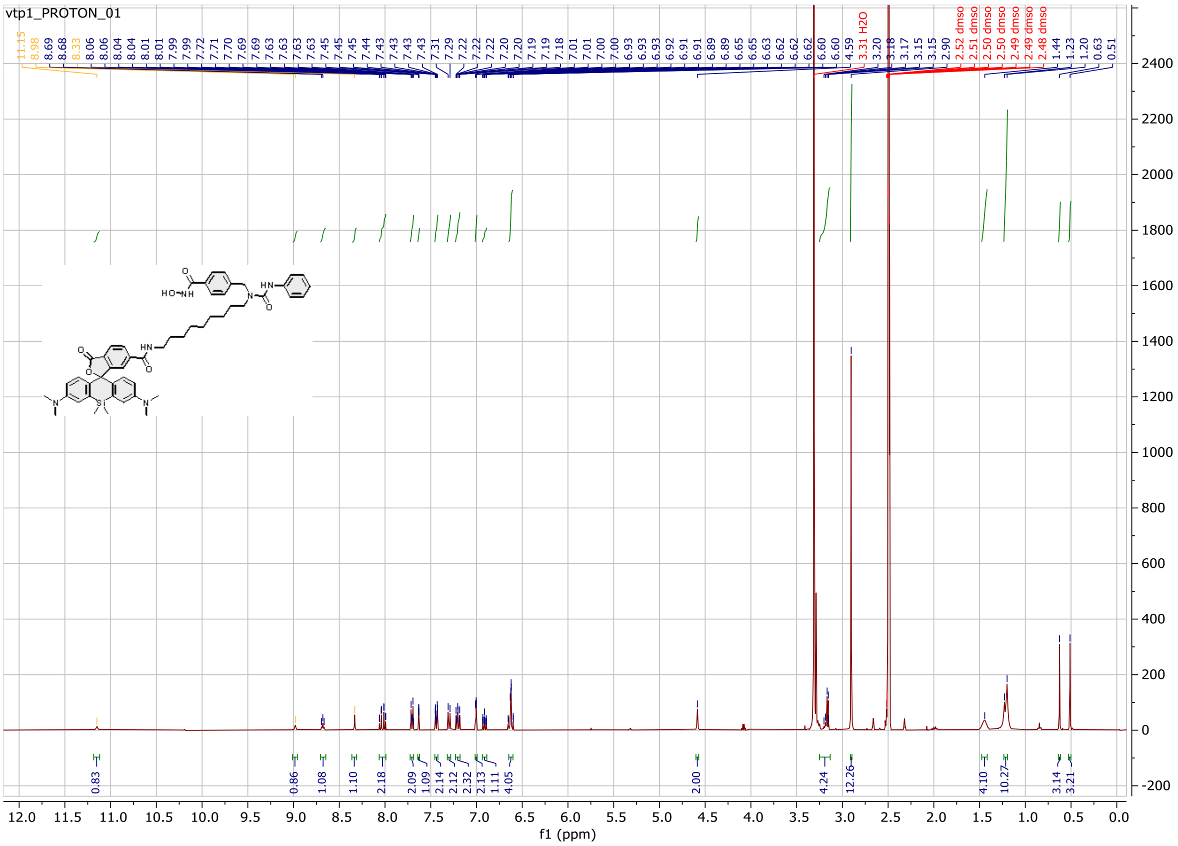

Probe **19**:

$^{1}$H NMR spectrum

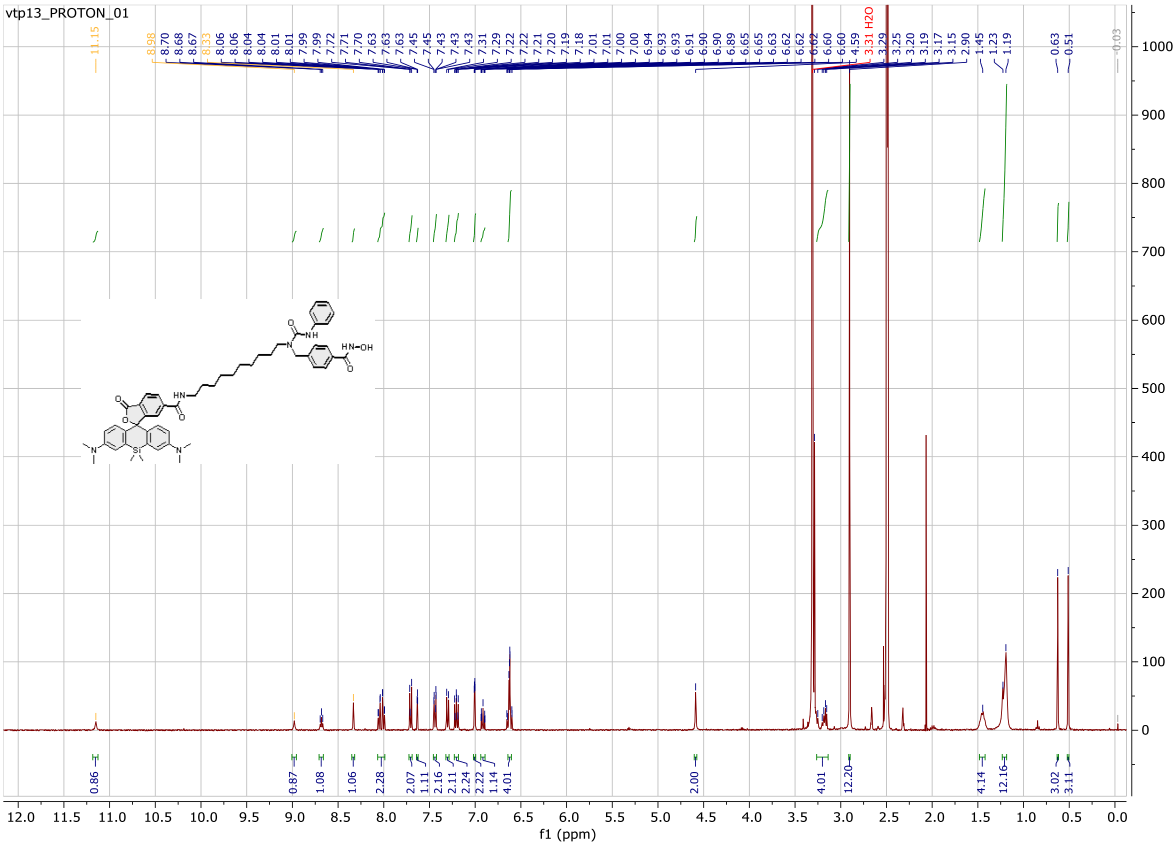

Probe **20**:

$^{1}$H NMR spectrum

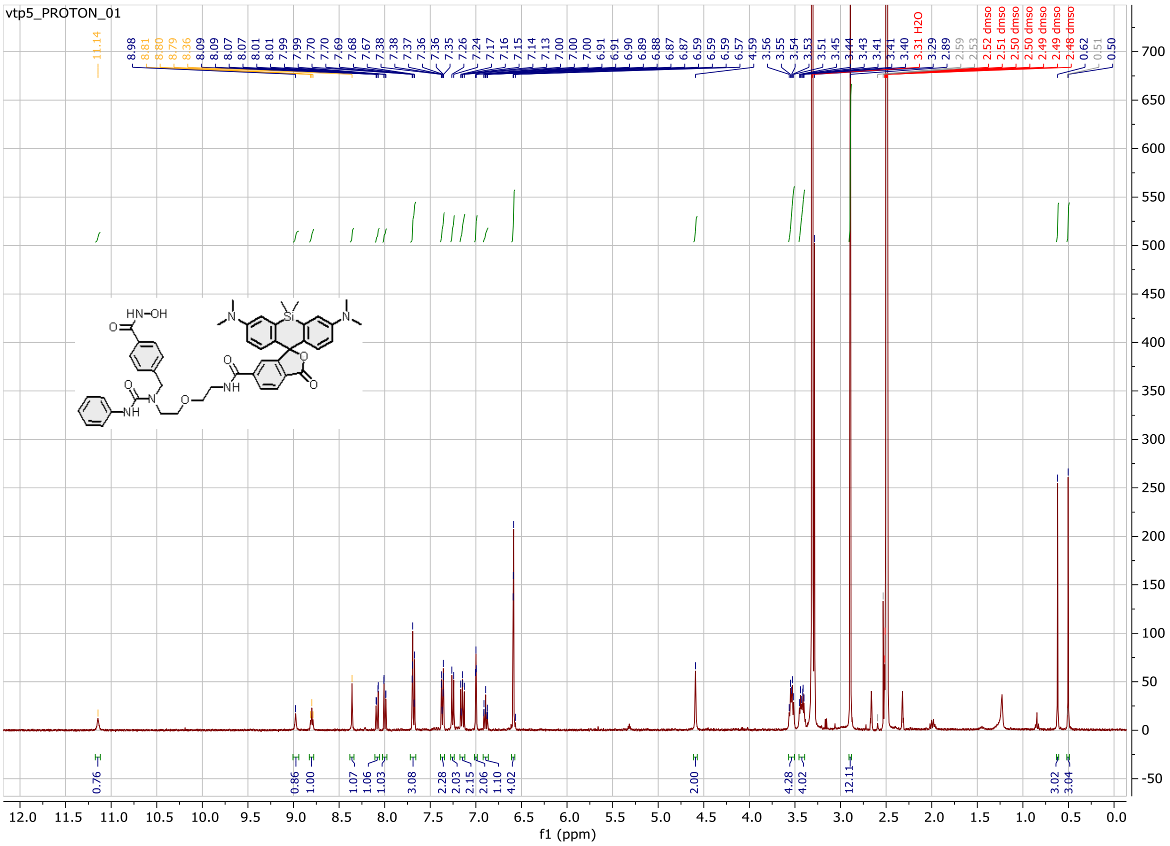

Probe **21**:

[1]H NMR spectrum

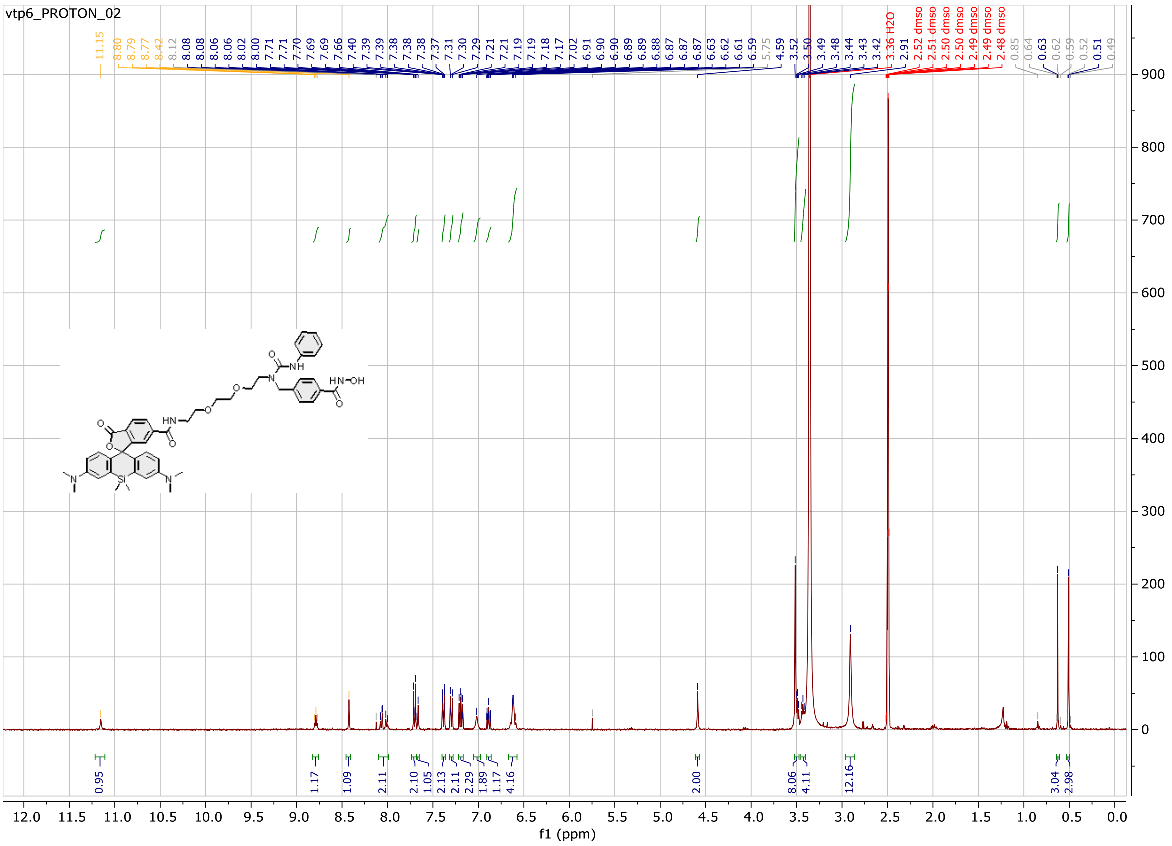

Probe **22**:

$^{1}$H NMR spectrum

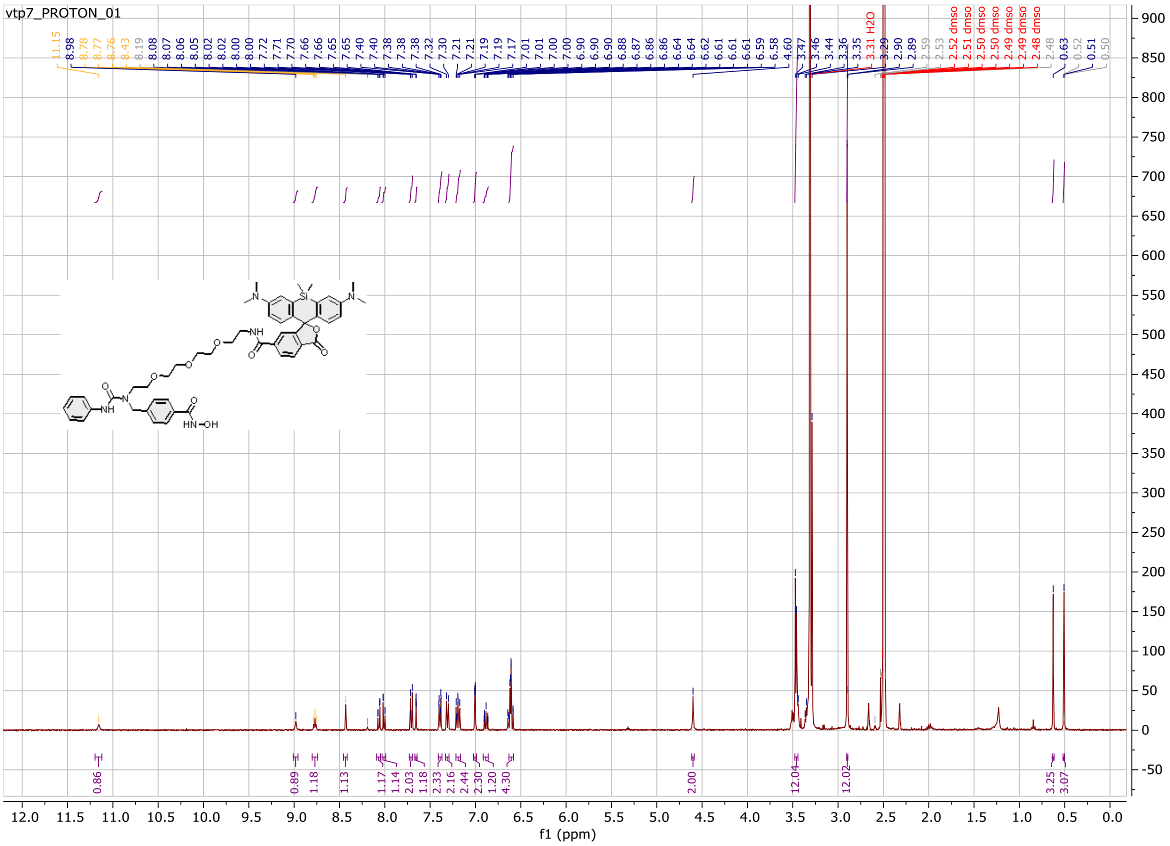

Probe **23**:

$^{1}$H NMR spectrum

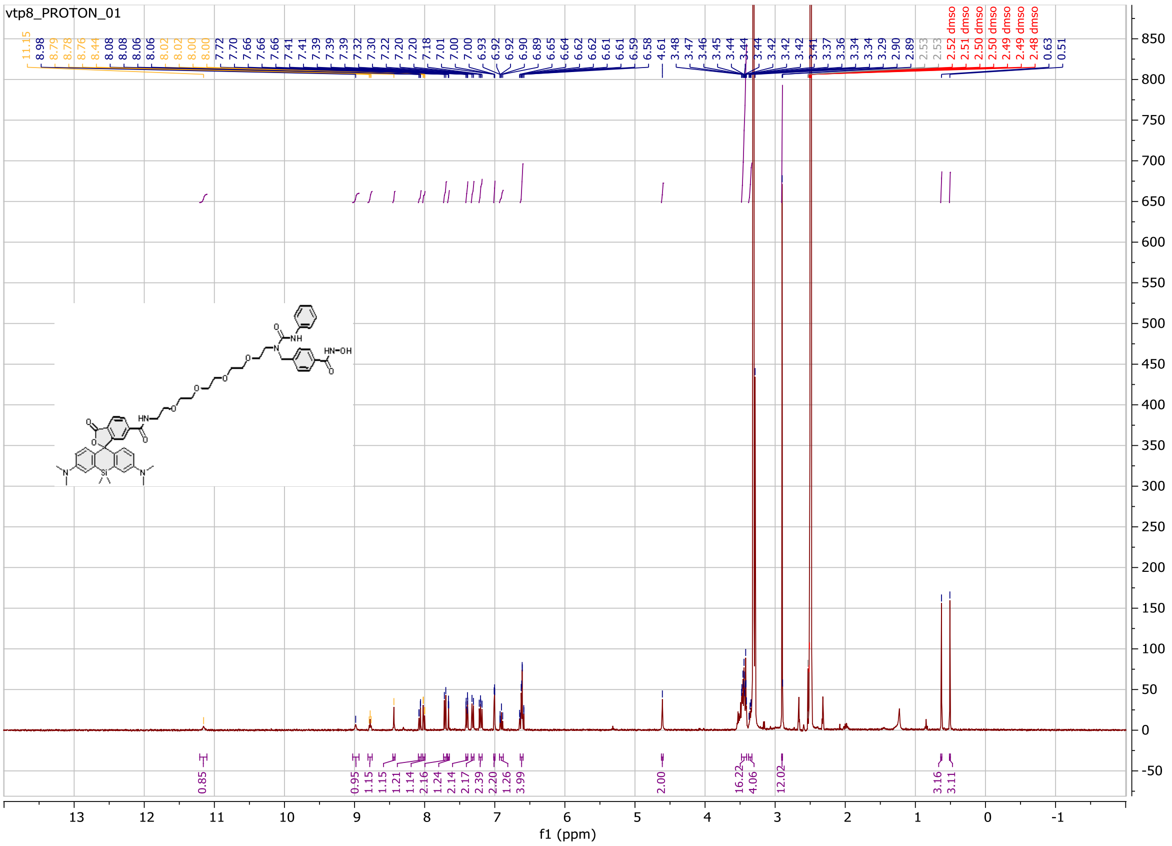
